# Exploring the evolutionary history of kinetic stability in the alpha-lytic protease family

**DOI:** 10.1101/2020.08.30.274340

**Authors:** Charlotte F. Nixon, Shion A. Lim, Zachary R. Sailer, Ivan N. Zheludev, Christine L. Gee, Brian A. Kelch, Michael J. Harms, Susan Marqusee

**Author notes:** SAL: Department of Pharmaceutical Chemistry, University of California, San Francisco, San Francisco, CA 94143. ZRS: Project Jupyter, Department of Physics, California Polytechnic State University, San Luis Obispo, 1 Grand Avenue, San Luis Obispo, CA 93407. INZ: Department of Biochemistry, Stanford University, Stanford, Stanford, CA, 94305.

## Abstract

In addition to encoding the final tertiary fold and stability, the primary sequence of a protein encodes the folding trajectory and kinetic barriers that determines the speed of folding. How these kinetic barriers are encoded by the sequence is not well understood. Here, we use evolutionary sequence variation in the alpha-lytic protease (αLP) protein family to probe the relationship between sequence and energy landscape. αLP has an unusual energy landscape: the native state of αLP is not the most thermodynamically favored conformation and, instead, it remains folded due to a large kinetic barrier preventing unfolding. In order to fold, αLP utilizes an N-terminal pro region of similar size to the protease itself that functions as a folding catalyst. Once folded, the pro region is removed, and the native state does not unfold on a biologically relevant timescale. Without the pro region, αLP folds on the order of millennia. A phylogenetic search uncovers αLP homologs with a wide range of pro-region sizes, including some with no pro region at all. In the resulting phylogenetic tree, these homologs cluster by pro-region size. Homologs naturally lacking pro regions are thermodynamically stable, fold much faster than αLP, yet retain the same fold as αLP. Key amino acids thought to contribute to αLP’s extreme kinetic stability are lost in these homologs, further supporting their role in kinetic stability. This study highlights how the entire energy landscape plays an important role in determining the evolutionary pressures on and changes to the protein sequence.

## Introduction

Proteins must remain functional in a wide array of environments, spanning from the protected cytosol of the cell to harsh extracellular environments that are often laden with proteases and other components that can rapidly degrade and inactivate a protein. For most proteins, achieving and maintaining the native three-dimensional conformation is a requirement for function, and even transient excursions from this folded conformation can allow for rapid proteolysis and destruction, particularly in harsh environments. As such, proteins that reside in extreme environments have employed mechanisms to prolong their folded state and their function.^1,2^ One such strategy is to encode a large unfolding barrier, resulting in less frequent sampling of unfolded or partially folded states. The resulting kinetic stability can help a protein stay folded and functional in extreme pH, temperature, and protease-rich environments, independent of thermodynamic stability.^3^ A high barrier to unfolding comes at a cost. It consequently increases the folding barrier, creating a paradox: how can a protein balance the need to fold at a biologically reasonable rate with the need to avoid unfolding upon reaching the folded state?

Members of the alpha-lytic protease family have evolved an elegant solution to this problem. These proteins encode their own intramolecular chaperone - a large pro region that accelerates folding by thermodynamically stabilizing both the transition state and the folded-state complex.^4–7^ Once folded, however, the pro region is proteolytically degraded, trapping the protein in its native, kinetically stable state.^8^ The homolog from *Lysobacter enzymogenes* (αLP) is the best characterized example of this family.^9^ αLP is a 198-amino acid serine protease that is secreted with an additional 166-amino acid N-terminal pro region that is degraded after secretion and folding, as described above. This pro region is required for proper folding, and can function both *in cis* or when provided exogenously *in trans*.^10,11^

Interestingly, the folded conformation of αLP is thermodynamically unstable.^4^ That is, the native state is less stable than its unfolded state; it remains folded due to the high kinetic barrier to unfolding, not the thermodynamic drive to the folded state. In the absence of a pro region, folding of αLP proceeds through a molten-globule intermediate that is unable to access its native fold within a biologically reasonable timescale (t_1/2_ folding >3,600 years at 25°C).^4^ Thus, as mentioned above, the pro region acts as an intramolecular chaperone and stabilizes the transition state and native state, enabling this thermodynamically unstable protein to reach its native fold. Once the pro region is degraded, the protease is kinetically trapped in its rigid folded state (t_1/2_ unfolding ∼70 days at 25°C) and does not even sample partially unfolded, protease-susceptible conformations.^12^ This unusual folding trajectory presumably provides a fitness advantage to the bacterial host by outlasting other proteases in the harsh soil environment, degrading proteins and providing nutrients for the host.

The homolog *Streptomyces griseus* Protease B (SGPB) has a smaller (76-amino acid) pro region and is less kinetically stable than αLP (t_1/2_ unfolding of 11 days, 35-fold faster than αLP)^5^, suggesting a potential evolutionary relationship between the size and structure of the pro region and the kinetic barrier. The sequence of the SGPB 76-residue pro region is homologous to the C-terminal domain of the αLP pro region. How the function of the pro region is encoded in these different domains and the role of sequence variation (both in the pro region and the protease) in determining the height of the kinetic barrier are unknown.

Here, we interrogate the evolution of kinetic stability in the αLP family of proteases and experimentally characterize the structure and energy landscape of additional homologs in this family. We find that among this αLP family, pro regions cluster into distinct sizes, and that the pro region is lost in a stepwise manner across the evolutionary tree. The energetics and structure of homologs that have no pro region reveal that they are thermodynamically stable and may have evolved novel roles as pseudoproteases. Our findings show how this protease family can encode a wide range of kinetic and thermodynamic properties while maintaining the same three-dimensional fold, and highlights how the kinetics, thermodynamics, and conformation of a protein can be uniquely encoded within a protein family and tuned over evolution.

## Materials & Methods

### Phylogenetics

250 unique sequences were used as BLAST^13^ queries against NCBI’s non-redundant (nr) protein database. Redundant sequences in the output were removed with a 90% sequence identity cutoff from seed sequences using CD-HIT 4.6.6^14^. This yielded a set of 363 protein sequences sampled from across 233 species. MSAProbs 0.9.7 was used to generate an initial alignment, followed by manual refinement in AliView 1.26^15^. The final alignment is available as Supplement 1. Secretion signal sequences were determined using the SignalP 5.0 server.^16^

Phylogenetic trees of the αLP family were generated with and without the pro region as part of the input MSA. In the MSA, residues upstream of position 584 are considered the pro region, excluding the signal sequence, while the residues downstream of position 584 are considered the protease region of each homolog. This cutoff was determined by the αLP sequence and structure. A maximum likelihood phylogenetic tree was constructed using PhyML 3.1^17^ and an AIC test^18^ was used to select a model that maximized the likelihood of the observed alignment while minimizing the number of included parameters. Of the models included in PhyML, the best model proved to be LG+Γ_8_^19^. Branch supports were calculated using the approximate likelihood ratio test^20^.

### Plasmid Cloning

αLP homologs were purchased as gBlock gene fragments from Integrated DNA Technologies. Subsequent restriction enzyme digestion and site-directed mutagenesis were used to clone the genes into a pET-27b(+) expression vector with no secretion signal and a C-terminal GSS-PreScission-GSS-His6 tag. The sequences were confirmed by Sanger sequencing.

### Protein Expression

Rosetta 2(DE3)pLysS Competent Cells (Novagen) were transformed with pET-27b(+) vectors containing αLP homolog DNA and plated on LB agar plates with 50 μg/ml kanamycin. Single colonies were used to inoculate starter cultures for overnight growth to saturation. 15 mL overnight culture was used to inoculate 1 L of LB growth culture with 50 μg/ml kanamycin. Cells were grown at 37°C, 250 RPM to an optical density (600 nm) of 0.6 to 0.8 and induced with 1 mM isopropyl-β-D-thiogalactopyranoside. Cells were harvested after 3 hours by centrifugation at 4000 x g for 15 min at 4°C. Cell pellets were resuspended in 30 mL buffer A (50 mM HEPES, 50 mM NaCl, 5 mM Imidazole, pH 7.5) and stored at −80°C.

### Protein purification

Frozen cell pellets were thawed, lysed by sonication on ice, and centrifuged at 15,000 x g for 30 min at 4°C. Inclusion bodies were visible, and the resulting pellets were resuspended, washed, and centrifuged again in buffer A with 1% triton, followed by buffer A twice. The washed inclusion bodies were resuspended in buffer A with 6 M GdmCl overnight and subsequently filtered through 0.2 μm filters before purification using Ni-NTA resin and elution with buffer B (buffer A, but with 300 mM imidazole) with 6 M GdmCl. Elutions were dialyzed to 50 mM NaOAc, 50 mM NaCl, 6 M GdmCl, pH 5. Elution protein concentration was measured by UV-Vis spectroscopy and the protein was drop-wise refolded to a final concentration of 0.04 mg/mL into 10 mM KOAc pH 5 while stirring at room temperature. Refolded protein was concentrated in an Amicon stirred cell ultrafiltration system to ∼1 mg/mL. Concentrated protein was dialyzed to 2 L 10 mM KOAc pH 5. Protein mass was confirmed by mass spectrometry.

### Equilibrium denaturation monitored by fluorescence

Protein samples were diluted to a final concentration of 0.04 mg/mL in 10 mM KOAc, pH 5 and a range of [GdmCl]. Using a FluoroMax-3 spectrofluorometer, samples were excited at 280 nm and emission spectra were collected from 305 to 450 nm, at 25°C. The ratio of emission at 374/324 nm was calculated and normalized using the equation: (y-y_D_)/(y_N_-y_D_), where y_N_ is the ratio at 0 M GdmCl and y_D_ is the ratio at the highest [GdmCl].

### Protease activity assay

Protease activity was measured by using an MBP-barstar variant containing a flexible linker with the sequence ENLYFQGPPPY/GS(25)/LEVLFQGPG as a substrate. 6.5 μM protein was added to 25 μg substrate in 50 mM KOAc, 10 mM CaCl_2_, 0.5 mM TCEP, pH 5 at 25°C overnight. Samples were run on SDS/PAGE to determine barstar cleavage. Thermolysin was used as a positive control.

### Folding kinetics monitored by fluorescence

For measuring unfolding kinetics, protein samples at 1.5 mg/mL in 0 M GdmCl 10 mM KOAc pH 5 were manually diluted 30-fold to a range of denaturant concentrations. For refolding traces, protein samples at 1.5 mg/mL in ∼6 M GdmCl, 10 mM KOAc, pH 5 were equilibrated overnight at 25°C. Samples were manually diluted 30-fold into 0 M GdmCl, 10 mM KOAc, pH 5. Samples were excited at 280 nm and emission was collected at 374 nm, with constant stirring at 25°C.

### X-ray crystallography

Protein crystals of N4 were grown using the sitting drop vapor diffusion method with drops containing 100 nL of protein solution (18 mg/mL N4, 10 mM KOAc, 50 mM KCl, 0.5 mM TCEP, pH 5) and 100 nL of well solution (1.5 M Lithium sulfate monohydrate, 0.1 M BIS-TRIS propane, pH 7.0). Drops were equilibrated against 45 μL well solution and incubated at 20°C. Crystals grew after 5 days and were harvested by looping and flash freezing in liquid nitrogen on day 20. The cryoprotectant was well solution with 20% glycerol.

Diffraction data were collected at the Advanced Light Source at Lawrence Berkeley National Laboratory, at beamline 8.2.2 using an ADSC Q315R detector. The images were integrated with XDS^21^ and then scaled and merged with Aimless^22^ in the CCP4 suite^23^. The structure was solved by molecular replacement with Phenix Phaser^24^ using SGPB (PDB 3SGB) as the search model. The model was refined in Coot^25^ and Phenix Refine^24^. All reflections were used in refinement, though completeness is low for 1.25 Å. Figures were rendered using Chimera^26^ and the PyMOL Molecular Graphics System, Version 2.0 Schrödinger, LLC.. The structure was deposited in the Protein Data Bank under PDB ID: 6XHZ.

## Results

### Pro regions of alpha-lytic protease homologs cluster into four distinct sizes, corresponding to structural domain cutoffs

Using NCBI’s non-redundant protein database, we identified 363 homologs of alpha-lytic protease (see Materials & Methods for details on methodology). Most of these homologs (n = 343) are from the gram-positive phylum Actinobacteria (which includes *Streptomyces griseus*, host of SGPB) and eleven are from the gram-negative phylum Proteobacteria (which includes *Lysobacter enzymogenes*, host of αLP). A multiple sequence alignment using all 363 sequences (see Materials & Methods, Supplement 1) reveals that the protease domain is highly conserved and aligns well to the known protease domain of αLP and SGPB (residues 586-869 in the MSA). The mean protease length is relatively constant at 194 ± 25 (standard deviation) amino acids. In contrast, the N-terminal pro regions of these homologs vary significantly by both amino-acid length and sequence. The pro region of each homolog is defined as the sequence between the predicted N-terminal signal sequence and the start of the conserved protease domain.^16^ A histogram of pro-region lengths reveals distinct clusters (Figure 1A). Fitting this histogram to a set of normal distributions identified five peaks, two of which were overlapping and thus combined as one distribution. These clusters are defined as: Large (ProL, 151-177 amino acids), Mid (ProM, 118 to 150 amino acids), Small (ProS, 68 to 85 amino acids), None (No-pro, 0 to 20 amino acids). Mapping the average amino acid length of each cluster to the structure of the αLP pro region shows that ProL corresponds to both N and C pro domains, ProM corresponds to the partial helix and beta strands of N pro domain and full C pro domain, ProS corresponds to C pro domain only, like SGPB, and No-pro corresponds to a loss of the pro region altogether (Figure 1B).

**Figure 1.**
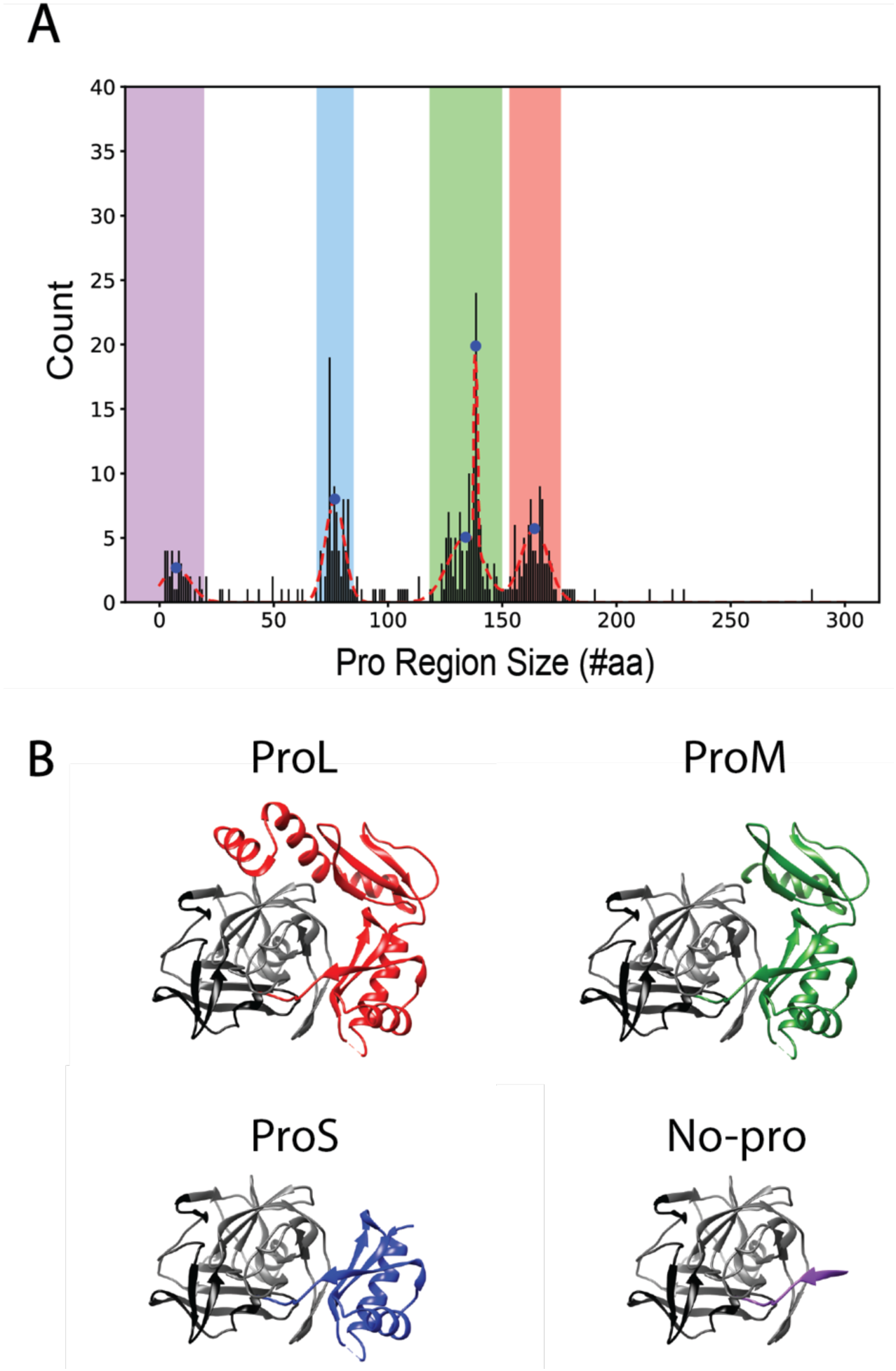
A) Histogram of αLP homolog pro region sizes fit to a set of normal distributions (red dashed line), with the peaks represented as blue dots. Pro region ranges (defined as peak ± two standard deviations) are highlighted as No-pro (purple), ProS (blue), ProM (green), and ProL (red). B) αLP complexed with its pro region (PDB ID: 4PRO) and pro region truncations to represent presumed structures (based on the structure of the pro region of αLP) of ProM, ProS, and No-pro homologs, based on peak pro-region size values. αLP protease N-terminal domain is in black and C-terminal domain is in grey.

The protease region shows high sequence identity^27^ across all the homologs (Table 1), with the average percent identity ranging from 44 to 60% identity within a given pro-region cluster, and 36 to 54% identity between clusters. The C-terminal protease domain, which contacts the pro region through a series of beta-strands^28^, is more conserved than the N-terminal protease domain. The pro regions are more diverse, with only 29 to 39% identity within each size cluster and 21 to 30% identity between clusters. This is true across all structural domains of the pro region.

**Table 1.**
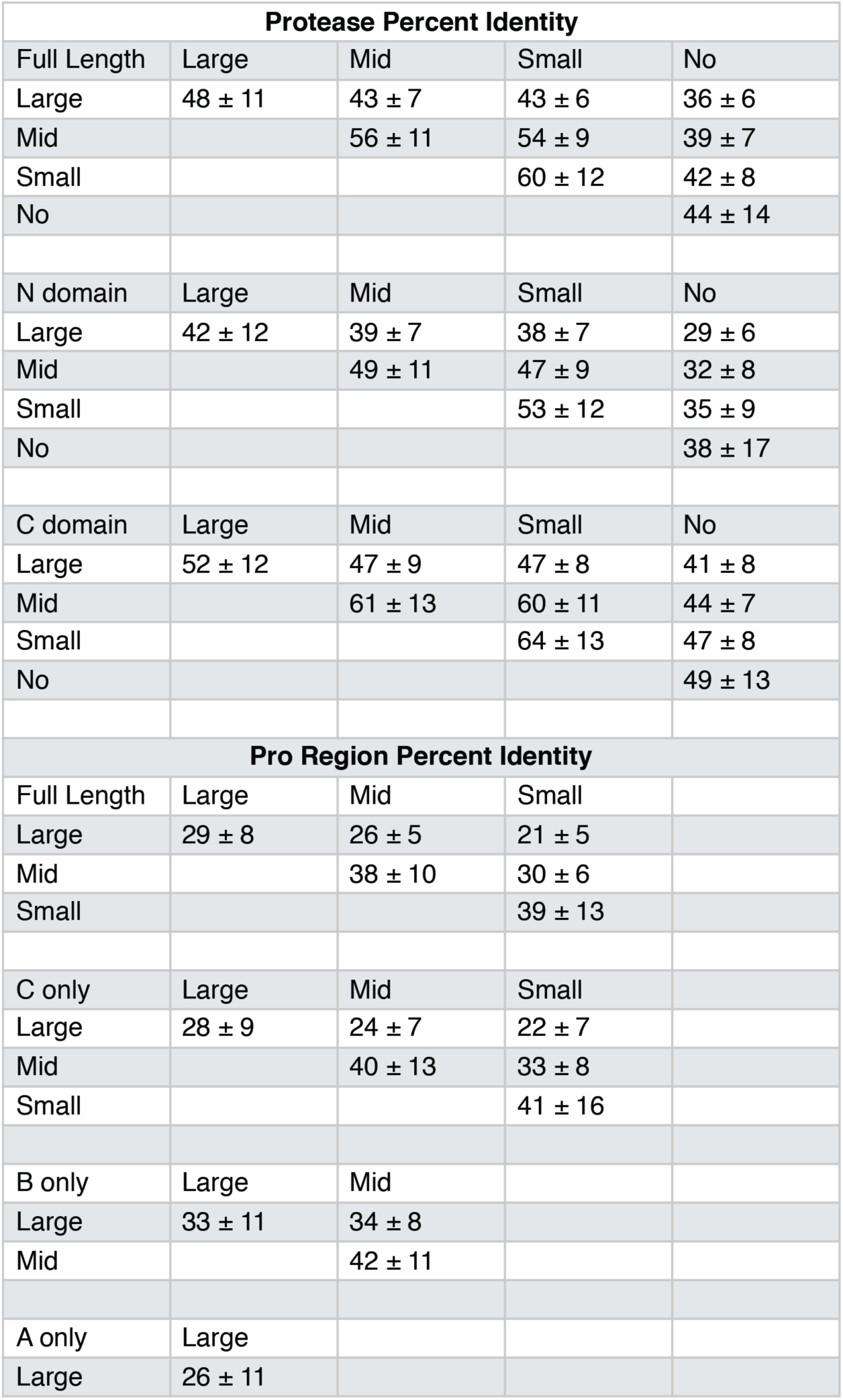
Average percent identities^27^ of pro and protease region for the different pro-region clusters. Analysis was done for both the full sequence and for specific sub-domains of the pro and protease region. (ProS = C domain only, ProM = BC domains, ProL = ABC domains; full length pro region percent identity calculation includes signal sequences).

| Protease Percent Identity |  |  |  |  |
| --- | --- | --- | --- | --- |
| Full Length | Large | Mid | Small | No |
| Large | 48 ± 11 | 43 ± 7 | 43 ± 6 | 36 ± 6 |
| Mid |  | 56 ± 11 | 54 ± 9 | 39 ± 7 |
| Small |  |  | 60 ± 12 | 42 ± 8 |
| No |  |  |  | 44 ± 14 |
| N domain | Large | Mid | Small | No |
| Large | 42 ± 12 | 39 ± 7 | 38 ± 7 | 29 ± 6 |
| Mid |  | 49 ± 11 | 47 ± 9 | 32 ± 8 |
| Small |  |  | 53 ± 12 | 35 ± 9 |
| No |  |  |  | 38 ± 17 |
| C domain | Large | Mid | Small | No |
| Large | 52 ± 12 | 47 ± 9 | 47 ± 8 | 41 ± 8 |
| Mid |  | 61 ± 13 | 60 ± 11 | 44 ± 7 |
| Small |  |  | 64 ± 13 | 47 ± 8 |
| No |  |  |  | 49 ± 13 |
| Pro Region Percent Identity |  |  |  |  |
| Full Length | Large | Mid | Small |  |
| Large | 29 ± 8 | 26 ± 5 | 21 ± 5 |  |
| Mid |  | 38 ± 10 | 30 ± 6 |  |
| Small |  |  | 39 ± 13 |  |
| C only | Large | Mid | Small |  |
| Large | 28 ± 9 | 24 ± 7 | 22 ± 7 |  |
| Mid |  | 40 ± 13 | 33 ± 8 |  |
| Small |  |  | 41 ± 16 |  |
| B only | Large | Mid |  |  |
| Large | 33 ± 11 | 34 ± 8 |  |  |
| Mid |  | 42 ± 11 |  |  |
| A only | Large |  |  |  |
| Large | 26 ± 11 |  |  |  |

### Phylogenetic analysis reveals that pro-region sizes are clustered evolutionarily

To probe the evolutionary relationship between these homologs, we used the multiple sequence alignment to construct a phylogenetic tree (see Materials & Methods, Figure 2A, Supplement 2). The overall topology of the tree remains the same when built with the full-length pro-protease sequences (Supplement 3) or the protease sequences only (Figure 2A). Homologs cluster according to their pro-region size; that is, homologs are closely related to proteases that have similar-sized pro regions. It is especially noteworthy that this clustering appears even when the tree is built without using the pro-region sequence. Therefore, the evolutionary relationships and sequence changes underlying pro-region size appear to be sufficiently encoded within the protease sequence alone.

**Figure 2.**
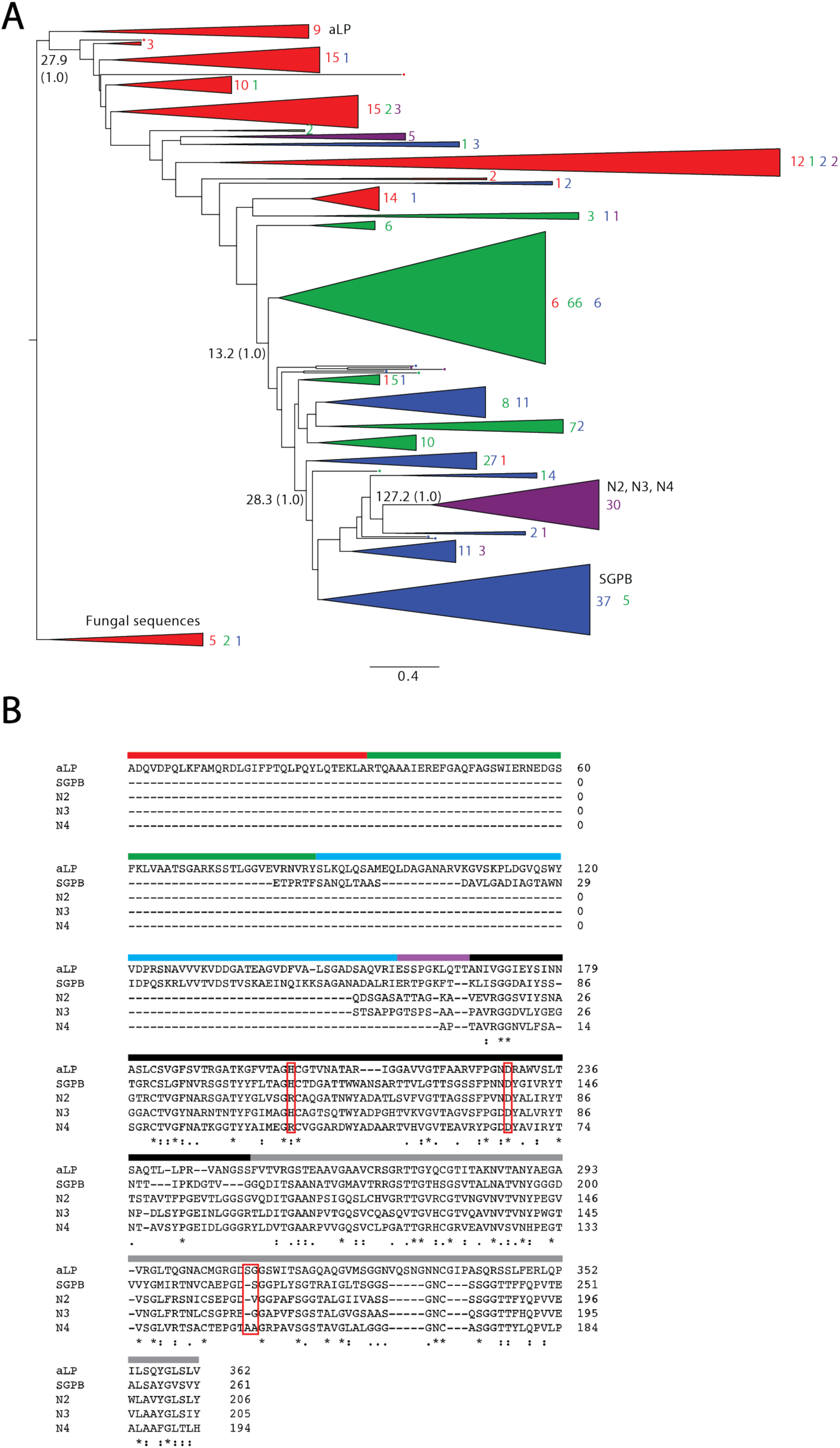
A) αLP phylogenetic tree with clades collapsed and colored based on pro regions found within that clade: No-pro (purple), ProS (blue), ProM (green), and ProL (red). Each clade is annotated with the number of sequences of each pro region cluster/color. The raw log-likelihood ratio and SH support (in parentheses) are listed for nodes where the pro region size changes. B) Alignment of selected homologs. The sequences are colored based on pro region and protease domains. Active site residues (canonical Asp102, His57, and Ser195) are noted in red boxes. (Asterisk) indicates fully conserved residues, (colon) indicates strongly similar residues, and (period) indicated weakly similar residues.^27^

To order these changes in pro-region length over time, we rooted the phylogenetic tree. We did this by three different methods: rooting with 8 fungal out-group sequences (Figure 2A), rooting by mid-point, and rooting by minimal ancestral deviation. These three methods gave slightly different roots; however, in all three methods the large pro-region was assigned as an ancestral feature of this family (Supplement 2). Proteases with a large pro region appear as an older trait in the phylogenetic tree, and the pro region decreases over time, resulting in αLP homologs without pro regions. While most of the homologs without pro regions are in a distinct clade, there are several No-pro homologs that branch off directly from the ProL clade, suggesting that it is possible for a pro region to be lost entirely in a short evolutionary span. In sum, the global trend of the evolutionary tree of the aLP family indicates a stepwise trend in pro region loss.

### Homologs lacking pro regions are cooperatively folded and thermodynamically stable

Since the pro region is a requirement for αLP folding, how do homologs in the No-pro cluster fold without any assistance from the pro region? To investigate this, we chose three No-pro homologs for biophysical analysis (N2, N3, N4). All three of these homologs are in the distinct No-pro clade at the terminal end of the tree and come from the Actinobacteria phylum. Notably, these No-pro homologs lack the canonical serine protease catalytic triad, in spite of their high sequence identity to αLP and SGPB (Table 2), suggesting that they are not proteolytically active (Table 2, Figure 2B).

**Table 2.** Pro and protease region percent identities among homologs studied.

|  |  | %Identity Pro Region |  |  |  |  |
| --- | --- | --- | --- | --- | --- | --- |
| %Identity Protease |  | ALP | SGPB | N2 | N3 | N4 |
|  | ALP |  | 26 | - | - | - |
|  | SGPB | 37 |  | - | - | - |
|  | N2 | 34 | 52 |  | - | - |
|  | N3 | 33 | 47 | 58 |  | - |
|  | N4 | 32 | 45 | 55 | 59 |  |

All three No-pro variants were expressed and purified easily using standard *E. coli* expression vectors (see Materials & Methods), which already suggests they are likely able to fold to a thermodynamically stable state. We monitored the stability (ΔG_unf_) for all three variants by carrying out GdmCl-induced denaturation monitored by intrinsic tryptophan fluorescence (pH 5, 25°C, Figure 3A). All three show a sigmoidal (cooperative) equilibrium unfolding transition. N2 is stable with a clear folded baseline, however the marginal stability of N3 and N4 resulted in a poorly defined folded baseline, which complicated the standard two-state analysis of the data. Therefore, after normalization (see Materials & Methods), N2 was fit using a two-state linear extrapolation model and N3 and N4 were fit globally using the baselines of N2 (Table 3).^29^ All three of these homologs are more thermodynamically stable than SGPB^5^ and αLP^4^, and their stabilities fall within a range typical of thermodynamically stable proteins.^30^ N2 is the most stable (ΔG_unf_ = 6.0 ± 0.2 kcal/mol), while N3 and N4 are notably less stable (ΔG_unf_ = 2.0 ± 0.2 kcal/mol and 2.2 ± 0.1 kcal/mol, respectively). The resulting *m*-values are less than expected for proteins of this size, perhaps indicating a breakdown of two-state behavior.^31^ Nevertheless, it is clear that all three of these No-pro homologs have evolved to be folded and thermodynamically stable independent of a pro region. Restoring the catalytic triad did not restore proteolytic activity using a known αLP substrate for any of these recombinant No-pro homologs (see Materials & Methods; Supplement 4).

**Figure 3.**
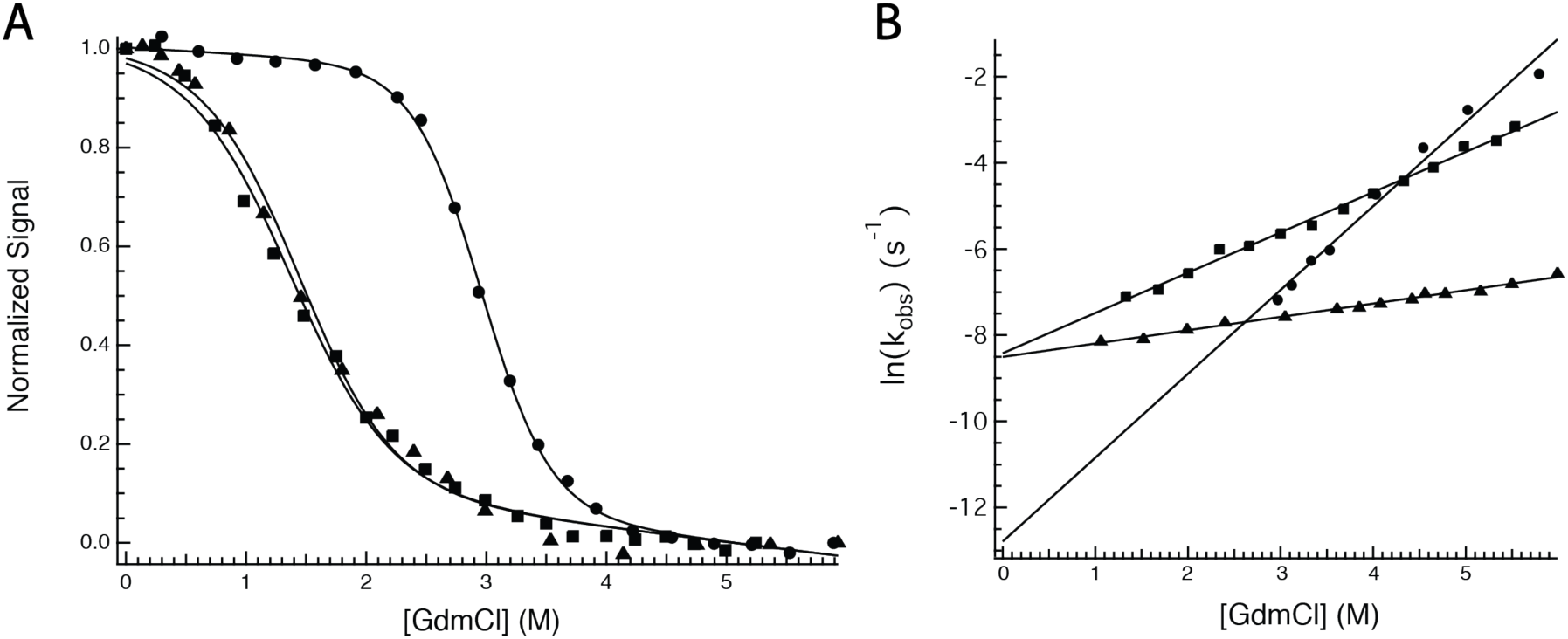
A) Normalized chemical denaturation melts monitored by tryptophan fluorescence of homologs N2 (circles), N3 (squares), and N4 (triangles). Fit lines represent a global fit of replicate melts with linked folded and unfolded baselines. B) ln(*k*_*obs*_) vs denaturant concentration for unfolding kinetic traces of homologs N2 (circles), N3 (squares), and N4 (triangles). Linear fits of the data were used to extrapolate the unfolding rate in 0 M denaturant for each homolog.

**Table 3.** Thermodynamic and kinetic parameters for αLP^4^, SGPB^5^, and homologs of interest (*at 0 M, **0.22 M, and **0.15 M GdmCl; equilibrium errors are reported as one standard deviation of replicates; kinetic errors are reported as one standard deviation of linear or exponential fits).

| | Parameter | αLP | SGPB | N2 | N3 | N4 |
| --- | --- | --- | --- | --- | --- | --- |
| **Equilibrium** | $\Delta G_{unf}$, kcal mol<sup>-1</sup> | - | - | 6.0 ± 0.2 | 2.0 ± 0.2 | 2.2 ± 0.1 |
| | $m$, kcal mol<sup>-1</sup> M<sup>-1</sup> | - | - | 2.0 ± 0.1 | 1.4 ± 0.1 | 1.4 ± 0.1 |
| | $C_m$, M | - | - | 3.0 ± 0.0 | 1.5 ± 0.1 | 1.5 ± 0.1 |
| **Kinetic** | $\Delta G_{unf}$, kcal mol<sup>-1</sup> | -4 ± 0.07 | 0.71 ± 0.04 | - | - | - |
| | $k_{unf}$ (sec<sup>-1</sup>) | 2.1 $\times 10^{-8}$ | 7.4 $\times 10^{-7}$ | 2.8 ± 1.2 $\times 10^{-6}$ | 2.2 ± 0.2 $\times 10^{-4}$ | 2.0 ± 0.1 $\times 10^{-4}$ |
| | $t_{1/2}$ unfolding | 70 days | 10.8 days | 2.9 ± 1.2 days | 52 ± 4 min | 57 ± 2 min |
| | $t_{1/2}$ folding | ~3600 yrs* | 2.97 days* | 77 ± 1 sec** | - | 31 ± 1 sec*** |

### Homologs lacking pro regions unfold on a biologically reasonable timescale

Given the reversible unfolding and thermodynamic stability of the No-pro homologs, we evaluated their unfolding/refolding kinetics monitoring intrinsic tryptophan fluorescence (see Materials & Methods, Supplement 5). Unfolding was initiated by rapid dilution from 0 M GdmCl and kinetics were monitored as a function of final GdmCl concentrations. The time-dependent signal was fit to a single exponential process and the resulting unfolding rates at 0 M GdmCl were then extrapolated from a linear fit of the log of the observed unfolding rate (*k*_*obs*_) versus denaturant concentrations (Figure 3B). This resulted in an extrapolated *k*_*unf*_ (sec^-1^) of 2.8 ± 1.2 × 10^−6^, 2.2 ± 0.2 × 10^−4^, and 2.0 ± 0.1 × 10^−4^ for N2, N3, and N4 respectively, corresponding to t_1/2_ unfolding of 2.9 ± 1.2 days, 52 ± 4 min, and 57 ± 2 min (Table 3). These rates are one to two orders of magnitude greater than SGPB^5^ and one to three orders of magnitude greater than αLP^4^.

Refolding was monitored by rapid dilution from ∼6 M GdmCl to a final concentration of 0.22 M and 0.15 M for N2 and N4, respectfully. The time course of refolding of both N2 and N4 fit well to a single exponential, resulting in a t_1/2_ folding of 77 ± 1 and 31 ± 1 seconds for N2 and N4, respectively (Table 3, Supplement 5). N3 refolds significantly faster, with the whole signal change occurring within the dead time of manual mixing techniques (∼10 seconds) and therefore resulted in no observed kinetics. In summary, all three proteins fold within seconds to minutes and are at least nine orders of magnitude faster than αLP’s refolding without the assistance of the pro region, which can only be measured by an enzymatic assay.^4^ Thus, N2, N3, and N4 can fold independently of a pro-region chaperone and have a much lower kinetic barrier to unfolding.

### N4, a homolog without a pro region, adopts the alpha-lytic protease fold

To compare the conformation of the No-pro homolog to αLP family homologs that require a pro region to fold, we determined the crystal structure of N4 to 1.25 Å resolution (see Materials & Methods, Table 4). The N4 structure reveals the same double beta-barrel fold conserved among the chymotrypsin-like serine proteases. The structure aligns to αLP (PDB ID: 4PRO) with an average Cα RMSD of 1.06 Å (Figure 4A) and SGPB (PDB ID: 3SGB) with an average Cα RMSD of 0.9 Å. There are a few notable differences between the N4 and αLP structures. Three loops in the C-domain that are adjacent to the “active”-site pocket show increased B factors relative to the rest of the N4 structure (Figure 4B). N4 also has less α/β secondary structure compared to αLP (as determined by the DSSP secondary structure assignment server^32,33^). Specifically, N4 has three loops that are longer than the corresponding loops in αLP (Figure 4C). αLP’s tight turns between secondary structure elements have been associated with its increased rigidity and kinetic stability^34^ and thus, the lengthening of these loops may contribute to the decreased kinetic stability of N4. Although we did not obtain a crystal structure for N2 and N3, based on high sequence identity, we expect these No-pro homologs to adopt a similar structure to N4 and the other αLP family homologs.

**Table 4.** N4 crystal and x-ray diffraction data collection and refinement statistics. Statistics for the highest-resolution shell are shown in parentheses.

|  | **N4** |
| --- | --- |
| **Wavelength** | 1 |
| **Data collection resolution range** | 41.70 – 1.25 (1.27 – 1.25) |
| **Space group** | P 1 21 1 |
| **Unit cell** | 39.121 41.695 51.175 90 98.618 90 |
| **Total reflections** | 133157 (957) |
| **Unique reflections** | 39417 (625) |
| **Multiplicity** | 3.4 (1.5) |
| **Completeness (%)** | 86.6 (27.6) |
| **Mean I/sigma(I)** | 14.9 (0.7) |
| **Wilson B-factor** | 11.303 |
| **R-merge** | 0.036 (1.034) |
| **R-meas** | 0.043 (1.462) |
| **R-pim** | 0.023 (1.034) |
| **CC1/2** | 0.999 (0.278) |
| **Refinement resolution range** | 32.18 - 1.249 (1.293 - 1.249) |
| **Reflections used in refinement** | 39381 (1584) |
| **Reflections used for R-free** | 1934 (90) |
| **R-work** | 0.1704 (0.4024) |
| **R-free** | 0.1873 (0.3902) |
| **Number of non-hydrogen atoms** | 1561 |
| **macromolecules** | 1372 |
| **ligands** | 5 |
| **solvent** | 184 |
| **Protein residues** | 192 |
| **RMS(bonds)** | 0.009 |
| **RMS(angles)** | 0.96 |
| **Ramachandran favored (%)** | 97.89 |
| **Ramachandran allowed (%)** | 2.11 |
| **Ramachandran outliers (%)** | 0.00 |
| **Rotamer outliers (%)** | 0.00 |
| **Clashscore** | 1.10 |
| **Average B-factor** | 19.93 |
| **macromolecules** | 18.35 |
| **ligands** | 41.35 |
| **solvent** | 31.14 |

**Figure 4.**
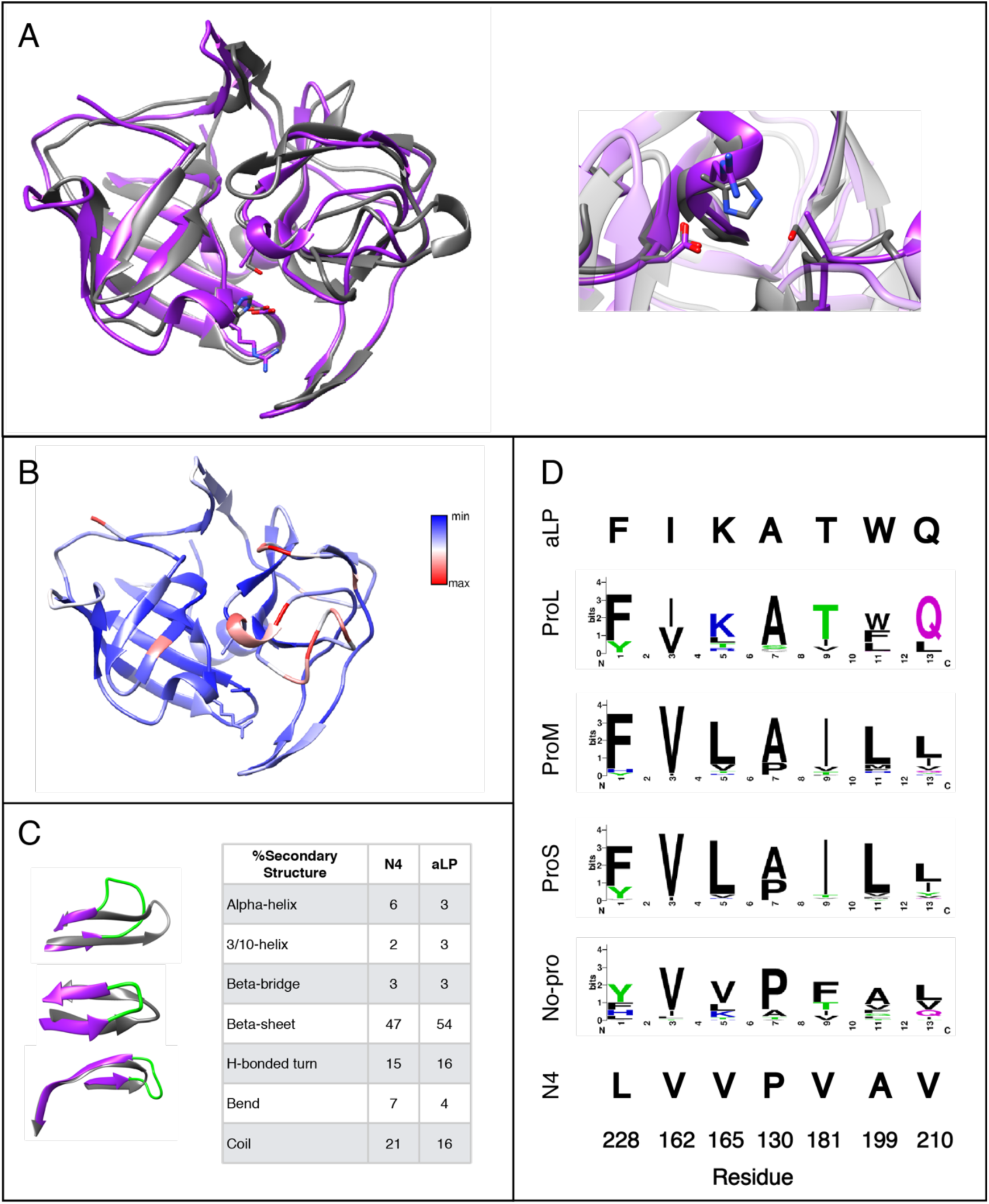
A) Superimposed ribbon diagram structures of N4 (purple) and αLP (PDB ID 4PRO; grey). Canonical catalytic triad residues are represented as atoms. Zoom inset of catalytic triad. B) N4 structure rendered by average residue B factor/residue (max 48.12, min 11.87). C) Three loops in N4 are longer than the respective loops in αLP. Overall random coil structure is 5% higher for N4 (21% vs. 16%). D) Sequence logo of Phe228 and co-varying residues for each pro region category and the individual residues of αLP and N4.

### Key residues and motifs that contribute to the unusual kinetic stability in αLP are absent in homologs without pro regions

Several unique aspects of the αLP sequence have been shown to contribute to its increased kinetic barrier. Strain caused by the out-of-plane distortion of Phe228 in αLP and other ProL homologs has been shown to correlate with increased kinetic stability.^35^ Co-varying residues (I162, K165, A130, T181, W199, Q210)^34,36^ create a tight-packed arrangement around Phe228 and are important for maintaining the out-of-plane strain in αLP. Our multiple sequence alignment together with the N4 structure confirm that Phe228 is not present in N4 and is replaced by a leucine (Supplement 1, Figure 4A). A sequence logo^37,38^ of our homolog clusters reveals this group of covarying residues are lost as the pro region decreases in size (Figure 4D). N2 and N3 contain a Tyr and His at position 228, respectively. While distortion cannot be directly evaluated without a structure, the loss of covarying residues in these homologs suggest that the Tyr and His are not distorted, as expected by the decreased kinetic stability of N2 and N3.

A beta hairpin at the pro:protease interface of αLP (residues 118–130) has also been suggested to contribute to its kinetic stability.^39^ These residues are conserved as type I turns among ProL homologs, and type I’ and I’’ turns for ProS homologs.^40^ VADAR analysis^41^ assigns the beta hairpin in N4 as a common and energetically favored “miscellaneous” type IV hairpin, indicating this region of the protein may have evolved in a way that lowers the kinetic barrier. Thus, sequence restraints that maintain the extreme kinetic stability became relaxed in the No-pro homologs as the αLP family became less kinetically stable over evolutionary time.

### Highly coupled residues in the pro:protease interface are maintained in the αLP evolutionary tree

We examined whether co-evolving residues and key contacts are maintained in the αLP family evolutionary tree by analyzing the sequences of αLP family homologs by the EVcouplings algorithm.^42^ EVcouplings on αLP (UniProtKB - P00778) revealed the top co-evolving residues for the αLP family (Figure 5A, 5B), which include five couplings between the pro region and protease sequences. Notably, the top interaction (63E:298A) has been previously shown to be important for folding catalysis.^43^ A sequence logo of these residues in the ProL and No-pro homologs shows that the sequence composition remains constant in the protease region of the sequence, with or without the presence of the pro region residues (Figure 5C). This suggests loss of the pro region is not correlated with amino acid changes at the strongly evolutionarily-coupled residues in the protease region. Mutations at other residues in the protease may be more important in encoding the energy landscape differences in αLP homologs.

**Figure 5.**
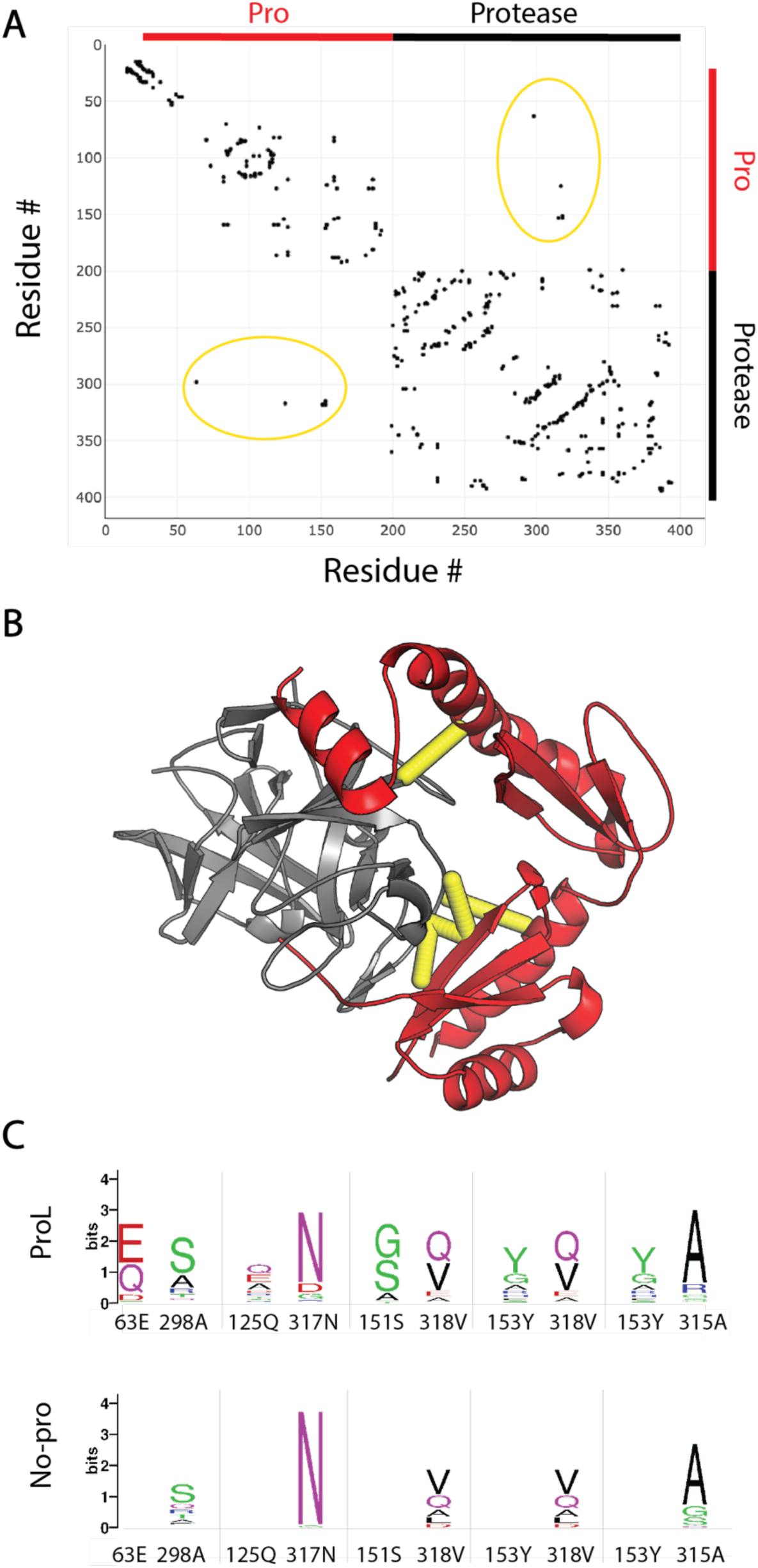
A) Contact map for αLP (UniProtKB - P00778) generated by EVcouplings (pro residues 25-199 and protease residues 200-397; numbering based on UniProtKB entry). Top intermolecular couplings are circled (yellow). B) αLP protease (grey) and pro region (red) structure with the top intermolecular couplings highlighted (yellow). Sequence logo of top couplings between the pro and protease region for ProL and No-pro homologs.

## Discussion

Using an evolutionary sequence-based approach coupled with biophysical characterization, we have uncovered evolutionary trends in the pro-region size and energy landscape of the αLP family of proteases. Having a large pro region is an ancient trait that was lost in a stepwise manner over time. While the pro region has been shown to chaperone folding to a kinetically stable and active protease, αLP homologs that have lost their pro regions are thermodynamically stable and able to fold on their own. They do not have an extremely high kinetic barrier and have likely lost their protease function. While the double beta-barrel protease fold remains conserved in these No-pro homologs, the sequence and structural motifs responsible for extreme kinetic stability have been lost.

αLP, SGPB, and N4 all show the same chymotrypsin-like serine protease fold; yet, they have very different kinetic and thermodynamic properties, highlighting that sequence encodes more than just the three-dimensional structure. Despite the recent successes in protein prediction and design^44^, we still have very little predictive power for these important dynamics. The dramatic changes in the observed folding kinetics can be a result of either a lowering of the kinetic barrier to folding, or the evolution of an alternative easier folding trajectory or barrier. Site-specific mutagenesis experiments can elucidate the residues involved in a transition state; while such experiments are feasible for the thermodynamically stable No-pro variants, the slow timescale of folding for αLP and SGPB preclude this kind of analysis. Nevertheless, an understanding of the regions determining the kinetic barrier for the No-pro homologs coupled with the MSA will yield insight into the mechanism of the kinetic barrier and its evolutionary changes.

Pro regions are biologically costly to produce, effectively acting as a single-use catalyst. Importantly, they can play the roles of acting as an inhibitor to control protease activity.^45^ This is a common feature of proteases (seen often as zymogens), ensuring that they remain inactive until they are in their intended location. The observed stepwise evolutionary loss of the domains in the pro region suggests each segment could have a specific role, through a binding interaction or another biophysical process. Each segment could be coupled to a specific aspect of the kinetic barrier, resulting in this relationship between pro-region size and foldability.

It is intriguing that the No-pro homologs have lost the canonical active-site triad needed for serine proteases (including the nucleophilic serine). This is perhaps not surprising given that the presumed reason for the high kinetic stability and requirement for a pro region is intimately tied to its function. Perhaps these No-pro homologs represent a set of pseudoenzymes (enzymes that lack some or all of their catalytic residues).^46,47^ While the best studied examples of pseudoenzymes are pseudokinases, 5-10% of enzymes across all kingdoms are thought to be pseudoenzymes.^48^ They have been found to have novel functions and act as allosteric modulators, scaffolds for signaling, and substrate sequesters. A key feature of a pseudoprotease often involves maintaining the capacity to bind the substrate. If this were the case for the αLP homologs, they could bind competing substrates, and rather than degrade them, sequester them from the environment. It would be interesting to know if any peptide binding capacity remains in the No-pro homologs studied here. Additionally, many of the No-pro homologs come from species that also have homologs with pro regions, suggesting they may be paralogs. The additional pseudoprotease could be used to “distract” competing proteases in the soil by being more susceptible to proteolysis than their kinetically stable homologs.

Protease inactivation, and thus loss of the need to be autoinhibited, may have driven the truncation of the pro region. Which features emerged first: protease inactivation, loss of pro region, increased thermodynamic stability, or decreased kinetic stability remains unknown. This intriguing question can be answered by studying ancestors in the αLP phylogenetic tree. Given the phylogenetic tree, the likely sequences of the ancestral nodes can be generated and characterizing their biophysical properties will give insight into the order of events for the αLP family. One may imagine that the protein evolved thermodynamic stability before the pro region was lost. Specifically, the last common ancestor between the ProS and No-pro homologs can tell us how the loss of a folding chaperone is coupled to thermodynamic stability.

Here, we have identified αLP homologs with a range of pro-region sizes and characterized their evolutionary history. We found several αLP homologs that fold without the assistance of a pro region, maintaining the chymotrypsin-like serine protease fold but losing key features presumed to drive kinetic stability in αLP. This protein family highlights how sequence changes and evolutionary processes can drive the height of the kinetic barrier, an important feature of a protein’s energy landscape. By studying proteins that function in extreme environments or show extreme phenotypes, we can learn not only how these “outliers” function, but also gain insights into how sequence drives the energy landscape of a variety of proteins in general.

## Accession codes

αLP: UniProtKB/Swiss-Prot: P00778.3

SGPB: NCBI Reference Sequence: WP_030706074.1

N2: GenBank: EST33180.1

N3: NCBI Reference Sequence: WP_051262886.1

N4: NCBI Reference Sequence: WP_030018218.1

## Supporting Information

Supplement 1. Full alignment of 363 homologs of αLP (PDF)

Supplement 2. Phylogenetic trees of the αLP family from protease sequences only with rooting by outgroup, midpoint, and minimal ancestral deviation (PDF)

Supplement 3. Phylogenetic trees of the αLP family from full-length pro-protease sequences with rooting by outgroup, midpoint, and minimal ancestral deviation (PDF)

Supplement 4. Protease activity assay of engineered No-pro homologs (PDF)

Supplement 5. Kinetic traces of A) N2 unfolding at 3.12 M GdmCl, B) N3 unfolding at 5.33 M GdmCl, C) N4 unfolding at 1.99 M GdmCl, D) N2 folding at 0.22M GdmCl, and E) N4 folding at 0.15M monitored by fluorescence emission at 374 nm, fit to single exponential curves (PDF)

## Acknowledgements

We thank Miriam Hood for co-evolutionary discussion, Helen Hobbs for assistance in crystal tray setting, Sophie Shoemaker for fitting pro-region distributions, and all members of the Marqusee laboratory for helpful feedback and discussion. This work was funded by National Institutes of Health Grant GM050945 (to S.M.) and a National Science Foundation Graduate Research Fellowship (to S.A.L.). S.M. is a Chan Zuckerberg Biohub Investigator.

The Berkeley Center for Structural Biology is supported in part by the Howard Hughes Medical Institute. The Advanced Light Source is a Department of Energy Office of Science User Facility under Contract No. DE-AC02-05CH11231. The ALS-ENABLE beamlines are supported in part by the National Institutes of Health, National Institute of General Medical Sciences, grant P30 GM124169.

