## Supplemental Information for "Exploring the evolutionary history of kinetic stability in the alpha-lytic protease family"

**Supplement 1.** Full alignment of 363 aLP homologs

>>id; name; signal_peptide_length; proregion_length; protease_length; alignment

>XX00000165 WP_018684498 22 1 170 MRFRA-------------------------------------
--------------------------------------------------------------------------------------------------------------
--------------------------------------------------------------------------------------------------------------
--------------------------------------------------------------------------------------------------------------
-----------------------ILCVLCVVTA-------------------------------------------------------------------------
----------------------------P--AASAT-V-TLVGGDS-VI-----V----G----GKRCVIGFNARTGSG-A--RV-
IITTGACAGT-SG--P--I-------------------G---IIPAPDVSTPLVRVG---------------------S-
GTTTVHGATEAAIGSSVCAYGPASGWRCGTVQARNQTV-NY-P--TG-------T-ITGLTRTTLCVEPGDA-G-
GPVLS-G---------N-GQAQGILIGGS--------GNCRT---------------GGVGYFAPVRPALSAYGLILY
>XX00000052 WP_019432718 19 2 191 MLFA---------------------------------------
--------------------------------------------------------------------------------------------------------------
--------------------------------------------------------------------------------------------------------------
--------------------------------------------------------------------------------------------------------------
-------------------LLTT-ALLLGATP---------------------------------------------------------------------------
----------------------------A-AAT--A-T-TVRGGDR-VY-----S----A----AGACVVAFNTQDG-SAQ--RY-
GLLPGGCGGP-GT--A--WY-A-----DPG-F-T--VPVGTTTASSFPGGGWSLLRYEP--A---VT-APGE--ISV--SG-
LPQPISSAATPTVGSRACVAGPTTGLRCGTVTAVNQTI-NV-G--GA-------V-YTGLFRTDVCVDPGTGPG-
TPAFS-G---------S--TGLGMLIGGS--------GSCAT---------------GGISYYLPLVQALGAFGLTLA
>XX00000386 WP_027772097 42 2 193 MNVPR------------------------------------
--------------------------------------------------------------------------------------------------------------
--------------------------------------------------------------------------------------------------------------
------------------------------------------------------------------------------------------------------------
APLTPYH----APPKPVPLTAAFAAVLLA-VLTALAPAA---------------------------------------------------------
-------------------------------------------------A-GAQ--E-Q-VIRGGDV-LY-----S---ST----GPSCTVGFNAGRG-
A-E--RY-ALMQGHCVRDAGL--V--WY-A-----DAA-R-T--VEVGRTEGVSFPGDDFAVIHYTN--P-A-FT-YPGE-
-LSSL-PG-GAIEITGAAGPSVGQQVCHAGGTTGLHCGTVVSVNDTV-AY-P--EG-------T-
VFGLFRSHACVEPGDG-A-VPAFS-G---------G--TALGLIVSGA---------SCSS---------------
GGATFYQPVVPALQAYGLTIS
>XX00000398 WP_030018218 31 2 192 MRPRSV-----------------------------------
--------------------------------------------------------------------------------------------------------------
--------------------------------------------------------------------------------------------------------------
--------------------------------------------------------------------------------------------------------------
--------------AVLRSLTVLAAAAVAALSTAPA----------------------------------------------------------------------
---------------------------------------A-AAA--P-T-AVRGGNV-LF-----S----A----SGRCTVGFNATKG-G-T--YY-
AIMEGRCVGG-AR--D--WY-A-----DAA-R-T--VHVGVTEAVRYPGDDYAVIRYTN--T-A-VS-YPGE--IDLG-
GG-RYLDVTGAARPVVGQSVCLPGATTGRHCGRVEAVNVSV-NH-P--EG-------T-VSGLVRTSACTEPGTA-
AGRPAVS-G---------S--TAVGLALGGG--------GNCAS---------------GGTTYLQPVLPALAAFGLTLH
>XX00000401 WP_031506745 29 2 184 MRRPP-------------------------------------
--------------------------------------------------------------------------------------------------------------

--------------------------------------------------------------------------------------------------------------------------------
--------------------------------------------------------------------------------------------------------------------------------
---------------LLATLTAILTALAAVFSAAPG------------------------------------------------------------------------------
-------------------------------------A-SAT--P-I-AIRGGDT-LY-----G----D----GTRCTVSFNARGG-G-T--YY-
GIIAGHCAGT-GT--T--WY-A-----DAA-R-T--VPVGTTAG-------NGLVRYTN--T-A-LS-YPGE--INLG-GG-
AVQDITGAAAPRYGQSVCHLGRSGGLRCGRVTGLNLTI-NY-G--GD-------I-VTGLFRTDACTEAGDA-G-
GPTFS-G---------T--TALGVIVGGS--------GNCSF---------------GGTTFHQPITEFLAAHGLSLY
>XX00000047 WP_035283421 19 3 169 MRVL-----------------------------------------
--------------------------------------------------------------------------------------------------------------------------------
--------------------------------------------------------------------------------------------------------------------------------
--------------------------------------------------------------------------------------------------------------------------------
------------------------I-ALVVLGAPV--------------------------------------------------------------------------------------
------------------------------A--AAAPT-A-AIVGGDT-VL-----V----N----GVPCRVAFNAKTPAN-A--HR-
VIVLGACTGS-AQ--P--V--------------------T--AAVPPPGTPTAQVRVG---------------------S-
RLVTVTGTRSPAIGSSVCTSSPTSGYRCGTVQAVNQTV-NF-P--GG-------A-ITGLTRTNTCFEFGDA-G-
SPVLA-G---------T--QAVGVLLAGS--------GNCTV---------------GGTSYYVPINQVLSQNGLALY
>XX00000059 WP_042185058 26 3 195 MTLPR---------------------------------------
--------------------------------------------------------------------------------------------------------------------------------
--------------------------------------------------------------------------------------------------------------------------------
--------------------------------------------------------------------------------------------------------------------------------
-------------------FRCLLGLFLVFWLVVPPAAQARVP------------------------------------------------------------------
---------------------------------------------------A-PLGGGSP-LET----G----S----SYRCTAGFAAVGG-S-T--GY-
LIASTACGRP-GD--T--VR-S-----A-----G--RVVGVVSGAPTPPTGAIVIRVTNTVD-W-R--LVGS--IPPV--D-
SRVTVTGSVEAPVGASVCKYGATSGWRCGVIQGKNQTV-TF-P--EG-------S-LHGLTRTNVCAEPGDT-G-
GPFVS-G---------T--QAQGVLVGAT--------GNCTS---------------GGTSYFLPVNRVLATYGLTIL
>XX00000131 WP_027742062 45 3 192 MNLPR---------------------------------------
--------------------------------------------------------------------------------------------------------------------------------
--------------------------------------------------------------------------------------------------------------------------------
-----------------------------------------------------------------------------------------------------------------------------
APLTPYHAPHGAPVKLAPLAAVLAAVLLA-VLTALAPPA------------------------------------------------------------
-------------------------------------------------------A-AAQ--E-Q-VVRGGDV-IY-----S---ST----
GRSCPVGFNAGRG-G-E--RY-ALMQGHCVGDSGA--V--WY-A-----DAA-R-T--
VEIGRTGGVSFPVSDYALIRYNN--P-D-FT-YPSE--LSTV-PG-
GAIEITGAAQPSVGQQVCRAGRTTGVHCGTVVSVNDTV-SY-P--EG-------T-VFGLFRATLCVEPGDS-A-SA-
YS-G---------G--TALGLLVSGS---------PCSS---------------GGTAFYQPVVPVLSAYGLSIP
>XX00000396 WP_054238350 31 3 193 MRRPHT-------------------------------------
--------------------------------------------------------------------------------------------------------------------------------
--------------------------------------------------------------------------------------------------------------------------------
--------------------------------------------------------------------------------------------------------------------------------
----------------RLTMFTTVVTA-LVALLFGTGTGA----------------------------------------------------------------------
------------------------------------------T-ALK--V-P-TIRGGTL-LY-----A---AT----GAQCVVGFNAVSG-S-N--RY-
AVMAGHCSGT-YT--T--WY-A-----DAA-H-T--LPVGVTAGASYPIDDYGVVRYTT--S-A-LV-LPGD--IALG-
GG-VYQDVTGAANPAIGQSVCHVGRTSGVHCGTVTAVNATV-NH-A--EG-------T-VHGLFRSTACSEPGDT-
G-APAYS-G---------S--IALGFAVGSS--------GNCAS---------------GGVTYYQPVAEVLSVFGLTLY

>XX00000399 WP_030667422 29 4 191 MRPRTA-------------------------------------
---------------------------------------------------------------------------------------------------------------------------------
---------------------------------------------------------------------------------------------------------------------------------
---------------------------------------------------------------------------------------------------------------------------------
----------------VARLLCAVLIG-AFALMAAPAAT-------------------------------------------------------------------------
-----------------------------------------A-APA--Q-T-PLRGGTL-LF-----S----P----AGRCVVSFNATNG-T-H--YY-
GIMAGHCGGV-GT--K--WY-A-----DPG-L-T--TQVGITETAVFPGKDYALVRFTN--P-D-YS-YPSE--VAA--GV-
QIIRINRAANPVVGQSVCRSGPTTGMRCGRVTATNVSI-TY-P--QG-------T-VTGLFAATVCAEPGDS-G-
GPTFS-G---------D--AALGFLVGGS--------GNCTS---------------GGTTYHQPLAPVLSAHALRVY
>XX00000408 WP_019731792 19 4 200 MAMV----------------------------------------
---------------------------------------------------------------------------------------------------------------------------------
---------------------------------------------------------------------------------------------------------------------------------
---------------------------------------------------------------------------------------------------------------------------------
--------------------------A-LLVAMTHPRAA-------------------------------------------------------------------------------
-----------------------------------A-ADV--R-I-PLGGGAG-IVV----N----G----DTMCTLTTIGGDAAG-D--LI-
GFTSAHCGGP-GA--Q--VA-A-----EGA-ENA--GILGTMVA-GNDNLDYAVIKFDP--A-K-V--T--P--VANF-
NG---FLISGIPDPAFGEIACKQGRTTGNSCGVT---------W-G--MG-------Q-TPGTIVMQVCGQPGDS-G-
APVTV-N---------N--QLVGMIHGAF--------SDNLP---------------TCVIKYIPLHTPVMSFNAILA
>XX00000088 WP_037752774 35 5 193 MSRRRT-------------------------------------
---------------------------------------------------------------------------------------------------------------------------------
---------------------------------------------------------------------------------------------------------------------------------
---------------------------------------------------------------------------------------------------------------------------------
---------APPLRTFATAAALALLGA-LLTVLAPPASA---------------------------------------------------------------------
---------------------------------------------Q-AAA--A-G-ELRGGDA-LY-----N---SG----GYSCTLGFNAVGS-G-A--
TY-GIIPGHCADV-GS--S--WS-A-----DVGGS-R--VNVGTTSGSSFPSNDYGVVRYTN--T-S-VS-YPGE--VKGS-
GG-APVDITGAVDPAPGMALCHYGRVSGYHCGTVQAVNVTV-NF-P--EG-------V-VSGLFRSNICSEPGDN-
G-GPAFS-G---------G--KAVGIIIGST--------GNCST---------------GGVTYYQPIKEILAVYGLTLS
>XX00000187 WP_030227892 37 5 190 MSTV------------------------------------------
---------------------------------------------------------------------------------------------------------------------------------
---------------------------------------------------------------------------------------------------------------------------------
---------------------------------------------------------------------------------------------------------------------------------
------PSEGRSLRTVLRAVVALAL-LLGWTAVSGSAAHA---------------------------------------------------------------
-------------------------------------------------G-REA--A-V-EVRGGDT-LY-----S----T----GTRCTVGFNARSG-S-T-
-TY-ALLSGRCAQG-AR--T--WY-A-----DPA-L-T--VPVGVTAGVSFPGNDYASIRYTN--T-A-VV-YPGE--
VSLGT---GARDITGAANPVVGQPICHVGRTTGYRCGTVQAVNLTV-NY-G--DG-------I-
VYGLFRSTICSEPGDA-G-GPAFS-G---------G--TALGIIVGST--------GTCAS---------------
GGFTYYQPVTEWLSAYGLTVY
>XX00000391 WP_027741836 35 5 193 MSRRRT-------------------------------------
---------------------------------------------------------------------------------------------------------------------------------
---------------------------------------------------------------------------------------------------------------------------------
---------------------------------------------------------------------------------------------------------------------------------
---------APPLRTFATAAALALLGA-LLTVLAPPASA---------------------------------------------------------------------
---------------------------------------------Q-AAA--A-G-ELRGGDA-LY-----N---SG----GYSCTLGFNAVGG-G-A-
-TY-GIIPGHCADV-GS--S--WS-A-----DVGGG-R--VHVGTTSGSSFPGNDHGVVRYTN--T-S-VS-YPGE--

VKGS-GG-APVDITGAVDPSPGMALCHYGRVSGYHCGTVQAVNVTV-NF-P--EG-------V-
VSGLFRSNICSEPGDN-G-GPAFS-G---------G--KAVGIIIGST--------GNCST--------------
GGVTYYQPIKEILAVYGLTLS
>XX00000397 WP_055511461 29 5 193 MRRLRT-------------------------------------
-------------------------------------------------------------------------------------------------------------------------------
-------------------------------------------------------------------------------------------------------------------------------
-------------------------------------------------------------------------------------------------------------------------------
----------------CLAMLTTAVTA-VVSLLFGTVASA----------------------------------------------------------------------
------------------------------------------A-ALQ--L-T-TIRGGTM-LY-----A---AT----GAQCVVGFNATGN-G-V--
FY-GVMVGHCSGT-HT--T--WY-A-----DAA-R-T--VQVGVTAGASFPIDDYGVVRYTS--N-T-LN-LPGD--
IALG-GG-VYQDITGMASPAVGQSLCHAGRTSGVHCGRVTAVNATV-NY-S--EG-------A-
VHGLVRSTTCSEAGDT-G-APAYS-G---------T--LAVGFLVGAS--------GNCTS--------------
GGVSYHQPVAEVLSNFGLTVY
>XX00000170 WP_053723080 20 6 184 MRTLL----------------------------------------
-------------------------------------------------------------------------------------------------------------------------------
-------------------------------------------------------------------------------------------------------------------------------
-------------------------------------------------------------------------------------------------------------------------------
---------------------LVLT-LLAAVATPATAAPAALP---------------------------------------------------------------------
-----------------------------------------------T-PLEAGTP-LYP----A----N----GLRCANGFNARG--------F-
LIVSPSCGIV-GT--V--VR-G-----P---G-L--VDIGPIVAVRQ---TYAIVRITNTAA-W-V--QRPT--IAG---Y-
AGTTIRGSAETPVGGQVCAAGRTTGLRCGAVQAKNVTV-NY-P--GG-------T-VYGLTRTSVCTEPGDQ-W-
APLFT-G---------A--QAQGHSLGGS--------GNCST--------------GGASYFEPLNRVLAAERLTLV
>XX00000096 WP_013020008 25 7 188 MKLAR---------------------------------------
-------------------------------------------------------------------------------------------------------------------------------
-------------------------------------------------------------------------------------------------------------------------------
-------------------------------------------------------------------------------------------------------------------------------
-------------------TLTMLAA-VVALTAVPTTAVAHEPEARG-----------------------------------------------------------
------------------------------------------------------G-VVTAGLP-FST----G----T----TSRCTTGFATEHA-SLG--DG-
FLAAGHCGSV-GD--R--AY-V-----N-----N--VAVGVVRLRASNGLTWLWVELDD--G-W-T--ASPA--LP------
GGGSITDDRETPIGGSVCRIGSTTGWHCGTVQAKNATV-SY-P--QG-------T-VTGLTRTNVCAEPGDN-G-
GPFVS-G---------S--SAQGIALGGS--------GNCSS--------------GGTTYFYPVRRVFSEAALSII
>XX00000389 WP_047018227 36 8 191 MNLRR---------------------------------------
-------------------------------------------------------------------------------------------------------------------------------
-------------------------------------------------------------------------------------------------------------------------------
----------------------------------------------------------------------------------------------------------------------------
APLTP----HAVPLRLTPAAAVLAAVLLA-VLTALAPPA----------------------------------------------------------------
---------------------------------------------------A-GAQ--D-Q-AVRGGDV-LY-----S---ST----GPSCPVGFNAGRG-
G-E--RY-ALMQGRCAAAAGP--V--WY-A-----DAA-R-T--VEVGRTEAVNFPGGDFALIRYTN--P-D-FG-YPSE-
-VSA--GI-GVIGITGAAQPSVGRQVCHAGRTTGVHCGTVVSVNDTV-SY-P--EG-------T-
VSGLFRSHVCVEPGDG-A-GPAWS-G---------G--TALGLIVSGG---------SCTS--------------
GGATFYQPVVPALQAYGLTLP
>XX00000394 WP_030360152 28 8 194 MRLRLI---------------------------------------
-------------------------------------------------------------------------------------------------------------------------------
-------------------------------------------------------------------------------------------------------------------------------

---------------------------------------------------------------------------------------------------------------------------------
---------------IAQLLCAALA-ALGLLAAPTATAAPA---------------------------------------------------------------------
-----------------------------------------P-ASA--Q-T-AIRGGDV-LY-----S----A----SGRCTVAFNVSNG-T-R--FY-
GIMGGHCGTV-GT--Q--WF-A-----DPR-L-T--VPVGVTETVVFPRSSYALVRYTN--T-N-LT-YPSE--IRA--GS-
QIIRINRAPQPVVGQQICRTGPTTGMHCGTIQSLNATV-TF-P--QG-------T-VYGLIRTNVCAEPGDA-G-
GPGFA-G---------D--AALGIVVGGS--------GNCSS---------------GGTTFFQPLPPILNAHGLRVT
>XX00000395 WP_058847733 28 8 192 MRLRTI--------------------------------------
---------------------------------------------------------------------------------------------------------------------------------
---------------------------------------------------------------------------------------------------------------------------------
---------------------------------------------------------------------------------------------------------------------------------
----------------LLQTLTTALA-ALTLAAAPAAGAAPA---------------------------------------------------------------------
------------------------------------------D-RAE--Q-G-VLRGGDR-LF-----A---PG----NARCSVAFNATDG-S-R--
FY-GLLPGHCGGR-GT--R--WF-A-----DPG-L-T--VPVGETEVSRFPGSDYALVRYTN--P-D-YS-YPSE--VSA--
GG-QSHRIDRAAQPTVGQRVCQVGSASGHRCGTVQAVNVSV-TH-P--EG-------V-VHGLFRSTLCLEPGDA-
G-APAFG-G---------N--AALGIAVGGS--------GNCRN---------------GGTTYHQPVAPVLAAHGLRVY
>XX00000400 WP_051307750 20 8 191 MTA-------------------------------------------
---------------------------------------------------------------------------------------------------------------------------------
---------------------------------------------------------------------------------------------------------------------------------
---------------------------------------------------------------------------------------------------------------------------------
---------------------------------------------------------------------------------------------------------LL-A----------------A-
LGA--------LALSAGPSAAQER-----N-AVQ--Q-Y-AVRGGDT-VY-----G---DH----GGACQVGFNARSSSG-T--
RY-GIVLGACAEG-AS--T--WY-A-----DAA-R-T--VEVGTA-GGGLPGPTAALIRYTN--P-E-VA-FPGE--ITV--
GG-DHVDITGSAAPRVGQNVCHAGRTSGLRCGVVQAVNVTI-QF-P--DG-------I-VYGAFRATDCAEPGDA-
G-APGFS-G---------T--VALGLVLGGS---------NCAA---------------GGSAYHLPVREVLSTYGLAVY
>XX00000372 WP_004936561 36 9 192 MSPMRE-------------------------------------
---------------------------------------------------------------------------------------------------------------------------------
---------------------------------------------------------------------------------------------------------------------------------
---------------------------------------------------------------------------------------------------------------------------------
-------------MPLRTIVRAVVVLAL-LLGGSAVVGPAAHAGQRA-----------------------------------------------------
---------------------------------------------------E-RQT--A-V-AVRGGDT-LY-----S----A----GARCTVGFNARSG-
T-T--LY-ALVPGRCAQG-SS--T--WY-A-----DPA-L-T--VAVGVTAGVSFPGNDYASIRYTN--T-A-VA-YPGE--
VSLGAGV-GTRDITNAANPVVGQSICHVGRTTGYRCGTVQAVNLTI-NY-G--GG-------T-
VYGLFRSTICSEPGDA-G-GPAFS-G---------G--TALGIIVGSS--------GNCSS---------------
GGVTYYQPVTEWLSAYGLSIY
>XX00000382 WP_055594556 36 9 192 MSPMRE-------------------------------------
---------------------------------------------------------------------------------------------------------------------------------
---------------------------------------------------------------------------------------------------------------------------------
---------------------------------------------------------------------------------------------------------------------------------
-------------RPLRTVVRAVVVLAL-LFGWSAAIGSAAHAGERA-----------------------------------------------------
-------------------------------------------------G-QQA--A-V-AVRGGDT-LY-----S----T----GTRCTVGFNARSG-
S-A--LY-ALVSGRCAQG-AR--T--WY-A-----DAA-L-T--VAVGVTAGVSFPGNDYASVRYTN--T-A-VA-YPGE--
VSLGADA-GTRDITNAANPVVGQSICHVGRTTGYRCGTVQAVNVTI-NY-G--GG-------T-
VYGLFRSTICSEPGDT-G-GPAFS-G---------G--TAIGIIVGSS--------GNCSS---------------
GGFTYYQPVAEWLSAYGLSVY

>XX00000388 WP_047015861 35 9 192 MSRRRT-------------------------------------
--------------------------------------------------------------------------------------------------------------------------------
--------------------------------------------------------------------------------------------------------------------------------
--------------------------------------------------------------------------------------------------------------------------------
---------APPLRTTATAAVLAVLGA-LLTVLAPPASASDAG-------------------------------------------------------------
-------------------------------------------------R-PAA--A-T-AVRGGEA-LY-----G----P----SYACTLGFNAVSG-G-
T--SY-GIIAGHCADV-GS--T--WA-V-----EAEGG-R--VSVGVTSGSVFPGSDYGVVRYTN--T-S-VS-YPGE--
VKGS-GG-APVDITGAADPAPGMSLCHYGRVGGYRCGTVQAVNLTV-NF-P--GG-------T-
VSGLFRSSICSEPGDT-G-GPAFS-G---------G--RAVGIIVGGS--------GNCTS---------------
GGVTYYQPVKEILAAYGLTLS
>XX00000205 WP_005153998 29 10 185 MFGC----------------------------------------
--------------------------------------------------------------------------------------------------------------------------------
--------------------------------------------------------------------------------------------------------------------------------
--------------------------------------------------------------------------------------------------------------------------------
---------------SRRKLI------SSAVSAAPSAV-------------------------------------------------------------------------------
-----------------------------------G-AAR--Q-S-TLKGGDA-MY-----T----VSGREPYYCSAGFAILVK-D-V--RY-
ILTAGHCTEH-GL--D--WE-G----------------IGPNVDTHFPVTDYGRIRDLS--G-S----GPSQ--VDLY-DG-
RTQRILRAGIPQVGQTVCKSGMTTKKTCGKVLGLNQTV-NY-G--KD------KQ-VIGTIATDIPIGSGDS-G-
GPLFY-G---------G--VGYGVLSGGN---------------------------SQVSFYQPLRTVLTAYGATLA
>XX00000220 WP_051674723 33 10 184 MRIGH----------------------------------------
--------------------------------------------------------------------------------------------------------------------------------
--------------------------------------------------------------------------------------------------------------------------------
--------------------------------------------------------------------------------------------------------------------------------
----------RSTVRAAVRRLGPAVV-ALAAFAAPTAAAGGQAA---------------------------------------------------------
---------------------------------------------------A-ATV--R-P-QVSAGDT-IY-----A----NN---GTACRTAANARSA-
S-T--YY-IIMPGHCTLG-TS--A--WY-T-----SAA-Y-T--TYIGPTVATSFPGNDYGLVRYDN--P-A-VP-HPGG-----
--------GFTSVGNPYAGERVCRTSPVSGLHCGTVTALNATV-NY-G--GG------QV-VSGLIATTVCTEPGDT-G-
GLLFS-G---------S--TALGLFSGGS--------GTCSS---------------GGTGYYQPLAEVLAAYGLSLY
>XX00000183 WP_053803022 31 11 190 MRPRPI---------------------------------------
--------------------------------------------------------------------------------------------------------------------------------
--------------------------------------------------------------------------------------------------------------------------------
--------------------------------------------------------------------------------------------------------------------------------
-------------AVRRFLAVVATAALTAPAAAPPAAAAPMGAP----------------------------------------------------------
-------------------------------------------------S-RTA--V-T-AIRGGNV-LY-----S----A----AGRCTVGFNATKA-G-
T--YY-ALMAGRCVGG-AK--D--WY-A-----DAA-R-T--VHVGVTEAVRFPGTDYAVIRYTN--P-A-VS-YPGE--
IDI--GG-SHLDVTGAAQPAVGQTVCLPGSTSAVHCGPVRALNVSV-SY-P--EG-------T-
VSGLVQTAACAEPGAV-G-RPAVS-G---------R--TAIGIVIGGS--------GSCAF---------------
GGATYLQPVVPALQAFGLTLH
>XX00000387 KOT81145 20 11 190 MA------------------------------------------------------
--------------------------------------------------------------------------------------------------------------------------------
--------------------------------------------------------------------------------------------------------------------------------
--------------------------------------------------------------------------------------------------------------------------------
-------VVATAAL-TAPAAAPPAAAAPMGAP----------------------------------------------------------------------------
--------------------------------S-RTA--V-T-AIRGGNV-LY-----S----A----AGRCTVGFNATKA-G-T--YY-

ALMAGRCVGG-AK--D--WY-A-----DAA-R-T--VHVGVTEAVRFPGTDYAVIRYTN--P-A-VS-YPGE--IDI--
GG-SHLDVTGAAQPAVGQTVCLPGSTSAVHCGPVRALNVSV-SY-P--EG-------T-VSGLVQTAACAEPGAV-
G-RPAVS-G---------R--TAIGIVIGGS--------GSCAF--------------GGATYLQPVVPALQAFGLTLH
>XX00000057 WP_051061688 43 12 187 MPRT--------------LRRTRPVERS---------
-------------------------------------------------------------------------------------------------------------------------------------
-------------------------------------------------------------------------------------------------------------------------------------
-------------------------------------------------------------------------------------------------------------------------------------
------------PHARRAAAVGAAALAAVFAA-ALLAAPAPAAADGAATGQVQQ--------------------------------------
--------------------------------------------------------------------------A-QIVAGDP-FIN----T-S-AP----
HSRCTIGAVVTV--------G-FLAASTCGSA-GD--R--IS-G-----R---D-G---GTGTVAWSSK-ESGVLLVEADD--N-
W-R--PTPY--INTH-GG-GRAVVRGTTEAPVGAGVCRTGSTTGWHCGIIQAKNQTI-RT-P--DG-------T-
VYNVTRTNVCAEPGDQ-G-GAFYS-G---------G--HLQGLTLGGS--------GNCAS--------------
GGTTFFLPIGPILARYALTPV
>XX00000390 WP_031011063 30 12 192 MRPGPT-------------------------------------
-------------------------------------------------------------------------------------------------------------------------------------
-------------------------------------------------------------------------------------------------------------------------------------
-------------------------------------------------------------------------------------------------------------------------------------
--------------AALRSLAALAAA-SAVLATAPPAGAAPGAGPS--------------------------------------------------------
-------------------------------------------------R-TAA--A-T-AVRGGNV-LY-----S----A----SGRCTVGFNATKG-G-
R--YY-AILEGRCVGG-AQ--D--WY-A-----DAA-R-T--VHAGVTEAVRFPGEDYALVRYTN--P-A-VS-SPGE--
VDLG-GG-RHLDITGAARPVVGQSVCLPGSTTGQRCGRVDAVNVGV-TY-P--EG-------T-
VSGLVRTSACTEPGTA-AGRPAVS-G---------S--SAVGLAVGGS--------GTCAS--------------
GGTTYLQPVVPVLQAYGLTLH
>XX00000192 CCH34430 24 13 182 MRPLR-------------------------------------------------
-------------------------------------------------------------------------------------------------------------------------------------
-------------------------------------------------------------------------------------------------------------------------------------
-------------------------------------------------------------------------------------------------------------------------------------
--------TPLPALAT-LIALLATPATATATATNNTNSPVP----------------------------------------------------------------
---------------------------------------------T-PLEAGTT-LGS----P----P----NARCANGFNVRG--------H-
LLISTRCANT-TD--P--VT-G-----P---G-G--GQIGPITRIRD---TYAVVKVQDPTK-W-N--QLPR--IV----G-
SATPITGSAEAPIGARVCAGSALQGWQCGTLQAKNQTV-HH-P--SG-------S-ITGLTRTNLCAQPGND-W-
LPVVS-G---------S--QAQGHLSGGS--------GTC-----------------TSYFYPINRILQAEAFTLV
>XX00000383 WP_051262886 35 13 192 MRRP----------------------------------------
-------------------------------------------------------------------------------------------------------------------------------------
-------------------------------------------------------------------------------------------------------------------------------------
-------------------------------------------------------------------------------------------------------------------------------------
--------------IVPAVIAAAVA-ALTLTAAPAAASTSAPPGT------------------------------------------------------------
---------------------------------------------S-PSA--A-P-AVRGGDV-LY-----G---EG----GGACTVGYNARNT-N-T--
YF-GIMAGHCAGT-SQ--T--WY-A-----DPG-H-T--VKVGVTAGVSFPGDDYALVRYTN--P-D-LS-YPGE--
INLG-GG-RTLDITGAANPVTGQSVCQASQVTGVHCGTVQAVNVTV-NY-P--WG-------T-
VNGLFRTNLCSGPREG-G-APVFS-G---------S--TALGVGSAAS--------GNCQS--------------
GGTTFHQPVVEVLAAYGLSIY
>XX00000380 KDS85990 27 15 193 MR---E-------------------------------------------------
-------------------------------------------------------------------------------------------------------------------------------------

------------------------------------------------------------------------------------------------------------------------------
------------------------------------------------------------------------------------------------------------------------------
RPLRTLVRAVVVLAL-LLGWSASAGPAAQAGEQA---------------------------------------------------------------
-------------------------------------G-QQA--A-S-TVRGGDT-LY-----S----N----GTRCTVGFNARSG-S-T--VY-
AFVSGRCAQGAGT--T--WY-A-----DAA-Q-T--VGVGVTAGVSFPGNDYASIRYTN--T-A-VA-HPGE--
ISLGAGA-GTQDITSAAHPVVGQSICHVGRTTGHQCGIVQAVNVTV-NY-S--GG-------T-
VYGLFRSTICSEPGDA-G-GPAFS-G---------G--TALGIIVGSS--------GNCSS--------------
GGVTYYQPVTEWLSAYGLSVY
>XX00000378 EST33180 40 17 193 MNPHR----------TERPFRRAFRTAAS-------------
------------------------------------------------------------------------------------------------------------------------------
------------------------------------------------------------------------------------------------------------------------------
------------------------------------------------------------------------------------------------------------------------------
-------------------VVAALTL-LLGWFTAAGSAAYAQDSGASAT---------------------------------------------------
---------------------------------------------T-AGK--A-V-EVRGGSV-IY-----S---NA----GTRCTVGFNARSG-A-
T--YY-GLVSGRCAQG-AT--N--WY-A-----DAT-L-S--VFVGTTAGSSFPVNDYALIRYTT--S-TAVT-FPGE--
VTLG-GS-GVQDITGAANPSIGQSLCHVGRTTGVRCGTVNGVNVTV-NY-P--EG-------V-
VSGLFRSNICSEPGDV-G-GPAFS-G---------G--TALGIIVASS--------GNCSS--------------
GGTTFYQPVVEWLAVYGLSLY
>XX00000379 WP_024488563 40 17 192 MSQHR-------------------------------------
------------------------------------------------------------------------------------------------------------------------------
------------------------------------------------------------------------------------------------------------------------------
------------------------------------------------------------------------------------------------------------------------------
-------------------ALAALTL-LLGWSAATGSTAYAQEASAPATAPTA--------------------------------------------
-----------------------------------------------A-NSR--A-V-EVRGGDI-LY-----G---SG----GIRCNVGFNAHSG-T-
A--TY-GMMPGHCALG-ST--N--WY-A-----DAA-R-T--IFVGTTAGSSFPSNDYSVISYVT--G---VN-ALGE--
VGLGPGG-GVQDIVSAANPTVGQSICHVGGTTGIHCGTVTGTNVTI-NY-P--EG-------M-
VSGLFSSNICSEPGDV-G-GPGFS-G---------T--TALGIIVASS--------GSCAS--------------
GGVTYYQPVVEWLSAYGLSIY
>XX00000376 WP_051438477 50 18 192 MRSLAT-------------------------------------
------------------------------------------------------------------------------------------------------------------------------
------------------------------------------------------------------------------------------------------------------------------
------------------------------------------------------------------------------------------------------------------------------
----------------GTVLALLGA-LLTLLAPPAAAAPPAAPAPAVPAA-----------------------------------------------
--------------------------------------------A-AGA--A-G-EVRGGDA-LY-----G----G----GFACTLGFNAVAG-G-
A--TY-GIIPGHCATA-AD--S--WA-A-----DVDGA-R--VSVGVTAGYTFPGSDYGLVRHTN--T-S-LS-YPGE--
VRGS-GG-VPVDITGATDPQPGMTLCHYGRVTGYQCGRLQQVGVTV-NY-P--GG-------V-
VYGLFRSDICSEPGDT-G-GPAFS-G---------G--RAVGIIVGSS--------GNCVT--------------
GGSTYYQPIKEILSAYGLTLP
>XX00000143 WP_017621136 26 20 195 MTPIP--------------------------------------
------------------------------------------------------------------------------------------------------------------------------
------------------------------------------------------------------------------------------------------------------------------
------------------------------------------------------------------------------------------------------------------------------
---------QRLLRLLGASLLALGM-LLAFAPAAAAAALGSPTPAPASDEP-------------------------------------------
---------------------------------------------------------A-RVIAGDP-IHL----D-G-ND----LNRCAIGAVVTTS-

DGT--YG-FLTSYMCGNP-GD--T--II-G-----P---D-G--RVIGTVVWSSR-QDGVAFVELHD--G-W-I--PTSY--
VRTS-PGPPTRSIQGSREASIGAAVCRAGSATGWHCGTIQAKNQSI-RF-D--WG-------T-
MYGLTRTSVCAEAGDL-G-GPFIS-G---------N--QIQGFLIGGT--------GSCRT---------------
GGTTYFVPINPILAKYNLHLF
>XX00000406 WP_010909015 32 20 302 MSRQPH--------R--S------------------------
-----------LWRSWLVSTLA-------------------------------------------------------------------------------------------
------------------------------------------------------------------------------------------------------------------------------
------------------------------------------------------------------------------------------------------------------------------
-----------------------------------AL-GLSLAVVPGSATALDRFSNRPPLP----------------------------------------------
-------------------------------------------------------------------------------------NMVAPISTRLGYNSAVGAS-S--
GV-VLTNNHVISG-AT--D--IS-A-------VGN-G--KTYGVDVVGYDRTQDVAVLQLRG--A-S-N--LPT-------------
-AV-IGGDVAIGEPIVALGNTGSVLPGRVVALNQTV-QA-S--EE-------T-LSGLIQVDAPIKPGDS-G-GPVVN-S-
-------RG--QVVGMNTAAT--------DNYKL---------------GGQGFAIPIGQAMEVVGAIRS
>XX00000381 WP_051265161 37 26 185 MLSRR------------------------------------------
------------------------------------------------------------------------------------------------------------------------------
------------------------------------------------------------------------------------------------------------------------------
------------------------------------------------------------------------------------------------------------------------------
---------------LLLLGAAPASA-ASAGTTTPGTATTARPGGEQP------------------------------------------------------
---------------------------------------------------A-AAQ--Q-A-VLRGGDR-IY-----A----P----GAVCTLGFNATDG-T-
Q--DY-GIASGRCLSS-AT--T--WY-A-----DAA-M-T--VPAGTTADTSFPGDDYGLLHYTN--P-D-VA-RPGQ--
ISA--GG-SIIDITGSASPMVGAVVCHAGHTTGMHCGTILSVNLTV-NY-P--EG-------S-
VHGLFSSNVSAGAGDT-G-VPAFS-G---------T--TALGFVTGAS---------------------------
GGSTFYQPVTEVLAAYGLTLL
>XX00000190 WP_028438174 25 27 186 MSRRR------------------------------------------
----L----------------------------------------------------------------------------------------------------------------------
------------------------------------------------------------------------------------------------------------------------------
------------------------------------------------------------------------------------------------------------------------------
--------------ATGGALLAVLLLGSLPAQAAHATASADASVAASTDAQQNRL-------------------------------------------
----------------------------------------------------------------P-ATQ--Q-V-TVRGGDS-VY-----A----A----
GRVCTVSFNATDG-T-N--DY-AISPGHCVEG-AT--T--WY-A-----DPA-L-T--
VVVGTTTGSSFPGNDYGLIRYTN--P-D-VS-RPGE--VNTG-SG-
GTVDITRAASPTVGRSMCHVGRVSGFQCGTVTAVNVSI-SY-P--EG-------T-VYGLFQSTAHSEPGDQ-G-
GPAFS-G---------G--TALGFIVGAS---------------------------AGSTFYQPITEVLSMYGLSLS
>XX00000128 WP_027762115 41 30 183 MSTS--------------------------------------------
------------------------------------------------------------------------------------------------------------------------------
------------------------------------------------------------------------------------------------------------------------------
------------------------------------------------------------------------------------------------------------------------------
---------------RKRLVAVLAGLLAATVWAPGAVAEAGPVLAGGGPAGTTA-------------------------------------------
-------------------------------------------------------------R-DAS--S-Q-TISPGDP-IF-----S----G----GARCTAGFNVTDG-
T-D--VF-IVTAGHCTSV-GS--T--WY-A-----DPQ-L-T--IPIGPTVSSSFPGNDYGLIRYDN--P-D-LT-PEGG-------
--------YSLGSPSVGQQVYIRTPQTGIHSGTITALNATI-DY-G--PD------GI-IYGLIRTNICLPVGAG-G-GTPLF-
G-------QND--QALGIASGGS--------GSCSS---------------GGTSYFQPLNEVLSTFGLTLY
>XX00000375 WP_052708401 40 38 186 MDFRRS----------------------------------------
------------------------------------------------------------------------------------------------------------------------------

--------------------------------------------------------------------------------------------------------------------------------
--------------------------------------------------------------------------------------------------------------------------------
----------------EPSSPAPPPTA-STAGPDASGRADSDRPSSQRP----------------------------------------------------
------------------------------------------------A-DAR--Q-S-VIRGGDP-VY-----A---PG----GLVCTIGFNATDG-S-
D--DY-GIAAGHCLGG-AE--T--WY-A-----DAA-L-T--IPAGTTAGASFPGNDYGLIRYTN--P-D-VA-TPGE--
IDT--AG-GPVDITGSGSPTVGSSVCQSGQVTGMHCGTVTALNITI-SY-P--EG-------V-
VSGLFSSNIQTEPREG-G-TPAFS-G---------N--TGLGIVVGSS---------------------------
AGATYYQPVAEVLAAYGLTLL
>XX00000374 WP_055486726 40 43 186 MRSR----------------------------------------
--------------------------------------------------------------------------------------------------------------------------------
--------------------------------------------------------------------------------------------------------------------------------
--------------------------------------------------------------------------------------------------------------------------------
--------------AQAAHAAHAAH-AAPDAGTPGTAGPATQDRDRDAATQDRD--------------------------------------
-----------------------------------------------------------P-AAG--Q-V-AVRGGDS-LY-----A----A----
GRVCTVGFNATNG-G-T--DY-AIVSGRCVTG-AD--T--WY-A-----DPA-M-T--
VPVGSTAGSSFPGNDYGLIRYTN--P-D-VA-RPGE--VETG-AG-
GPVDITRSASPAVGQSMCHAGRVSGLQCGTVQAVNVSV-NY-P--EG-------T-VSGLFRSTAHSEPGDE-G-
GPAFS-G---------G--TALGFIVGAG---------------------------SGSTFYQPVGEVLAAYGLALS
>XX00000042 WP_020521700 25 49 192 MRHHRL------------------------------------
--------------------------------------------------------------------------------------------------------------------------------
--------------------------------------------------------------------------------------------------------------------------------
----------------------------------------------------------------------------TA-L----V-FAC--ALVLLPA-AP-AQ-AAS-------
--------------------------------------------------------------------------LN------TP-GLSWGPDPVTGEVVVTADATV-T-
---EA--Q-VAA-LRRT---------------YPV---------RR---------V-----D-GQF--R-T-LIAGGQA-IY-----S----A----
SSRCSLGANVRKG-S-V--AY-FVTAGHCTNT-GA--T--WY-S-----NAA-R-T--
TVLGSRTGTSFPGNDYGIVRYSN--T-S-IA-HPSA--VYTY-PG--
TVTIRGAGNATVGMSVCRSGSTTGVRCGVVTGLNQTV-YY-S--EG-------V-VYGLIRTNICAEPGDS-G-
GPLYV-A---------STGVIIGILSGGS--------GNCTS---------------GGTTFYQPIAEILAAYGVTIP
>XX00000144 WP_018349638 21 49 186 MRLS-----------------------------------------
--------------------------------------------------------------------------------------------------------------------------------
--------------------------------------------------------------------------------------------------------------------------------
-----------------------------------------------------------------------------------RL--AAAVILA-ATALLVPGE------------
-------------------------------------------------------------------RP------VP-GTAWWTDPATGQLTAAVDSTV-T----
GP--A-LAR-VEAT-------------A-ARV---------ER---------Y-----P-GRF--T-E-HLSGGDT-FL-----N----G----
SYRCTVGFNVVDNLG-V--LY-FLTAAHCVGA-VG--S--PV---------------PNIGVVTGR-NTTYDYAIVKYTN--P-
N-IP-KPGN--VTLH-NG-SFQDIPPASNGLVGQAVRRSGGTTGVRSGTITAVNVTV-NY-P--SG-------T-
VYGLIRTNICTEPGDS-G-APLFS-G---------T--KAIGLASGGS--------GNCTS---------------
GGISFYSPVTRPLSAYGVNVY
>XX00000027 WP_027343883 26 53 190 MRRTL--------------------------------------
--------------------------------------------------------------------------------------------------------------------------------
--------------------------------------------------------------------------------------------------------------------------------
--------------------------------------------------------------S---A--VAAAAA-L----A-VTV--LTGPPAH-AA-DP-RAV-
------------------------------------------------------------------------------LA------TP-
GVAWILDPATGRTEVTVDSTV-P----AS--A-QAV-LRSA---------------YAV---------RH---------D-----P-GRL--

R-I-LIAGGDG-VY-----A----T-----GRCSLGANVHAG-T-T--YY-FVTSGHCTGT-GT--T--FY-A-----DAA-H-R--
TVLGTRTGTSFPVNDYGIVRYTG--T---VA-HPSA--VDTY-PG--
LLPIKGVALAYVGMTVCRSGVTTGVRCGIVTALNATV-AY-A--EG-------V-VTGLIRTNICAEPGDS-G-
GPLYD-P---------ATGRILGVLSGGT--------GDCTT--------------GGTTYYQPIGEILSAYGVTLP
>XX00000210  WP_051799105  13  56  196  MLT-----------------------------------------
--------------------------------------------------------------------------------------------------------------------------
--------------------------------------------------------------------------------------------------------------------------
------------------------------------------------------------------------------LV--TAPAPAF-AGPAAGPDP----------
-----------------------------------------------------------------ET------LP-GTAWWTDPATGTRTVSADDTV-A----
PR--D-LAR-LSRT-------------G-ATV---------VR---------E-----P-GTR--T-K-RLAGGDS-IY-----P---TG----
GYRCTIGFNARSTSK-E--YY-FITSGHCVGP-VG--S--TV-R-----IIA-N-G--TIIGTVVQR-LEPRDFALVRYVS--T-
VTVP-HPSA--VNRY-DG-TLQPITTFGTATVGQQVQRAGSTTGLWNGTVTAVNVTV-NY-A--DG-------
TRISGLIRTNLCSEPGDS-G-GPLFS-T---------T--IGLGLASGGS--------GNCTS--------------
GGVTYFTPVTPLAAAWSVGPF
>XX00000368  WP_015655451  44  60  187  MSVRRT------------------------------------
--------------------------------------------------------------------------------------------------------------------------
----------------S-------L-GR--FR--SAAC--------------V-LSLV-V-GA-----V-L-----G-------LSG-TG-S--A-V---
GPVAGPAR-AASAVEA----------------------------------------------------------------------------------------------
-------------------------------------------------------------------------------------------LG------IA-
GTAWGVDQRTGTLRVDVDSSV-S----AG--D-LAR-LRQVAE-RS----D-V-R-VVL---------QR---------E-----E-
GRL--R-T-LVSGGDG-VF-----A----A----GLRCSAGVNVRSG-T-T--YY-FVTAGHCTDA-AS--T--WY-T-----
TSA-Q-T--TSIGPTTSSSFPGNDFGVVRYAN--A-A-VP-HPGT--I-------
GTVDITGTATAYVGQSVCRRGSTTGVRCGVVTGLNATV-NY-G--GG-------SIVYGLIRTNICAEPGDS-G-
GPLYA-G---------D--KIIGILSGGS--------GNCTT--------------GGTTYYQPIQEVLSSQGLSVY
>XX00000373  WP_033367693  23  62  192  MRRTL--------------------------------------
--------------------------------------------------------------------------------------------------------------------------
--------------------------------------------------------------------------------------------------------------------------
--------------------------------------------------LGVL--AAAL---L--PATDAA-A--ASG-----------TATAAPV-RAF-
--------------------------------------------------------------------------AA------TP-GTAWWHD---
GRLTVSVDDSV-R----PA--V-AAG-LTRAVT-R-------A-G-GTV---------LR---------E-----P-GTL--A-P-
HIAGGDT-FN-----A----GT--LGGRCLIGFNARAG-A-T--YF-FLAAAHCVPA-VG--T--VY-A-----GTG-T-G--
TVLGVV-AARDPSYDSALVRYTN--A-T-IA-KPSA--VDLY-AG-GLQPITSFGAGTVGQAVRRTGP-
NGVRTGTITALNVTI-NY-A--GG-------T-VYNMIRTTVCSEPGES-G-GPLFA-G---------T--VGIGMNWGGS--
------GNCAT--------------GGVSYYSSARRAAVAYGVAPY
>XX00000013  WP_007461051  24  68  191  MRRSPH------------------------------------
--------------------------------------------------------------------------------------------------------------------------
--------------------------------------------------------------------------------------------------------------------------
-----------------------------------------------------VALVLTAL---L--LVAPGA-A----Q-AAV--PDRAAAD-PA-AV-
LSE---------------------------------------------------------------------------VR------TP-
GAAWGLDPATGRMTVTVDDTV-T----AA--D-LAR-LRAAAD-RA----G-----ALL---------RR---------E-----P-
GML--R-P-LIAAGQG-IY-----G----G----GSRCSLGANVRSG-S-T--YY-VVTAGHCTAA-VS--A--WY-A-----
DSA-Q-T--TVLGTRTATSYPGNDYGLIRYTG--R---IA-HPSA--VYTY-PG--
LLTIYGAGTAYVGQAVCRSGTTTGVRCGSVTGLNQTV-NY-A--TG-------V-IYGLIRTNICAEAGDS-G-
GPLYV-A---------STGTILGILSGGT--------GNCTA--------------GGTTYYQPIGEVLAAYGLSLP

>XX00000048  WP_025354457  31  70  178  MVTVGS-----------------------------------
----------------------------------------------------------------------------------------------------------------------
----------------------------------------------------------------------------------------------------------------------
-----------------------------ELRRLG------------S-ATLLAGVLTAMA----------APGA-Q--A-----A--PMAALVT-
AQATL-DAV-------------------------------------------------------------------------TG------VP-
NTAWGVDESSHQLVVTISDQA-K----GP--G-MGR-LLDLVR-GM----G-P-L-ARV---------RH---------T-----A-
RPL--S-P-QLLGGQE-IR-----T----D----TSICSAGFNVTKG-G-Q--LY-VLTAGHCTQG-GG--D--WQ-G---------
------LGPTAGSVFPGSDHGLIRNDT--G-D----GPGE--VDRY-DG-
GTQRISTVGNARVGEQACKSGRTTGLTCGQVTGLNRTV-NY-G--NG------QV-VRGLIEARVHCEGGDS-G-
GALFD-G---------G--TALGTVSGGD---------------------------SSTTYFQPVGPALSAYGVTLA
>XX00000064  WP_051844355  36  70  186  MLTPE--------------------------------------
----------------------------------------------------------------------------------------------------------------------
----------------EAVLRFK-R-TT--FA--TAIA--------------S-ALIA-A-GA-----L-A-----GP-----------------
AQAAPSPAQQAKS-AASAVDA-------------------------------------------------------------------------
-----------------------------------------------------------------------------------------TG------IE-
GISWGYDKQ-GRLFVTADSTV-S----RA--D-LAT-LKQTAA-RY----G-D-A-VRV---------ER---------T-----S-
GTF--S-P-KVGPGDA-IW-----G----V----GYRCSLGFNVVKN-G-T--YY-FLTAGHCAKD-VS--T--WY-A-----
DQN-R-N--TLIGSNVGYSFPGDDYALVRYDN--T-S-LS-HSGG--------------
YTAADAYVGETVTRDGSTTGVHSGKVTALNATV-RY-S--KG------GGTVYGLIQTTVCAEGGDS-G-GPLYD-
G---------N--KAIGLTSGGS--------GDCTS---------------GGTTFFQPVTEALSVYGVSLL
>XX00000105  WP_051897389  29  70  193  MRN-----------------------------------------
----------------------------------------------------------------------------------------------------------------------
----------------------------------------------------------------------------------------------------------------------
-----------------------------------------AH-RKITRKKAIIAAAGVALA--AAAAFA-I--PQA------SAHNSRP-
AVPSV-DQM-------------------------------------------------------------------------RG------MV-
GTAWATDPETGQTLITADSTV-N----SD--Q-WAS-LLT----AR----S-D-K-VQV---------QR---------A-----P-
GQF--R-L-FAEGGDA-IF-----A----G----QGRCSLGFNVTMSNG-A--PG-ILTAGHCTAA-GN--Q--WS-L-----
TQ--G-G--RPVATVQESVFPGNDFALLSYNN--A-N-TN-APSA--VDTG-NG--
TVQIQRAANPRVNQNVLRMGSTTGLNDGRVTGLNATV-NY-P--EG-------R-VTGLIQTTVCAEGGDS-G-
GPLFTQD---------G--TALGLTSGGA--------GDCTQ---------------GGVTFFQPVTEALQAFGAQIG
>XX00000366  WP_030952243  35  70  187  MTARRT------------------------------------
----------------------------------------------------------------------------------------------------------------------
----------------TK-------G-RS--QL--IAYV---------------L-SLLG-V-AA-----L-T-----V-------SGT-SA-T--A-A---
PATGPAPG-VASAVAA------------------------------------------------------------------------------
-------------------------------------------------------------------------------------LG------ID-
GTAWTVDERSGRLRVLADSTV-S----DT--E-LAR-LRRTAA-RF----D-G-G-LTL---------ER---------L-----D-
GRL--R-T-LLSGGDG-IY-----A----A----GRRCTAGVNVQSG-S-T--YY-FVTAGHCTEG-LP--T--WY-T-----
GAA-L-D--TAVGPTTGSSFPGNDFGVVRYAN--P-D-VP-HPGT--V-------
GTVDVTGTASAHVGQRVCTRGATTGVRCGVVTALNATV-NY-G--GG-------DIVTGLIQTNICAEPGDS-G-
GPLYA-G---------D--KVIGILSGGS--------GNCTT---------------GGTTYYQPIQEALSAYGLSVY
>XX00000331  WP_040247686  40  72  192  MKLRRS------------------------------------
----------------------------------------------------------------------------------------------------------------------
----------------SSPRRSR-T-LR--RQ--ALAA--------------A-GLLT-A-AF-----F-L-----TP-----------G-TGAS-
AAQEPTGAAAAH-AAEAVLD--------------------------------------------------------------------------

------------------------------------------------------------------------------------------------AD------VP-
GTAWSLDPATGTLALTADSTV-D----AA--E-LAR-LKRATA-PY----A-Q-A-VSL---------DR---------V-----D-
GTL--R-P-HVSGGSS-VY-----G----A----GARCTAGFNVHNG-S-D--YY-FLTAGHCTEA-AQ--D--WY-A-----
DPA-L-T--VYIGSTAGSSFPGNDYGLVRYQN--P-A-VP-HPSD--VRNA-TG-
VVTQITGAGQATVGQQVCTTDRTTGVHCGTVTGLNATV-NY-G--NG-------QVVSGLAQTTLCSEPGSS-G-
SPVFS-G---------T--IALGLISGGS--------GNCTS---------------GGTSFYQPVPEVLSAYGLTI-
>XX00000367 WP_054225693 32 72 187 MRIKSI-------------------------------------
-------------------------------------------------------------------------------------------------------------------------
---------------T---------RT--RL--LAVA---------------A-GLAA-V-AG-----L-A-----AP-------T----AA--S-
ADTGFSAARLAA-AGASVQR---------------------------------------------------------------------------------------
------------------------------------------------------------------------------------------------AD------VA-
GTAWYTDPASGTLVVTADSTV-T----AA--D-LAR-IRTEAG-AD----A-A-A-LRI---------ER---------T-----P-GKL-
-R-K-LISGGDA-IY-----T----S----GWRCSLGFNVRSG-N-T--YY-VLTAGHCTDG-AG--T--WW-T-----SSA-H-
T--TTIGATVGSSFPNNDYGLVRYDN--T-S-LA-HAGT--V-------
GSQDITSAANPTVGMSVTRRGSTTGTHGGSVTGLNATV-NY-G--GG-------DIVYGMIRTNVCAEPGDS-G-
GPLYS-G---------T--KAIGLTSGGS--------GDCTS---------------GGTTFFQPVVEALNAYGVSVY
>XX00000083 WP_015621935 24 73 191 MRRRL-------------------------------------
-------------------------------------------------------------------------------------------------------------------------
-------------------------------------------------------------------------------------------------------------------------
--------------------------------------------------IAGP--PTAG---G--VSAAGA-R--PGT------------AAVAGRP-
AVA--------------------------------------------------------------------------GG------QP-
GTALFRDPVTGRMTLVADDTV-V----HP--V-A------------------D-G-VDV---------RR---------E-----P-GRL--R-
L-LIAGGEG-VY-----A----G----AARCTLGANVRQG-T-T--AF-FISSGHCAGT-GT--T--WF-A-----SPG-Q-G--
TLLGTRTGLSFPGNDYAIVRYTN--P-A-VA-HPSA--VQT--PG-
GLLPLTGWAGPVVGQAVCRSGATTGVRCGTVTALNATV-TY-A--EG-------T-VTGLIRTNICAEPGDS-G-
GPLYT-T--------AG--VLIGILSGGS--------GTCTT---------------GGTTYFQPVGEILSAYGVTVP
>XX00000176 WP_051787611 53 73 188 MPTS---------------------------------------
-------------------------------------------------------------------------------------------------------------------------
-----------------------------------------------------------------------------------------------------------------------
A-GGLL-T-----LP------------------------------------------------------TAPA-T--ARN-SPH--RPVDVAE-ARAHV-
E-A---------------------------------------------------------------------------LA------IG-
GTSWSVDPDSHTLVVDADQSV-T----DA--E-WAE-LISATA-RY----G-G-A-VEV---------QR---------V-----R-
GTF--T-R-TISGGNA-VY-----A----S----WKRCSVGFNVRNAAG-T--HY-LLTAGHCARG-TT--N--WY-T-----
NSA-R-S--TWIGPVSGYSFPGNDYALVRYAN--T-A-LS-HPGM--V-------
GAVDITRAGNAYVGQSVTRRGSTTGSRSGKVTALNVTV-NY-G--DG-------LVVYGLVKTNICVEPGDS-G-
GPLHS-G---------G--TALGLTSGGR--------GNCTT---------------GGVSFFQPVTEALSAYRVNVY
>XX00000315 WP_027770685 45 73 196 MSTRRT-----------------------------------
-------------------------------------------------------------------------------------------------------------------------
-------------------------------------------------------------------------------------------------------------------------
-----------------------------NPLRRTVRRLRRPATALT-ALAVLALPAVLGLG---G--PQA--A-Q--AQQ-RTF--
SATELAA-AGDAV-L-A-------------------------------------------------------------------------------AD------IA-
GTAWGVDRATGTVLVTVDERV-S----AA--E-IAA-LRRSAG-AQ----A-D-A-LTF---------ER---------T-----T-
GTF--R-P-HLQGGEP-VW-----S----S----AGRCTVGFNVRAG-S-A--DY-FVTAGHCTLG-SP--T--WY-T-----
NSS-F-T--TVVGPTAGSSFPGNDYGIVRYTN--T-A-VP-RPGT--VN-C-DG-

TIIDITGPANAGVGQTIWMAGSTSGCHSGAVTGLNATV-NY-G--DG--------IVSGLIRTNLCSEPGDS-G-
APVFI-R---TGDGTTG--LAVGVLSGGS--------GNCAA--------------GGTSFVQPINEILAAYGAVLT
>XX00000323 WP_059009593 40 73 193 MSLKRT---------------------------------------
-----------------------------------------------------------------------------------------------------------------------------------
----------------NPHRRMS-R-RL--RL--TAAA--------------S-GLLA-A-VA-----F-F-----VP-------SP-AS-A--A--
PVTFSTAQLTS-ASGAVLA----------------------------------------------------------------------------------------------
----------------------------------------------------------------------------------------------------AD------VA-
GTAWTVDKSTNRVVVTVDETV-S----SA--E-IAE-IRRSAG-AL----S-G-A-VTI---------EH---------T-----E-GQL-
-E-R-YIAGGDA-IY-----A----SL---GWRCSLGFNVQIG-S-A--YY-FLTAGHCTEG-FP--T--WY-T-----NSS-Q-T--
TLIGPTTGSSFPGNDYGIVRYDN--T-A-VS-HPGT--VNLY-NG-
TTRDITGAANPVNGQAVQRSGSTTGLRSGSVTGLNYTV-NY-G--GG-------DIVSGLTRTNVCAEPGDS-G-
GPFFS-G---------N--TALGLTSGGS--------GNCTT--------------GGTTYFQPVVEALNAYGAVVY
>XX00000356 WP_031511140 41 73 184 MRIKRT---------------------------------------
-----------------------------------------------------------------------------------------------------------------------------------
--------------SPHTGPARR-G-RI--RL--LAAA--------------S-GIVA-A-AA-----L-T-----IP-------AS-AS-A--E--
PAPSPAAQVSM-ANKAMKT------------------------------------------------------------------------------------------------
-------------------------------------------------------------------------------------------------------AD------VA-
GTAWYTDQETNTLVVTVDSTV-S----KA--E-IAK-IKREAG-AG----A-E-A-IRI---------ER---------T-----P-GKF--
N-K-LISGGDA-IY-----S----G----GGRCSLGFNVRSSSG-V--DY-FLTAGHCTNG-TS--T--WT-N---------G-S--
TTLGTTAGSSFPNNDYGIVRYTS--S-I--S-RPGQ--V-------
GNQDITRAANPSVGQTVYRRGSTTGTHSGRVTGLNATV-NY-G--GG-------DIVYGMIQTNVCAEPGDS-G-
GPLYY-G---------T--TAYGLTSGGS--------GNCRT--------------GGTTFFQPVVEALNAYGVSVY
>XX00000033 WP_049577616 39 74 193 MRVKRT---------------------------------------
-----------------------------------------------------------------------------------------------------------------------------------
-----------------NPRGRVA-R-SL--RA--AAVV--------------T-GLVA-V-AA-----F-L-----TP-------G---TAL--
ADEAAAFGTAQLEA-AGEAVDA-------------------------------------------------------------------------------------------
-------------------------------------------------------------------------------------------------------------AD------VG-
GTAWSVDEKAGTVLVLADESV-S----EA--E-IAE-ITASAG-DL----A-G-A-LSI---------ER---------T-----P-GTF--
E-T-YLQGGDP-IY-----A----DI---GWSCSLGFNVRAG-S-T--FY-FLTAGHCTED-PP--L--WH-D---------G-S--
AWIGFTAASNFPGDDFGLVQYDP--Q-P-AS-VPSS--VS------
GGGAITGFGNPFVGQSACRSGLTTGVRCGSVTGLNWTV-DY-G--GG-------DVVYGMIQTNICAEPGDS-G-
GPLWS-G---------T--TGLGLTSGGS--------GNCSV--------------GGTTFFQPVAEAAAYYGVGIP
>XX00000085 WP_046502477 38 74 182 MRTKRT---------------------------------------
-----------------------------------------------------------------------------------------------------------------------------------
-----------------NRRAGGL-R-RT--RT--LAVV--------------T-GLLA-A-VA-----L-A-----IP-------AA----S--A-
HTAAGTGADRSAAVGDAVLA---------------------------------------------------------------------------------------------
------------------------------------------------------------------------------------------------------------AG------VA-
GTAWHTDPATGEVVVTADRTV-S----AG--E-IAA-IERAAG-PH----A-G-A-LRF---------ER---------T-----V-
GTL--A-P-HLSGGDA-VY-----S----G----GARCAAGFNVVSG-S-T--YY-FVTAGHCTSG-AS--T--WY-A-----
DPS-G-T--TLLGTTAGTTFPSRDFGLVRYTN--P-S-VP-VPGT--V-------
GTVDITGAGNATVGMSVTFHGPVGGVRSGTVISLNNTV-NY-G--GG-------QTVTGLIRTNICTYPGES-G-
APLYS-G---------G--IAIGVLSGGS--------G------------------CTSYFQPIIPVLGAYGVSVY
>XX00000113 WP_031231424 31 74 185 MNRTP----------------------------------------
-----------------------------------------------------------------------------------------------------------------------------------

----------------------A-P-RA--RL--IAVA---------------S-GVLA-A-TA-----L-T-----AP-------GA----T--
ADDAVRPTAAQLTA-AHNALAR-------------------------------------------------------------------------------------
---------------------------------------------------------------------------------------------------AD------VT-
GSAWYTDAATGTVVVTVDSTV-D----AG--E-LAA-LTKAAT-TP----A-G-A-PEI---------NR---------T-----T-
GRF--T-K-HIAGGEY-IL-----L---PG----GIRCLIGFNVQDSAG-V--KY-ALTAGHCTDT-GD--V-------------------
TSLGTTVNSSFPGNDYALIRYSN--Q-S-A--AEGS--VYLQ-NG-
TYQDITSAGSPFVGQRITTSAPTTGVRSGTVTGLNATV-NY-G--AD-------GIVYGLIQSNLCSEPGSS-G-
GPVYS-G---------T--QAMGLISGGS--------GNCST---------------GGTTFAQPVVEPLSAYGVSVF
>XX00000174 WP_052850046 39 74 193 MGLKRT-------------------------------------
----------------------------------------------------------------------------------------------------------------
-----------------NPHGRMS-R-RL--RL--AAAA---------------S-GLLA-A-AA-----F-V-----MP-------ST-AS-A--E-
PVSTFSSSQLSA-ASAAVLD------------------------------------------------------------------------------------
-------------------------------------------------------------------------------------------AD------VA-
GTAWTVDKQSGRVVVTADETV-S----NA--E-IAE-LRSSAG-AL----S-G-A-LTV---------ER---------T-----E-
GTL--E-R-YLAGGDP-IY-----A----SL---GWRCSAGFNVSIG-G-A--AY-FLTAGHCTDG-YP--G--WY-T-----
NPS-L-T--TYIGPTVGSSFPGNDYGLVRYDN--A-S-VP-RPGV--VNLY-NG-
TTVDITGAANPSVGQAAQRSGSTTGLRSGSITGLNYTV-NY-G--GG-------DIVSGLIRTNICAEPGDS-G-
GPLFS-G---------R--TALGLTSGGN--------GNCSV---------------GGTTYFQPVVEALSAYGATIL
>XX00000208 WP_006381951 38 74 190 MRIKRT-------------------------------------
----------------------------------------------------------------------------------------------------------------
----------------TPRSGIT-R-RT--RL--IAVS---------------T-GLVA-A-AA-----I-A-----IP-------NA-TA----A-
DVTTFSTAQLTK-ASGSVLK--------------------------------------------------------------------------------------
-------------------------------------------------------------------------------------------AD------VP-
GTAWAVDAKTNRVVVTADSTV-S----QA--E-IAK-IKRQAG-ST----A-D-A-ITV---------KR---------T-----L-
GKF--S-K-LISGGDA-IY-----A----S----SWRCSLGFNVRSG-S-T--YY-FLTAGHCTDG-AG--N--WW-S-----
NSA-R-T--TLLGATSGSSFPTNDYGIVKYAS--S-Y-PV-SSGT--V-------
GSVDITSAATPSVNTTVYRRGSTTGIHSGRVTALNATV-NY-G--GG-------DVVYGMIQTTVCAEPGDS-G-
GPLYS-N--------SG--VAYGLTSGGS--------GNCSS---------------GGTTFFQPVTEALSAYGVSVF
>XX00000314 WP_053723396 40 74 187 MRIKRT-------------------------------------
----------------------------------------------------------------------------------------------------------------
--------------TPTPPTGLA-R-RT--RL--AAVA---------------T-GLLA-A-VA-----I-A-----VP-------NA-SA----T-
PARTFGVAALTA-ASDAVRG-----------------------------------------------------------------------------------
---------------------------------------------------------------------------------------------AD------VA-
GTAWSVDSATGEVVVTADSTV-S----AA--G-LAK-IKREAG-AN----A-G-A-LRI---------ER---------T-----P-
GKF--T-K-LLSGGDA-IY-----T----T----SWRCSLGFNVKKG-S-T--YY-FLTAGHCTQG-KP--N--YY-T-----NSS-
H-S--TSIGPTVGTSFPGNDYGLVQYTN--T-S-LA-HPSA--V-------
GSQHISSAANPTVGQTVTRRGSTTGVHSGKVTALNQTV-NY-G--NG-------DIVSGLIRTTVCAEPGDS-G-
GPLYA-G---------S--TALGLTSGGS--------GDCTS---------------GGTTFFQPVVEAMNAYGVTLF
>XX00000317 AJF64415 31 74 188 MTT---------------------------------------------------
----------------------------------------------------------------------------------------------------------------
----------PT-K-KF--RL--LALT---------------A-GLVAAA-AT-----L-G-----VP-------TA-S--A--D-
SAQTFSATQLSA-AGDAVLA------------------------------------------------------------------------------------
---------------------------------------------------------------------------------------------AD------VA-
GTAWRVDTATGQVVVTADSTV-S----QA--E-IAR-IKHKAG-AN----A-G-A-LRI---------ER---------T-----P-

GTF--N-K-LISGGDA-IY-----A----S----SWRCSLGFNVKDSAG-N--YY-FLTAGHCTDG-AG--T--WW-S-----
NSS-H-S--TVLGSTAGSSFPTNDYGIVRYTN--T-S-VT-KSGT--V-------
GSVDITSAANATVGMSVTRRGSTTGIHSGTVTGLNATV-NY-G--GG-------DIVYGMIQTNVCAEPGDS-G-
GPLYS-G---------S--KAIGLTSGGS--------GNCSS---------------GGTTFFQPVTEALSAYGVNVY
>XX00000339 WP_005319313 39 74 187 MRIKRT-------------------------------------
-----------------------------------------------------------------------------------------------------------------
---------------IPRTRTTA-R-RT--RL--LAVA---------------T-GLTA-A-AA-----L-A-----VP-------TA-SA----E-
ETQTFSTHQLTA-ASDAVLQ-----------------------------------------------------------------------------------------
---------------------------------------------------------------------------------------------------AD------VP-
GTAWHVDQATGTLVVTADSTV-S----RA--E-IAK-IKREAG-TN----A-G-A-LRI---------ER---------T-----P-
GTF--S-K-LISGGDA-IY-----A----S----SWRCSLGFNVRSG-S-T--YY-FLTAGHCTDG-AG--T--WY-T-----NSS-
R-T--TSIGPTTGSSFPGNDYGIVRYSN--T-S-LS-HPGT--V-------
GGTDITSAANATVGMSVTRRGSTTGTHSGTVTGLNATV-NY-G--GG-------DIVYGMIRTNVCAEPGDS-G-
GPLYS-G---------S--RAIGLTSGGS--------GNCST---------------GGTTFFQPVTEALSRYGVSVY
>XX00000342 WP_030356095 38 74 187 MRIKRT-------------------------------------
-----------------------------------------------------------------------------------------------------------------
----------------TPRSGIA-R-HS--RT--VAIA---------------T-GLVA-A-AA-----L-A-----VP-------AA-SA----D-
EARTYSAADLTA-ASDAVLE------------------------------------------------------------------------------------------
---------------------------------------------------------------------------------------------------AD------VA-
GTAWNVDPKTKQVVVTADSTV-S----KA--E-IAK-LKQAAG-DK----A-G-A-LKI---------ER---------T-----P-
GKF--Q-K-YISGGDA-TY-----A----S----SWRCSLGFNVRSG-S-T--YY-FLTAGHCTDG-AG--T--WW-A-----
NSA-K-T--TVLGTTHGSSFPTNDYGIVRYTN--S-S-IT-KSGT--V-------
GSQDITRAADPTVNQSVTRRGSTTGTHSGRVTGLNATV-NY-G--NG-------DIVYGMIRTTVCAEPGDS-G-
GPLYA-G---------S--TALGLTSGGS--------GNCSS---------------GGTTFFQPVTEALRAYNVSVY
>XX00000343 WP_030578655 38 74 189 MRTQRT-----------------------------------
-----------------------------------------------------------------------------------------------------------------
-----------------TPRSGVA-R-RT--RL--IAVA---------------A-GLVA-A-AA-----V-A-----VP-------NA-NA----S-
DAQTFSTTELKQ-ASSSILQ--------------------------------------------------------------------------------------
--------------------------------------------------------------------------------------------------AD------VP-
GTAWAVDQKTGKVLLTVDSTV-S----EA--E-LAK-IQKAAG-DN----A-G-A-LEV---------KH---------T-----P-
GKF--S-K-LIQGGDA-IY-----S----G----SGRCSLGFNVRSTSG-V--DY-FLTAGHCTSG-SG--T--WY-G-----NSS-
R-T--TVLGPAAGTSFPGNDYGIVRYSN--T-S-IA-KPGT--A-------
NGVDITRAATPSVGTTVIRDGSTTGTHSGRVTALNATV-NY-G--SG-------QVVSGLIQTTVCAEPGDS-G-
GPLYG-S--------NG--TAYGLTSGGS--------GNCSS---------------GGTTFFQPVTEALSAYGVNLT
>XX00000345 WP_033248983 39 74 187 MRIKRS-------------------------------------
-----------------------------------------------------------------------------------------------------------------
---------------TPRNGGIT-R-RG--RI--TAVA---------------A-GFVA-V-AA-----L-A-----VP-------AA-DA----D-
QSGTYSTSQLTA-AGAAVLR-----------------------------------------------------------------------------------------
--------------------------------------------------------------------------------------------------AD------VP-
GTAWNADPATGRLLVTVDSTV-S----QT--E-IKK-IRKAAG-AD----A-G-A-LRI---------ER---------T-----P-GTL-
-S-K-LISGGDP-VY-----A----S----GWRCSLGFNVRSG-T-T--YY-FLTAGHCTEG-AD--T--WW-A-----DAA-H-
T--TVLGTTAGSSFPGDDYGIVRYTN--A-T-VA-KPGT--V-------
GSQDITGAADATIGMSVTRRGSTTGLHSGTVTGLNATV-NY-G--GG-------DIVYGMIRTNVCAEPGDS-G-
GPLYS-G---------T--RAIGLTSGGS--------GNCTS--------------GGTTFFQPVTEALNAYGVSVF

>XX00000347 WP_031075620 38 74 188 MRIKRT-------------------------------------
---------------------------------------------------------------------------------------------------------------
---------------TPRSGIA-R-RT--RL--LAVS--------------V-GLVS-A-AA-----L-A-----VP-------AA-NA----D-
SPRTYSAADLSA-AGSAVLA-------------------------------------------------------------------------------------
---------------------------------------------------------------------------------------AD------VP-
GTAWHVDPGSNTLVVTADSTV-S----RD--E-IAR-IKREAG-TH----A-G-A-LRI---------ER---------T-----P-GKF-
-T-K-LISGGDA-VY-----A----S----SWRCSLGFNVRDSAG-A--YY-FLTAGHCTDG-AG--T--WY-A-----NSG-R-
T--TVLGSTSGSSFPNNDYGIVRYTN--S-S-IT-KSGT--V-------
GNVDITSAANATVGMSVTRRGSTTGTHSGSVTGLNATV-NY-G--GG-------DIVYGMIRTNVCAEPGDS-G-
GPLYS-G---------S--RAIGLTSGGS--------GNCSS--------------GGTTFFQPVTEALNAYRVSVY
>XX00000348 AAM96214 38 74 188 MRIKRT---------------------------------------------
---------------------------------------------------------------------------------------------------------------
------TPTSGIA-R-RT--RL--IAVA--------------T-GFVA-A-AA-----I-A-----VP-------NA-NA----S-
DVHTFSANQLTK-ASDSVLN-------------------------------------------------------------------------------------
-----------------------------------------------------------------------------------------SD------VA-
GTAWAVDPATNRVVVTVDSTV-S----KA--E-IAK-IKKTAG-SN----A-G-A-LTI---------KH---------T-----P-
GKF--N-K-LISGGDA-IY-----A----S----SWRCSLGFNVQDSSG-N--YY-FLTAGHCTDG-AG--T--WW-S-----
NSS-H-T--TTLGTTAGSSFPGNDYGIVRYTN--S-S-VA-KSGA--V-------
GSQDITSAATPSVGTTVYRRGSTTGTHSGRVTALNATV-NY-G--NG-------EIVYGLIQTTVCAEPGDS-G-
GPLYG-G---------S--TAYGLTSGGS--------GNCTS--------------GGTTFFQPVTEALSAYGVHVY
>XX00000349 WP_009314945 38 74 188 MRITRT-------------------------------------
---------------------------------------------------------------------------------------------------------------
---------------TPRSGIA-R-RT--RL--IAVA--------------A-GLAA-A-AA-----V-T-----VP-------TA-NA----A-
GTQTFSAAELKS-ASSSVLK-------------------------------------------------------------------------------------
-------------------------------------------------------------------------------------AD------VP-
GTAWAIDPKTDKVLLTIDSTV-S----DA--E-LAK-IKEAAG-DN----A-D-A-LTV---------KR---------T-----P-GKF-
-N-K-LIKGGDA-IY-----A----S----SWRCSLGFNVRSG-S-T--YY-FLTAGHCTDG-AG--T--WY-S-----NSG-R-T--
TVLGPTTGSSFPTNDYGIVRYSN--T-S-IA-KDGT--A-------
GSVDITSAATPSVGTNVIRTGSTTGTRTGRVTALNATV-NY-G--GG-------DIVYGMIQTTVCAEPGDS-G-
GPLYG-S--------NG--VAYGLTSGGS--------GNCTS--------------GGTTFFQPVTEALSAYGVSVF
>XX00000350 WP_028815637 38 74 188 MRIKRT-------------------------------------
---------------------------------------------------------------------------------------------------------------
---------------TPRSGIA-R-RS--RL--IAVT--------------A-GLVA-T-AA-----L-T-----VP-------SA-SA----D-
TASTYSANQLTS-ASDAVLD-------------------------------------------------------------------------------------
---------------------------------------------------------------------------------------AD------VA-
GTAWHIDKAKNTLVVTADSTV-S----AA--E-IAK-IKREAG-TN----A-G-A-IRI---------ER---------T-----P-GKL--
S-K-LLSGGDA-IY-----A----T----SWRCSAGFNVRNSAG-T--YF-FVTAGHCTDG-NP--P--WY-T-----NSS-R-T-
-TSIGPTAGSSFPGNDYGLVRYAN--T-S-LA-HPGT--V-------
GNQDITSAANATVGMSVTRRGSTTGTHSGSVTGLNATV-NY-G--GG-------DVVYGMIRTNVCAEPGDS-G-
GPLYS-G---------T--RAIGLTSGGS--------GNCSS--------------GGTTFFQPVTEALSAYRVSVY
>XX00000352 WP_030583931 38 74 187 MRIKRT-------------------------------------
---------------------------------------------------------------------------------------------------------------
---------------NNRSNAA-R-RV--RT--TAVV--------------A-GLAA-V-AA-----M-A-----IP-------TA-NA----E-
TPRTFSANQLTA-ASDAVLG-------------------------------------------------------------------------------------

-----------------------------------------------------------------------------------------------AD------IA-
GTAWNIDPQSKRLVVTVDSTV-S----KA--E-INQ-IKKSAG-AN----A-D-A-LRI---------ER---------T-----P-GKF-
-T-K-LISGGDA-IY-----S----S----TGRCSLGFNVRSG-S-T--YY-FLTAGHCTDG-AS--T--WW-A-----NSA-R-T--
TVLGTTAGSSFPNNDYGIVRYTN--T-T-IP-KDGT--V-------
GGQDITSAANATVGMSVTRRGSTTGTHSGSVTGLNATV-NY-G--GG-------DVVYGMIRTNVCAEPGDS-G-
GPLYS-G---------T--RAIGLTSGGS--------GNCSF---------------GGTTFFQPVTEALSAYGVSVY
>XX00000353 WP_028427405 40 74 185 MKTRRT------------------------------------
------------------------------------------------------------------------------------------------------------------
----------------TTPLGGMS-K-RA--RL--IATA---------------S-GLAV-A-AA-----L-M-----SP-------AV-AQ-A--E-
EPSTFSKAQLNQ-ASDAVLE------------------------------------------------------------------------------------
-----------------------------------------------------------------------------------------------AD------VA-
GTAWRVDEKTGTLVVKADRTV-S----NA--E-IAE-IKKAAG-SN----A-G-A-VEI---------ER---------T-----P-
GKF--T-K-YISGGDA-IT-----T----G----SGRCSLGFNVRSSSG-A--SY-FLTAGHCTDG-FP--T--WY-T-----GS----
G--AYLGPTAASSFPGNDYGIVQHSN--S-Q--S-KPGT--V-------
GGQDITTAANPSVGQSVTRRGSTTGTHGGQVTGLNATV-NY-G--GG-------DVVYGMIQTNVCAEPGDS-G-
GPLYS-G---------T--TALGLTSGGS--------GNCTS---------------GGTTFFQPVVEALNAYGVSVY
>XX00000354 WP_030696597 38 74 187 MRIKRT-------------------------------------
------------------------------------------------------------------------------------------------------------------
----------------TPRSSVA-R-RT--RL--VALA---------------A-GLAA-V-GA-----L-A-----TP-------SA-HA----G-
ESATFSSAELTS-AAEAVLD--------------------------------------------------------------------------------------
----------------------------------------------------------------------------------------------TD------VA-
GTAWYVDKAAGKVVVTADSTV-S----KA--E-IAK-IQKAAG-GE----A-G-A-LEI---------NR---------T-----E-
GTF--S-P-LLSGGDA-IY-----S----S----SSRCSLGFNVRSG-S-T--YY-ALTAGHCTSG-TS--T--WY-T-----NSA-R-
T--TAFGTVAGSSFPNNDYGLIRYTN--S-S-VP-ASGT--V-------
GSVDITRAANPTVGQTVTRRGSTTGTHSGRVTGLNATV-NY-G--SG-------QVVYGMIQTNVCAEPGDS-G-
GPLYS-G---------S--TALGLTSGGS--------GNCSS---------------GGTTFFQPVVEALNAYGVSVF
>XX00000405 WP_030706074 38 74 187 MRIKRT-------------------------------------
------------------------------------------------------------------------------------------------------------------
----------------SNRSNAV-R-RV--RT--TAVL---------------A-GLAA-V-AA-----L-A-----VP-------TA-NA----E-
TPRTFSANQLTA-ASDAVLG------------------------------------------------------------------------------------
-----------------------------------------------------------------------------------------------AD------IA-
GTAWNIDPQSKRLVVTVDSTV-S----KA--E-INQ-IKKSAG-AN----A-D-A-LRI---------ER---------T-----P-GKF-
-T-K-LISGGDA-IY-----S----S----TGRCSLGFNVRSG-S-T--YY-FLTAGHCTDG-AT--T--WW-A-----NSA-R-T--
TVLGTTSGSSFPNNDYGIVRYTN--T-T-IP-KDGT--V-------
GGQDITSAANATVGMAVTRRGSTTGTHSGSVTALNATV-NY-G--GG-------DVVYGMIRTNVCAEPGDS-G-
GPLYS-G---------T--RAIGLTSGGS--------GNCSS---------------GGTTFFQPVTEALSAYGVSVY
>XX00000110 WP_014173668 45 75 191 MRIK---------------------------------------
------------------------------------------------------------------------------------------------------------------
----------RSGPASRPARAA-A-RL--RT--AALI---------------A-GLAA-V-FA-----L-L-----AP-------GA-AA----
ADPAQTYSASELKA-ASDAILA------------------------------------------------------------------------------
--------------------------------------------------------------------------------------------------AD------VG-
GTAWAVDEDSGSVRVDVDRTV-S----SS--E-LAT-LKQAAG-PR----S-G-A-LHI---------QR---------T-----Q-
GRI--T-Q-LLSGGDP-IY-----S----SN---GFRCSAGFNVHIG-T-S--YY-ILTAGHCTNG-YP--N--WS-A-----RL----
G--VPIGPTAYSSFPVNDYGYVSYTN--P-L-LP-HPSN--VYLY-NG-

TYRAITGAGNAFVGMLVQRSGSTTGLHGGTVTGLNYTV-NY-G--NG-------NVVYQMIRTNVCAEPGDS-G-
GPLFS-G---------N--TAIGLTSGGS--------GNCQT---------------GGVTYYQPVVEALNNHGAAIP
>XX00000137 WP_003957238 34 75 188 MKIHKV-------------------------------------
---------------------------------------------------------------------------------------------------------------------
--------------------SLS-Q-RA--RM--VAVV---------------A-GLVT-V-SA-----L-I-----AP-------TA-VA----
AESAPKPGAAQLAQ-VSDAVLD-----------------------------------------------------------------------------------
-------------------------------------------------------------------------------------------------AD------IS-
GTAWYTDAKSGKVVVTADSTV-S----AA--E-IDR-IKKQAG-AE----S-G-A-LKI---------NR---------T-----P-
GTF--S-K-MIAGGEA-IY-----T----A----GARCTLGFNVVRA-G-V--FY-ALTAGHCTNI-GP--S--WT-T-----AV---
----GPLGNRFASVFPGSDHGVIQHAI--P-A-N--ADGR--VYLH-NG-
TYRDMVNASTPLIGQFVERSGSTTGYHTGRVTGLNATV-NY-G--GG-------QLVYGLIQTNVCGQPGDS-G-
GPLFS-G---------N--TAHGLLSGGS--------GNCTV---------------GGTTFYEPVLRPLTQYGLTIF
>XX00000336 WP_055416160 38 75 188 MRIKRT-------------------------------------
---------------------------------------------------------------------------------------------------------------------
----------------TPRSGIT-R-RT--RL--IAVS---------------S-GLVA-A-AA-----I-A-----IP-------SA-NA-A--P-
TPATFSAAELNS-ASGAVLE---------------------------------------------------------------------------------------
-------------------------------------------------------------------------------------------AD------VP-
GTAWAVDSKTGRVLLTVDSTV-S----QA--E-IAK-IKKEAG-DK----A-G-A-FTI---------KR---------T-----P-GTF-
-N-K-LIQGGDA-IY-----A----S----SWRCSLGFNVRSSSG-A--EY-FLTAGHCTDG-AG--T--WY-S-----NSG-R-T-
-SVIGSTAGSSFPGNDYGIVRYSG--S-V--S-RPGT--A-------
NGVDITRAATPSVGTTVIRDGSTTGTHSGRVTALNATV-NY-G--GG-------DVVGGLIQTTVCAEPGDS-G-
GALYGSN---------G--TAYGLTSGGS--------GNCRS---------------GGTTFFQPVTEALNAYGVSVY
>XX00000338 WP_031064229 38 75 188 MRNKRI-------------------------------------
---------------------------------------------------------------------------------------------------------------------
----------------TPRSGVT-R-RT--RL--IAVA---------------S-GLLV-A-GA-----V-A-----VP-------TA-SA-D--G-
SVTTFSASQLSA-ASDSVRA---------------------------------------------------------------------------------------
-------------------------------------------------------------------------------------------AD------VA-
GTAWAVDKKTNALVVTVDKTV-S----QA--E-IAK-IKKEAG-VN----A-G-A-IRI---------ER---------T-----A-
GKF--Q-K-YISGGDA-IY-----A----T----SWRCSLGFNVRNSSG-T--YY-FLTAGHCTDG-NP--P--WY-T-----
NSS-R-T--TSIGPTAGSSFPNNDYGIVKYTN--T-S-VS-HAGT--V-------
GSQDITSAGNPTVGQSVTRRGSTTGIHSGTVTALNATV-NY-G--GG-------DIVYGMIQTTVCAEPGDS-G-
GPLYS-G---------T--KALGLTSGGS--------GDCTS---------------GGTTFFQPVTEALSAYGVSVY
>XX00000046 WP_003964236 38 76 183 MTFKRF-------------------------------------
---------------------------------------------------------------------------------------------------------------------
----------------SPLSSTS-R-YA--RL--LAVA---------------S-GLVA-A-AA-----L-A-----TP-------SA-VA-A--
PEAESKATVSQLAD-ASSAILA-----------------------------------------------------------------------------------
-----------------------------------------------------------------------------------------------AD------VA-
GTAWYTEASTGKIVLTADSTV-S----KA--E-LAK-VSNALA-GS----K-A-K-LTV---------KR---------A-----E-GKF-
-T-P-LIAGGEA-IT-----T----G----GSRCSLGFNVSVN-G-V--AH-ALTAGHCTNI-SA--S--W-----------------
SIGTRTGTSFPNNDYGIIRHSN--P-A-A--ADGR--VYLY-NG-
SYQDITTAGNAFVGQAVQRSGSTTGLRSGSVTGLNATV-NY-G--SS-------GIVYGMIQTNVCAEPGDS-G-
GSLFA-G---------S--TALGLTSGGS--------GNCRT---------------GGTTFYQPVTEALSAYGATVL
>XX00000111 WP_054243206 20 76 187 MRKLA--------------------------------------
---------------------------------------------------------------------------------------------------------------------

----------------------------------------------------------------------------------------------------------------------------
---------------------------------V-GAAGVVLA-------------ASGL---V--AQSIPS-E--------A--PDASVKD-ARKVL-
KQE----------------------------------------------------------------------------AA------IP-
GTSWSVDPETDKLVVTADKTV-K----GD--D-LDQ-LKDVVD-EL----G-D-N-VEL---------EQ---------S-----K-
TVI--Q-P-YAVGGDA-IY-----A---------SERCSAGLNVLMA-G-N--PF-LLTAGHCGKT-GS--L--WS-D-----KK---
-G--LTVGTMVESVFPVDDYALIKYED--S-T-T--PESA--VNLY-EG-VN-
KVFTVGTPTEGMQVERSGSTTGVQRGRITGLNATV-NY-G--NG------IS-VDGLIQTDVCAEPGDS-G-GPLFS-
G---------G--MALGLLSGGY--------GNCKT---------------GGATYYQSLNEVFSKYNLTLP
>XX00000121 WP_004949237 43 76 192 MRNKR--------------------------------------
----------------------------------------------------------------------------------------------------------------------------
----------------------------------------------------------------------------------------------------------------------------
L-AV-S-A-----LT-----------------------------------------------------ST-PAG-AAPGAP-EVP--SAARLAA-LSTAV-
G-E-------------------------------------------------------------------------------AD------VP-
GTAWAVDERSHEVVVTADATV-P----AE--G-VAR-IRRAAG-PD----T-D-A-IRV---------ER---------A-----P-
GAF--R-P-LIAGGDA-VH-----A----G----DWRCSAGFNVRKG-T-T--YY-FLTAGHCTQG-KP--A--YF-V-----
DPG-H-R--MSVGPTAGSTFPGADHGLVRYAN--G-S-LP-HPSA--VDLY-DG-
RSQRITSAGTPRVGQSVRRSGSTTGVRDGRVTGLNATV-NY-G--NG-------QVVRGLIKTDVCAEPGDS-G-
GALFS-G---------S--TALGLASGGS--------GDCRK---------------GGTTFYQPVVAALDAYGVRLF
>XX00000328 WP_030728136 37 76 192 MSSERP--------------------------------------
----------------------------------------------------------------------------------------------------------------------------
----------------NPHRRLV-K-RL--RL--IAAA---------------S-GVAA-L-TA-----F-V-----LP-------ST-AA-A--E-
PVQTFSTAQLAT-ADSAVLA-----------------------------------------------------------------------------------------
-------------------------------------------------------------------------------------------------------AD------IA-
GTAWTVDEETGQVVVTVDSTV-T----RE--E-IAE-IKDSAG-SL----A-G-A-ISI---------ER---------T-----P-GTL--
E-N-YLSGGDA-IY-----A----S----GWRCSAGFNATRG-G-T--TV-LLTAGHCTDG-AG--T--WW-A-----NSA-R-
T--TVIGSTIGSSFPVNDYGLISYTN--S-S-LA-RPSN--VYLY-NG-
TYRTITGAANATNGLAVQRSGSTTGLRSGSVTGLNYTV-NY-G--GG-------DIVYGMIRTNVCAEPGDS-G-
GPLFS-G---------N--TAIGLTSGGS--------GNCSV---------------GGTTYFQPVTAALSAYGASIP
>XX00000333 WP_018537096 39 76 187 MRNERT-------------------------------------
----------------------------------------------------------------------------------------------------------------------------
-----------------TPRSGVA-R-RT--RL--IAVA---------------S-GLVA-V-GA-----V-A-----VP-------SAAGAQT--P-
APETFSAARLTA-ASDAVRA-----------------------------------------------------------------------------------------
-------------------------------------------------------------------------------------------------------AD------VA-
GTAWQIDPATHTLVVTADSTV-S----TA--R-LGR-IRQAAG-SN----A-G-A-LRI---------ER---------T-----A-
GRF--R-K-LISGGDA-IY-----A----P----SWRCSLGFNVRSG-S-T--YY-FLTAGHCTQG-KP--P--WY-T-----NSS-
D-S--TSIGPTTGSSFPGNDYGIVKYTN--G-S-LA-HAGT--V-------
GSQDITSAGNPTVGQTVTRRGSTTGVHRGRVTALNATV-NY-G--NG-------DIVSGLIRTTVCAEPGDS-G-
GPLYA-G---------T--KALGLTSGGS--------GDCTS---------------GGTTFFQPVTEALSAYGVSVY
>XX00000334 WP_055500617 41 76 186 MRIKRT-------------------------------------
----------------------------------------------------------------------------------------------------------------------------
--------------TADTGTDHR-R-RN--RL--IAAA---------------T-GLVA-V-GA-----L-V-----VP-------ATASAQA--P-
SPHTYSATQLSA-AGDAVRS-----------------------------------------------------------------------------------------
-------------------------------------------------------------------------------------------------------AD------VA-
GTAWHVDKDSGTVEVTADSRV-S----SA--D-LAR-IRQAAG-KN----A-G-A-LRV---------ER---------V-----K-

GTF--K-K-YVSGGDA-IY-----A----G----QYRCSLGFNVRKG-D-T--YY-FLTAGHCTEG-GG--N--WY-T-----
DSG-H-S--TLIGPTSGSSFPGNDYGLVRYSS--N---VA-HPGT--V-------
GSVDITGAGNATVGQTVTRRGSTTGTHSGRVTALNATV-NY-G--GG-------DIVSGLIRTTVCAEPGDS-G-
GPLYA-G---------G--TALGLTSGGS--------GNCSS---------------GGTTFFQPVREALDAYGVSVY
>XX00000357 WP_055612483 35 76 187 MTARGT-------------------------------------
------------------------------------------------------------------------------------------------------------------------
-----------------T-----I-R-RS--RL--AASV---------------L-SLLV-V-AA-----L-A-----VS-GTNATAAP-TA-G--A-A--
-PSAASVSG-AASAVGS-------------------------------------------------------------------------------------------------
-----------------------------------------------------------------------------------------------LG------VA-
GTSWAVDERTGQLRVLADSTV-G----EA--D-LAA-VRRATG-RF----A-G-A-VTV---------ER---------V-----D-
GRL--R-T-LLSGGDA-IY-----A----S----SWRCSVGVNVQSG-S-T--YY-FVTAGHCTDG-LP--T--WY-T-----SSG-
L-T--TMVGPTTGTSFPGNDFGVVRYSN--P-A-VP-HPGT--I-------
GTVDVTGTATAYVGQSVCRRGSTTGVRCGRVTALNATV-NY-G--SG-------DIVYGLIQTNICAEPGDS-G-
GPLYA-G---------D--KVIGILSGGS--------GNCTS---------------GGTTFYQPIQEVLSAYGLTVY
>XX00000358 WP_031126128 38 76 183 MTFKRF--------------------------------------
------------------------------------------------------------------------------------------------------------------------
----------------SLLTGTT-R-RA--RL--VAVA---------------S-GLVA-G-LA-----L-T-----AP-------SA-VA-A--
PEPQVKATASQLAD-ASAAMLG-------------------------------------------------------------------------------------
---------------------------------------------------------------------------------------------------------VD------VA-
GTAWYTEASTGKIVVTVDSTV-S----KA--E-LAK-VKGALA-NS----K-A-E-LTV---------KR---------T-----T-GKF-
-T-P-LIAGGEA-IT-----T----G----GSRCSLGFNVSVG-G-V--AH-ALTAGHCTNI-SA--S--W-----------------
SIGTRTGTSFPNNDYGIIRHSN--P-A-A--ADGR--VYLY-NG-
SYQDITTAGNAFVGQAVRRSGSTTGLRSGTVTGLNATV-NY-G--SS-------GIVYGMIQTNVCAEPGDS-G-
GSLFA-G---------S--TALGLTSGGS--------GNCRT---------------GGTTFYQPVTEALSAYGATVL
>XX00000359 KND28857 26 76 189 MSKR--------------------------------------------------
------------------------------------------------------------------------------------------------------------------------
----------------A--RL--VAVA---------------S-AFVA-A-TA-----V-T-----VP-------AA-AA-D--A-
APAKLSGAQLAQ-VTDAVET-------------------------------------------------------------------------------------------
---------------------------------------------------------------------------------------------------AD------VV-
GTAWYTDPASGKVVVTADATV-S----AA--E-IAE-IKAAAG-DR----A-G-G-LQI---------NR---------T-----P-
GTF--S-K-LIAGGDA-IY-----S---------GGRCSLGFNVRSG-N-T--YY-ALTAGHCTAI-GS--T--WY-T-----NSG-
Q-S--TTLGTRTGSSFPGNDYGIIRHAN--A-S-A--ADGR--VYLY-NG-
SYRDITAAGNATVGQSVQRSGSTTGLRSGSVTGLNATV-NY-A--EG-------T-VSGLIQTNVCAEPGDS-G-
GSMFS-G---------T--TALGLTSGGS--------GNCTS---------------GGVTFFQPVTEALNAYGVSVF
>XX00000073 WP_018386864 18 77 186 MVS-------------------------------------------
------------------------------------------------------------------------------------------------------------------------
-------------------------------------------------------GLLA-A-VA-----L-A-----VP---GAAAAT-DS-P--A-G---
AAAPAART-AAAQLDT-----------------------------------------------------------------------------------------------
------------------------------------------------------------------------------------------VR------GT-
GVAWGADASGDTLRIQVDPTV-R----GA--D-LAA-VQRVAD-RY----G-S-A-VRV---------ER---------L-----D-
TPL--T-L-RLSGGDA-AY-----T----S----AYRCTVGVNVRKG-T-T--DY-FVTAGHCAGV-GS--T--WY-A-----
DAG-R-T--TTVGPTVSVDFPGNDYALVAYAN--G-A-VP-RPGT--I-------
GSVDITGVQDPVVGLSVCMRGGTSGVHCGVILALNQSV-SF-P--EG-------T-ITGLIRTNICSEPGDS-G-
APLYS-G---------D--KVIGILVGGS--------GNCAS---------------GGTSYFQPIREVLNAYGVSVY

>XX00000149  WP_051084883  26  77  190  MTASR-------------------------------------
-------------------------------------------------------------------------------------------------------------------------
-------------------------------------------------------------------------------------------------------------------------
-------------------------------------GVALLV-T-GALALTALPA--AT------AVSAVP-A--KPE-P------RTVAA-
ITAAL-NKD---------------------------------------------------------------------------VE------IP-GTAWMTRPD-
GKILVSYDSTV-V----GA--K-LQS-LTKVTK-QF----G-N-Q-VVL---------EK---------V-----P-GKF--K-K-
YLGGGGA-IY-----G----G----DFKCSLGFNVTRK-G-R--YY-FLTAGHCGND-AA--N--WY-H-----DSN-H-T--
NRIGTVHHSSYPWNDYALVVYRT--G-I-T--PGGS--VNLY-NG-
RSQNITTAANPGIGQLVYRTGATTGLASGRVTGLGATV-NY-D--DG-------R-VAGLIRTSVCAEPGDS-G-
GPLFY-R---------T--WAFGLTSGGN--------GDCTS---------------GGVTFFQPVVEALNVYGVNVY
>XX00000326  WP_030900885  38  77  190  MTSKRL------------------------------------
-------------------------------------------------------------------------------------------------------------------------
---------------TPRRTVS-R-RA--RY--LAVA---------------A-GVVA-A-AA-----F-T-----AP-------SALAQPA--
PVASPTATAAALAA-TAGAVNQ------------------------------------------------------------------------------------
----------------------------------------------------------------------------------------------------TA------AP-
GTAWYTDPASGKVVMTVDSTV-S----PA--Q-VAK-IKSAVP-D-----S-S-K-LVV---------KH---------A-----A-
GEF--K-P-LVAAGDA-IQ-----N----S----EYRCSLGFNVRSG-S-T--YY-ALTAGHCTAE-GT--S--WS-T-----PS----
G--TPVGTTADSQFPGNDYGLIRLSS--T-P-SA-SDGR--VDLY-NG-
SYQDITSAGNATVGEHVTRSGSTSGVHSGTVQAVNTTV-RY-A--EG--------TVSGLIQTNVCAEPGDS-G-
GPLFD-G---------S--KALGLTSGGS--------GDCTS---------------GGTTFFQPVTEALNAYGLSVF
>XX00000337  WP_030590950  38  77  187  MRHERT------------------------------------
-------------------------------------------------------------------------------------------------------------------------
----------------TLRSGAA-R-RG--RL--IAVA---------------S-GLAA-I-GA-----M-T-----LP-------TAHAGEP--A-
PPATFGASQLAA-VSDAVRS----------------------------------------------------------------------------------
----------------------------------------------------------------------------------------------------AD------VA-
GTAWVVDPKTHRVVVQADSRV-T----DA--Q-LDR-IKQRAG-AN----A-G-A-LKV---------ER---------V-----P-
GVM--R-K-LISGGDA-VY-----A----T----SWRCSLGFNVKKG-S-T--YY-FLTAGHCTDG-NP--P--WY-T-----
NSS-H-S--TSIGPTAGSSFPGNDYGLVRYSN--S-S-VS-HEGT--V-------
GGQDITKAGNATVGQTVTRRGSTTGTHSGKVTGLNATV-NY-G--NG-------DIVYGMIKTTVCAEPGDS-G-
GPLYA-G---------S--TALGLTSGGS--------GNCRS---------------GGTTYFQPVTEAISAYGVSIY
>XX00000341  WP_045861952  37  77  187  MRIRRT------------------------------------
-------------------------------------------------------------------------------------------------------------------------
---------------TPTKGIR-G-RT--RL--IAGL---------------S-GLAA-A-AA-----F-T-----AP-------AVAAE-A--P-
ATPRFSASQLSA-ASDAVLG-------------------------------------------------------------------------------------
---------------------------------------------------------------------------------------------------AD------VA-
GTAWHVDKKSNTLVVTADSSV-S----RA--E-IAR-IEKAAG-PQ----A-A-A-LKI---------ER---------T-----P-GKI--
R-K-LTQGGDA-IY-----A----D----SWRCSAGFNVRSG-S-T--YY-IVTAGHCTDG-AG--T--WY-S-----NSG-R-T-
-SVIGPTAGSSFPTNDYGLVRYSS--S-S-AS-HPGT--A-------
GGADITSAANATVGMSVTRHGSTTGTHSGTVTGLNATV-NY-G--GG-------DIVYGMIQTNVCAEPGDS-G-
GSLRN-G---------S--RAIGLTSGGS--------GNCTT---------------GGTTFFQPVVEALNAYGVSVY
>XX00000355  WP_007380272  35  77  187  MSRART------------------------------------
-------------------------------------------------------------------------------------------------------------------------
----------------D-----I-R-RP--RL--IAAV---------------V-SLLV-V-TA-----L-S-----VS-DTAAAAAP-PA-S--S-A---
PSTTSVSG-AASVVAS-----------------------------------------------------------------------------------------

-------------------------------------------------------------------------------------------LG------IA-
GTTWTVDGRTGKLRISADSTV-A----DP--D-LSR-IRQATG-RF----P-G-A-ATV---------ER---------L-----D-
GRL--R-T-LVSGGDG-IH-----A----S----AWRCSAGVNVRGG-S-T--YY-FVTAGHCTEA-MP--T--WY-T-----
SSA-L-T--TEIGPTTGTSFPGDDFGVVRYAN--P-A-VP-HPGT--I-------
GTVDVTGTATAYVGQSVCRRGATTGVRCGTVTALNATV-NY-G--DG-------STVSGLIRTNICAEPGDS-G-
GPLYA-G---------D--KVLGILSGGS--------GNCAT---------------GGTTYYQPIQEALNAYGLSVY
>XX00000369 WP_051841350 25 77 187 MSRI-----------------------------------------
--------------------------------------------------------------------------------------------------------------
---------------------------RL--LAVA---------------A-GLAA-V-AG-----L-A-----TP-------AAAGA-H--P-
ADEGFSAARLAA-AGASVLR--------------------------------------------------------------------------------------
------------------------------------------------------------------------------------------------AD------VA-
GTAWHTDPATGTLVVTADSTV-T----EA--D-LAR-IRHEAG-PD----A-G-A-LRI---------ER---------T-----P-
GKL--R-K-LLSGGDA-IY-----T----D----SWRCSLGFNVRSG-S-T--YY-VLTAGHCTDG-AG--T--WW-T-----
SSS-H-T--TTVGSTVGSSFPTNDYGLIKYAS--N-S-PV-PPGT--V-------
GSQDITSAVDATNGMSVTRRGSTTGIHGGTVTGLNATV-NY-G--GG-------DIVYGMIRTNVCAEPGDS-G-
GPLYS-G---------S--RAVGLTSGGS--------GNCTS---------------GGTTFFQPVVEALNAYGVSVY
>XX00000016 WP_037044357 40 78 182 MAAL----------------------------------------
--------------------------------------------------------------------------------------------------------------
--------------------------------------------------------------------------------------------------------------
-------------------------DGQRTAVR-LPGFRTVL--R-ALLVLTAAAALTGA---G--VAAASP-P--DPK-AAA--
PLAAPAA-VMALL-DAR--------------------------------------------------------------------------MP------AG-
LADYGIEGDVVDVRLVGDP--------SP--E-VTA-LF--DG-ID----P-A-L-LRL---------ET---------G-----A-GVP-
QH-Q-ALRGGQR-IT-----G----G----NTRCTLGFSAGSG-A-Q--DW-IITAGHCTRE-SD--E--WL-D-----DT----
G--EVIGDGAKTAGNGVDVGAVPVTS--S-T-AA-LP-E--------V-
ASVPVRGTVPAPVGSAVCLYGSTSGKACGTVSARNRTV-NF-D--GQ-------Q-QRGMVVASICTAQGDS-G-
GPYIT-A---------D-GQAQGVHSGAG------------N---------------GCTAYFTPIDQALSSLGLTLR
>XX00000203 CEJ94882 0 78 186 MS---------------------------------------------------
--------------------------------------------------------------------------------------------------------------
--------------------------------------------------------------------------------------------------------------
--------------------------------------------------------------------L--TLSKLEK-AQKSL-NK-LMKVD---------------
-----------------------------------------------------------KA-----SAG-IASHVVDIATNKLIIETLADS-R----DH--A--
QQ-LASKVG-LA----E-S-E-FAV---------RT---------V-----S-QMPSTR-A-SVISGDA-FLN---PD----A----
SLRCSIGFAVTN--------G-FISAGHCGRR-GQ--R--AT-D-----S---N-G--NPLGTFYDSIFPGNDMSYIQAVQ--S-
D-T--LYGY--VDGY-SG-SYYPIYGSQEAAIGASICRSGSTSGPTGGTIQAKGQTV-NY-A--EG-------T-
LYGLTQTNACSDHSDS-G-GSFLT-S---------G--QAQGVLSGGD--------GQC------------------
TSYYQEVNPILQKYGLTLV
>XX00000332 WP_019355338 39 78 187 MKIKRT-------------------------------------
--------------------------------------------------------------------------------------------------------------
--------------KANDSTAGR-R-RN--RL--IAVA---------------S-GMVA-A-GA-----L-I-----VP-------ATAAAEA--P-
SPHTFTASQLGS-ASDAVRA--------------------------------------------------------------------------------------
-----------------------------------------------------------------------------------------------AD------VA-
GTAWHVDKKSDKVVVAVDSRV-S----DA--D-IAR-IRHSAG-KN----A-G-A-VEI---------EH---------V-----K-
GTF--K-K-YLSGGDA-IY-----S----G----SYRCSLGFNVRSG-S-T--YY-FLTAGHCTEG-GG--D--WY-T-----NSS-
H-S--TRIGPTVGSSFPGNDYGIVRYDN--T-S-IS-HPGT--V-------

GGQDITSAGTPSVGQTVTRRGSTTGTHSGQVTGLNATV-NY-G--GG-------DIVYGLIRTTVCAEPGDS-G-
GPLYR-G---------S--TALGLTSGGS--------GNCSS---------------GGTTFFQPVTEALNAYGVNVY
>XX00000371 WP_014173463 16 78 188 MAAV---------------------------------------
-----------------------------------------------------------------------------------------------------------
-----------------------------------------------------------------------------------------------------------
------SSLFTAAA--------------------------------------------VAT---A--GTSAAA-A--AE----A--SPASVSQ-AASAI-
E-S----------------------------------------------------------------------LN------VP-
GTAWTVDERTGTLRVSVGSTV-G----GD--D-LAR-LDRTAA-RS----G-G-A-VTV---------ER---------L-----G-
GPL--R-T-LLSGGDG-AY-----A---SA----GWRCSVGVNVQSG-A-T--YY-FITAGHCTDG-LP--T--WY-T-----
GAD-L-T--TAVGPTTGTSFPGDDYGVVRYAN--A-A-VP-HPGT--V-------
GTVDITGTATAYVGQEVCRRGATTGVHCGRVTSLNATV-NY-G--NG-------EIVYGLIQTNICAEPGDS-G-
GPLYA-G---------D--KIIGILSGGT--------GDCAT---------------GGTTYYQPIQEVLSAYGLTVY
>XX00000134 WP_040754605 30 79 181 MRFTRS-------------------------------------
-----------------------------------------------------------------------------------------------------------
-------------------------RN--RL--LAAA---------------A-GGLA-A-SA-----A-T-----MP-------AA-PP-E--E----
RPSARALAA-VDSAVNR--------------------------------------------------------------------------------------
------------------------------------------------------------------------------------------SA------LS-
GIAWYTDEATGKVVVTADSTV-P----AS--A-LSR-LTDAVG-VD----A-D-A-ITL---------KR---------A-----K-
GEF--K-P-FVGAGDA-IY-----G----T----GYRCSVGFNVRSG-G-T--HY-FLTAGHCADD-VE--T--WY-T-----
NAG-E-T--TRIGPTVKASFPGNDYALVRYAN--T-S-LT-HAGG---------------
YSVGDAHVGQRVTRDGSSTGVHSGRVEALDVSV-RY-R--GS-------GTVRGMIQTDICAESGDS-G-GPLYA-
G---------S--TALGITSGGT--------GDCSS---------------GGTTFYQPVREAVRAYDVNVY
>XX00000364 WP_059198448 28 79 187 MFRNRT------------------------------------
-----------------------------------------------------------------------------------------------------------
-----------------T-----L-R-RS--RL--CASA---------------L-GLLI-L-AA-----------------VAGSGTG-AA-A--A-A---
TTRASLTG-AAVAVES---------------------------------------------------------------------------------------
----------------------------------------------------------------------------------------LG------IA-
GTSWVADERTGRLRVFADAAV-S----DP--D-LTR-LRQTTG-RF----T-G-A-ATL---------TR---------L-----D-
GRL--R-T-LVSGGDG-IY-----T----A----GRRCSAGVNVQSG-S-T--YY-FITAGHCTDG-AA--T--WY-T-----GSA-
L-T--TAIGPTTATSFPGDDFGVVRYTN--S-A-VP-HPGT--I-------
GTVDITGTATAYVGQAVCHRGATTGVHCGRVTAVNATV-NY-G--SG-------DVVYGLIQTNICAEPGDS-G-
GPLYA-G---------D--KVIGILSGGS--------GNCTS---------------GGTTYYQPIQEALTAYGLSVY
>XX00000044 WP_013675653 31 80 189 MPSRL--------------------------------------
-----------------------------------------------------------------------------------------------------------
-----------------------------------------------------------------------------------------------------------
----------------RRALCGVALVAATVTTALG-MTVSAGAA--D-AAPVTALLDP----------------------------------AA-
VMGLL-DAR-------------------------------------------------------------------------LP------AE-
IADYGVDEQAHRVVVDLTGAG-R----TP--A-VDT-LL--SG-ID----P-S-V-LQV---------HT---------G-----V-
PAPHHD-A-ALEGGDT-IV-----S----G----DLRCTAGFPATGG-G-R--TW-LLTAGHCTRG-TG--T--WS-V-----
ETA-D-GELVPIGSGARTAAAGTDVGAVPVTA--G-G-WH-VR-A-------AA-
GTVAVRGATEAGPGAAVCLYGSTSGKVCGTVIARNRTV-NF-D--GT-------V-QYGMTATTICSAEGDS-G-
GPYVT-P---------S-GQAQGIHSGGG--------I--GS---------------GCTSYFTPVSVAVSAFGLRLS
>XX00000072 WP_028428730 32 80 192 MRSTRR-----------------------------------
-----------------------------------------------------------------------------------------------------------

----------------TPRGARA-G-RL--AA--LGLA--------------V-LLPA-G-PA-----L-A-----AS----------G-PGGP-
APESFTGAQLDL-ASEAVRV-----------------------------------------------------------------------------------------
-----------------------------------------------------------------------------------------------AD------VA-
GTAWHVDEAANRLVVTVDSTV-D----EV--D-VDR-LRQVAG-PR----S-G-A-MRF---------ER---------T-----P-
GVL--A-P-LVSGGQA-VH-----T----E----EGRCSLGFNVRGD-G-A--DG-FLTAGHCAVG-SD--T--WY-A-----
DAN-R-A--EELGATEESSFPGDDYALVRYSG--E-D-TE-RPGT--VDLH-NG-
RSQDITEAGEATVGQAVRRSGSTTGLRAGEVTGLDATV-NY-G--DG-------RIVRGLIRTDICAESGDS-G-
GPLFA-D---------G--TALGLTSGGS--------GNCRT--------------GGTTFFQPVSPVLERFGVDVY
>XX00000193  EPH42135       18        80        190      MAI---------------------------------------------------
-------------------------------------------------------------------------------------------------------------------------
-------------------------------------------------------------------------------------------------------------------------
-------------------------------------TALVSTAFTFQ---H--AEAAGT-G--ARV-ADQ-STLAELKD-ARAEF-DRR----
----------------------------------------------------------------------SS------TP-GTSWVISPAAGKLVVTADRTV-
R----GA--E-LRE-LKAVVE-DL----G-D-K-ARL---------TY---------T-----K-GEF--K-R-HIAGGDG-VY-----D----A-
---SGYCSLGFNVVDD-G-N--PY-FLTAGHCTNE-SA--D--WS-E-----TE--G-G--
PPIGTTADSSYPGDDYGIVEYTA--D-V-P--HPSA--VNLY-NG-
TTRPISDAADPYIGQYVHRSGATTGLHDGQVTNLDAAV-DY-G--DG-------P-VYGLIETDVCAEPGDS-G-
GSLFA-G---------E--TALGLTSGGT--------GDCAS--------------GGTTYFQPVTEPLAVYGVSIG
>XX00000316  WP_047019641          38        80        196      MSTERT------------------------------------
-------------------------------------------------------------------------------------------------------------------------
-------------------------------------------------------------------------------------------------------------------------
----------------------------NPLRRTARRFRRPVTALT-ALAVLALPAGLGLA---G--PQA--A-Q--AQP-RTF--
STGELAA-AGDAV-L-A--------------------------------------------------------------------------AD------IA-
GTAWGVDQATGTVLVTVDERV-S----AA--E-IAE-LRRSAG-AQ----A-G-A-LTF---------ER---------T-----A-
GTF--R-P-YLQGGEP-VW-----S----T----TGRCTAGFNVRVS-N-A--DY-FVTAGHCTAG-FP--T--WY-T-----
NSS-R-T--TLVGPTAASSYPGNDFGIVRYTN--P-A-VP-RPGT--VN-C-NG-
TIIDITGPANLVVGQSVSMAGGATGCHGGGITGINVTV-NY-G--DA--------IVSGLIRTTLCCGPGDS-G-
SPVFT-L---TGDGTTA--LAVGVLSGGS--------GNCTS--------------GGTSFVQPINEILTAYGAVLT
>XX00000330  WP_031088079          37        80        187      MRIKRT------------------------------------
-------------------------------------------------------------------------------------------------------------------------
----------TTPHDGHPKRTR-T-AA--AV--IAVV--------------A-GLFA-V-TA-----Q-A-----VP------AA-GA-D--
P-A-GPYSADQLGA-AGQAVLD-------------------------------------------------------------------------------------
--------------------------------------------------------------------------------------------------AD------VA-
GTAWHLDPVGKRLVVTVDSTV-T----PA--D-IDR-IKRTAG-AQ----A-G-A-LRI---------EH---------T-----P-
GKL--R-A-LISGGDP-VH-----T----D----GWRCSLGFNVRKG-D-A--HY-FLTAGHCTEG-AG--T--WY-A-----
DSG-K-T--TVLGTTYGSSFPGDDYGIVRYTD--D-S-VP-KEGT--V-------
GGQDITGAATATVGMSVTRRGSTTGIHSGTVTGLNATV-NY-G--NG-------DIIQGLIRTDVCAEPGDS-G-
GPLHS-G---------P--RAIGLTSGGS--------GDCTT--------------GGTTFFQPVTEALSAYGVSVY
>XX00000346  WP_030812438          32        80        188      MRIKRT------------------------------------
-------------------------------------------------------------------------------------------------------------------------
---------------THRGRIA-R-RT--QL--AAAV--------------S-GLLA-V-AA-----F-A-----AP-------TA-NA----S-
DVHTFSAGQLTK-ASDSIRQ-----------------------------------------------------------------------------------------
-----------------------------------------------------------------------------------------------AD------VP-
GTAWAVDSKTNRVLVTVDSTV-S----QA--E-IAT-IKRQAG-AN----A-G-A-LTI---------KH---------T-----P-

GKF--Q-K-LISGGDA-IY-----A----S----SWRCSLGFNVQDSAG-N--YY-FLTAGHCTDG-AG--T--WW-S-----
NSS-H-T--TTLGSTAGSSFPGNDYGIVRYTN--S-S-VT-KSGA--V-------
GSQDITSASTPSVGTSVTRRGSTTGIHSGTVQALNATV-NY-G--GG-------DIVYGLIQTNVCAEPGDS-G-
GPLYG-G---------T--RAYGLTSGGS--------GDCTS---------------GGTTFFQPVTEALNAYGVHVY
>XX00000351 WP_046424140 32 80 187 MRIKRT-------------------------------------
--------------------------------------------------------------------------------------------------------------
----------------TAPSGPA-R-RT--RL--IAVA---------------A-GFLA-A-AA-----F-A-----AP-------TA-NA----S-
DARPFSAAQLTR-ASDSVMK---------------------------------------------------------------------------------------
---------------------------------------------------------------------------------------------AD------IP-
GTAWAVDAKNNHVVVTVDNTV-S----KA--E-IAK-IRKQAG-AD----A-G-A-LTV---------KR---------T-----P-
GTF--N-K-LISGGDA-IY-----G----G----QYRCSLGFNVRSG-S-T--YY-FLTAGHCGQV-AS--T--WY-S-----NSG-
H-T--TVLGTNVGYSFPGNDFALVRYTN--S-S-IA-HPSA--V-------
GSQTISSAATPSVGQTVYRRGSTTGTHSGKVTALNATV-NY-G--GG-------DIVSGLIQTTVCAEGGDS-G-
GPLYN-G---------S--VAYGLTSGGS--------GNCTS---------------GGTTFFQPVTEALSYYGVTLP
>XX00000365 WP_033254233 31 80 181 MRTTRT-------------------------------------
--------------------------------------------------------------------------------------------------------------
------------------TTAPRR-T-GG--RL--LAVA---------------A-ALAA-A-TA-----L-A-----VP-------AA-GA-A--T--
AAPDATAALAR-VSDSVEF-------------------------------------------------------------------------------------
------------------------------------------------------------------------------------------SG------VN-
GIAWHTDPASGKVVVTADSTV-T----PA--A-LAR-LTAAAG-SD----S-A-R-LKV---------ER---------T-----P-
GTF--S-P-RLSAGAA-IY-----G----G----GYRCSLGFNVVKG-S-T--YY-FLTAGHCGNI-AS--T--WY-T-----NSG-
Q-S--TVAGTTAGSSFPGNDYALVRYTN--T-S-LS-HPGG---------------
FTAANAYVGEAVKRTGSTTGTHSGKVTALNATV-RY-S--DG-------GTVSGLIQTNVCAEGGDS-G-GPLYD-G-
--------T--KALGLTSGGS--------GNCTT---------------GGTTFFQPVTEALSAYGVSIY
>XX00000109 WP_036963149 33 81 182 MNARPT-------------------------------------
--------------------------------------------------------------------------------------------------------------
------------ASRPAA----TK-R-AR--LA--VLAM---------------ILGLMA-P-AT-----A-A-----PA-------QA-RD-S--D-
---EPTDAELAE-LHEAIEE----------------------------------------------------------------------------------------
-------------------------------------------------------------------------------------------SD------VE-
GVAWYTDEAAGEVVVTADSTV-T----GA--E-RNA-VRRAAG-DK----I-G-A-LDL---------QR---------M-----E-
GEF--KKR-SMPPGAA-IY-----G----S----GTRCSLGFNARRG-N-K--HY-LITAGHCGND-VA--T--WR-A-----G--
-N-Q--KRIGPTVSSRFPGVDYALVRYDT--R-S-LK-RPGG---------------
YLPGTARVGQKVTRVGSTTGQHSGVVTAVGVTV-RY-G--GG-------AMVGNLIQTNICAEPGDS-G-GPLYS-
G---------R--RALGITSGAA--------GACPA---------------HGISLYQPIKPVLAAYGVQLY
>XX00000325 WP_047014981 32 81 192 MRIKRT-------------------------------------
--------------------------------------------------------------------------------------------------------------
----------------PPLSGTK-K-RL--RL--VAAL---------------T-AALA-G-AA-----L-A-----VP-------AA-GA-A--Q-
PPEGFSAATLAR-ADDAVLA-----------------------------------------------------------------------------------------
---------------------------------------------------------------------------------------------AD------VG-
GTAWGVDKKTGTLVVSIDETV-T----NA--E-VAR-IKRTAG-AD----A-G-A-LRI---------ER---------M-----K-
GKL--Q-S-YINGGQA-IY-----A----S----SWRCSLGFNVRSG-S-T--YY-FVTAGHCTDG-AG--T--WY-S-----
NSG-R-T--TTIGPTVGSSFPTNDYGLVRYSN--T-S-QT-HPGT--VNLY-NG-
SSQDITRAANATVGQSVQRSGSTTGLHGGSVTALNQTV-NY-G--GG-------DIVYGMIRTNVCAEPGDS-G-
GPLFS-G---------T--TALGLTSGGS--------GNCST---------------GGTTFFQPVVEALNAYGVSVY

>XX00000329 WP_018849390 32 81 191 MNLKRL-----------------------------------
--------------------------------------------------------------------------------------------------------------------------
----------------SPLGGTA-K-GA--RL--IAVA--------------S-ALVA-A-TA-----L-A-----AP-------T--AL-A--
EPAPARATAAQLAQ-AGDAVAK-------------------------------------------------------------------------------------
------------------------------------------------------------------------------------------------------AD------VA-
GTAWYTDTATGKVVVTVDSTV-T----AA--E-IKR-IKQQAG-NR----A-G-A-LEI---------NR---------T-----E-
GTF--T-K-LIAGGQA-IY-----A----G----GGRCSLGFNVRSG-S-T--YY-ALTAGHCTDS-AS--T--WY-T-----NSA-
N-T--ALLGSRAGSSFPVNDYGLIRHSN--S-A-A--ADGR--VYLY-NG-
SYRDITSAANPSVGQSVQRSGSTTGLRSGTVTGLNATV-NY-G--GG-------DVVYGMIQTNVCAEPGDS-G-
GPLFS-G---------S--TALGLTSGGS--------GNCRT--------------GGTTFFQPVVEALNAYGVSVF
>XX00000361 WP_012918700 23 81 192 MRIRRA------------------------------------
--------------------------------------------------------------------------------------------------------------------------
--------------------------------------------------------------------------------------------------------------------------
-----------------------------------VALLATA-G-LATTTVQLAA--PA------NAAPGG-E--APA-V------TSASS-
ITATL-AKE--------------------------------------------------------------------------AS------IP-
GTAWMTDEKSGRIIVSYDDTV-S----GG--K-FAA-LTAVTK-RF----G-S-Q-VVL---------EK---------L-----P-
GVL--S-K-RISGGQA-IY-----G----G----GYRCSLGFNVRDSAG-T--YY-FITAGHCTNS-AS--T--WY-A-----NSS-
Q-S--TVLGTRTGSSFPGNDYGIVRYST--S-Y-TN-HPGN--VYLY-NG-
SYQDITTAGNASVGQAVRRSGSTTGLRSGSVTGVNATV-NY-P--EG-------S-VSGLIRTNVCAEGGDS-G-
GSLFA-G---------S--TALGLTSGGS--------GNCST--------------GGTTYFQPVIEVLNRYGVNVY
>XX00000006 WP_045549200 29 82 179 MRFERT-----------------------------------
--------------------------------------------------------------------------------------------------------------------------
----------------T----FR-P-RT--LA--ALVA--------------GVGLAA-S-AA-----V-A-----VP-------AA-AA-D--A----
EFSKAEVAS-VKSAVAE-----------------------------------------------------------------------------------------
--------------------------------------------------------------------------------------SA------PV-
GSSWYTDPATNKVVITVDETV-S----AS--A-IAE-VKKAAA-EKTNQGD-Q-A-VTV---------KH---------T-----K-
GEF--T-P-LIGAGDA-IY-----G----G----GYRCSLGFNVKVG-G-A--DY-FLTAGHCTND-VS--T--WY-A-----NSS-
Q-T--TLLGSTAGSSFPGNDYGIVRYNS--T-I--S-KTGG--------------
FTAGTARVGQSVTRDGSTTGVHTGTVTAVNASV-RY-A--EG--------TVSGLIQTNVCAEPGDS-G-GALYS-G-
--------S--TALGLTSGGS--------GNCTT--------------GGTTFFQPVTEALSVYGATIY
>XX00000168 WP_030231674 31 82 182 MRTKRR-----------------------------------
--------------------------------------------------------------------------------------------------------------------------
---------------TPRTGTTR-R-RT--RL--LAVA--------------L-GLLT-A-TA-----V-G-----VP-------GA-AA----A-
PAHTAEPEQLAA-ISDAVLK-------------------------------------------------------------------------------------
-----------------------------------------------------------------------------------------------AE------VN-
GTAWYVDEAAGRVVVTADSTV-S----PA--G-LAK-IMRTAG-TG----A-A-T-LTV---------HR---------V-----P-
GVF--T-P-LLAAGDA-IY-----G----G----KYRCSLGFNVVGG-S-T--HY-FLTAGHCGNV-AP--D--WY-T-----
DAA-H-G--TRIGPTVSSTFPGHDYALVRYDN--S-A-LA-HPG-------------
GFTSAPNAVVGESVKRTGSTSGTHGGQVTGLNATV-RY-T--DG-------GTVRGLIQTNVCAEPGDS-G-
GPLYD-G---------T--KALGLTSGGS--------GDCTS--------------GGTTFFQPVNAALAAYGVSVY
>XX00000318 WP_031070896 36 82 191 MEIKRF------------------------------------
--------------------------------------------------------------------------------------------------------------------------
---------------TPHGGVA-R-GL--RL--TAVA--------------S-ALLT-A-AA-----L-A-----AP-------SA-AA-D--
SAPAPRATTAQLAA-ASDAVLG-------------------------------------------------------------------------------------

-----------------------------------------------------------------------------------------------------------AD------VP-
GTAWYTDSASGKLVLTADSTV-S----AA--E-LAK-IKKALG-SR----S-G-T-LEV---------KR---------T-----P-GTF--
N-K-LIAGGQA-IY-----A----G----GGRCSLGFNVRSG-S-T--YY-ALTAGHCTNS-GS--T--WY-T-----NSA-N-T--
AVLGTRTGSSFPGNDYGIIRHSN--A-S-A--ADGR--VYLY-NG-
SYRDITGAGNASVGQSVQRSGSTTGLRSGTVTGLNATV-NY-G--NG-------DVVYGLIQTNVCAEPGDS-G-
GALFS-G---------T--TALGLTSGGS--------GNCSS---------------GGTTFFQPVTEALSAYGVSVF
>XX00000320 WP_023590119 32 82 193 MNFKRF------------------------------------
---------------------------------------------------------------------------------------------------------------------------
-----------------TPRGGFA-R-GA--RL--TAIA--------------A-AMVT-A-TA-----L-A-----AP-------SA-GA-E--
TADATRASVAELAQ-VSEAVLD------------------------------------------------------------------------------------------
-----------------------------------------------------------------------------------------------------------AD------VP-
GTAWYTDAKSGKLVITADSSV-S----AA--E-LAQ-LKKAAG-DK----A-G-A-VEI---------KR---------T-----P-
GTF--N-K-LIAGGEA-IY-----A---SG----GGRCSLGFNVRNSSG-A--TY-ALTAGHCTEI-AS--T--WY-T-----NSG-
Q-T--TLLGSRAGTSFPGNDYGIIRHSN--S-A-A--ADGR--VYLY-NG-
SYRDITGAGNAYVGQNVQRSGSTTGLHSGRVTGLNATV-NY-G--GG-------DIVSGLIQTNVCAEPGDS-G-
GALFS-G---------S--TALGLTSGGS--------GNCRT---------------GGTTFFQPVTEALSAYGVSII
>XX00000321 WP_010474987 32 82 193 MNFKRF------------------------------------
---------------------------------------------------------------------------------------------------------------------------
-----------------TPRNGLS-R-GA--RL--TAIA---------------A-ALVS-A-TA-----L-A-----AP-------SA-GA-E--
TPDTTRATAAQLAQ-VNEAVLE-----------------------------------------------------------------------------------------
------------------------------------------------------------------------------------------------------------AD------VP-
GTAWYTDAKSGKLVVTADSTV-S----AA--E-LAK-LKKAAG-DR----A-G-A-LEI---------KR---------T-----P-
GKF--N-K-LIAGGQA-IY-----G---NG----GGRCSLGFNVRTSSG-T--PY-VVTAGHCTEG-TS--T--WY-T-----
NSS-Q-T--ATLGRTAASSFPNNDYGVISHAN--P-S-A--ADGR--VYLY-NG-
TYRDITGAGNAYVGQTVQRSGSTTGLHSGRVTGLNATV-NY-G--NG-------DVVYGLIQTTVCAEPGDS-G-
GAMFA-G---------S--TALGLTSGGS--------GDCRS---------------GGTTFFQPVTEVLNAYGLTII
>XX00000327 WP_049572611 33 82 190 MNPQQ-------------------------------------
---------------------------------------------------------------------------------------------------------------------------
---------------TSNSGRPTA-R-WG--RL--IAAA---------------S-CLLA-L-AA-----M-A-----IP-------GA--AQA--
VQAPSAFTSAELAR-AERAVLA-----------------------------------------------------------------------------------------
-----------------------------------------------------------------------------------------------------------AD------VA-
GTAWGVDPATGTVRVLADETV-S----AA--E-IGR-VRRGAG-DL----S-P-A-LTI---------TR---------T-----P-
GEL--R-T-FLQGGEP-IY-----S----T----AGRCAAGFNGTAG-G-Q--DV-LITAGHCTSA-FP--T--WY-A-----DSA-
G-T--ALIGPTAASSFPGNDYGLIRYVS--S-L--S-RPGT--V-IC-NG-QVIDIIGARNPTIGETLWVA-
APTGCTSGGVTALNLTI-NY-G--GA-------DIVHGLGRLTACTEPGSS-G-APVFT-M--------SG--
YAVGIVSGGS--------GNCST---------------GGTTYFQPILEILNAYGVTLT
>XX00000340 WP_014059671 32 82 187 MRIKRN------------------------------------
---------------------------------------------------------------------------------------------------------------------------
----------------APHNKQA-R-RI--AL--IAAT---------------S-ALVA-G-AA-----L-A-----AP-------AY-AG-S--D-
GARTFSAAEAKS-ASDAVLK-------------------------------------------------------------------------------------------
-------------------------------------------------------------------------------------------------------AD------VA-
GTAWYVDKATNKLHVTADSTV-S----QS--E-IAK-IKNTAG-AS----A-H-A-IEV---------KR---------T-----S-GKI-
-Q-K-LISGGDA-IY-----A----D----SWRCSLGFNVRNSSG-A--NY-FVTAGHCTDG-AG--T--WW-S-----NSG-
H-S--TTIGPTAGSSFPTNDYGLVRYSG--S-A--T-PQGT--V-------

GSQDITSAANPTVGQTVTRRGSTTGIHSGQVTALNATV-NY-G--NG-------DIVYGMIQTTVCAEPGDS-G-
GPLYA-G---------S--TALGLTSGGS--------GNCTS---------------GGTTFFQPVVEALNAYGVSVY
>XX00000344 WP_030474858 27 82 191 MTSTL------------------------------------------
---------------------------------------------------------------------------------------------------------------------------------------
---------------------------------------------------------------------------------------------------------------------------------------
-----------------------------------------RILLAAV-ALLLGAVITPTTAL---A--SDAAHA-T--PRF-VTR--SAAVLQA-
ATDML-RRE-----------------------------------------------------------------------------------AT------IP-
GTSWSTDPVTNRVRVSADSTV-T----GT--K-LAQ-LQAVLA-KL----G-D-T-ARL---------ER---------L-----P-
GTL--R-T-TITGGDA-IY-----G----G----PYRCSLGFNVNRPGS-E--AY-FLTAGHCANA-AV--T--WS-A-----A---
G-G--QVIGTRAGSSFPGNDYALIRYTS--S-I-A--RPGA--VNLY-NG-
TQRDISGAGNAYVGQTVGRSGSTTGFRTGKVTALNVTV-NY-S--DG------TT-VTGMIATTVCAEGGDS-G-
GALFA-G---------N--TALGLTSGGS--------GNCTS---------------GGQTFFQPIVEALNAYGVRIY
>XX00000360 WP_053757918 27 83 187 MRARRT---------------------------------------
---------------------------------------------------------------------------------------------------------------------------------------
-----------------T-----V-R-HS--RL--IAAV---------------L-SMFT-M-AA-----L-AVSGTGA-------AAT-PG-S--G-T-
--ASVAAADR-AASAVAS-------------------------------------------------------------------------------------------------------
-----------------------------------------------------------------------------------------------------LG------IA-
GTAWTADRRTGAVRVFVDATV-G----DA--D-LAA-LRHATG-RL----P-G-T-VVL---------ER---------L-----D-
GRL--R-T-LVSGGDG-IH-----A----S----GRRCSAGVNVRAG-S-T--YY-FVTAGHCADG-AP--T--WY-T-----
SSA-M-T--TVIGPTGGSSFPGNDFGVVRYAN--P-A-VP-HPGT--V-------
GTVDVTGAGTAFVGQSVCRRGATTGVRCGVVTALNVTV-NY-G--SG-------ATVHGLIQTNICAEPGDS-G-
GPLYA-G---------D--KVIGILSGGS--------GNCTT---------------GGTSYYQPIQEVLNTYGVSVY
>XX00000362 WP_030205217 25 84 187 MRKKRI--------------------------------------
---------------------------------------------------------------------------------------------------------------------------------------
----------------T---------PT--RL--TALA---------------A-GLAA-C-VA-----L-A-----AP-------AGASGGT--D-
GGGGYSPARLAA-ADASVLR--------------------------------------------------------------------------------------------------
----------------------------------------------------------------------------------------------------------AD------VA-
GTAWHTDPATGTLVVVADSRV-S----PA--D-IAR-IRAAAG-AN----A-G-A-LRI---------ER---------T-----P-
GRF--T-K-LISGGDA-IY-----A----S----SWRCSLGFNVRSG-T-T--YY-VLTAGHCTDG-AG--T--WW-T-----
NSS-H-T--TTIGATVGSSFPVDDYGLVRYDN--T-S-LS-HPGA--V-------
GSQDITSATNATVGMSVTRRGSTTGIHSGSVTGLNATV-NY-G--GG-------DVVYGMIRTNVCAEPGDS-G-
GPLYS-G---------T--RAVGLTSGGS--------GNCTS---------------GGTTFFQPVTEALAAYGVSVY
>XX00000041 WP_050515494 33 85 200 MQRSLS--------------------------------------
---------------------------------------------------------------------------------------------------------------------------------------
----------------------FR-R--------------------------------------------------------------------------------------------------
-------------------AALAASTLVLAAA------------------SLAPFSTD-AATA---R--ARSAGD-E--AQN-VRG---
PDPLMA-TMATL-DRS----------------------------------------------------------------------------EL------IP-
GTAWGVDPRLKRVVVTADPTV-K----GA--R-LAR-LTKLVR-PL----G-K-N-VEL---------RR---------T-----T-
TKL--T-T-FIEGGDA-IH-----G---RN----AATCSLGFNVRRAGA-P--DA-FLTAGHCGNL-VK--S--WS-M-----
TR--R-G--PQFARTADSRFPGNDYALARYTA--A-V-P--HGSA--VDLH-NG-
RWLPVTGARDAVVGDRVGRSGATTGAHLGTVTALNQTV-TY-P--EG-------R-VYGLTRTNVCAEPGDS-G-
GPLFT-Y-DPGNPPVTT--IALGLLSGGS--------GDCSR---------------GGTTYYQPVAEALDAYEAVIP
>XX00000172 WP_053161351 28 86 189 MKTFRL--------------------------------------
---------------------------------------------------------------------------------------------------------------------------------------

----------------S-----L-R----------------------------------------------------------------------------------------------
----------------------ALPATAVALAAVTP-PQQAGASP---------------TAPA---A--VTAAGG-A--------A--
TAARLAA-ARRTL-DTM---------------------------------------------------------------------------AA------IP-
GTAWVTDPRSHRVVVTADPTV-T----GT--R-LAR-LTEVTT-GL----G-D-A-VEV---------RR---------T-----A-
TPL--V-R-YAAGGDA-VW-----G----G----AARCSLGFNVSRG-G-R--AA-FLTAGHCGKA-TA--E--WA-E-----
DK--G-G--SRAAVTETSVFPGHDYALARYTD--G-T-P--RPSA--VDLH-GG-
GTRVITRAGEARPGERVQRSGSTTGVRGGTVTGVDATV-NY-E--EG-------R-VEGLIETDVCAEPGDS-G-
GPLFD-G---------G--TALGITSGGT--------GNCRT---------------GGKTYYQPVREALRALGASIG
>XX00000335 WP_041662198 28 86 188 MRTMRT----------------------------------
----------------------------------------------------------------------------------------------------------------------
------------------T----PT-R-TV--RL--LAVA--------------A-GLAA-A-AA-----L-A-----APT-ASAGTAD-S--A--R-
STRTFDAAALSA-TGDAVRA-----------------------------------------------------------------------------------
---------------------------------------------------------------------------------------------------------AD------VA-
GTAWYADTATGELVVTADSTV-T----PA--G-IAK-IKRQAG-AN----A-D-A-IRV---------ER---------T-----P-
GKF--T-K-LISGGDA-IY-----A----T----SWRCSLGFNVRDSAG-N--YY-FLTAGHCTDG-AG--T--WY-S-----
NSS-R-T--TVLGTTAGSSFPGNDYGLVRYTN--S-S-VT-KSGT--V-------
GSVDITSAANATVGMSVTRRGSTTGIHSGSVTGLNATV-NY-G--GG-------DIVSGLIRTNVCAEPGDS-G-
GPLYS-G---------S--RAVGLTSGGS--------GNCST---------------GGTTFFQPVTEALSAYGVSVF
>XX00000087 WP_052744163 19 88 188 MGV------------------------------------------
----------------------------------------------------------------------------------------------------------------------
----------------------------------------------------------------------------------------------------------------------
------------------------------------------------------------LTALAAG---I--LAPSPA-SGRTTA-GTF--SVAELNT-VRGAV-L-
E--------------------------------------------------------------------------------AD------VV-
GTAWSADPATGKVLVSVDETV-S----DQ--E-IGH-IRRTAG-RL----A-A-A-LKF---------ER---------I-----
PGGEM--T-P-FVKGGEY-IY-----G----F----DTRASLGFNTVYN-G-Q--DY-FVTAGHCTND-VS--H--WY-L-----
DSS-L-T--TYIGPTVGSSYPGNDFGVVRYDN--S-T-IP-RPGT--VR-C-GQ-NEVDITGSAEPRVGMPFNIAGYN-
-CRPATVQAVNVSV-NY-A--NG--------AVHGLFRANSCAAPGDS-G-AASFS-G---------S--LALGLLSGGT-----
---TSCTT---------------SGITYWQPVNEVLTAYGLTLT
>XX00000166 WP_006345345 0 93 196 MTG------------------------------------------
----------------------------------------------------------------------------------------------------------------------
----------------------------------------------------------------------------------------------------------------------
----------------------------------------------------------GTVLFGPE---A--VARDGS----P---TVF--STAQLTS-AGRAL-D-
G----------------------------------------------------------------------------------SG------VT-
GIAWAVDRKAGVVQVLADDTV-R----AA--Q-LAQ-LRSAAG-PL----A-D-A-LRV---------ER---------A-----P-
GRL--A-ERGPLGGDG-FR-----S----S----LGPCTFAFNVRIG-S-G--WY-VITAGHCILR-VQ--Y--IY-E-----RPP-
P-E--PPLGQVVRVDYPGNDFGLIRYLQ--P-P-QD-ALGS--VNLH-NG-
TVQDITTAGDPALGQPVCFSGSTSGRGCGQVTGLNWTV-SL-A--GG--------SVHGLIRTNICSAAGDS-G-
GPLFA-G---------T--TGLGMASVGS--------GSCAT---------------GGVTFFQPLGEALATYGATVY
>XX00000115 WP_018333832 31 94 205 MPLTAPAAR--------------------------------
----------------------------------------------------------------------------------------------------------------------
----------------RVGAAVALVLTAVLALA-----------------------V-P--------------------------------------------------
-----------------------------------------------------------------------ALAQTP---P-PLPSPGS-T--TRP-TPV--
EPGAGQA-AKQRL-DER--------------------------------------------------------------------------AP------AN-
VAAWFVDPTTNQTVVAVVGPA-T----KA--A--TD-FA--AD-EN----P-A-A-VRV---------QP---------M-----S-

APIRAA-A-PLVGGTA-ITT----T----T----GIRCSDGFSVRRN-T-T--TY-LLTAGHCTQN-TS--S--WS-G-----P---
N-G--QRIGNVALTAWPTNDYGIIQVANTAA-W-R--GSGQ--VR------
GGPTVTGATEAPVGTQVCRSGSTSGYHCGVIEAKNVTV-NY-G--NG------VT-VQGLTQTSACAESGDS-G-
GSFLA-P---------GTAQAQGLLSGGS--------GDCTT--------------GGTTYFQPLRPILNQYGLTLV
>XX00000324 CDR16288 22 96 188 MIAAM-------------------------------------------------
---------------------------------------------------------------------------------------------------------------------------------
----------------------S--GLFV---------------T-ATLASS-AS-----T-ATSSASIA-PSPSPSPS-TS-P--S-P---
SASTSVSG-AASAVAS------------------------------------------------------------------------------------------------------
---------------------------------------------------------------------------------------------LD------VP-
GTAWTVDERTGTLRVLVGSTV-R----GA--D-LAR-IDRTAE-RF----G-G-A-VTV---------ER---------V-----D-
GPL--R-T-LLSGGDG-IY-----S---ST----GVRCSAGVNVQSG-A-T--YY-FVTAGHCADG-NA--T--WY-T-----
GSG-T-T--TPVGPTTGTSFPGNDYGVVRYTN--T-A-VP-HPGT--V-------
GTVDITGTATAHVGQQVCRRGATTGVRCGVVTALNATV-NY-G--GG-------DIVYGLIQTNICAEPGDS-G-
GPLYA-G---------D--KIIGILSGGS--------GDCVS--------------GGTTYYQPIQEVLSAYGLTVY
>XX00000020 WP_037255569 21 97 217 MLSAC----------------------------------------
----------ALIISTLLTTGATA-QAAPNTAAIEYVMRADGVSRAEAIRRLDA-------------------------------------------
---------------------------------------------------------------------------------------------------------------------------------
-------------------------Q-----------------------PGQLRTAARLTGELGAARTTGAAIE--D-GELVVNVLDSQAAT---
K--VRAAGA-R--PRI-VSK--STVDVP-------------------------------------------------------------------------------------
---------------------ADPVTTVAV---------------------------------------------------------------------------------Q-
NVTPGLT-QV-----N----G----SEPCTAGFAAKDSFG-F--NY-MFTAAHCVKQ-VV--P--VV----------N-G--
VPMGDVVRFD-IGHDVALVRNLA--P-F-HL-QPHV--FNHE-TG-
KFVTVLDELVIVPGATVCKEGITSKHHCGRVNAINQPV-TT-K--FG-------V-VTGLIETDLCVKGGDS-G-
GPLYV-A---------S--SAAGITEGSK--------LFCGQ--------------PNVGYFVPIHPYLSAFGVSLI
>XX00000312 WP_037955997 29 98 188 MRRFR---------------------------------------
---------------------------------------------------------------------------------------------------------------------------------
-------------------------------L--IAAV---------------C-GLF--S-AA-----A-SSASTATS-PSASTPVS-AS-T--S-A---
SASTPVSG-AASAVES------------------------------------------------------------------------------------------------------
---------------------------------------------------------------------------------------------MD------VP-
GTAWTVDERTGTLRVLAGSTV-R----GA--D-LAR-LDRTAE-RF----G-G-A-VTV---------ER---------L-----D-
GPL--R-T-LLSGGDG-IY-----S---ST----GVRCSAGVNVQSG-A-T--YY-FVTAGHCADT-NA--T--WY-T-----
GSG-T-A--TPVGPTTGTSFPGDDYGVVRYTN--T-A-VP-HPGT--V-------
GTVDITGTATAHVGQQVCRRGATTGVRCGVVTALNATV-NY-G--GG-------DIVYGLIQTNICAEPGDS-G-
GPLYA-G---------D--KIIGILSGGS--------GDCVS--------------GGTTYYQPIQEVLSAYGLTVY
>XX00000039 WP_051299576 33 104 207 MARLR---------------------------------------
---------------------------------------------------------------------------------------------------------------------------------
---------------------------------------------------------------------------------------------------------------------------------
----------------PRRSYGVLLLAIALSVTLA-AIAFSTTLA-A-SGASMAQSNPAGFD---S--GGQDRS-P--SRD-
AAP--AAWQLAD-VQSQL-TRF-----------------------------------------------------------------------------------------------GA------
GQ-VQNWYVDQDRNSVVLMVRAGA-D----DA--K-TRA-FVEKAQ-SF----G-H-R-VDV---------ER---------V--
---R-SQA--R-A-YLYGGQP-VTL----S----N----GKQCSIGFNARTSSG-E--SI-FLTAGHCAAG-GP--I--AI-R-------
--N-G--SRIGHTIGRSFPGNDYAAMTINRPNA-W-N--PQPA--VDKY-NG-
YARVVRGHREPPVGSRVCKGGESTGWTCGTIQSYNQAV-NY-N--GN-------I-VYGLVRFSACVQSGDS-G-
GAVMR-G---------N--YAVGIVSGAC--------GQ-TA--------------PNVSFYQPIGEVLNAYDATLV

>XX00000038 WP_051814127 26 105 176 MARKVL--------G--AL--------------LP-----
-------AVVLA----------F--------------------------------------------------G--------------------------------------------------
-------------------------------------------------------------------------------------------------------------------------------
-----------------------------------------------------------AA--------A-VPA-----AAVPVEEP------QCGPA--SLSRLDG-
LMAEL-DARAA--------------------------------------------------------------------------------AG----QAGQ-
AQYWYVDPQACRLGVAVLRGA-DDERTEG--F-VGA-ATAVAG-QS----A-L-T-TLT-ELD-TPVVRRIA----
LVTAPDDS-I-ARM--P-G-TLHGGSV-IY-----S----GTSGTVYQCTNGFNNYRA-G-----T-TTTAGHCADA-AK--
T--WY-D-----A---K-G--NKIGSVSAWRFPGSDWAVITPVA--G-W-T--LPND--V----VG-
DEEEITAFGKATLGEPVCGRGSTTGTQCGTVTALNVTV-NY-P--DG-------L-VRGLVRSSQSGGEGDS-G-
GPVYH-G---------G--TGLGLISGGP--------A---N---------------GSTTFFQPMNF----------
>XX00000095 WP_035795974 29 106 179 MKLTAK--------R--SL--------------LP------
------IALLA----------L------------------------------------------------------G-----------------------------------------------------
-------------------------------------------------------------------------------------------------------------------------------
------------------------------------------------------AAAVPATPAAA-ATA-----IPPAVRGP------VCLPA--
QLDQLQG-VQSRL-DDLAR-----------------------------------------------------------------------------------SG----RAGN-
SQYWYVDQGTCKVAVAVLAGA-Q----DR--S-TRD-FLADLD-PQ----L-L-H-VTE------VASAVSSRVA-
TVPAPLDAGT-PAA--D-L-TFNGGTP-IF-----S----GTTGTVYECTSGFNNYRE-G-----K-ATTAGHCGKA-AP--
K--WY-D-----R---S-K--ELLGSVTADKFPGHDWAVITPTD--K-W-K--LNST--V-LD-DG-
VVRDVTGFALPKLNERVCGTGATSGTQCGSIEATNVTV-NY-P--DG-------L-VTGLAESTQDGNAGDS-G-
GPVFD-R---------E--LGLGLVSGGP--------K--GG---------------GGPTFFQPLNF----------
>XX00000370 WP_019816540 0 107 176 #NAME?
>XX00000407 WP_051865846 0 108 230 MLE---------------------------------------------
---------------------------------------------------------------------------------------------------------
ALQRDLGLTREQAQTRLLNE------------------------------------------------------------------------------------------
------------------------------------------------------ARLRPVEAKLRRRLGS-RFGGSWFAGSA-QTLVVATTDSADIP-
--Q--VLAAGA-R--GEV-VNR--STTED------------------------------------------------------------------------------------
-------------------------------II-------------------------KTIG-VD----A-S-A-VSV---------LP---------S-----D-
EQPRPL-S-DVVGGAP-YYV----G----V----TSRCSIGFSVLRG-T-Q--NG-FVSAGHCGKT-GA--V--TA-G-----V--
-N-R--VALGVFQASNFPDNDFGWVAVNA--D-W-T--PKPL--VDNG-TG-
GTVAVAGARVAIEGASVCRSGSTTGWHCGTIQQRNASI-TY-P--QG-------T-VSEVTRTNVCAEPGDS-G-
GSFIS-I---------A--S---ISTSRN--------GTVAS---------------GTPTPARPATTAPSRTGLTVD
>XX00000151 WP_020520449 24 113 184 MR--I--------------------------------------------
-------------------------------------------------------------------------------------------------------------------------------
------------------------------------------------------TL-------------------------------------------------------------------
--------VAATALATIGLSAPAQAADLSGLSGL-ADAGTYYS--D-GKLVAAVTSPEAAA---Q--IRAKGG-T--AKV-
VAR--TLGQLNS-MVTDL-D-------------------------------------------------------------------------------PH------IA-
GTAWSVDPATNQVVIDIDISV-T----GA--N-YDS-LHAIAA-KS----N-G-A-IRI---------EK---------M-----N-GLL-
-T-R-LIRGGNA-IY-----T----G----GSRCSLGFNVRSG-S-V--YS-FITAGHCTNI-GS--T--WN-D-----G-----S--
TTLGSRTGTSFPTNDYGIVRYNT--S-Y-TN-HPGS--IA---N---
GQDITSAGNASVGQSVRRHGSTTGNRGGSVTGLNATV-NY-A--QG-------S-VYQMIRTNVCAEPGDS-G-
GPLYA-G---------T--VALGLTSGGS--------GNCSS---------------GGTTFFQPVPEVLSRYGVAVY
>XX00000164 WP_050564149 0 113 205 M---------------------------------------------
-------------------------------------------------------------------------------------------------------------------------------
-------------------------------------------------------------------------------------------------------------------------------

-------------------------VDLLGD-RAPGMYLDD-D-GEFVVTVTDDAAAQ---Q--VSAQGV-Q--AKR-VNR--
DAKQLRD-IRATL-KE--------------------------------------------------------------------------MG------IP-
GTSWWVDPATNQVVLGYDTRV-T----DE--E-LET-LRGVAD-DA----D-G-A-LRL---------EA---------M-----N-
GDL--S-A-LAMGGEA-IY-----S----S----DFRCSLGFNVTNS-D-RSAFF-SLTAGHCTDG-SA--R--WY-T-----
DLP-D-K--NLLGFDLDSSFPGDDYGLIWTGS--E-A-Q--DWGM--VTLG-DG-
NVRPIYDHGEAYVGQSVEQYGSTSGLLSGEVTQVDATA-VY-R----GEDGEDDV-VEGLIGTDICSQKGDS-G-
GPLFA-G---------N--TALGMTSGGV--------ILCLT---------------ASWSFYQPIGEVLDEYGVELY
>XX00000051 WP_010358284 34 118 187 MIRA-----------------------------------------
---------------------------------------------------------------------------------------------------------------------------
-------------------------------------------------------------------------------------------------------------------------
P-VR-A-ASAISGTA-------------------------------------------AAS---P--ARATTA-I--PG----T--AATTPAH-
ATTTV-Q-A---------------------------------------------------------------------------LD------VA-
GTAWVPDPRTGQLHVLADPTV-T----AP--D-LAR-IQRATG---------R-S-ASV---------ER---------L-----G-GPL-
-R-T-LLSGGDG-IY-----T----S----AWRCSVGVNVQAG-T-A--YY-FITAGHCTDG-TS--T--WY-T-----DAA-G-S-
-TVAGTTVSTRFPVDDFGIVRYTN--T-S-VP-HPGT--I-------
GTVDITGAATAYVGESVCHRGGVSGVHCGRVTGLNATV-NY-G--SG-------DVVSGLIQTNICAEPGDS-G-
GPLYA-G---------D--KVLGIISGGS--------GNCTS---------------GGTTYYQPIQEALNAYGVSVY
>XX00000311 WP_051772592 27 119 187 MRTLKN--------------------------------------
---------------------------------------------------------------------------------------------------------------------------
----------------------------A--LV--IA----------------ATAI------------------------------------------------------------------
---------------TAAALVPTTATAAAVSPTDLLTSLGPDRTAGVHVD--A-GRAIVNVTDPAAAT---T--VRTAGA-T-
-PRL-VTR--SARQLED-VKAAL-D------------------------------------------------------------------------------------TA----
--TP-GTALAVDPITNQVVVTADSSV-G----TS--A-LAK-LRKSVA-DT----G-G-A-ARV---------ER---------A-----
A-GTF--A-P-LTTGGDA-IY-----G----G----QYRCSLGFNVRSG-S-T--YY-FLTAGHCTDS-AT--E--WF-A-----
DSG-H-N--TKLGDRTGSSFPGNDYGIVRYTN--T-S-VT-ISGS--V-----G--
GQDITSAGDAVVGESVTRRGSTTGTRSGTVTALDVTV-HY-S--GG-------GTVRGMIQTTVCAEPGDS-G-
GPLYD-G---------G--KALGLTSGGS--------GDCRR---------------GGTTFYQPVTEALGKYGVSVF
>XX00000019 WP_045744296 20 123 178 MLV-------------------------------------------
---------------------------------------------------------------------------------------------------------------------------
-------------------------------------------------------------------------G-G-----AL-------------------------------------------A-
------A-S---PVSAAPKAPALDASTLASSLDTQFGT-RDAGSYIDG-A-GKVVVNVTDAATAA---E--VKAAGA-T--
PRY-VAR--TGATLAA-ADSSL-KTS------------------------------------------------------------------------------LK-----
-TP-GTAFAVDPVSNQVVVSVDESV-T----GA--K-LAA-VEAIVA-KL----G-D-N-ARI---------ES---------I-----P-
GTL--S-P-NIAAGDA-IY-----T----G----GSRCSLGFNVRSG-S-T--YY-FLTAGHCTNA-GT--T--WT-N-----G-----
S--ATLGTTAGSSFPTNDYGIVRYTN--T-S-IT-KT-------------
GGFTSAATPSVGAAVTRRGSTTGTRSGTVTALNSTV-NY-S--QG-------S-VSGLIRTTVCAEPGDS-G-GPLYR-
G---------T--VAHGLTSGGS--------GNCTS---------------GGVTYFQPVVEALSVYGVSIF
>XX00000129 WP_033363971 33 123 188 MRLTRS--------------------------------------
---------------------------------------------------------------------------------------------------------------------------
----------------R-----L-R-RA--VV--IAAG---------------T-ALVA-G-SF-----LTP-----SP-------------------------------
------------A---------------------QAAPIGSTDALVEQLGL-ATAGAYWDAQL-GKMVVTVTDANTAK---S--
VESTGA-V--ARK-VKH--SGAELAR-ITATL-GAK-------------------------------------------------------------------------
-------VS------VP-GTAWAVDPVANQVVVSIDPTV-T----AE--H-VKT-IEATVA-SF----G-S-A-AKI---------ER---
------L-----P-GEL--R-S-TISGGQA-IY-----S----G----GSRCSLGFNVVSG-S-T--YY-FLTAGHCTNI-GS--T--WT-

T-----SG----G--SVLGTRAGTSFPGNDYGIVRYAS--G-V-A--HPGN--VSLY-NG-
SFQDITSAGNATVGQSVRRSGSTTGVRSGSVTQLNATV-TY-S--QG-------Q-VSGLIRTTVCAEPGDS-G-
GSLFA-G---------S--VALGLTSGGS--------GNCSS---------------GGVTYFQPVTEPLSVYGVSVY
>XX00000178 WP_018382917 44 123 188 MKSNT--------------------------------------
-----------------------------------------------------------------------------------------------------------------------
----------------VATRSTA-R-RV--RL--IAAA---------------A-GMIA-A-SA----FV-V-----VP--------------------------
-----------------TA---T----A---S------STASASPEEATALVAELGTTDTAGTYLDQAT-GKMVVNVTTDEAAE---
A--VRDSGA-V--PKF-VEH--SSAELAK-IDKAV-LGS------------------------------------------------------------
-----------VD------VA-GTSWGVDPVSNKLVISVDSTV-K----GA--D-LAK-VNKFAA-QY----G-D-A-VEV--------
-DR---------M-----Q-GKL--N-K-LISGGDA-IY-----A---DA----GWRCSLGFNVVKG-S-T--YY-FVTAGHCTDG-
YP--D--WY-T-----NSS-L-T--TWIGPTSGSSFPVNDYGLVRYAN--T-S-VS-HPGT--V-------
GSIDITRAANPVVGATASRRGSTTGTHSGTVTALNQTV-NY-G--GG-------DIVYGMIQTNICAEPGDS-G-
GPMYS-G---------S--TGYGLTSGGS--------GNCSS---------------GGTTFFQPLVEAINAYGVSII
>XX00000076 WP_029126509 32 124 192 MKSTMP------------------------------------
-----------------------------------------------------------------------------------------------------------------------
------------------L-----F-R-RV--GA--VAAA---------------G-TLVA-S--G-----LLG-----AP----------------------------
---------------T-------Q-A---A---------PASPETATNLAERLGD-RTAGIYADH-N-GTMIITVTDETTAR---Q--
VRAEGA-T--PKL-VTR--SAAQLRA-ATTEL-ERS---------------------------------------------------------------------
-------AK------IP-GTAWWTDPTTNQVVLSIDSTI-T----GD--K-LTQ-VKEAAA-RT----N-G-A-VRI---------EA---
------H-----N-GTL--T-T-DIIGGEA-VW-----S----V----SSWCSLGFNVQTSAD-D--YY-FLTAGHCIKD-TG--E--
WY-G-----DSG-R-T--DVIGTVVGKSFPGDDYGLVRYEP--G-K-E--GYGH--VKV--DN-
SWRDLTASAEAYVGQDVMRTGATTGTHSGTVTALNATA-NY-A--AG-------P-VYGLIVTDMCAEGGDS-G-
GPVYA-D---------N--FALALHSGSS-----H-TGACEN---------------DIYSYHQPVVEALAAYDAWVY
>XX00000058 WP_012183279 32 125 187 MRPTRF------------------------------------
-----------------------------------------------------------------------------------------------------------------------
-----------------V-----F-R-RA--AA--VAAA---------------G-TLVV-G--A-----LIG-----SP-----------------------------
-------------A-------Q-A---A-------P-DASPEAAADLAEQLGD-RAAGIYADD-S-GKVVVAVTDSAAAR---Q--
VTRAGA-T--PRM-VKR--GAAELKR-ATAEL-ERS--------------------------------------------------------------------
---------AK------IP-GTAWWVDPETNQVVVSVDSTV-T----GT--K-LAR-VKAAAA-RT----G-G-A-VRI---------
EP---------E-----A-GVL--S-T-QITGGER-IT-----G---AS----GGTCSLGFNVRSG-S-N--YY-FLTAGHCTDV-VS--
S--WY-D-----N-----G--SLLGPTAGSSFPGDDYGIVRLNN--G-Y-E---PGY--VYLY-NG-
GYQDITTAGNAFVGQSVRRSGQTTGLHSGSVTGLNATV-NY-V--EG-------T-VYGLIRTNVCAERGDS-G-
GSLFS-G---------S--TALGLTSGGN--------GNCTF---------------GGTTYFQPVIEALNRYGVDVY
>XX00000288 WP_043967906 32 125 192 MRPTRS------------------------------------
-----------------------------------------------------------------------------------------------------------------------
-----------------T-----L-L-RA--VS--VAAA---------------G-TLVA-T--A-----LIG-----AP------------------------------
------------A-------Q-A---A-------P-AASPDAAATLAERLGD-RSAGTYADA-S-GRMVVTVTDAAAAR---Q--
VRASGA-I--AKL-VKR--GAAELDR-ATAEL-ERS---------------------------------------------------------------------
-------AK------IP-GTAWWTDPVTNQVVVSVDSTV-T----GA--K-LER-VKAAAA-RT----G-G-A-VRI---------
EA---------E-----A-GVL--S-T-RISGGQA-IY-----A---AG----GGRCSLGFNVRSSSG-V--TY-FLTAGHCTNI-AS--
T--WY-S-----NSG-Q-T--SVLGTRAGTSFPGNDYGIVRHSN--S-A-N--ATGN--VSLY-NG-
SYQDVTGAANAFVGQSVQRSGSTTGLRGGTVGALNATV-NY-A--EG-------S-VSGLIRTNVCAEPGDS-G-
GSLFS-G---------T--SAIGLTSGGS--------GNCTR---------------GGTTYFQPVTEALSVYGVSII

>XX00000294 WP_053658538 32 125 191 MRPTRS------------------------------------
-------------------------------------------------------------------------------------------------------------------
----------------S-----F-R-VA--AT--VAVS--------------G-TLVV-G--S-----LIG-----AP-----------------------------
------------A-------Q-A---A-------P-AASADTAAALSAELGD-RSAGSYIDA-S-GKTVVTVTDAAAAR---K--
VSNAGA-S--VKY-VTR--GSAELNR-ATADL-ERS---------------------------------------------------------------
--------AK------IP-GTAWWSDPVTNQVVVSVDSTV-T----GA--K-LER-VKAAAA-RA----N-G-A-VRV---------
EA---------E-----A-GVL--S-T-RISGGQA-IY-----A---GG----GGRCSLGFNVRSG-S-T--YY-FLTAGHCTNI-SA--
S--WY-S-----NSA-Q-T--SLLGSRTGTSFPGNDYGIVRHNN--S-A-N--AAGN--VSLY-NG-
SFRDITGSGNASVGQSVQRSGSTTGLRSGSVTATNATV-NY-A--EG-------S-VSGLIRTNVCAEPGDS-G-
GSLFA-G---------G--TALGLTSGGS--------GNCRT--------------GGTTFFQPVTEALSRYGVSVF
>XX00000303 WP_033669275 32 125 188 MRPTRF------------------------------------
-------------------------------------------------------------------------------------------------------------------
----------------M-----M-R-RA--AA--VAVA--------------G-TLVV-G--A-----LIG-----SP--------------------------
----------------A-------H-A---A-------P-NASPEAAVDLAEQLGD-RAAGVFADA-N-GAVTVAVTDDDAAR---
Q--VAKAGA-T--PKI-VKR--DAAELRR-ATAKL-ESL------------------------------------------------------------
-----------TT------IP-GTAWWTDPETNQVVVSVDSTV-T----GS--K-LAR-VESVVE-RI----G-G-A-VRI---------
ES---------E-----T-GVY--S-T-QISGGEA-IW-----G----G----ESVCSLGFNVRSG-S-N--YY-ALTAGHCTDP-IS--
Y--WY-E-----DSA-H-N--TYLGPTVGSSFPGNDYGIVRLSG--Y---E---PGN--VYLY-NG-
RYQDITSAGNAYIGQSVQRSGRTTGLHSGSVTGLNATV-NY-E--EG-------T-VRGLIRTNVCAQRGDS-G-
GALFS-G---------S--TAVGLTSGGS--------GNCFF--------------GGTTFFQPVVEALNAYGVDVY
>XX00000306 WP_029021503 32 125 187 MRPTRF------------------------------------
-------------------------------------------------------------------------------------------------------------------
----------------V-----F-R-RA--AA--VAAA--------------G-TLVV-G--A-----LIG-----SP-----------------------------
-------------A-------Q-A---A-------P-DASPEAAADLAEQLGD-RAAGIYADD-S-GKVVVAVTDSAAAR---Q--
VTRAGA-T--PRM-VKR--GAAELKR-ATAEL-ERS---------------------------------------------------------------
---------AK------IP-GTAWWVDPETNQVVVSVDSTV-T----GT--K-LAR-VKAAAA-RT----G-G-A-VRI---------
EP---------E-----A-GVL--S-T-QITGGEQ-IT-----G---AS----GGVCSLGFNVRSG-S-N--YY-FLTAGHCTDV-VS--
S--WY-D-----N-----G--SLLGPTAGSSFPGDDYGIVRLNN--G-Y-E---PGY--VYLY-NG-
GYQDITTAGNAFVGQSVRRSGQTTGLHSGSVTGLNATV-NY-A--EG-------T-VYGLIRTNVCAERGDS-G-
GSLFS-G---------S--TALGLTSGGN--------GNCTF--------------GGTTYFQPVIEALNRYGVDVY
>XX00000054 WP_033342256 37 126 189 MSIFR---------------------------------------
-------------------------------------------------------------------------------------------------------------------
------------------------RLHVT--LTRR--------------A-ALAL-T-LP-----AVA-----VA--------------------------------
--------ALLT---PALAQA---G------STPTLDPAGAAALSDRLGD-RSAGAYLDG---DAIVVTVTDDAAAR---T--
AEDAGA-V--ARH-VTR--GAGALAA-VAADL-RAT---------------------------------------------------------------
-----------AA------IP-GTAWAVRPAENLIVVTVDDTV-T----GA--G-LSA-VEAAVA-RS----G-G-A-ARL---------
ER---------V-----P-GRL--G-T-RISGGEA-IY-----G----G----AYRCSLGFTVTDG-V-A--SY-FLTAGHCTEV-SE--
V--WY-A-----DLD-H-T--AQLGTVRGGSFPGDDYGIVRYSA--D-D----HPSA--VDLY-DG-
TTQEITSAGTAVAGQAVQRSGSTTGLHAGTVLEVDATV-IY-P--EG-------T-VSGLIRTDVCAEGGDS-G-
GSLFA-G---------P--VALGLTSGGL--------GDCFT--------------GGTTYFQPVDEVLAVYGVNIP
>XX00000287 WP_043720310 32 126 192 MRPTRS------------------------------------
-------------------------------------------------------------------------------------------------------------------
----------------M-----L-R-RA--VT--VAAA--------------G-TLVA-G--V-----LAG-----TP--------------------------
--------------A-------Q-A---A------PA-SATPDSAAALAERLGA-RAAGAYADA-A-GKMVVAVTDSLAAR---Q--

VASAGA-T--PKL-VKR--GAAELNR-ATAEL-DRS---------------------------------------------------------------------
--------AA------IP-GTAWWVDPATNQVVVSVDSTV-T----GA--K-LER-VKAAAA-RT----N-G-A-VRI---------
EA---------E-----A-GVL--S-T-RISGGEA-IY-----A---QG----GGRCSLGFNVRAG-S-T--YY-FITAGHCTNI-AA--
N--WY-S-----NSS-Q-T--SLLGSRTGSVFPGSDYGIVRYSN--Q-S-VV-QPGN--VSLY-NG-
TFRDITASGNAYVGQSAQRSGSTTGVRSGSVTALNATV-NY-A--EG-------Q-VRGLIRTNICAQPGDS-G-
GSLFS-G---------T--IALGLTSGGS--------GNCSI---------------GGTTYFQPVGPVLSRYGVSVY
>XX00000290  WP_013285478         32        126      191      MRPTRS------------------------------------
-----------------------------------------------------------------------------------------------------------------------------
-----------------T-----L-R-RA--AA--VAVA---------------G-TLVT-G--S-----LLG-----AP-----------------------------
-------------A-------Q-A---A-------PTALSPDAAATLVEKLGA-RAAGTYADA-S-GKMVVTVTDTATAR---Q--
VRAAGA-I--PKI-VAR--GADKLNA-ATSEL-ERS----------------------------------------------------------------------
-------AK------IP-GTAWWTDPATNQVVVSVDSTV-T----GA--K-LER-VKAAAA-RH----N-G-A-IRI---------EA-
--------E-----A-GTL--S-T-RISGGQA-IY-----A---GG----GGRCSLGFNVRSG-S-T--YY-FLTAGHCTNI-SS--S--
WY-S-----NSA-Q-T--SLLGTRTGTSFPGNDYGIVRHNN--S-A-N--AAGN--VSLY-NG-
SFQDITSAGNAYVGQSVRRSGSTTGLRSGSVTALNATV-NY-A--EG-------S-VSGLIRTNVCAEPGDS-G-
GSLFA-G---------T--VALGLTSGGS--------GNCRT---------------GGTTYFQPVTEALSRYGVSVF
>XX00000291  KXK60454       32        126      191      MRPTRS------------------------------------------------
-----------------------------------------------------------------------------------------------------------------------------
-------S-----L-R-RA--AT--VVVA---------------G-TLVA-G--S-----LIG-----AP-----------------------------------------
---A-------Q-A---A-------PAVASPDAAAGLAERLGD-RSAGTYADG-T-GKMVVTVTDAAAAR---Q--
VTANGA-T--PKL-VKR--GAAELAR-ATAEL-DRS---------------------------------------------------------------------
--------AR------IP-GTAWWTDPATNQVVVSVDSTV-T----GA--K-LER-VKAAAA-KS----N-G-A-VRV---------
ES---------T-----P-GVL--S-T-TISGGQA-IY-----A---GG----GGRCSLGFNVRSG-S-T--YY-FLTAGHCTNI-SS--S-
-WY-S-----NSG-Q-T--SLLGTRTGTSFPGNDYGIVRHNN--S-A-N--AAGN--VSLY-NG-
TFRDITAAGNAFVGQSVQRSGSTTGLRSGSVTGLNATV-NY-A--EG-------T-VTGMIRTNVCAQPGDS-G-
GSLFS-G---------T--TALGLTSGGS--------GNCTT---------------GGTTFFQPVTEALSAYGVSVF
>XX00000296  WP_025618159         32        126      189      MRPTRF------------------------------------
-----------------------------------------------------------------------------------------------------------------------------
-----------------T-----L-R-RA--AA--VAAA---------------G-TLVV-G--V-----LVG-----SP----------------------------
--------------A------QA-A---A-------P-NASPEAAADLAEQLGD-RAAGVYAEA-D-GSIVVTVTDAAAAQ---Q--
VSTAGA-T--PKI-VER--GAAELQR-ATAEL-ERS----------------------------------------------------------------------
-------AK------IP-GTAWWTDPATNQVVVSVDSTV-T----GA--K-LAQ-VEAAVA-RS----G-G-A-VRI---------
ES---------E-----A-GVY--S-T-RITGGEA-IY-----G----S----AGVCSLGFNVRSG-S-N--YY-FLTAGHCTDV-IS--N-
-WY-A-----NSS-L-S--NYLGPTAGSSFPGNDYGIVSLSG--G-Y-Q---PGY--VYLY-NG-
NHQDITSAGNAYVGQSVQRSGRTTGLRSGSVTGLNATV-NY-A--EG-------T-VYGLIRTNVCAERGDS-G-
GSLFS-G---------S--TALGLTSGGS--------GNCSF---------------GGTTFFQPVVEALNVYGVNVY
>XX00000300  WP_018794778         32        126      188      MRPTRS------------------------------------
-----------------------------------------------------------------------------------------------------------------------------
-----------------L-----L-C-RA--AS--LAAV---------------G-ALVA-G--T-----LSS-----AP-----------------------------
------------A-------QAS---P-------A-PATPEVAASLSERLGS-RSAGAYANT-D-GTMVVTVTDAAAAR---Q--
VHKAGA-T--PKI-VTR--SASALHH-AATEL-ERS----------------------------------------------------------------------
-------AS------FP-GTAWWTDPATNQVVVSVDSTI-T----GD--K-LNS-VKTVVA-RL----G-G-A-IRL---------ES--
-------A-----A-GVY--S-T-RISGGEA-IY-----A---DR----GGRCSLGFNVRRG-G-T--YY-FLTAGHCTDG-AS--G--
WS-S-----N-----T--ERLGVTAGSSFPGDDYGIVRYDS--N-T-A--QPGN--VYLY-NG-

RYQEITSASDAYIGQRVQRSGSTTGLHSGTVTALNATV-NY-R--EG-------T-VRGLIRTTVCAEPGDS-G-
GSLFA-G---------T--TALGLTSGGS--------GNCFF---------------GGTTFFQPVTEALNRYGVDIY
>XX00000302 WP_052311973 18 126 184 MGS-----------------------------------------
--------------------------------------------------------------------------------------------------------------------------
--------------------------------------------------------------A-----L-V-----AA------------------------------------------
S-------P-A---SAAPALADPSFAPSALAGRIDAQLGT-RDAGSYLDA-S-GKLVVNVTDAATAA---D--VTKAGA-
T--PRY-VAR--SGATLAA-ADASL-KVN---------------------------------------------------------------------------
LK------TP-GTAFAVDPVSNQVVVSVDESV-T----GA--K-LAA-VKATVA-KL----G-D-A-ARL---------ET---------
I-----A-GRT--S-T-RISGGDA-IY-----T----S----SARCSLGFNVRDSAG-T--YY-FLTAGHCTNI-GA--T--WT-N----
-G-----S--STLGTRAGTSFPGNDFGIVRYTN--T-S-VT-KSGA--V-----G--
SQDITSAGTPAVGATVYRRGSTTGIHSGRVTALNSTV-NY-S--EG-------T-VSGLIRTTVCAEPGDS-G-GSLYS-
G---------T--TALGLTSGGS--------GNCTS---------------GGVTYFQPVVEPLSVYGVSVY
>XX00000005 WP_018353253 25 127 187 MESTKS--------------------------------------
--------------------------------------------------------------------------------------------------------------------------
----------------R-----L-SVTL--AA--TL-----------------LAGT------------------------------------------------------------
------------------FAVQAPVQAAPADGSRLAALASGLGEHRTAGVYRDA-S-GQLIVNVTDAAAAS---A--
VRAAGG-Q--ARI-VGR--STHTLNS-LAATL-D--------------------------------------------------------------------
-------PG------VA-GTAWSVDVAGSRLLVELDPAL-S----GQ--R-LAK-VLDAVA-RS----G-G-A-ATT---------
AW---------L-----P-GPI--T-T-LTSGGDA-IY-----G----G----QYRCSLGFNVRSG-T-T--YS-FLTAGHCTDI-AG--
E--WF-S-----DGQ-H-R--TKLGDTAGGSFPGNDYGIVRYTN--T-S-IT-ISGT--V-----G--
GQDIASAAEAFVGESVKRTGSTTGTKSGTVTGLNATV-HY-S--GG-------GTVKGMIRTNVCAEGGDS-G-
GPLYD-G---------S--KALGLTSGGS--------GNCQS---------------GGTTFYQPVTEALGRYNVSIF
>XX00000094 WP_034590401 34 127 192 MSFTKL--------------------------------------
--------------------------------------------------------------------------------------------------------------------------
----------------S-----L-R-HAAVIG--AAAT-----------------AFFF----------ATA-----VA-------------------------------
-----------P------GAAN---A------AAPHATALGAANLVAGLGD-RSAGFYLDQVT-GKPVVTVTDDAAAA---L--
VRSSGA-T--ARL-VRH--SAAALSS-AAGQV-DRT-------------------------------------------------------------------
---------IP------GS-GIAWGVDPASNQVVITVDSTV-T----GA--R-YAQ-VQAAAA-KI----G-D-T-VRV---------
EQ---------T-----P-GVF--S-L-RINGGTA-IY-----G----G----QYRCSLGFNVRSG-S-T--YY-FLTAGHCGNI-AS--
T--WY-A-----NSA-H-T--SLLGTTSGSSFPGNDYAIVKFAS--G-V-A--HPGT--VNLY-SG--
SQTISSSANAYVGEAVKRSGSTTGVHSGTVQQTGATV-RY-A--EG-------T-VTGLIRTNVCAEGGDS-G-
GSLFD-G---------S--IALGLTSGGS--------GNCSS---------------GGTTYFQPVTEALSVYGVTVS
>XX00000188 WP_003070424 36 127 187 MSTAH---------------------------------------
--------------------------------------------------------------------------------------------------------------------------
----------------------PASPVRRGG--TICA--------------A-ALLA----------------------------------------------------
-------------AGTALLPPSAAFAAQPMNVRAATALGGQLTNRLGD-RTVGAYYDH---GHLVVTVTDAAAAA---
D--VRTAGA-E--PRP-VPF--RRSELAA-VTGRL-NRS-----------------------------------------------------------------
------------DR------VP-GTAWRIDPRTGQVRVTADPTV-A----GA--A-LAK-LTAEVR-GF----G-A-K-ARF--------
-ER---------T-------ARL--R-P-LLTGGDA-IW-----G----Q----GLRCSLGFNVHDSSG-N--PG-FITAGHCGVA-
TN--D--WW-A-----DSG-N-Y--EHIASTQRADFPGTDYSYAAYDP--G-A-D--GPSA--VN------
TGQEITGAGDASVGEPVTRSGSTSGVYSGTVTGLDATV-NY-E--EG-------S-VYGLIDTDVCAEPGDS-G-
GALFD-G---------A--TAIGLTSGGS--------GDCTS---------------GGETFFQPVPAVLQAYGLSLP
>XX00000289 WP_043963113 32 127 191 MRLSGS------------------------------------
--------------------------------------------------------------------------------------------------------------------------

----------------P-----L-R-RA--TA--ASVA--------------G-ALVV-G--A-----LTG-----AP----------------------------
-------------A-------Q-A---A------PATPASPDQVTALAAALGD-RSGGAYARP-D-GTMVVTVTDDAAAR---K-
-VRAAGA-T--PKL-VTR--GADDLAR-ATAAL-DSS----------------------------------------------------------------
----------AR------IP-GTAWWTDPATNQVVVSVDSSV-T----GT--K-LQR-VEAVVS-RL----G-G-S-VRI---------
ER---------E-----A-GTL--S-I-RAAGGEA-IW-----A---EG----GGRCSLGFNVRSG-D-T--YY-FLTAGHCTDI-AA--
N--WY-A-----DSG-Q-S--SLLGSGGTGSFPGDDYGIVAHSD--A-A-A--ADGS--VSRY-DG-
TTQDITAAADAFVGQSVQRSGSTTGLHDGTVEAVDATV-NY-V--EG-------S-VTGLIRTTVCAEPGDS-G-
GALFS-G---------S--TALGLTSGGN--------GNCSS--------------GGTTYFQPVTEALSVYGVEIV
>XX00000309 WP_045312430 24 127 191 MKSRL-------------------------------------
-----------------------------------------------------------------------------------------------------------------
-----------------------------------------------------------------------------------------------------------
AALTATLL----LGVALTPSASADVAAATTDLATAVADRLGD-RTAGSYQDSAT-GRLVVAVTDTETAH---A--
VRAAGA-V--PRF-VTR--SGAVLKA-ATEKL-DRS------------------------------------------------------------------
--------AA------IP-GTSWGIDPRTNQVLVSADSTV-T----GE--K-LAK-LRGVLA-GL----G-D-T-ARL---------EH--
-------V-----P-GAF--T-T-TITGGDA-IY-----G----G----PYRCSLGFNVNRPGS-T--AW-FLTAGHCGNA-AA--T--
WS-A-----A---G-G--QVIGTREGSSFPGNDYALIRYTS--G-I-A--RPGA--VNLY-NG-
TTRDISRAADPYVGQSVGRSGSTTGFRTGQVTGLNVTV-NY-S--DG------TT-VYGLIRTTVCAEGGDS-G-
GSLFA-G---------N--TALGLTSGGS--------GNCSS--------------GGTTFFQPVVEALNAYGVRVY
>XX00000090 WP_012852788 29 128 192 MRPVRL-----------------------------------
-----------------------------------------------------------------------------------------------------------------
---------------------------A--AL--AAAA--------------A-TATA-T-TL-----L-T-----AP--------------------------------
----------A-------ATA---A------PAPSDPGRLAATLAERLGS-RSAGSYYDA-R-GKLVVTVSDDAAAQ---T--
VARAGA-L--PRR-VAR--SGAELQR-IMADF-DRS------------------------------------------------------------------
---------ID------VA-GTAWAIQPASNQVVLTLDATV-T----GA--E-LAK-VRAIAA-KY----G-T-A-VRI---------EQ-
--------V-----S-GTF--R-T-TIAGGQA-IY-----A----S----GARCSLGFNVRDNSG-N--LY-FLTAGHCTNI-SS--T--
WY-A-----NSS-G-T--TVLGTRVGTSFPGNDYGIVRYTN--T-G-IA-KPGT--VYLY-GA-
GEQDITSAGNAYIGQSVRRSGSTTGVRSGIVTALNQTV-RY-A--EG-------T-VSGLIRTTVCAEPGDS-G-
GPLFA-G---------T--TALGLTSGGS--------GNCLW--------------GGTTFFQPVTEPLSVYGVSVY
>XX00000248 WP_051818456 40 128 194 MPTQVK-----------------------------------
-----------------------------------------------------------------------------------------------------------------
----------------VKTLRTA-R-RS--GL--IALA--------------S-GLLA-----------V-----SL-------------------------------
-----------TTTTAQ----A---A------PSAKASPEAAAKLTKTLGTTLTAGSYYDAET-KSVVVNVTTRKAAE---A--
VRQAGA-V--PRL-VTY--SSAQLAK-AGAAT-K-A-------------------------------------------------------------------
--------AD------IP-GTAWAVDPRSNKLVVSADSNV-S----RA--D-LAE-LKSATA-SY----G-D-A-VTV---------KR-
--------V-----S-GTF--R-K-LLNGGTA-IY-----A---DG----GWRCSVGFNVRSSSG-A--YY-FLTAGHCTDG-YP--
N--WY-T-----NSG-L-S--TYVGPTIDSSFPTNDYGIVRYDN--T-G-VS-HPGT--VNLY-NG-
TSQDITSAANAYVGESVKRSGSTTGVHGGTVQALNATV-NY-G--GG-------DVVYGLIQTNVCAEPGDS-G-
GSLFD-G---------T--KAIGLTSGGS--------GDCTS--------------GGTTFFQPVTEALSAYGVSVY
>XX00000014 AJT68962 0 129 299 MTFS----------------------------------------------
-----------------------------------------------------------------------------------------------------------------
-----------------------------------------------------------------------------------------------------------------
-------APSAARLATSLKADLGD-KSAGWYLDGAT-GHLVMNVLDKEDAE---R--VASKGA-V--ARV-VRN--
SMTALKA-GTQSL-REN-----------------------------------------------------------------------AS------VP-
GTAWSIDPKTNRIVVLADRTV-T----GQ--K-MAT-LNKATK-GM----G-G-M-VTV---------KR---------S-----A-

GEF--R-----VGGSA-IF-----G----G----NARCSLGFNVTVQ-G-A--PA-FLTAGHCGND-SK--T--WS-A-----
DQG-G-Q--QPLGTVTDSQFPKNDFALVTYDD--A-G-AQ-PQSA--VDLG-NG-
KTQPITKAAEAAVGMKVQRSGSTTGVHDGTVTGLDATV-NY-G--NG-------DIVNGLIQTDVCAEPGDS-G-
GSMFS-G---------D--SAVGLTSGGS--------GDCTQ--------------GGETFFQPVTAALQATGAEIG
>XX00000184  WP_033342257  33  129  185  MRSTSS-------------------------------------
-----------------------------------------------------------------------------------------------------------------------------
---------------S-----I-R-R--------------------SATILAAPAVIVG--A-----L---------------------------------------------
------------IAPT---NAQAAPTLLTEAFADKATTLSDSLGADRTAGTYLDAET-GRMVVTVTDATAAR---A--
VRTQGA-V--AET-VSF--STAQLDA-VIADL-NK-----------------------------------------------------------------------
-------TP------FA-GTSWATDVINNQVVVSIDGTV-S----TA--D-AAV-IKAKVA-KA----G-A-A-ARV---------ET--
-------T-------DKI--S-T-TIAGGAA-IY-----T----G----GSRCSLGFNVRSG-S-T--YY-FLTAGHCTNA-GA--T--
WT-N---------G-S--ATLGTRSGTSFPGNDYGIVRYTN--T-S-IT-HPSA--V-------
GSTTISSAATPAVGTAVTRSGSTTGTRTGSVTALNATV-NY-S--QG-------S-VSGLIRTTVCAEPGDS-G-
GPLYR-G---------S--VAYGLTSGGS--------GNCTS--------------GGVTYFQPVTEALSVYGVSVG
>XX00000206  WP_043827530  22  129  176  MRSTRF-------------------------------------
-----------------------------------------------------------------------------------------------------------------------------
----------------------------------------------------RLSVLAVGSFGTLALAMTTVP-------------------------------------
----------A----------------------ASAAATPAAVMSSLGS-ATGGHYLDD-S-GASVVTVLNNADAA---K--
VRASGL-N--AQV-VKY--SLSQLST-VKQQL-DAT--------------------------------------------------------------------
---------GG------VP-SAGWGLDLAHNQVVVDVYDAT-P----KA--Q-KDA-LLETAR-QH----G-D-M-VRV-------
--EH---------H-----P-GAM--S-L-FIKGGDK-IS-----N----G----QASCSNGFNVEKD-G-K--KL-LLSAGHCEKL-
GGGGP--WN------------------DGKTVDAVFPDEDNMLVENAS--G-E----GPSE--IN---D---
GTKITEFADATVGEEMKRAGITSGVTSGKVTKVDYTF-EA-E--GY-------T-VYHEFCTTAHSDHGDS-G-
GPAYD-G---------S--KGLGTLSGGD---------------------------TQTSCFFPATLSAKRYGVTLP
>XX00000284  WP_026400347  34  129  189  MSKRHG------------------------------------
-----------------------------------------------------------------------------------------------------------------------------
-----------------R-----A-RAAA--AT--AAAA--------------A-GLAA-A--A-----LVT-----AP--------------------------
----------------P---------A------AAAPAAPGAPDQVAAQLAERLGA-RSAGAYLDSST-DKLVVTVTDAAAAR---
T--VRAAGA-Q--PKT-VER--SGADLKE-VMAEL-KRD---------------------------------------------------------------
-------------AT------VP-GTAWAIDPVSNKVVLTMDSTV-K----GA--N-LEK-VRGEAA-QH----G-A-A-VRT-----
----ER---------V-----A-GEF--R-M-FTAGGQA-IY-----A----G----GGRCSLGFNVRSG-S-T--YY-FLTAGHCTAI-
GS--T--WT-D-----GS----G--RTLGRNYASSFPGNDYGVVRYTS--T-P-QD-TRGV--VHLY-GG-
GTQDITRAGNATVGQSVTRSGSTTGVHRGQVTALNQTV-NY-A--EG-------S-VSGLIRTTVCAEPGDS-G-
GSLFS-G---------S--TALGLTSGGS--------GNCTW--------------GGTTFFQPVTEPLSRYGLSVY
>XX00000310  WP_051468057  19  129  189  MTV-----------------------------------------
-----------------------------------------------------------------------------------------------------------------------------
------------------------------------------------------L----------------------------------------------------------------
AAVALT----AAAAPGAHAAQPKPHDPATTARELAKQLGG-RSAGAYVDRAS-HKLVVTVTNDADAR---T--
VRAAGV-E--ART-VGR--DGSALRA-ATARL-GRT--------------------------------------------------------------------
---------AK------VP-GTAWSIDPRTDQVVLSVDRTV-T----GA--R-LAK-VKAAAA-EL----G-P-A-VRI---------
KH---------V-----R-GAF--R-P-LISGGDA-IY-----G----G----NIRCSLGFNVHDG-G-T--DY-FITAGHCGNE-AA--
T--WT-D-----E---S-G--NVLGQTYDSRFPGTDYAIVQYTG--N-V-D--HPSD--VDLY-DG-
TTQPITQAADAYVGESVQRSGSTSGVHGGTVQGLDATV-NY-Q--EG-------T-VYGLIDTDVCAEPGDS-G-
GSLFD-G---------T--SALGLTSGGS--------GDCSS--------------GGETFFQPVGPVLGAYGVDVG

>XX00000285 CNG71536 33 130 188 MKNRHR--------------------------------------------
----------------------------------------------------------------------------------------------------------------------
--------R-------AGAA--AV--TTAT---------------V-GLVA-A--T-----LAA-----AP-------------------------------------
-----A---------A---SASPAAPAAPAAPGTLATSLAQRLGT-QTAGSYLK--G-GKLVVTVTSAQAAQ---A--
VRAAGA-V--PKT-VAR--SGADLAK-VMKAL-ERE-------------------------------------------------------------------
----------AT------VP-GTAWAVDPATNQVLLTMDSTV-K----GA--K-LTK-VRAAVA-KQ----G-A-A-VRT--------
-ER---------V-----A-GQF--Q-K-FTRGGEA-IY-----T----G----GARCSLGFNVRSG-S-T--YY-FVTAGHCTNI-GT-
-T--WT-D-----SS----G--RTLGRTVASSFPGNDYGVVQYTS--T-P-TD-TQGS--VSLY-SG--
TRDITSAGNAAVGQTVYRSGSTTGLHSGQVTALNQRV-NY-A--EG-------S-VSGLIRTTVCAEPGDS-G-
GSLFA-G---------S--TALGLTSGGS--------GNCSW---------------GGTTFFQPVTEPLSRYGLSVY
>XX00000000 WP_014141698 35 131 187 MRIKRV--------------------------------------
----------------------------------------------------------------------------------------------------------------------
----------------TPPAATG-R-RL--RL--LAVA---------------C-GLVA-A-TA-----V-S-----LP----------------------------
---------------AA---D----A---A------PAARATPDAAAGVARTLGAAATAGTYYDAAA-KATVVNVTSDAAAR---
Q--VRAAGA-V--PRL-VKY--SSAQLDT-AGAAV-Q-R------------------------------------------------------------
-------------LG------IA-GTAWAQDPRTDQLVVSADHGV-P----AG--R-YAA-LRTTAA-RY----G-D-A-VRV------
---QR---------V-----A-GRF--R-E-LLSGGDA-IY-----G----G----GYRCSLGFNVHSG-G-T--YY-FLTAGHCGKA-
VS--T--WY-T-----SSG-Q-S--TVIGPTTGYTFPGNDYALVRYSN--T-S-LP-HPSA--V-------
GGQTITGAGNAYVGESVTRRGSTTGVHSGTVTGLNATV-NY-G--SD-------GIVRGLIQTNVCAEPGDS-G-
GSLYS-G---------S--TAVGLTSGGS--------GDCTS---------------GGTTFFQPVTAALSHFGVSLP
>XX00000068 WP_020662213 30 131 177 MTSRKT--------------------------------------
-----------------LVAGITT-LSA-----------------------------------A-------A-----L--------------------------------
TAGMCVNAASASPLAFAQMQ-------------------------------------------------------------------------------
----------------------------------------------------AQAITQATQVADSLGA-GNGGIYLE--N-
GKAVVNVTDAVAQE---K--AKAAGF-M--TKQ-VKY--SFTALTN-AKNQM-DAV-----------------------------------
------------------------------------------KN------VP-QTSWGIDTKNNQVVVKIYDSA-S----QA--A-KDK-
VAAAAA-KL----G-D-A-ARV---------EH---------R-----S-GKL--A-T-YIADGDS-IN-----N----G----
QFTCSLGFNVTKG-G-Q--PY-MLTAGHCTNE-GG--T--WS-G-----G-----D--
VSGAQVVKSDCPGADSGLLTRPN--G-T----GDGA--IN---T---
GQKISSAGTPTVGEQMQKMGQTTGGGSGQVTSVDESV-NF-D--VG-------V-LNHEFGTTAHTDHGDS-G-
GPAYD-G---------S--KGLGTLSGGD---------------------------TQTSYFYPLTLELQSYGLELA
>XX00000273 WP_030624624 33 131 189 MKHRRI--------------------------------------
----------------------------------------------------------------------------------------------------------------------
----------------P-----K-R-RA--AL--AGAG---------------VLALA------------------A------------------------------------
-----SATFALSAN----A------SPAPEPLSLTPAAAGKLAAQLTD-GTAGSYYDAKA-KKLVVNVVDEAAAD---T--
VRAKGA-E--PRV-VQH--TLAQLND-ARQTL-KSK------------------------------------------------------------------
----------AT------IP-GTSWGIDPRSNKVVVTADRTV-K----GA--K-LAK-LTKVVD-GL----G-S-K-VQL---------
HR---------S-----K-GEF--K-P-FIAGGDA-IW-----G----S----GSRCSLGFNVVKD-G-Q--PY-FLTAGHCGNP-VK-
-S--WS-D-----SQ--G-G--GEIGATEESSFPGNDYALVKYTA--D-T-D--HPSA--VDLY-NG-
STQEISGAAEATVGEKVQRSGSTTQVHDGTVKALNASV-NY-Q--EG-------T-VDGLIQTDVCAEPGDS-G-
GALFA-G---------D--KALGLTSGGS--------GDCSS---------------GGETFFQPVTEALEAYGAQIG
>XX00000277 WP_030636241 34 131 189 MKHRRI--------------------------------------
----------------------------------------------------------------------------------------------------------------------
----------------P-----Q-R-RA--VL--AGAG---------------ALALA------------------T------------------------------------

-----TATLSLASAN----A------ADGPAVKQLSPAAATALASQLSG-HAAGSYYDARA-KKLVVNVVDRKAAD---
A--VRAKGA-E--ARM-VRH--SLAQLDA-ARATL-KQR-----------------------------------------------------------
-------------AT------IP-GTAWAMDPRANKVVVTADRTV-S----GA--K-LDK-LNKVVK-SL----G-S-K-AEL-------
--RR---------T-----A-AKF--T-P-FIAGGDA-IW-----G----S----SSRCSLGFNVTKG-G-Q--PY-FLTAGHCGNA-
VK--S--WS-D-----KQ--G-G--QEIAGTEDSKFPGNDYAIAKYTA--D-T-A--HPSE--VNTY-GG-
APQKITKAAEATVGQKVKRSGSTTHVHDGTVKALDASV-NY-Q--EG-------T-VEGLIQTDVCAEPGDS-G-
GALYD-G---------E--SALGLTSGGS--------GNCSS--------------GGETYFQPVPAALKAVGAQIG
>XX00000279 WP_055588229 33 131 190 MRNSRT----------------------------------
---------------------------------------------------------------------------------------------------------------------
----------------H-----G-R-RT--AL--LATA--------------A-AVLT-A-TA-----L-A-----LP------------------------
------------QA---QAAEAQ---A------LSPQQVSTLSAKVQQQLGD-ATAGSYVR--N-GRLVVTVTDSASAA---T-
-VRSEGA-V--PQL-VTR--SGAQLKA-ATDTL-DRT---------------------------------------------------------------
----------AR------IP-GTAWAVDPSTNQVLISADSTV-K----GA--K-LAR-LQKAAK-AL----G-A-S-VRV---------
EN---------V-----P-GTL--S-L-RLSGGQA-IY-----G----G----GYRCSLGFNVKAG-S-T--YY-FLTAGHCGNV-AS--
T--WY-S-----NSS-Q-S--TRIGTTYDSWFPGNDFALVQYTT--S-G-T--PAGN--VSLY-NG-
SYQDIASAGTPSVGQTVYRSGSTTGVHSGTVTALNSTV-NY-A--EG-------T-VSGLIRTTVCAEGGDS-G-
GSLYA-G---------T--VAYGLTSGGS--------GNCTS--------------GGVTYFQPVTEALSYYGVSVY
>XX00000304 WP_030449912 25 131 189 MRPTRT----------------------------------
---------------------------------------------------------------------------------------------------------------------
----------------S-----F-R----LAA--VAAA----------------ALLT----------TGA-----VA-------------------------------
----------T------TAAQ---A------APKTPDRAGVAALVDGLGT-HTAGSYLDH---GTQIVTVTDATSAA---K--
VRTAGL-T--PKL-VRY--STAHLHS-ITAEL-GEK-------------------------------------------------------------------
------AT------IP-GTAWAIDPRNDRVNVTADSTV-S----KA--D-LAK-VRQIAG-QY----G-A-A-AHV---------RV-
--------V-----S-GTF--S-T-KIAGGDA-IY-----G----G----GYRCSLGFNVTDG-S-T--AY-FLTAGHCGNV-AD--T--
WY-S-----DSG-S-S--NEIGTTQDSQFPGNDFAIVKYDS--G-V-S--HPST--VDLY-GG--
TQNITGAANAQVGESVKRSGSTTQVHSGSVTGLDATV-NY-Q--EG-------T-VSGLIDTNVCAEGGDS-G-
GPLFD-G---------S--TALGLTSGGS--------GDCSS--------------GGETFFQPVTEPLSTYHVQIG
>XX00000308 WP_018537098 23 131 189 ML-----------------------------------------
---------------------------------------------------------------------------------------------------------------------
------------------------------AGAG--------------ALALA------------------A-------------------------------------
TATVTLANAH----A------APAPTVASLSPAAATTLASQLKS-GTAGAFYDAQD-RKLVVNVVDEASAA---A--
VRAKGA-E--ARI-VKH--SMAQLDA-ARQTL-KDR---------------------------------------------------------------
----------AT------IP-GTAWAMDPRANKVVVSADRTV-T----GA--K-LDR-LTKVAR-SL----G-D-T-VEL--------
RR---------T-----Q-GEF--K-P-LIAGGDA-IW-----G----S----SARCSLGFNVTKG-G-Q--PY-ILTAGHCGNA-VK-
-E--WS-D-----QQ--G-G--QTIATTEDSKFPGNDYSIAKYTG--N-T-D--HPSE--VDLY-NG-
STQKITKAADATVGEKVKRSGSTTQVHDGTVKALNASV-NY-Q--EG-------T-VNGLIQTDVCAEPGDS-G-
GALFD-G---------E--SALGLTSGGS--------GDCSQ--------------GGETFFQPVPAALQATGTQIG
>XX00000032 WP_030449650 28 132 174 MHRRL------------------------------------
---------------------------------------------------------------------------------------------------------------------
-------------------------------------------------AGLAILGVGVLAAAGIGSTPAQAAPPL-------------------------
----------SAAV-Q---------------------AAAVPQANTVAQRLGS-QNAGVYLDR-N-GHAVVTVTSAAAAR---T--
VRAAGL-R--SRV-VHY--SAASLTS-TKNRL-DRL-----------------------------------------------------------------
-------AG------VP-DTAWGIDTAHNQVVVTISDAA-P----KA--G-AAR-VLAAAQ-RY----G-S-E-VRV---------
EH---------T-----T-GHF--S-T-YVRGGDA-IQ-----N----S----QARCSDGFNVRRN-G-Q--LM-VLTAGHCTNL-

GG--T--WS-P-----M----------GGQVVASNSPGGDEGLITNPS--G-N----GPSQ--IN---T---
GQTISRIGQPTQGEQVTKSGSTTGVTGGTIEAVEQTV-NF-D--VG-------V-IYHLFATDVYSDHGDS-G-
GPGYD-G---------S--TGLGTLTGGD---------------------------TQTTFFYPAWREFNDYGLTLP
>XX00000089 WP_037790920 42 132 184 MEER----------------------------------------
------------------------------------------------------------------------------------------------------------------------
---------VKVRPPV----STQTVKRGKGI--AALA--------------T-GLVA-A-VV-----L-T-----SP-----------------------
------------------TA---T----A-A----PASAPSERATPKAAMAVADTLGS-AQTGGAYKADS-
GRMVVTITDKADAA---K--VEKAGA-V--AKV-VEH--SQAELAR-AAKAI-EHQ------------------------------------
----------------------------------------AD------IK-GTAWSVDPVANTLSVAADSTV-D----AK--E-MAT-
LRKVAA-RF----D-G-A-VTI---------ER---------T-----K-GEF--T-K-LIRGGDA-IY-----M---ST----
GGRCSAGFNARIG-S-T--YY-VITAGHCTEG-YP--N--FS-----------------GIGPTAASSFPGDDYGVIRNDS--S-S-
---TPGV--VNLY-NG-STRDITGVGNAYVNQYVQRSGSTTGLHSGYVTGLNATV-NY-G--GG-------
DVVYGMIRTNVCAEPGDS-G-GSLFA-G---------N--TALGMTSGGS--------GNCSS--------------
GGTTFFQPVTEVTNRYGLVVY
>XX00000276 WP_030285084 34 132 189 MKHRRI--------------------------------------
------------------------------------------------------------------------------------------------------------------------
----------------P-----Q-R-RA--VL--AGAG--------------ALALA------------------A----------------------------------
-----TATLTLNSAN----A------ASVPQPTKLSPAAATTLASQVNA-GTAGSYYDAQA-KKLVVNVVNDAAAK---
T--VRAKGA-E--ARM-VKY--SMAQLDS-ARRTL-KAD---------------------------------------------------------------
--------------AT------IP-GTSWGVDPRTNKVVVSADRTV-K----GD--A-LAK-ITKVTK-GL----G-D-K-VTF--------
-RR---------T-----Q-GEL--K-P-LIAGGDA-IW-----G----S----SARCSLGFNVVKG-G-Q--PY-FLTAGHCGNA-
VK--S--WS-D-----QQ--G-G--QEIATTEDSTFPGNDFAIVKYTG--D-T-E--HPSE--VDLY-SG-
SPQKITKAADATVGEKVKRSGSTTQVHDGTVKALNATV-NY-Q--EG-------S-VEGLIQTDVCAEPGDS-G-
GALFD-G---------E--SALGLTSGGS--------GDCSQ---------------GGETFFQPVPAALQKFGAEIG
>XX00000278 WP_030356732 29 132 189 MRRTTC-------------------------------------
------------------------------------------------------------------------------------------------------------------------
-----------------A----------R--LG--LS-------------------A------------------LLVT-------------------------------------
G-----SLALG----VTPAGAQSEPAPSATRLNALNAQVERQLGD-DSAGTYLDRRT-GNLVVTVTSDAAAE---R--
VRATGA-T--PQR-VER--SAAQLDA-AMDTL-ESR---------------------------------------------------------------
----------AK------IT-GTSWGIDPRTNQIAVEADRSV-S----TR--E-MAR-LHRVAD-SL----D-G-A-VRV---------
SR---------V-----P-GTF--H-R-EVAGGDA-IY-----G----G----GSRCSAAFNVSKG-T-T--KY-FVTAGHCTNL-SA--
N--WS-A-----TS--G-G--AAVGVREGTSFPTNDYGIVRYTN--G-S-S--PAGN--VNLY-NG-
SYQEISSAADAVVGQAIKKSGSTTKVTSGSVTAVNVTV-NY-G--DG-------P-VHGMVRTTACSAGGDS-G-
GAHFA-G---------T--VALGIHSGSS--------GCTGT---------------NGSAIHQPVREALSAYGVSVY
>XX00000275 WP_026414926 32 133 190 MRTPHS-------------------------------------
------------------------------------------------------------------------------------------------------------------------
-----------------R-----A-R-RV--AL--VGSA--------------A-ALLT-A--A-----L-S-----TP------------------------------
------------AA------H-A---SATSPAPSPADRQNRLADSLARTLGD-RSAGSYLDA-S-GKLVVTVSDAASAN---A--
VRAAGA-Q--PRT-VAR--TGAVLNR-AMDAL-KRD---------------------------------------------------------------
-----------AT------VP-GTAWAVDPKTNQVVLSVDSTV-T----GA--K-LAK-VKKAAA-KQ----G-T-A-VRT--------
-ER---------V-----A-GAF--R-T-FTAGGDA-IY-----T----G----NYRCSLGFNVKVG-G-A--YY-FLTAGHCTNL-GS-
-T--WY-S-----NSA-H-T--AVLGTRAGSTFPGHDYGIVKYSS--T-P-TD-TTGV--VDTY-GG--
SQNITSAANPTVGQSVRRSGSTTHVHSGTVTALNQTV-NY-A--EG-------T-VTGMIRTTVCAEGGDS-G-
GSLYS-G---------T--TALGLTSGGS--------GNCTS--------------GGTTFFQPVTGALSAYGASVY

>XX00000075 WP_018635935 0 134 209 M--------------------------------------------------------------------------------------------------------------
------------------------------------------------------------------------------------------------------------------------------------
------------------------------------------------------------------------------------------------------------------------------------
--------------------------RGKLGS-SFGGSWFDAQS-GRLVVGVTSEASAA---Q--VTELGA-T--PKV-VAR--
SYAALES-IATEL-DTLAGRAPEKAGTRTAAGKPQ------------------------------------------------------------AA-----
-VQGLAGWHIDPTTNSVVVSVGRDR-L----SR--R-----AQENLT-RY----G-D-A-VRT---------EY---------L-----P-
AAPTTA-D-YMDGGDQ-IN---------------GGSCSAGFNLRNPTG-K--GY-LLTAGHCVSL-NQ--T--VT-G-----Q-
--G-G--SAFGKVVSRWFLTYDDALVEATN--P-G-YWIQGPWVDTNPS-NG-
GFINITGYTDAPVGTYVCKSGVKTKMTCGNITGKDETV-TF-D--GV------NT-VYGLTRRSACVEKGDS-G-
GANYA-P------GAPN--RAEGVTSGAVLY---GTGLRCGSA-----------IGQPTISWHFPIADSIVAYGANLW
>XX00000147 WP_018381585 0 134 191 MHG----------------------------------------------------------------------------------------------------------
------------------------------------------------------------------------------------------------------------------------------------
------------------------------------------------------------------------------------------------------------------------------------
-ATA---A------PADELEVRRVATLESLLGP-ESTAGYYRS-V--GRMVVNVTSSTAAE---Q--VRASGA-T--PRV-
VRR--SGVELAA-ALSTL-EKS--------------------------------------------------------------------------------AR------IP-
GTAWAADPQSNQVVVRADETV-S----DT--E-LAQ-LNGVIT-TL----G-D-A-ARL---------ER---------H-----R-
GRF--S-P-YLAGGEP-IY-----A----G----NIRCSAGFNVHRG-N-R--HF-TLTAGHCTAG-RG--K--WF-T-----
DPR-G-Q--EELGDTVGSSFPGNDYGLLERTT--A-W-VP-RPGD--VSLH-DG-
RSQKIDKVGTAFVGQSVQRSGSTTGVRGGHVISVGTTV-NY-P--QG-------T-VTGLVGTDACAEPGDS-G-
GPFFH-G---------S--QALGLTSGGN--------GDCAS---------------GGITYFQPVDEALNAYDVSLN
>XX00000256 WP_053656779 36 134 190 MAQRRT---------------------------------------------------------------------------------------------------
------------------------------------------------------------------------------------------------------------------------------------
------------------A-----L-T-TR--AL--TGAS---------------VAVLAA------------G-----AAL-----------------------------
--------LPGAGPATAAP----S------PAAATPAAARAASAATLVARLGD-RAAGSYYDAAA-
HRFVVNITDPSAAG---Q--VRATGA-V--PRT-VRY--STPQLRG-VLGTL-AQH----------------------------------------
----------------------------------------AR------IA-GTAWSIDPRQDKVVVTVDPTV-A----GA--R-RTR-LNGVVD-
SL----G-D-K-VVV---------RR---------T-----A-SAF--K-P-FIAGGDA-IW-----G----S----TVRCSLGFNVTVN-G-
A--PY-FLTAGHCGNA-SS--T--WS-D-----SQ--G-G--GEVGQTVDSQFPGTDYALVQYDD--A-GSD--HASV--
VDLY-GG-GSQQITHAADAYVGESVQRSGSTSGVHGGSVTGLDATV-NY-E--EG-------S-
VSGLIDTDVCAEPGDS-G-GALFD-S---------D--AAIGLTSGGS--------GDCTS---------------
GGETFFQPVPAALSAEGATLP
>XX00000268 WP_030067877 34 134 189 MKHRRI-----------------------------------------------------------------------------------------------------
------------------------------------------------------------------------------------------------------------------------------------
----------------P-----H-R-RA--VL--AGAG---------------ALALA------------------ATA-------------------------------
-----TVTLANAHAAP----A------PTIPKPLKLTSAAATALASQLKS-NAAGAYYDARA-KKLVVNVVDETAAK---
A--VRAKGA-Q--ARL-VKH--SLAQLDA-ARQTL-KSR--------------------------------------------------------------
-------------AT------IP-GTAWGVDPVSNKVVVTADRTV-K----GA--K-LDR-LTKVAR-SL----G-D-K-VEL--------
-RH---------S-----Q-GEF--K-P-FIGGGDA-IW-----G----D----SARCSLGFNVVKD-G-Q--PY-FLTAGHCGNA-
VK--S--WS-D-----TQ--G-G--SEIATTEASTFPGHDYSIVKYTG--N-T-D--HPSE--VDLY-NG-
STQKITQAADATVGEKVQRSGSTTQVHDGTVKALNASV-NY-Q--EG-------T-VDGLIQTDVCAEPGDS-G-
GALFD-G---------E--SAVGLTSGGS--------GDCTN---------------GGETFFQPVPEALSAYGAQIG
>XX00000008 CEJ82719 0 135 171 MK-------------------------------------------------------------------------------------------------------------
------------------------------------------------------------------------------------------------
RDLGHNAEQAVARVAHD----------------------------------------------------------------------------------------

---------------------------------------------FHASRLIEKLRGSVGP-SFAG------------------AD---T--ISAQGA-
V--PVI-MTT--PLSKFQE-AKSAL-DS-IFLGHNRSKRS------------------------------------------------------------
--TN------DA-VIGFYVDEAANKVIFKILAHG-R----AQ--A--ED-LAKQAG-VS----A-S-E-FGV---------KI---------
E-----D-EIPTLM-S-SVRGGDI-YHI----D----S----KHAYSVGFSVNG--------G-FISAAHCVKK-GI--N--VT-D-----
I---E-G--NLLGTAVESSFGGEDSSYIKTID--G-T-N--LTGY--VNSY------------------------
GQAHFHCGTITAKNVTI-DF-D--FR------HP-STGLTETDVCAEPGDS-G-GSFFS-G---------D--
QAQGTLSGGS--------GECAR--------------NGQQYFPPINKSLEAFNLTLI
>XX00000015  WP_033265882  30  135  190  MSPST------------------------------------
---------------------------------------------------------------------------------------------------------------------
----------------------P-RT--VL--AGAG--------------AVVLIT-T--A-----L-A-----LP------------------------------
---------HA---T----A-A-PRPVPDPPTAATAAQRAKQIGSTLGN-EGAGSYYDAEN-HKLIVNVTSESAAA---K--
ARGAGA-D--VKI-VKH--SLASLDA-ARATL-EQH---------------------------------------------------------------
---------AS------IP-GTSWAMDPRSNKVIVTADRTV-R----GD--R-LER-LREVTS-SL----G-D-R-AAL---------RL-
--------S-----S-GTL--R-P-LLAGGDA-IW-----G----A----TARCSLGFNVTKA-G-Q--PY-FLTAGHCTHA-VR--S--
WS-A-----TQ--G-G--PETAVSEGGSFPGDDFGIVKYTA--A-DMV--HPGE--VNLY-NG-
SMQRITGVGEAIVGQRVRRSGSTSHVHDGEVLAVDVTA-NY-Q--QG-------P-VEGLIQTSVCAEAGDS-G-
GPLFE-D---------A--AALGLTSGGR--------GDCTS--------------GGESFYQPVGEALARTGTQLG
>XX00000037  WP_030728132  33  135  189  MTNRRM---------------------------------
---------------------------------------------------------------------------------------------------------------------
----------------S-----A-R-RI--TM--AATG--------------VAALLA-G--T-----F-T-----MS-------------------------
--------------QA---N----A-QTPTFTPDVLSSQAAVELATTLEDTLSA-NTAGTFYDAQT-ETLVVNVTDTAAVS--
-E--VQATGA-E--ARL-VDH--TLAQLEA-AADEV-EE------------------------------------------------------------
------------LA------VP-GTAWGIDPVSNAVKVTVDSTV-T----GD--A-LAE-LRSGVE-AL----G-G-Q-AVL--------
-EQ---------V-----E-GEF--S-P-YIAGGDA-IY-----T----S----GARCSLGFNVTIG-G-Q--RG-FLTAGHCGST-GS--
S--WS-A-----TS--G-G--AAFGTLSTSVFPGSDYAAGRYSS--S-I-A--SPSS--VNLY-NG-
STRSITGAANPVVGQTVERSGSTTGRHSGTVTALNVSV-TY-P--QG-------T-VRGTIQTTVCAEPGDS-G-
GALFS-G---------N--TAHGLTSGGS--------GNCSS--------------GGTTFFQPVVAALNAVGATIP
>XX00000080  WP_014627479  48  135  191  MTKRR------------------------------------
---------------------------------------------------------------------------------------------------------------------
--AFVRSAVPTVRRAGP-----ARS-GA--AV--LA------------------V------------------AAVA----------------------
------------LPAALLPAAHP----ARGAAPPEPTPVTQAAALKIASDLTARLGD-RAAGVYYDATA-
RQLVVNITDPANAD---T--VRAAGA-V--PRT-VKF--TTAQLHQ-ATVTL-GEQ----------------------------------
--------------------------------------AR------IP-GTAWAVDPRRNKVVITADPTV-T----GA--R-RTQ-LDQAVA-
QL----G-E-K-AVL---------RQ---------G-----K-AVF--R-P-FIAGGDA-IY-----G----G----VYRCSLGFNVTKG-G-
A--SY-FLTAGHCGNV-AA--S--WS-D-----AQ--A-G--QPFATTVDSRFPGTDFALVKYDD--PQT-A--HPST--
VDLY-GG-GTQQITQAAEATVGERVRRSGSTSQVHEGQVTGLDATV-NY-Q--EG-------T-
VTGLIDTNVCAEPGDS-G-GSLFD-G---------A--SAIGLTSGGS--------GDCTA--------------
GGETFYQPVPAALRAEGAQLP
>XX00000104  WP_004948276  33  135  189  MTTRRT----------------------------------
---------------------------------------------------------------------------------------------------------------------
---------------R-----A-R-HA--AL--AATA--------------AAVLTA-T--A-----L-------GA---------------------------
------------QG---A----G-ADPAPDPAPLTDTQAGALAHRITHDLGD-RAAGAYFDAAT-
GKLVVNVVDAKAAA---E--VRAAGA-E--PRT-VTR--SAARLAA-ATKTL-DAQ----------------------------------
--------------------------------------AR------IP-GTAWAVDPRTNRVVVKADSTV-R----GE--R-LAR-LDKVVT-

SL----G-D-A-ATL---------RR---------V-----P-GAF--R-P-TLAGGDA-IW-----A----P----RARCSLGFNVTKN-G-
Q--PH-FLTAGHCGNI-SS--T--WS-D-----RQ--G-G--AAIGRTVTSVFPGHDYALVRYTA--N-I-A--HPSA--
VDLY-DG-GQQTITGAAEAAVGQSVQRSGSTTGVYGGQVTALDATV-NY-Q--EG-------S-
VTGMIDTTVCAEPGDS-G-GALFA-D---------S--SAVGLTSGGS--------GDCSS--------------
GGETFFQPVTDALSALGAQLP
>XX00000155 WP_028436376 23 135 189 MVI-------------------------------------------
--------------------------------------------------------------------------------------------------------------------------
------------------------------AGAG--------------VAALVA-G--S-----VTL-----ST-----------------------------------
------AN---A----A-EDSAPSLKTLTSASASKLATTLVSDVK--GDAGSYYDAKA-KKLVVNVTDQDAKK---Q--
VEAAGA-E--AKI-VQH--TMSQLKS-VKDAF-TA-------------------------------------------------------------------
--------KD------IA-GTARMVDVKSNKVVITADRTV-K----GA--K-LAQ-LKKQAK-AE----G-S-K-VEL---------NR-
--------T-----A-GKY--T-T-KIAGGDA-IH-----G----D----GGRCSLGFNVTLD-G-Q--PA-FLTAGHCTTA-IA--S--
WS-D-----SE--G-G--QEIAKSGDGEFPGADYGIASYTA--E-V-D--HPSS--VNLY-DG-
GTQEITGAAEAEVGQQVTRSGSTTQVHEGTVKAVDASV-TY-P--EG-------T-VNGLIQTDVCAEPGDS-G-
GSLFA-G---------D--KALGMTSGGS--------GDCSG---------------GGETFFQPVTTALEATGAQLP
>XX00000222 WP_037742097 33 135 190 MRYRAI--------------------------------------
--------------------------------------------------------------------------------------------------------------------------
---------------S-----K-S-RY--AI--AGVG--------------AAALIA-G--S-----L-T-----LA--------------------------------
----------TA---N----A-TPSEPVADTLSSTAASELAGTLVQDLKG-GAAGAYYDAQA-KELVVNVVNDEAAE---
Q--AEAAGA-R--ART-VDN--TTKELRA-AKADL-TQ---------------------------------------------------------------
-------------GA------TA-GTSRALDPVTNQVVVTADSTV-K----GA--A-LAQ-LKKEVA-AQ----D-G-K-AVL-------
--KR---------S-----E-GTF--K-P-YISGGDA-IY-----T----G----GSRCSLGFNVTKG-G-E--PY-FLTAGHCVEL-
GGS-S--WS-E-----TS--G-G--PVIGEAEGATFPGNDRALVKYTA--D-V-E--HPSE--VNTY-DG-
SAQEITGAREATVGETVQRSGSTTQLHDGTVVALDVSV-TY-P--EG-------T-VDGLIQTDVCAEPGDS-G-
GSLFA-G---------S--DALGLTSGGS--------GDCTN---------------GGETFFEPVVAALDEYGATIP
>XX00000231 CCB73658 22 135 191 ML----------------------------------------------------
--------------------------------------------------------------------------------------------------------------------------
----------------------A-------------------V------------------AAVA------------------------------------
LPAALLPAAHP----ARGAAPPEPTPVTQAAALKIASDLTARLGD-RAAGVYYDATA-RQLVVNITDPANAD---
T--VRAAGA-V--PRT-VKF--TTAQLHQ-ATVTL-GEQ------------------------------------------------------------
-------------AR------IP-GTAWAVDPRRNKVVITADPTV-T----GA--R-RTQ-LDQAVA-QL----G-E-K-AVL-------
--RQ---------G-----K-AVF--R-P-FIAGGDA-IY-----G----G----VYRCSLGFNVTKG-G-A--SY-FLTAGHCGNV-
AA--S--WS-D-----AQ--A-G--QPFATTVDSRFPGTDFALVKYDD--PQT-A--HPST--VDLY-GG-
GTQQITQAAEATVGERVRRSGSTSQVHEGQVTGLDATV-NY-Q--EG-------T-VTGLIDTNVCAEPGDS-G-
GSLFD-G---------A--SAIGLTSGGS--------GDCTA--------------GGETFYQPVPAALRAEGAQLP
>XX00000271 KUJ41359 33 135 188 MKRTTV-------------------------------------------
--------------------------------------------------------------------------------------------------------------------------
-------R-----M-R-RT--AA--AGAA--------------VVALTT-A--T-----L-T-----PQ-------------------------------------
-----SA---D----A-A-PAADPDPMSAAAAGQLAENLHSDLGD-DAAGAYYDPGR-RTLVVNVMSEEAGE---K--
ARAAGA-E--AKM-VRH--SLASLEE-ARRTL-KDK------------------------------------------------------------------
---------AA------IP-GTSWAMNPKLNQVVVTADSTV-T----GA--D-LAR-IDEVVT-SL----G-D-R-AAM---------
RR---------S-----A-TPL--R-P-LISGGDA-IW-----G----S----GARCSLGFNVTKG-G-E--PY-FLTAGHCTNM-VD-
-S--WS-D-----TQ--G-G--TQVAATAGSSFPGDDYGIAAYTT--T-I-A--HPSA--VDLY-SG--

SQQITQAGDPIVGQQVQRSGSTTHVHSGDVTGLDATV-NY-Q--EG-------T-VEGLIQTDVCAEPGDS-G-
GSLYS-G---------T--TALGLTSGGS--------GNCTS---------------GGETFFQPVTEALRATGSQIG
>XX00000272 WP_030886733 33 135 188 MKHRRM-----------------------------------
--------------------------------------------------------------------------------------------------------------------------
------------------P-----K-R-RV--AL--AGAG---------------VAVLAA-A--S-----L-T-----LQ---------------------------
---------------NA---N----A-A-PGPTPDTLSVTSAGKLAGELSAGLKG-DAAGSYYDAKA-KKLVVNVLNEKAAD-
--Q--VRRAGA-E--PKV-VKH--SLAQLES-ARRTL-DAD-----------------------------------------------------------
--------------AA------VP-GTAWATDPKTNQVVVTADRTV-K----GA--A-LSK-VSKVVS-SL----G-D-K-AAL-----
----KR---------S-----A-GEF--R-P-LIAGGDA-IW-----G----S----SARCSLGFNVTKG-G-Q--PY-FLTAGHCGNA-
VK--S--WS-D-----QQ--G-G--QEIATTEASTFPGRDYSLAKYTG--Q-T-D--HPSE--VNLY-GG--
TQKITKAAEATVGEQVKRSGSTTQVHDGTVKALNATV-NY-Q--EG-------Q-VDGLIQTDVCAEPGDS-G-
GALFD-G---------E--SALGLTSGGS--------GDCSQ---------------GGETFFQPVPEALKEYGAEIG
>XX00000086 WP_018657892 26 136 191 MKYRSA------------------------------------
--------------------------------------------------------------------------------------------------------------------------
-----------------R-----L--RGP--AA--IVAA---------------T-GLAL-T--A-----LAA-----TP------------------------------
----------------------G---AASAASPAAPADPARVAAALTAQLGA-RSAGTYLD--G-GALVVTVTDSGAAA---A--
VRAAGA-R--PRT-VAR--GGDDLKK-AEAAL-KRR-----------------------------------------------------------------
----------AS------VP-GSAWAVDPAANKVVVTLDATV-T----GA--R-LAQ-VRKTAA-AL----G-D-A-VHV--------
-RQ---------T-----K-GTY--R-T-YTAGGEA-IY-----A----G----TGRCSLGFNVEKG-G-E--YY-FLTAGHCTDG-RG-
-A--WY-A-----DAA-N-S--SKLGDTAGSDFPGTDYGLVKYSS--A-P-AD-TRGV--VSLY-GG--
TQDITGAGSATVGQSVTRSGSTTQVHTGTVTALNKTV-NY-D--DG------AT-VSGLIETTVCAEPGDS-G-
GPLFT-G---------P--TALGLTSGGS--------GDCTL---------------GGTTFFQPVGPALSAYGVTLY
>XX00000153 WP_054055370 28 136 198 MSRRSL--------S---------------------------
--------------TTLSLACAAVS-IAALAT-------------------------------P--------------------A-----------------------------
--AAAPSALDAAADP------------------------------------------------------------------------------------------
-----------------------------------------ARLATAEQTLREQLGG-AYANSWLDATT-GKFVVAVTDANRAG---E--
VRAAGA-V--AKV-VRH--SAATLSA-VQSTL-DARTAS-------------------------------------------------------------
-----------VP------DS-VTGWYVDAPTNQVVVSVLRGD-P----A---G--LA-WA--NA-SG----N-K-A-VRV---------
EA---------V-----T-DAPRPL-W-NIIGGQA-IY-----F----G----SGRCSVGFNARNASG-T--RY-VITAGHCTNL-
GG--T--VS-G---------T-G--GTIGPVAGSSFPTNDYGIIRVTS--S-AAV--STPL--VDRY-SG-
SDVTVAGSTVTPVGGAVCRSGSTTGWRCGTVQAFNQTV-NY-G--GG-------QIVGGLTRTSACAQPGDS-G-
GSFVS-A----VVSGRV--QAQGMTSGGS--------GNCTS---------------GGTTFFQPVREAMSAYGLTLY
>XX00000224 WP_055553223 33 136 189 MKHRRI-------------------------------------
--------------------------------------------------------------------------------------------------------------------------
----------------P-----K-R-RA--AI--AGTG---------------AVALVA-A--A-----L-T-----LQ------------------------------
------------NA---S----A-APAEPRPHTLSVNAAGKLANTLASALGG-DSAGSYYDAKA-GKLVVNVVNKAAAE---
R--VRDAGA-E--AKV-VEH--TLAQLGR-ASATL-KDK---------------------------------------------------------------
-------------AT------IP-GTSWSVDPRANKVVVTADSSV-T----GS--R-MAT-LGKVAK-SL----G-D-K-VEV--------
-KR---------S-----A-GTF--S-T-FLAGGDA-IW-----G----G----GGRCSLGFNVVKD-G-Q--PY-FLTAGHCTES-IK-
-S--WS-D-----SQ--D-G--DEIGANEGSDFPGNDYGLVKYTK--G-V-P--HPSA--VNLY-DG-
GTRAISKAGEATVGQKVTRSGSTTQVHDGEVTGLDATV-NY-Q--EG-------S-VSGLIRTNVCAEPGDS-G-
GALFA-G---------G--TALGLTSGGR--------GDCSS---------------GGETFFQPVPEALKAYGAEIG
>XX00000226 WP_043236868 33 136 189 MKHRRI-------------------------------------
--------------------------------------------------------------------------------------------------------------------------

----------------P-----K-R-RV--AI--AGAG---------------IAALVA-T--G-----I-T-----LQ--------------------------------
-----------SA---N----A-APGETQPDTLSVGAAGKLASTLTSSLKS-DTAGAYYDAKA-KKLVVNVVNKGVVD---
K--VRDAGA-E--VKV-VKH--TLAELSK-ARQTL-KEK-----------------------------------------------------------------
------------AT------IP-GTSWSVDPRSNKVVVTADSSV-K----GA--G-MAT-LDKVAG-AL----G-D-K-VEV--------
-KH---------S-----A-GKF--S-T-FIAGGDA-IW-----G----N----GGRCSLGFNVVKG-G-Q--PY-FLTAGHCTEA-IS-
-S--WS-D-----SQ--G-G--EEIGTNEGSDFPGNDYGLVKYTK--D-T-D--HPSA--VDLY-NG-
STQAISKAGDATVGQKVTRSGSTTQVHDGEVTGLNATV-NY-Q--EG-------S-VEGLIQTNVCAEPGDS-G-
GALFA-G---------D--TALGLTSGGS--------GDCSS---------------GGETFFQPVPEALQAFGAQIG
>XX00000254 WP_043469023 33 136 189 MKHRRI--------------------------------------
------------------------------------------------------------------------------------------------------------------------
----------------P-----S-H-RI--AI--AGAG---------------VTALVA-G--A-----F-T-----LN-------------------------------
------------SA---N----A-TTAEPAPTTLSAASADKLAGTLLSDLGD-EAAGSYYDAKE-RKLVVNVVDEGAAE---
D--VRERGA-V--ARV-VKH--TLAELKS-ARATL-KAE-----------------------------------------------------------------
------------AT------IP-GTSWSVDPKTNKVVVTADSTV-K----GA--K-LEK-LNKVVD-GL----D-G-R-AAV---------
ER---------V-----A-GEF--K-P-FAAGGDA-IH-----S----G----GGRCSLGFNVVKG-G-E--PY-FITAGHCGSA-GS-
-T--WS-A-----SS--G-G--SAIGTMEESSFPGNDYALVKYAS--G-V-D--HPSV--VNLY-NG-
STQQITQAGNATVGQQVQRSGSTTQVHGGEVTGLNETV-NY-Q--EG-------S-VSGLIKTTVCAEPGDS-G-
GSLFA-G---------S--TALGLTSGGS--------GNCSS---------------GGTTFFQPVPEALSAFGASIG
>XX00000022 WP_051344734 46 137 257 MRRRS---------------------------------------
------------------------------------------------------------------------------------------------------------------------
---TPREVSELSHRRKD-----KKR-AY--VI--AGAG---------------TAALAT-A--A-----I-M-----LP---------------------
----------------------Y---AN----AGQQEQPGPRTMSAKAAAAMGTKLADENGA-RTAGWFIDSKA-
KRLVMNVLDEETAQ---R--VQESGA-Q--ARI-VQN--SKAELKK-VSDTL-ANG--------------------------------------
----------------------------------------AS------VA-GSAWSVDPRTNKVNVLADSTV-D----DE--E-WAS-
LNKTAA-GM----N-G-M-MSV---------KR---------T-----A-GKF--Q----VLGGDA-IF-----A----E----
GTRCSLGFNVQVD-G-A--PG-FLTAGHCGNA-AQ--N--WT-S-----DEA-G-A--
QALGSITDSQFPETDFAIATLDD--A-QAE--ATSA--VNLG-QG-
GTQEITGAAEAAVGMQVQRSGSTTGVTDGEVTALDATV-DY-G--GG-------DVVNGMIQTNVCAEPGDS-
G-GAMFS-E---------D--QAVGLTSGGS--------GDCTQ---------------GGETFFQPVTDALEATGATIG
>XX00000029 WP_030547957 33 137 258 MSHRRV--------------------------------------
------------------------------------------------------------------------------------------------------------------------
-----------------N-----K-R-TS--AI--AAAS---------------AAALAG-A--V-----I------LLP-------------------------------
------------NA--NA----S-DEKEAAPRTFAGPAAAETGTALAADLGA-DAAGWYFDGDS-
GRLVMNVLDEEAAE---T--ARAKGA-E--ARI-VDN--SMAELKA-ATQTL-SDQ------------------------------------
-----------------------------------------AS------IP-GTAWSIDPKTNKVSVLADRTV-T----GD--K-LAE-LEQATD-
GM----A-G-L-VSV---------KR---------S-----Q-GEF--K-P-FADGGDA-IF-----G----G----GSRCSLGFNVDVN-
G-A--PG-FLTAGHCGGV-GQ--T--WT-A-----DQG-G-A--QPLGATAESVFPVADFSLVMYDD--P-ATE--
APST--VDLQ-DG-TTQEITEAAEASVGMEVQRSGSTTGLSDGQVTGLNATV-NY-G--NG-------
DIVNGLIQTNVCAEPGDS-G-GAMFS-G---------T--AAVGLTSGGS--------GDCTV---------------
GGETFFQPVTTALEAVGATIG
>XX00000093 WP_052808965 43 137 321 MSPRQ---------------------------------------
------------------------------------------------------------------------------------------------------------------------
------HSRRQAGHRQV-----NKR-TG--AI--AGAS---------------AAAIAA-A--A-----F-L-----LP---------------------
----------------------NAHA----S-TDTTPTLRTFSAPSAARLAATLKSDLGD-KAAGWYLDGST-

GHLVMNVLSKEEAE---R--VTAKGA-V--AKV-VRH--SMAALEQ-GTRTL-RAT-------------------------------------
-------------------------------------------AS------VP-GTAWSIDPRTNRIVVLADRTV-T----GR--K-MAA-LAKATG-
QM----G-E-M-VTV---------KR---------S-----A-GEF--R----PAGGDA-IF-----G----G----NARCSLGFNVTVQ-G-
A--PA-FLTAGHCGND-SK--T--WS-A-----DQ--G-G--QQLGTVKDSKFPKTDFALVTYDD--A-SAK--PQSA--
VTLG-NG-KTQPITKAAEAAVGMKVQRSGSTTGVHDGTVTGLDATV-NY-G--NG-------
DIVNGLIQTDVCAEPGDS-G-GSMFS-G---------D--SAVGLTSGGS--------GDCTK---------------
GGETFFQPVTKALEATGAKLG
>XX00000123 WP_033319317 32 137 304 MRHARR-------------------------------------
---------------------------------------------------------------------------------------------------------------------------------
------------------N-----V-Q-RF--AR--FAAI---------------G-GLVC-G--G-----L-----------------------------------------
----------MVSQ---AMASET---SAESDRPGAVSQAADTGRGLVSELGTERTAGTWIGA-D-
GRPMVAVTDEEAAA---D--VRRAGA-G--AKM-VDH--NMRELRS-ATATL-KDS----------------------------------
----------------------------------------------PR------VS-GTSWSMDYANNQVVVRADSTV-S----AD--E-WSR-
LTKVAE-SI----G-K-S-VRM---------ER---------I-----D-GTY--T-T-RLNGAEP-LF-----A----G----
AGRCSAGFNVTNG-R-A--DF-ILTAGHCGPE-GT--T--WF-S-----DQQ-G-T--
QQVGTTTDVEFPGTDYSLVRYDT--G-T-VLEGPDV--VAVG-NG-
QGVRITGAADPAVGQKVFRSGSTTGLQSGEVTGLDATV-NY-P--QG-------T-VTGLIETTVCAEPGDS-G-
GPMFA-D---------G--TALGITSGGN--------GDCDS---------------GGTTFFQPVTKALDTLGVQLA
>XX00000146 WP_031079649 32 137 267 MRHTRR--------------------------------------
---------------------------------------------------------------------------------------------------------------------------------
-----------------K-----L-R-RI--AR--LTAV---------------G-GMLC-G--G-----L-------------------------------------------
--------MVNS---AMAGES---PGPGGGGGHVARPPESGAHLVARLGPSRTAGTWIGA-D-
GRPVVAVTDTEAAA---E--VERAGA-R--AKV-VRH--SMRRLRA-ATETL-SEA--------------------------------------
------------------------------------------PR------VP-GTAWVMDYASNRVVVRADSTV-S----AA--D-WSR-
MSDLAE-GI----G-G-F-VHM---------QR---------T-----K-GAF--S-I-RLNGADP-IF-----A----G----
NGRCSAGFNVTDG-R-Q--NF-VLTAGHCGPR-GT--T--WF-T-----DAQ-G-S--
ARIGTTTASAFPGSDFSLVAYEN--G-D-GSGARSV--VGVG-PG-
REAGIGAAADPVVGQEVFRSGSTTGVRSGRVTALNATV-NY-R--EG-------T-VTGLIETTVCAERGDS-G-
GPLFA-D---------G--LALGLTSGGN--------GDCTT---------------GGTTYFQPVTTAMAALGVRLT
>XX00000152 WP_014140823 36 137 189 MRHRRI---------------------------------------
---------------------------------------------------------------------------------------------------------------------------------
----------------H-----T-P-RI--AL--AGAG---------------AAALAA-T--A-----V-L-----LPG-----------------------------
--------AGAATAASAHT----S------ATSHAQLSANAARATALVSRLGD-RAAGTYYDNAA-
GRLVVTITDPSAAG---S--VRSIGA-I--PRT-VKY--SAAQLHS-ATATL-AHQ------------------------------------------
--------------------------------------AR------IP-GTAWSIDPRQDKVVVTADPSV-T----GA--K-RAR-LDAALA-RL--
--G-D-K-ATL---------RH---------V-----T-SRL--R-P-LLAGGDA-IW-----G----S----DVRCSLGFNVTVN-G-S--
PY-FLTAGHCGNA-AS--S--WS-D-----SQ--G-G--GEIGQTVDSHFPGTDYALVQYTD--G-G-D--HPSA--
VDTY-DG-GSQSISQAADAYVGEQVQRSGSTTGVHGGQVTGLDATV-NY-E--EG-------S-
VSGLIDTTVCAEPGDS-G-GSLFD-G---------A--SAVGLTSGGS--------GDCSS---------------
GGETFFQPVTAALSAEGATLP
>XX00000255 KIX78759 33 137 190 MNHRRI------------------------------------------------
---------------------------------------------------------------------------------------------------------------------------------
-------P-----K-R-RV--AV--TGAG---------------IAALVV-A--G-----L-T-----FQ----------------------------------------
----TA--NA----S-EAPSAEPRTLSVASAGKLADTLAGDLGK-AAAGTYYDAKA-KNLVVNVLDEAAAT---E--

VEASGA-K--ARI-VEH--SLTALND-ARATL-KDR---------------------------------------------------------------------
-------AA------IA-GTSWAIDPTSNKVVVTADRTV-E----GK--E-LAK-LTKVVD-AL----G-S-K-AEL---------KH---
------T-----K-GEF--K-R-FIAGGDA-IH-----G----N----GGRCSLGFNVVKG-G-E--PY-FLTAGHCTEG-IS--S--
WS-E-----TS--G-G--AEIGTTVESNFPGDDYGLVQYTG--S-T-E--HPSE--VNLY-DG-
STQAITEAGDATVGQSVTRSGSTTQVSTGEVTALDATV-NY-G--GG-------DIVEGLIQTTVCAEPGDS-G-
GSLFA-G---------S--TALGLTSGGS--------GDCSS---------------GGETFFQPVPEALSAYGVEIG
>XX00000262 WP_055470807 33 137 189 MKHRRT-------------------------------------
-------------------------------------------------------------------------------------------------------------------
-----------------S-----TTR-RT--AV--AGAG---------------AAAVAL-T--A-----F-T-----LS-----------------------------
-------------ANA---A----P-A-PAPSPKALSATAAQQLAGALASGLGG-DAAGSYYDAEA-
RKLVVNVVDEAAAK---E--VRAKGA-E--AKV-VRH--SLAELDS-ARETL-ADR--------------------------------------
---------------------------------------AT------IP-GTSWSVDPRTNKVVVTADSTV-K----GE--R-LAE-LDKVVK-
SL----G-D-R-AAV---------KR---------T-----A-GEF--S-P-LIAGGDA-IW-----G----G----SGRCSLGFNVTVG-G-
Q--PH-FITAGHCTEA-IS--S--WS-D-----AQ--G-G--SEIGANADSSFPDNDYGLVKYTA--D-T-A--HPSE--
VNLY-NG-STQAIAKAGDATVGQQVQRSGSTTQVHDGEVTALDATV-NY-A--EG-------T-
VNGLIQTTVCAEPGDS-G-GSLFA-G---------D--TALGLTSGGS--------GDCSS---------------
GGETFFQPVPEALQAFGAEIG
>XX00000274 WP_012378904 29 137 189 MRRNSR-------------------------------------
-------------------------------------------------------------------------------------------------------------------
-----------------A----------R--LG--VS--------------------L-------------------LLVA--------------------------------------
GALGLGAAPST----AADTPPAAPSAIPAPSAYALDAAVERQLGA-ATAGTYLDAKT-GGLVVTVTTDRAEE---
Q--ARAAGA-T--VRR-VAR--SAAQLDA-AMATL-EAE-------------------------------------------------------------
---------------AK------IT-GTSWGVDPRTNRVAVEADSSV-S----AR--D-MAR-LEAVAE-RL----G-S-A-VDI------
---KR---------V-----P-GVF--H-R-EVLGGGA-IY-----G----G----GSRCSAAFNVTKG-G-A--RY-FVTAGHCTNI-
SA--N--WS-A-----SS--G-G--SVVGVREGTSFPTNDYGIVRYTD--G-S-S--PAGT--VDLY-NG-
STQDISSAANAVVGQAIKKSGSTTKVTSGTVTAVNVTV-NY-G--DG-------P-VYNMVRTTACSAGGDS-G-
GAHFA-G---------S--VALGIHSGSS--------GCSGT---------------AGSAIHQPVTEALSAYGVTVY
>XX00000286 WP_057235065 23 137 190 MRNRRP-------------------------------------
-------------------------------------------------------------------------------------------------------------------
---------------------------A--VL--LAAV---------------A-MLVT-G-SA-----L-A-----LP--------------------------------
-----------S----AAVARA---T------PVGEQAAAVATALQQSLGE-ESAGSYLDG-S-GRLVVTVTDQSAAQ---Q--
VRATGA-V--AQL-VSR--SGAQLAA-ATAEL-DRS-----------------------------------------------------------------
---------AA------VP-GTAWSVDPVSNQVLLSVDESV-N----GA--K-LTQ-VTEAAQ-RL----G-A-A-VRV---------
EQ---------V-----A-GRF--S-T-KLAGGDA-IY-----G----G----GYRCSLGFNVRSG-S-T--YS-FLTAGHCGNA-VS--
S--WY-T-----SQS-Q-T--TKLGNTTNSRFPGDDFAIVRYTN--G-S-I--PPSS--VDLY-NG-
TTRRITAASTPAVGATVYRSGSTTKVHSGQVTALNATV-NY-A--EG-------S-VRGLIRTTVCAEPGDS-G-
GALYS-G---------N--TAHGLTSGGS--------GNCSS---------------GGTTFFQPVVEALSAYGVSVY
>XX00000307 WP_033176777 23 137 184 MKSTR--------------------------------------
-------------------------------------------------------------------------------------------------------------------
--------------------------RI--QL--LAVT---------------T-GLLA-A-TA-----F-A-----LP------------------------------------
--------SASATQ----A---A------GAAKATPEAAASLAAHLGADRTAGTYYDSAT-KTTVVNVTSQATAD---A--
VRAAGA-T--PRL-VTH--SAAQLAK-AGDIT-K-A--------------------------------------------------------------------
--------AD------IA-GTAWSVDPRTDTLEVSADSRV-T----AA--Q-LAA-LTKSTA-AY----G-D-A-VHV---------TR-
--------V-----A-GTF--S-K-LLSGGDP-IL-----T----D----QWRCSVGFNVRSG-S-T--YY-FLTAGHCTQG-YP--A--

YY-T-----SS----G--SYIGPTVGTSFPGNDYGIVRYDT--S-V--A-HDGT--V-------
GSQDITSAANAFVGESVKRRGSTTGIHSGTVQALNATV-NY-G--AD-------GIVSGLIRTNVCAEPGDS-G-
GSLYD-G---------T--KAIGLTSGGS--------GNCSS--------------GGTTFFQPVTEALSAYGVSVY
>XX00000004 WP_011027321 33 138 283 MRHARR-----------------------------------
------------------------------------------------------------------------------------------------------------------------------
------------------R----IV-R-RV--AR--LAAV--------------G-GLLG-G--T-----M--------------------------------------
-----------VTRA---VASEPP---DASTAPRTFAQTPSGAGGDLVSRLGTGRTAGHWVDS-E-
GRPVVAVTDERAAE---A--VRRAGA-T--AKR-VRH--SMSELRS-AASTL-RSA------------------------------------
--------------------------------------PR------VA-GTAWALDYRSNEVVVRGDRTV-S----GS--D-WSR-
LTAVAE-GI----G-G-F-VRM---------ER---------T-----E-GTF--T-T-RLNGAEP-IL-----S----T----
AGRCSAGFNVTDG-T-S--DF-ILTAGHCGPT-GS--V--WF-G-----DRP-G-D--
GQVGRTVAGSFPGDDFSLVEYAN--G-K-AGDGADV--VAVG-DG-
KGVRITGAGEPAVGQRVFRSGSTSGLRDGRVTALDATV-NY-P--EG-------T-VTGLIETDVCAEPGDS-G-
GPMFS-E---------G--VALGVTSGGS--------GDCAK--------------GGTTFFQPLPEAMASLGVRLI
>XX00000010 WP_006606557 28 138 377 MNKRTG-----------------------------------
------------------------------------------------------------------------------------------------------------------------------
------------------------------AI--AGAA--------------AVAIAG-A--A-----I-L-----LP--------------------------------
---------NA---N----ASQDQPAAPKTLSAHSATQLAASLKADLGD-QSAGWYLDSGN-GHLVMNVLSKDDAA--
-S--VTAKGA-V--AKV-VQN--SLTALKA-GAKTL-RDN-------------------------------------------------------------
-------------AS------IP-GTAWSIDPKTNKIVVVADRTV-T----GA--K-MAT-LSKTTG-GM----GAG-M-VTV-----
----KR---------S-----Q-GEF--K----PVGGSA-IF-----G----G----NARCSLGFNVTVK-G-A--PA-FLTAGHCGND-
SK--T--WT-A-----DQG-G-S--QPLGTVADSKFPKTDFALVKYDD--A-AAK--PESS--VDLG-NG-
STQKITKAAEAAVGMKVQRSGSTTGVHDGTVTGLDATV-NY-G--NG-------DIVNGLIQTDVCAEPGDS-G-
GSMFA-D---------D--SAVGLTSGGS--------GDCTQ--------------GGETFFQPVTDALKATGAEIG
>XX00000034 WP_005484987 21 138 188 MT------------------------------------------
------------------------------------------------------------------------------------------------------------------------------
-------------------------------GAG--------------IAALVA-A--G-----V-T----FQT------------------------------------
------AN--AS----E-SASAPELKALTVMAAGKLARTLDADLGA-DAAGTYYDAKS-KLLVVNVLDESAAE---T--
VEAAGA-K--ARL-VEN--SLAELKS-ARTTL-TSD------------------------------------------------------------------
-------AT------IP-GTAWATDPTTNKVVVTADRTV-S----KA--E-LAR-LTEVVD-GL----G-A-K-AEL---------KR---
------T-----E-GEY--K-P-FVAGGDA-IT-----G----G----TGRCSLGFNVVKG-G-E--PF-FLTAGHCTEG-IS--T--
WS-A---------D-G--QVIGENADSSFPGDDYGLVKYSA--D-V-D--HPSE--VNLY-NG-
SAQAISGAAEATVGMKVTRSGSTTQVHSGTVTGLDATV-NY-G--NG-------DIVNGLVQTDVCAEPGDS-G-
GSLFA-G---------D--KAIGLTSGGS--------GDCTS--------------GGETFFQPVTEALSATGTQIG
>XX00000045 WP_031511141 31 138 190 MKHQRT-----------------------------------
------------------------------------------------------------------------------------------------------------------------------
-----------------F-----A-W-RA--AA--AGAG--------------AAAFLC-G--A-----LVP-----AS-------------------------
-----------------AD---P----A-PSAGAASAPLTAAAATELATSLEGDLKD-DMAGAYYDAGA-
GKLVVNVLGEQAAE---K--ARKAGA-E--PRI-VRY--SLARLGT-VQKDL-AA---------------------------------------
--------------------------------------TS------LP-GTARAVDHRLNKVVVSADSSV-E----GE--A-LSA-LRKQVA-
AQ----D-G-K-AVL---------KR---------V-----K-GEF--R-P-FVAGGDA-IW-----R---GD----NARCTLGLNVTRN-
G-L--PY-FLTAGHCGEL-LD--G--WS-T-----SR--G-G--RNIGITVASSFPGDDHSLVRYTA--R-V-D--HPSR--
VNLH-NG-TSQAITRAADPIVGQRVWRSGSTTGVRTGEVVATGVSV-TY-P--QG-------T-

VHDLIQTTVCAEPGDS-G-GPLFS-G---------E--TAHGLTSGGS--------GNCRT---------------
GGTTFHQPVTEALRAHGVRLG
>XX00000177 WP_053557080 37 138 276 MTHAR-------------------------------------
------------------------------------------------------------------------------------------------------------------------
-----------LRGVRRK-----L-R-R--AVR--LAAA---------------VG-LLS-G--G-----L--------M--------------------------
--------------VSGA---MAGEPP---AAGPPAAADPRAADDPAAGLVSRLGPARTAGSWTGP-D-
GKTVVAVTDQEAAE---E--VRRAGG-E--ARV-VRH--SMERLRS-ATETL-RSA-------------------------------------
---------------------------------------PR------VP-GTAWAVDYAANSITVRADSTV-S----AG--D-WSR-
MTGLAE-EI----G-G-F-VRM---------ER---------T-----T-GTF--T-T-RVNGGTA-VF-----G----D----
QRRCTAGFNVTNG-Q-D--IF-ILTAGHCGPQ-DS--T--WF-R-----DRQ-G-D--
ELVGTTTAASFPGDDYSLVRYED--V-D-VLDRTNV--VDIG-GG-
RGVQIVGAGEPAVGQRVFRSGSTTGFRSGEVTAVNATV-NY-P--EG-------T-VTGLIETTVCAEPGDS-G-
GPLFS-E---------G--LALGVTSGGD--------GDCTT---------------GGTTYFQPVTTAMEKLGVDLT
>XX00000182 WP_009314915 33 138 188 MKHRRI-------------------------------------
------------------------------------------------------------------------------------------------------------------------
----------------P-----G-R-RA--VV--TGAG---------------IAALVA-A--G-----V-T----FQT----------------------------
--------------AN--AS----E-VAPTPEPKALSVAAAGKLALNLDADLGA-DAAGTYYDAKS-KLLVVNVLDEAAAE--
-T--VEAAGA-K--ARV-VEN--SLAELKS-ARSTL-KED-------------------------------------------------------------
------------AT------IP-GTSWVTDPTTNKVVVTADRTV-S----KA--E-LTR-LTKVVD-DL----G-A-T-AEL---------
RR---------T-----K-GEY--K-P-FVAGGDA-IT-----G----G----SGRCSLGFNVVKD-G-R--PY-FLTAGHCTES-IS--
T--WS-A---------G-G--QVIGQNEDSSFPDNDYGLVKYTA--N-V-D--HPSE--VNLY-NG-
STQAITGAAEATVGMKVTRSGSTTQVHSGTVTGLDATV-NY-G--NG-------DIVNGLIQTDVCAEPGDS-G-
GSLFS-G---------D--KAIGLTSGGS--------GDCTS---------------GGETFFQPVTEALSVYGAEIG
>XX00000225 WP_037941100 21 138 189 MA------------------------------------------
------------------------------------------------------------------------------------------------------------------------
--------------------------------GAG---------------IAALAA-G--A-----V-T----FQT------------------------------------
------AN--AS----E-VPADAAPRTLSFQAAGKLASTLAQDLGA-DAAGTYYDAKT-KSLVVNVLDEAAAE---T--
VESAGA-E--ARV-VAH--SLAELKS-ARTTL-KQD-------------------------------------------------------------------
--------AT------IP-GTSWVTDPTTNKVVVTADRTV-S----EA--E-WAK-LTEVVD-GL----G-G-K-AEL---------
QR---------T-----K-GEY--K-P-FVAGGDA-IT-----G----N----GGRCSLGFNVTKG-G-E--PH-FLTAGHCTEG-IS--
T--WS-D-----S---S-G--NVIGENAASSFPGDDYGLVKYTA--D-V-A--HPSE--VNLY-DG-
SAQAISGAAEATVGMQVTRSGSTTQVHSGAVTGLDATV-NY-G--NG-------DIVNGLIQTDVCAEPGDS-G-
GSLFS-G---------D--KAVGLTSGGS--------GDCTS---------------GGTTFFQPVTEALSATGTQIG
>XX00000227 WP_037646379 21 138 189 MA------------------------------------------
------------------------------------------------------------------------------------------------------------------------
--------------------------------GAG---------------IAALVA-A--G-----V-T----FQT------------------------------------
------AN--AS----E-APKTEAPHTLSLSAAGKLASTLGKDLGT-DAAGTYYDAKA-KHLVVNVLDETAAK---T--
VEAAGA-K--ARV-VRN--SLAELTS-ARTTL-KQD-------------------------------------------------------------------
--------AT------IP-GTSWATDPETNKVVVTADRTV-S----KA--E-WAT-LTKVVD-GL----G-Q-R-AEL---------
QR---------T-----K-GEY--K-P-FIAGGDA-IT-----G----G----GGRCSLGFNVVKG-G-Q--PY-FITAGHCTES-IS--
T--WS-D-----S---S-G--SQIGTNEQSSFPGNDFGLVKYTS--N-A-D--HPSE--VDLY-NG-
STQPITKAGDATVGQKVTRSGSTTQVHSGTVTGLDATV-NY-G--NG-------DIVNGLIQTDVCAEPGDS-G-
GSLFA-A---------D--TAIGLTSGGS--------GDCTS---------------GGETFFQPVTEALSTFGAQIG

>XX00000249 WP_031075608 33 138 190 MKHRRI------------------------------------
------------------------------------------------------------------------------------------------------------------------
---------------P-----K-R-RA--AL--AGGA--------------AVALLG-A--G-----I-T----FQT----------------------------
------------AS--AS----E-EKPAPEVKALSATAAGNLASTLVERLGG-DAAGAYYDVKT-RALVVNVVDENAAD--
-A--VREAGG-R--ARI-VEH--SLAELTR-ARQTL-KTR----------------------------------------------------------
-----------AT------MP-GTSWAVDPVTNKVVVTADRTV-D----GA--S-LDK-LTQVVK-GL----G-A-K-AEL-------
--KR---------A-----A-GVF--R-P-FIAGGDA-IH-----S----G----GGRCSLGFNVVKD-G-Q--PH-FITAGHCGEA-
NS--Q--WS-E-----TQ--G-G--AAIGTMVDSQFPGNDFALVKYDG--D-T-A--HPSE--VDLY-NG-
STQQITKAGEATVGMKVTRSGSTTQVHDGEVTGLNATV-NY-G--SG-------QVVEGLIQTTVCAEPGDS-G-
GALFA-G---------D--TAIGLTSGGS--------GDCTS--------------GGVTFFQPVPEALEAFGAQIG
>XX00000250 WP_030124784 33 138 190 MKHRRI------------------------------------
------------------------------------------------------------------------------------------------------------------------
---------------S-----R-K-RV--TL--AGSA--------------VLALVA-A--G-----V-T----FQT----------------------------
------------AN--AS----D-DVPQFSAKTLTAHSAGSLAATLGKSLGG-DAAGSYYDAEA-KNLVVNVVDEAAAE-
--Q--VRQAGG-K--ARL-VQH--TLAELTT-ARQTL-TDK---------------------------------------------------------
--------------AT------IP-GTSWAVDPVSNKVVVTADSTV-H----GA--A-WDK-LGKVVE-GL----G-S-K-AEL-----
----KK---------T-----A-GEF--K-P-LIAGGDA-IW-----G----N----GGRCSLGFNVVKG-G-E--PY-FLTAGHCTES-
IT--S--WS-D-----SQ--G-G--SEIGANEGSSFPDNDYGLVKYTS--D-V-E--HPSE--VDLY-DG-
STQAITKAGDATVGMSVTRSGSTTQVHDGEVTALDATV-NY-G--NG-------DIVNGLIQTTVCAEPGDS-G-
GSLFS-G---------D--TAIGLTSGGS--------GDCSA--------------GGETFFQPVPEALEAFGAEIG
>XX00000251 WP_018893014 33 138 190 MKHRRI------------------------------------
------------------------------------------------------------------------------------------------------------------------
---------------P-----K-R-RA--AA--AGGA--------------VAALVA-A--G-----V-T----FQT--------------------------
--------------AN--AG----E-DKSAVAVKTLSASAAGNLASTLHESLGG-SAAGAYYDAEA-
KALVMNVVDEAAAD---A--VREAGG-R--ARI-VEN--SPAELRT-ARQTL-TDR----------------------------------
-----------------------------------AT------IP-GTSWATDPVTNKVVVTADRTV-E----GA--D-LAK-LREVVE-
GL----G-E-K-AEL---------KR---------T-----A-GEF--K-P-FAAGGDA-IH-----S----G----GGRCSLGFNVVKD-G-
A--PH-FVTAGHCGET-GS--E--WS-D-----SA--G-G--AAIGSMLDSRFPEDDFALVKYSD--G-T-A--QPSE--
VNLY-DG-TTRQITRAGDATVGMQVTRSGSTTQVHDGEVTGLDATV-NY-G--DG-------
QIVNGLIQTNVCAEPGDS-G-GSLFA-E---------D--TAIGLTSGGS--------GDCSS--------------
GGETFFQPVTEALQALGAEIG
>XX00000252 WP_030569936 33 138 190 MKHRRI------------------------------------
------------------------------------------------------------------------------------------------------------------------
---------------S-----R-K-RA--TV--AGSA--------------VAALVV-A--G-----F-T----FQS---------------------------
------------AN--AS----D-EVPQFTVKTLNAAAAGKLATTLDRDLGA-DAAGSYYDTKA-KALVVNVVDEAAAE-
--Q--VRQAGG-K--ARV-VEH--SLAELKS-ARQTL-TDK---------------------------------------------------------
--------------AT------IP-GTSWAVDPVSNKVLVTADSTV-D----GT--A-WKK-LSAVVE-GL----G-G-K-AEL------
---NK---------A-----A-GEF--K-P-LIAGGDA-IF-----G----S----GSRCSLGFNVVKD-G-E--PY-FLTAGHCTEG-
VT--S--WS-D-----TQ--G-G--AEIGANEGSSFPENDYGLVKYTS--D-T-A--HPSE--VNLY-NG-
STQAITQAGDATVGQQVTRSGSTTQVHDGEVTALDATV-NY-G--NG-------DIVNGLIQTTVCAEPGDS-G-
GALFA-G---------D--TALGLTSGGS--------GDCSS--------------GGTTFFQPVPEALAAYGAEIG
>XX00000253 WP_040875107 33 138 190 MKHRRI------------------------------------
------------------------------------------------------------------------------------------------------------------------
---------------P-----K-R-HA--AV--AGGA--------------VVALVA-A--G-----V-T----FQT--------------------------

---------------AN--AS----E-EPAGFAVKKLSATAAGNLASTLGDDLGD-RSAGAYYDVKS-
QALVVNVVDEAAAD---S--VRKAGG-Q--ARI-VEN--SVAELTT-ARQTL-KSR------------------------------------
----------------------------------------AT------LP-GTSWAVDPVTNKVVVTADRTV-K----GK--A-WST-LTKVVE-
GL----G-S-K-AVL---------KK---------S-----A-GEF--K-P-FIAGGEA-IH-----S----G----SGRCSLGFNVVKD-G-Q-
-PH-FITAGHCGES-GS--K--WA-E-----SQ--G-G--PEIGTMVDSQFPGNDFALVKYDG--D-T-E--HPST--VDLY-
NG-STQEITKAGEATVGMKVTRSGSTTQVHDGQVTGLDATV-NY-G--NG-------
QIVEGLVQTDVCAEPGDS-G-GSLFA-G---------D--TAIGLTSGGS--------GDCTS---------------
GGTTFFQPVPEALAVFGAEIG
>XX00000258 WP_046502510 33 138 189 MEHRRI-------------------------------------
----------------------------------------------------------------------------------------------------------------------
----------------S-----R-K-RA--TM--AGGA---------------AVALAV-A--A-----V-T----FQT----------------------------
---------------AN--AS----E-DTPTFTVKSLTAAAAGNLASALGEDLGG-DAAGSYYDATS-
RALVMNVVDERAAE---T--VRQAGG-R--ARL-VQN--SLAELKS-ARQTL-ADR------------------------------------
-----------------------------------------AA------VP-GTSWALDPVTNKVVVTADRTV-T----GE--A-WDR-
LSGVVE-GL----G-G-K-AEL---------KK---------T-----A-GEF--K-P-FVAGGDA-IT-----G----S----
GGRCSLGFNVVKD-G-A--PY-FLTAGHCTEA-IQ--T--WS-D-----A---Q-G--
NEIGTNEGSSFPGDDYGLVRYTA--D-V-A--HPSE--VNLY-NG-
SSQPITRAAEATVGQQVTRSGSTTQVHDGQVTGLDATV-NY-G--NG-------DIVDGLIQTSVCAEPGDS-G-
GSLFA-G---------D--TALGLTSGGS--------GDCSS---------------GGVTFFQPVTEALTVFGAQIG
>XX00000259 WP_018532397 33 138 189 MKHRRI-------------------------------------
----------------------------------------------------------------------------------------------------------------------
----------------P-----K-R-RV--AV--VGAG---------------ITALVA-A--G-----I-T----FQT----------------------------
-------------AN--AS----T-ETPDVALKNLSVAKAGNLASALHKDLGA-DTAGSYYDAKA-KHLVVNVLSEDVAQ--
-A--VEAAGA-R--ARV-VQN--SLAALKS-ARTTL-KSD------------------------------------------------------------
-------------AT------IP-GTSWATDPATNKVVVTADRTV-K----GA--D-WQK-LNQVVD-GL----G-A-K-AEL-----
----QR---------T-----Q-GEF--K-P-FVAGGDA-IS-----G----G----GGRCSLGFNVVKD-G-Q--PY-FITAGHCTEA-
IS--T--WS-D-----S---S-G--QEIGANEQSSFPDNDFGLVKYTA--D-V-P--HPSE--VNLY-NG-
SAQQITQAGDATVGMQVTRSGSTTQVHDGSVTGLDATV-NY-G--NG-------DIVNGLIQTDVCAEPGDS-G-
GSLFS-G---------D--TAIGLTSGGS--------GDCTS---------------GGETFFQPVTEALSTFGAEIG
>XX00000260 WP_030940931 33 138 189 MKHRRI-------------------------------------
----------------------------------------------------------------------------------------------------------------------
----------------P-----G-R-RV--AV--AGAG---------------IAALVA-A--G-----V-T----FQT----------------------------
--------------AN--AS----E-PAKSPASATLSALAAGKLASTLGQDLGA-DAAGTFYDAKT-RSLVVNVVDEAAAK--
-T--VEAAGA-K--ARL-VEN--SLAELKS-ARGTL-KKD------------------------------------------------------------
------------AT------IP-GTAWVTDPTTNKVVVTADRTI-S----DA--E-WAK-LTKVVG-GL----G-A-T-TEL---------
QR---------S-----K-GEF--K-P-LVAGGDA-IT-----G----S----GGRCSLGFNVVKG-G-E--PY-FITAGHCTEA-IS--
T--WS-D-----S---S-G--DAIGQNEQSSFPDNDYGLVKYTA--G-V-A--HPSE--VNLY-DG-
STQKITGAAAATVGMKVTRSGSTTQVHSGTVTGLDATV-NY-G--NG-------DIVNGLIQTDVCAEPGDS-G-
GSLFS-G---------S--NAIGLTSGGS--------GDCTS---------------GGETFFQPVTEALSALGAQIG
>XX00000261 WP_059196658 33 138 189 MKHRRI-------------------------------------
----------------------------------------------------------------------------------------------------------------------
----------------P-----G-R-RA--AV--AGAG---------------IAALVA-A--G-----V-T----FQT----------------------------
--------------AN--AS----E-PANDPTPKVLSALAAGSLASSLDKELGT-DAAGTYYDAKS-KALVVNVLDEAAAS---
V--VEEAGA-Q--ARV-VEN--SLAELRS-ARTTL-KDD------------------------------------------------------------

------------AT------IP-GTSWATDPTTNKVVVTADRTV-S----DA--E-WAK-LTKVVD-GL----G-A-T-AEL---------
QR---------T-----K-GEF--K-P-FIAGGDA-IT-----G----G----SGRCSLGFNVSKG-G-E--AY-FLTAGHCTEA-IS--
T--WS-D-----S---S-G--TQIGTNEVSSFPDNDYGLVKYTA--S-V-D--HPSA--VDLY-NG-
SSQAITGAAAATVGMKVTRSGSTTQVHDGSVTGLDATV-NY-G--NG-------DIVNGLIQTDVCAEPGDS-G-
GSLFS-G---------S--NAIGLTSGGS--------GDCTS---------------GGETFFQPVTEALSATGTSIG
>XX00000263 WP_030947398 33 138 188 MKHRRI--------------------------------------
------------------------------------------------------------------------------------------------------------------------
----------------P-----K-R-RA--AV--VGAG---------------IAALVA-A--G-----V-T----FQT-----------------------------
--------------AN--AS----E-PSKTSAPETLSAVAAGKLASTLGKNLGA-DAAGTYYDAKA-KSLVVNVLDQAAAK--
-T--VEAAGA-K--ARV-VEN--SLAELDA-ARTTL-TKD-----------------------------------------------------------------
-------------AT------IP-GTSWSTDPVSNKVVVTADKTV-S----KA--E-WAK-LSKVVD-GL----G-A-K-AEL--------
-KR---------S-----K-GEF--K-P-FIAGGDA-IS-----G----S----GGRCSLGFNVTKG-G-E--AY-FLTAGHCTDA-IK--
T--WS-D-----S---S-G--NQIGENADSQFPGNDFGLVKYTA--D-V-D--HPSE--VDLY-NG-
STQKITKAAEATVGEKVTRSGSTTQVHTGTVTGLNATV-NY-S--EG-------S-VSGLIQTDVCAEPGDS-G-
GSLFD-G---------E--SAIGLTSGGS--------GDCSA---------------GGETFFQPVTEALSAEGASIG
>XX00000265 WP_053324283 33 138 188 MNNRRI--------------------------------------
------------------------------------------------------------------------------------------------------------------------
-----------------S-----M-R-RK--AL--AGTG---------------IAVLIA-A--G-----M-S----FQT----------------------------
---------------AN--AS----E-SAQVPQAKVLSASEAGNFASTLLKDIDG-ATAGTYYDAES-KALVVNVLDKGTAQ-
--K--VEQAGA-K--ARI-VRH--TLAELDA-AQATL-AKD------------------------------------------------------------------
--------------AT------IP-GTSWVTDPVTNKVVVTVDSTV-T----GD--K-WAK-LSKVVD-TL----G-D-K-AEI-------
--KR---------T-----Q-GEF--R-S-FIAGGDA-IT-----G----S----GGRCSLGFNVVKN-G-Q--PY-FLTAGHCTDA-IS-
-T--WS-D-----S---S-G--RVIGRNAASSFPGNDYGLVEYTA--D-V-D--HPSA--VNLY-NG-
STQQITRAGNATVGEQVTRSGSTTQVHTGTVTGLNATV-NY-A--EG-------T-VSGLIQTTVCAEPGDS-G-
GALFD-G---------S--TALGLTSGGS--------GDCTS---------------GGTTFFQPVTEALSATGTQIG
>XX00000293 WP_019522793 21 138 189 MA--------------------------------------------
------------------------------------------------------------------------------------------------------------------------
---------------------------------GAG---------------IAALAA-G--A-----V-T----FQT--------------------------------------
------AN--AS----E-VPADAAPRALSVQAAGNLASTLLQGLGA-DAAGTYYDAKS-KSLVVNVLDETAAE---A--
VESAGA-K--ARV-VAN--SLAELKS-ARTTL-QQD-------------------------------------------------------------------
---------AT------IP-GTSWVTDPVSNKVVVTADRTV-S----KA--E-WAK-LTEVAE-GL----G-T-T-VEL---------
KR---------T-----Q-SEF--K-P-FIAGGDA-IT-----G----G----GGRCSLGFNVTKG-G-E--PH-FLTAGHCTES-IS--
T--WS-D-----S---S-G--NVIGTNAASSFPGDDYGLVKYTA--D-V-A--HPSQ--VNLY-NG-
SSQSISGAAEATVGMQVTRSGSTTQVHSGTVTGLDATV-NY-G--NG-------DIVNGLIQTDVCAEPGDS-G-
GSLFS-G---------D--KAVGLTSGGS--------GDCTS---------------GGTTFFQPVTEALSATGTQIG
>XX00000295 WP_059125791 21 138 188 MAG-----------------------------------------
------------------------------------------------------------------------------------------------------------------------
-----------------------------------GA---------------VAALIA-A--G-----V-T----FQN--------------------------------------
------AN--AS----E-APKTGTPRTLSAEAAGKLASTLGQDLGS-GAAGTYYDAKT-KQLVVNVLDESAAQ---S--
ARTAGA-T--ARI-VQN--SLAELAG-ARTTL-KKD-------------------------------------------------------------------
--------AT------IP-GTSWATDPRTNKVVVTADSTV-S----KA--E-WAK-LSKVVD-GL----G-G-K-AEI---------KR-
--------S-----P-GEF--K-P-FVAGGDA-IT-----G----S----GGRCSLGFNVVKG-G-Q--PY-FITAGHCTEA-IS--S--
WS-D-----S---S-G--NVIGQNEQSSFPGNDFGLVKYTA--E-V-D--HPSA--VDLY-NG-

STQAITKAGDATVGEKVTRSGSTTQVHSGTVTGLDATV-NY-A--EG-------T-VSGLIQTDVCAEPGDS-G-
GSLFD-G---------D--TAIGLTSGGS--------GDCTS--------------GGETFFQPVTEALSAFGAQIG
>XX00000297  WP_037738987   21   138   188   MTG-------------------------------------------
--------------------------------------------------------------------------------------------------------------------------------
-----------------------------------AG---------------IAALVI-A--G-----V-T----LQN---------------------------------------
-----AN--AS----E-TPTAAAPHTLSALAAGKLASTLGRNLGT-DAAGTFYDAKT-KNLVVNVLDQTAAK---T--
VREAGA-K--ARI-VEN--SLAELES-ARTAL-KQD------------------------------------------------------------------------
-------AT------IP-GTSWATDPVTDKVVVTADRTV-S----KT--E-WAK-LTKVVD-GL----G-T-K-AEL---------QR-
--------T-----K-GEF--K-T-FIAGGDA-IS-----G----S----GGRCSLGFNVVKG-G-Q--PY-FITAGHCTQA-IS--T--
WS-D-----S---D-G--NVIGQNEQSSFPGNDFGLVKYTA--D-V-D--HPSA--VDLY-DG-
SSQPITQAGDATVGETVTRSGSTTQVHSGTVTGLNATV-NY-S--EG-------T-VSGLIQTDVCAEPGDS-G-
GSLFD-G---------S--TAIGLTSGGS--------GDCTS--------------GGETFFQPVTEALSAFGAQIG
>XX00000298  WP_015624444   26   138   183   MRNPGS-------------------------------------
--------------------------------------------------------------------------------------------------------------------------------
-------------------------KLGR--PI--VGLA--------------A-VLLA-G-SA-----F-A-----SP---------------------------------
-----------A-------S-A---APAAPAADPSFAPSALASKLDTQFGT-KAAGSYIDS-A-GKLVVNVTDAATAA---S--
VKASGA-T--PRY-VTR--SGAELAA-ADSTL-KST---------------------------------------------------------------------
-------LK------TP-GTSFGVDPVTNQILVTADSTV-T----GT--K-LDA-VKAIVA-KL----G-D-K-ARL---------EQ---
-----T-----P-GTL--S-T-RISGGDA-IY-----T----S----SARCSLGFNVRSG-S-T--YY-FLTAGHCTNI-GT--T--WT-
N-----G-----S--TTLGTRSGTSFPGNDYGIVRYTT--S-S-VT-ISGA--V-----G--
SQDITSAGTPAVGATVYRRGSTTGIHSGQVTALNSTV-NY-A--EG-------S-VSGLIRTTVCAEPGDS-G-GSLYS-
G---------T--TALGLTSGGS--------GNCSS--------------GGTTYFQPVTEPLSVYGVSVF
>XX00000301  WP_043438960   21   138   187   MA--------------------------------------------
--------------------------------------------------------------------------------------------------------------------------------
--------------------------------GAG--------------VAALLV-A--G-----F-T----FQN--------------------------------------
------AN--AS----E-PAKAPEARTLSSAAAGKLASTLMRDLGA-DAAGTYYDATT-KNLVVNVLDPSAAE---T--
VERAGA-K--ARV-VRN--SLAELTG-ARTTL-EKD------------------------------------------------------------------------
--------AA------IP-GTSWATDPVTNKVVVTADSTV-S----GA--E-WNK-LSKVVD-GL----G-E-K-AEL---------
KR---------S-----E-GEF--K-P-FVAGGDA-IT-----G----S----GGRCSLGFNVVKG-G-Q--PY-FLTAGHCTEA-IS--
T--WS-A---------G-G--TVIGQNEQSSFPGNDFGLVRYTA--S-V-D--HPSV--VNLY-NG-
STQTITRAGDATVGERVTRSGSTSQVHSGTVTGLNATV-NY-A--EG-------T-VSGLIQTDVCAEPGDS-G-
GSLFD-G---------D--TAIGLTSGGS--------GDCTS--------------GGVTFFQPVTEALSATGTQIG
>XX00000060  WP_045303793   26   139   192   MSNRRT-------------------------------------
--------------------------------------------------------------------------------------------------------------------------------
----------------S-------R-RA--TL--LAVA--------------A-TLLA-G-TA-----L-A-----LP---------------------------------
---------TPA----VALPIP---N------PTGEAAVAVANDLQQQLGE-DTAGSYLDS-S-GRPVVTITHAADAQ---R--
VQSAGA-V--PRL-VTR--SAAQLAA-ATAEL-DRR----------------------------------------------------------------------
---------AA------VP-GTAWAVDPLTDQVVVSADSTV-R----GA--A-LDR-LTRTAD-GL----G-S-A-VRL---------
EQ---------V-----P-GVF--S-T-RVGGGDA-IY-----G----G----GYRCSLGFNVRSS-G-T--YY-FLTAGHCGNI-AS--
A--WY-A-----SES-M-T--DPTGSTTSSTFPDHDYALVQYTD--G-S-V--PASA--VDLY-DG-
TTRAINTAANPSVGQTVSRSGSTSHVHSGIVTGLNTTV-HY-A--QG-------T-VRGLIKTTVCAEEGDS-G-
GSLFA-G---------H--TALGLTSGGN--------GNCTL--------------GGTTFFQPVTAALAAYGVGLV
>XX00000079  WP_030564119   28   139   190   MSTQRA-------------------------------------
--------------------------------------------------------------------------------------------------------------------------------

-----------------S-----R-R-RL--TL--IGVG---------------VAALVA-G--P-----L-T-----AA-------------------------------
------------NT---S----A-ADPAPAPATLTSSSASKLARTLTAD-PA-RTAGAYYDTPG-KKLVVNVLDDDEAD---R-
-VRDAGA-E--PRH-VKH--SQAQLNA-VQRSL-GE-------------------------------------------------------------------
-----------RA------VP-GTARGVNPVLNKVVVTADSTV-G----GA--A-LNQ-LKRHVA-AQ----D-G-K-AVL-------
--KR---------A-----A-GTF--R-P-LIRGGDA-IY-----S----G----DSRCSLGFNVVKD-G-E--PY-FLTAGHCAAL-
AGR-I--WS-E-----TK--G-G--PVIGVVDDYRFPGSDDAIVEYTA--D-V-P--HPSE--VNLY-NG-
SAQEITGAREAVVGEQVQRSGSTTQLHDGSVQALNASV-TY-P--QG-------T-VHGLIQTDVCAEPGDS-G-
GAMFA-D---------R--DALGLTSGGT--------GDCTS---------------GGQTFFAPVTEALAHYGAEIG
>XX00000114 WP_050515473 35 139 229 MSKNHH-----------------------------------
----------------------------------------------------------------------------------------------------------------------------------
------------------V-----NKR-TG--VL--VGGA---------------AAALLA-A--A-----I-V-----LP----------------------------
---------------QA---N----ASPERPRPLRTFSSESAARVASSLKADLGADRTAGWYLDSGK-
GQLVMNVLSAEDAG---R--VEAEGA-V--ARV-VRN--SMGELQA-ATRSL-GNS----------------------------------
-------------------------------------------AA------IP-GTAWSIDPRTNRVSVTADRTV-T----GE--K-LAT-LTRAVE-
KL----G-S-GVASV---------RR---------S-----S-GEF--R-----VGGSA-IF-----G----G----NARCSLGFNVTVQ-G-A-
-PA-FLTAGHCGKA-SA--T--WS-A-----DQA-G-A--RQLGTVADAQFPESDFALVKYDD--P-GAR--PESA--
VDLQ-NG-GTQRISRAAEAAVGMRVQRSGSTTGVSDGTVTGLNATV-NY-G--NG-------
DIVNGLIQTDVCAEPGDS-G-GAMFA-A---------D--AAVGLTSGGS--------GDCTQ--------------
GGETFFQPVTKALAATGAQIG
>XX00000139 WP_030909936 28 139 189 MYYRQI--------------------------------------
----------------------------------------------------------------------------------------------------------------------------------
----------------S-----T-R-RA--RL--AGAC---------------ALALA--------------------TT----------------------------------
---AALAASNAHAA----A------PAAPGPQGLSAASAGSLAARFGD-TSQGSFYDSAH-RRLVVNVLDESAAD---
T--VRAGGA-T--PRL-VSN--SLAALRS-ATATL-KAK--------------------------------------------------------------
-----------AN------IV-GTAWVIDPRADKVVVTADRTV-D----DD--R-LAR-LNKIVG-DL----G-S-R-AVL---------
ER---------T-----P-GKL--S-L-SIAGGDA-IW-----G----D----TARCSLGFNVVKD-G-E--PY-FLTAGHCGKA-VK--
T--WS-S-----TQ--D-G--AKIAQTETVSFPTNDYSMAKYTD--T-A-A--HPSE--VDLY-NG-
TSQPITEAGTPLVGQSVERSGSTTKTHTGKVTGLDATV-NY-Q--EG-------T-VSGLIKTDVCAEPGDS-G-
GPLFD-S---------S--TALGLTSGGS--------GDCTS---------------GGETYFQPVPEALTALGASIG
>XX00000247 WP_039829465 33 139 190 MKHRRI--------------------------------------
----------------------------------------------------------------------------------------------------------------------------------
----------------P-----K-R-RA--AL--AGGA---------------VVALLG-A--A-----V-T----FQT-----------------------------
--------------AN--AS----D-EVPQFEAKTLSATAAGKLASSLDARLGDGDIAGAYYKVES-RTLVVNVVDEAAAD-
--T--VRRAGG-E--ARV-VEH--SLAELKS-ARETL-KDR-------------------------------------------------------------
-------------AT------IP-GTSWAVDPVTNKVVVTADRTV-K----GE--A-WEK-LSQVVK-SL----D-D-K-AEL--------
-KR---------A-----A-GTF--T-P-FASGGDA-IH-----T----S----GGRCSLGFNVVKD-G-Q--PH-FLTAGHCGGT-
GS--K--WS-D-----KQ--G-G--EPVGTMVDSQFPGNDYALVKYDR--E-V-D--HPSA--VNLY-NG-
SAQQIARAAEATVGMKVTRSGSTTQVHEGEVTGLDATV-NY-G--NG-------QIVEGLIQTTVCAEPGDS-G-
GSLFS-G---------D--AAVGLTSGGS--------GDCTS---------------GGVTFFQPVTEALTEYGAQIG
>XX00000257 KUM91808 33 139 188 MKHRRI-----------------------------------------------
----------------------------------------------------------------------------------------------------------------------------------
-------P-----G-R-RA--AL--AGAG---------------IAALVA-A--G-----VTF----QTA------------------------------------
------NA--SE----P-SKSAAAPQVLSAPAAGKLASTLVKDLGS-DAAGSYYDAQA-KSLVVNVLDAGAAK---T--
VQAAGA-K--ARI-VAH--SLAELDG-ARATL-KAD-------------------------------------------------------------------

---------AT------IP-GTSWATDPVTNKVVVTADKTV-S----GA--Q-WAK-LGKVVD-GL----G-T-K-AEL---------
KR---------S-----K-GEF--K-P-FIAGGDA-IS-----G----S----GGRCSLGFNVVKG-G-Q--PY-FLTAGHCTHA-IT--
S--WS-D-----S---S-G--NVIGKNEQSSFPGNDFGLVKYTS--D-T-D--HPSE--VDLY-NG-
STQKITGAAEATVGMKVTRSGSTTQVHDGTVTGLNATV-NY-S--EG-------T-VSGLIQTDVCAEPGDS-G-
GSLFS-G---------S--NAIGLTSGGS--------GDCTS---------------GGETFFQPVTEALSATGTQIG
>XX00000267 WP_027772022 30 139 189 MRPQMS-----------------------------------
------------------------------------------------------------------------------------------------------------
-----------------A------RL-GG--TL--AA-------------------L------------------LLLG-----------------------------------
--TAALGSTAAAA----PPPADRPDQGVKADAATVALSADLTRALGA-DSAGTYLDADT-GELVVTVTDKAAAA-
--E--ARAAGA-R--AEL-VEH--SASELRS-AMGAF-EDR---------------------------------------------------------------
---------------AK------IT-GTSWGVDPATNQIAVQADSTV-S----DA--E-YAK-LKRVAA-SL----D-G-A-ARV------
---ER---------I-----A-GTF--Q-K-EVLGGDA-IY-----G----G----GYRCSAAFNVTKN-N-A--RY-FVTAGHCTNA-
GA--S--WS-A-----TS--G-G--AQIGTREGTSFPTNDYGIVRYTD--G-S-S--PAGT--VNLY-NG-
STRDITSAANAVVGQAIQKSGSTTQVTSGTVTAVNVTV-NY-S--DG-------P-VYNMVRTTACSAGGDS-G-
GAHFS-G---------S--VAYGIHSGSA--------GCSGS---------------NGSAIHQPVTEALSAYGVSVY
>XX00000280 WP_018352447 25 139 190 MRISKL---------------------------------------
------------------------------------------------------------------------------------------------------------
-----------------------------------IA---------------AGALVA-------------------G------------------------------------
TLLVPAIAHAA----PTAAIPSAEAGIRAGLGAEATALLAGSLGD-RSAGSYLDTAS-GRMIVTVTDAEAAA---A--
VYAAGA-L--PHM-VTR--SGADLRA-ATSEL-DRS--------------------------------------------------------------------
---------AK------IP-GTTWAVDPATNQILVTADSTV-T----GE--K-LAT-MKRAVA-GL----N-G-A-ARL---------
VQ---------E-----P-GVL--N-S-YISGGNA-IY-----M----G----SSRCSLGFNVNRG-G-V--NH-FITAGHCTNL-GT-
-T--FW-S-----NSS-H-T--TVIGSRIGTSFPTNDYGIVRYSG--S-I-T--RPGN--VYLW-NG-
AYQDITSAGNAFVGQAIKRSGSTTGLRSGSVTALNATV-NY-P--QG-------T-VYQMIRTNACAEGGDS-G-
GSMFA-G---------S--VALGLTSGGS--------GNCSS---------------GGTTYFQPVPEVLSVYAASIY
>XX00000050 WP_010235465 30 140 184 MPLRRP-----------------------------------
------------------------------------------------------------------------------------------------------------
----------------------HRLLLAALAGTAAIGLGVA-----------------------------------VP-------------------------------
-----------A-------T-A---APVDLPILTEDAAAGLLPAVGEAAGT-SLAGSWFE--D-GRLMVGTWDPALAP---V--
LTRLGA-T--PVV-RDE--PRRDPGA-VLGEL-TAR------------------------------------------------------------------
--------MP------AG-LASYGIDPRTEQVVLEVVDGA-G----AD--A-LAGALT--AG-LD----P-S-A-VRV---------ER-
--------V-----P-AGPRQQ-A-AVAGGDT-IT-----D----G----SRRCTAGFAASDSSG-D--DW-VLTAGHCTRG-SS-
-T--WY-D-----SE----Q--STIGTGARSAGGAVDVGAIPVAE--G-S-A--SP-T--------V-
SGTRVAGTSAAPVGSTVCLYGSTSGRSCGSVERTDMTV-NF-D--GQ-------Q-QSGLTAVGACAQEGDS-G-
APYIT-S---------S-GQAQGVHTGAG--------G--SD---------------NCTSYFTPIGTATSALGLTLT
>XX00000062 WP_051073018 0 140 192 MV--------------------------------------------
-----------------------------------------------------------------------------------------------------
RDLGLDPATAEQLLEAQ--------------------------------------------------------------------------------------
---------------------------------------------EEALTVQESAASAAGQ-AYVDALFDTET-LSLTVLVSDPAAVA---A--
VEATGA-N--TRV-IDE--DPAATMA----AL-EQVP---------------------------------------------------------------
--------VP------EG-VTGWYPDTEQGLVVVEAVAP-------AA--A--DA-LLEEAG-TD----S-G-A-VRV---------VE-
--------V-----D-HAPMPH-S-ELIGGHP-FY-----V----G----SAQCVIGFSAVASGG-Q--GG-FITAGHCGRV-GA--
T--AL-D-----S---E-G--AALGVFQRSQFPGSDGAFVRVGP--E-W-T--LTPW--IELWSSG-

SYHSVTGSQEAPIGSVVCASAPINGWRCGVIEARGQTV-SY-P--SG-------T-VTGLTRTTLCADPGES-G-
APVVA-G---------S--QAQGVISGGS--------GNCAT--------------GGTTFFQRLNPMLTSWGLTLV
>XX00000098 WP_051468293 32 140 190 MRHTRG-----------------------------------
--------------------------------------------------------------------------------------------------------------------------------
-------------------------RGP--VV--LCAA---------------LGLTAAGLAVVPSL-ANA-----TP------------------------
------------------RS------E-D---VAQTASDSPMLAQSHLARGLSTKLGG-ATAGSYVDDAT-
GRLVVTVTNATAAE---S--VRAAGA-Q--PRT-VSR--SGGDLKK-VMAAL-KRD-----------------------------------
-----------------------------------------AT------VP-GTAWAADPKTDQVVLSVDRTV-T----GA--R-LAK-
VKKAAA-KH----G-A-A-VRL---------SH---------V-----G-GKF--R-P-LTAGGEA-IW-----T----D----
GGRCSLGFNVKKG-S-D--YY-FLTAGHCTAI-GK--S--WS-A-----TQ--G-G--
TPLGDTVDGTFPGHDYGLVKYAS--A-P-SD-TKGV--VSLY-GK-
GEQDITGAGEAVVGQTVTRSGSTTQVHDGKVTALDQTV-NY-Q--EG-------S-VSGLIQTTVCAEPGDS-G-
GSLFA-G---------D--KALGLTSGGS--------GDCTS--------------GGETFFQPVGPALEAFGAELY
>XX00000136 WP_028425804 37 140 190 MSQH---------------------------------------
--------------------------------------------------------------------------------------------------------------------------------
----------------TPARRPGRRRPLA-LA--AATA--------------ALALLP-A--------------------------------------------
-----------LSGTAQAAD----PAPGSDAGPSAAPTAAELRLADSLVDRLGA-RTTGSFLHPET-
GRLTVNVTDATSAE---Q--VRKAGA-E--PRK-VAR--GAAELAR-AADTL-ESE-------------------------------------
--------------------------------------VQ------IP-GTSWGTDPVTNRVEVQADETV-S----AH--E-LAR-LEEVVD-
GL----D-G-A-AEV---------TR---------V-----E-GAF--T-P-EVNGGDA-IY-----T----G----GSRCSAAFNVAKN-G-
V--RY-FLTAGHCTNI-AA--N--WS-A-----SS--G-G--HTIGVREGTSFPTNDYGIVRYTD--G-S-Q--APGH--
VNLY-NG-SFQDIARAANPVAGQAISKSGSTTGVTSGTVTATNVTI-NY-S--QG-------S-
VYGMVRTNLCSAGGDS-G-GAHFR-S---------D--VAYGIHSGGT--------GSCVN--------------
GGGAIFQPVTEALNVYGVSVY
>XX00000181 WP_016333982 23 140 201 MIRHPL------------------------------------
-------------RTLTALIGAAA-AVTAFAA-----------------------------P--------------------A---------------------------
---SAATSVLNEAAGP--------------------------------------------------------------------------------------------
-----------------------------------------AALAAAQQKLGTSLGT-AFAGTWLDTTT-GELVVGTTDARQSA---R--
IRSAGA-I--PKV-LKH--SAAELKA-IQSTL-DNRTEG---------------------------------------------------------------
--------LP------GS-VAGWYVDVPANEVVVSVVGGD-P----A---G--RA-W---AG-SA----G-V-P-VRV---------
EH---------V-----K-SAPRPL-W-SVVGGQG-LY-----F----S----GGACSVGFNAYDD-D-D--HY-VITAGHCTEL-
GG--T--VR-G---------V-G--GTIGKVARSSFPGNDYGTVEVTH--S-DVS--TPPR--VDRY-EG-
SDVRIEGADVVGVGGRVCRSGITTHWHCGRVEALDQTV-NY-G--NG-------NIVRGLTQTDACAEPGDS-G-
GSFVS-RPSSGSGTKLV--QAQGMTSGGS--------GDCGE--------------GGTTFFQPVKEVLDRYDLKLE
>XX00000269 WP_051172873 31 140 187 MCMIR-------------------------------------
--------------------------------------------------------------------------------------------------------------------------------
----------------------HGGRGGIAF--AAAV--------------L-AAGS---------------------------------------------------
---------------VFAASGVATAQPSAAPLNARAATAISANLAKSLGD-RTVGSFYDQAS-SRLTVTVTDQAAAD---
Q--VRAAGA-T--PKF-VKY--RASQLKA-VTAEL-ART---------------------------------------------------------------
------------AS------VP-GTAWRVDPRTGQVLVTADSTV-T----GA--K-LSQ-VTNAVH-RF----G-D-K-ARF-------
--AQ---------T-----K-GKL--R-P-LISGGDA-IW-----G----Q----GLRCSLGFNVHTSSG-Q--AA-ILTAGHCGVA-
TN--D--WW-A-----DQG-N-S--QHIATTSAANFPGTDYSYATYDA--G-V-D--APSA--VN------
TGQQITAAGDATVGETVTRSGSTTGVHDGQVTALDATV-NY-Q--EG-------S-VSGLIDTTVCAEPGDS-G-
GSLFD-G---------A--TALGLTSGGS--------GDCNS--------------GGETFFQPVPAALSAYGLVLP

>XX00000264 WP_005309034 29 141 189 MRRTLR----------------------------------
----------------------------------------------------------------------------------------------------------------------
----------------A-----RIC-LP--AL--LA-------------------L------------------TAVG----------------------------------
-LGSSTAGAADA----ADTGGNPSPGVAASPAQYSLHTSLERDLGD-RTAGSYFDTAS-GELVVTVIDDEAAE---
E--VRATGA-R--AQL-VTR--SMAQLRA-AMDTL-ESK------------------------------------------------------------
-------------AK------IT-GTSWGIDPSTNKIAVEADSTV-S----PA--S-MAK-LRAAAA-RL----G-D-A-VSI---------
KR---------V-----A-GEF--K-K-EVAGGDA-IY-----G----G----GYRCSAAFNVSKA-T-T--RY-FLTAGHCTNS-AS--
S--WS-A-----TS--G-G--PAIGTREGTSFPTNDYGIVRYTD--G-S-Q--PAGN--VNLY-NG-
SHQDITSAADAIVGQSIRKSGSTTKLTSGTVTAVNVTV-NY-G--DG-------P-VYNLVRTTACSAGGDS-G-
GAHFS-G---------S--TALGIHSGSA--------GCTGT--------------NGSALHQPVKEALSAYGVAVY
>XX00000266 WP_018848369 29 141 189 MRRTLR----------------------------------
----------------------------------------------------------------------------------------------------------------------
----------------A-----RIC-LP--AL--LA-------------------L------------------TALG-----------------------------------
LGAHTSTAAPG----PDGAPSPGPGATATAAQYSLNSSLTQRLGD-RSAGSYLDSAT-GELVVTVTDAEAAT---
A--AREAGA-R--AEL-VRH--STAELRS-AMDTL-RAR-------------------------------------------------------------
------------AA------IT-GTSWGIDPSSNAVAVEADSSV-S----AR--D-LAR-LKKVAA-SL----G-D-R-VSI---------
KR---------V-----P-GVF--T-K-EVAGGDA-IY-----G----G----GSRCSAAFNVSKG-T-A--RY-FVTAGHCTNL-SA-
-S--WS-A-----TS--G-G--AAVGTREGTSFPTNDYGLVRYTN--G-S-Q--PAGN--VNLY-NG-
GYQDITSAADAVVGQSISKSGSTTRVTSGTVTAVNVTV-NY-S--DG-------P-VYNMVRTTACSAGGDS-G-
GAHYS-G---------T--TALGIHSGSS--------GCSGT--------------AGSAIHQPVKEALSAYGVAVY
>XX00000100 WP_030544818 32 142 190 MKHRI-------------------------------------
----------------------------------------------------------------------------------------------------------------------
----------PRFARR-----PST-RA--VL--AGAG--------------AAVLAL-T--A-----P-A-----LP-------------------------
--------------RAG---A----A-V-PRPAPAVISPAAAGALSAQLAGRLGN-EAAGAYYEAGS-
GRLVVNVLTEGAAD---T--VRAAGA-E--PKL-VRH--SAAELDA-ARRVL-RER-------------------------------------
------------------------------------AT------IP-GTAWSADPRANKVVVTADRTV-T----GV--R-LDR-LRDVVD-
RL----G-A-M-ATL---------ER---------T-----D-HEL--R-P-LIMGGDP-IW-----T----G----RERCSLGFNVTVN-G-
S--PH-FLTAGHCGGR-GT--S--WS-A-----SR--G-G--PRIGVVTHARFPGEDYALVRITS--S-LVE--TPSR--
VGLF-WG-QTQRITRAAAPVVGQRARHKGSTSGLRTGRVTGVGITV-NY-R--QG-------S-
VHGLIETNMCAEPGDS-G-GPLFA-G---------T--TALGLTSGGI--------GDCTR--------------
GGMTYYQPVAGAVRATGVTIG
>XX00000102 ELP66831 19 143 307 MGSV--------------------------------------------
----------------------------------------------------------------------------------------------------------------------
----------------------AALG--------------AAAL-----------------------------------------------------------
LLPNAMASQTDAAAGAAPKTLKATDASDLASQLQQLLGD-AFAGSYYDTDK-QQLVVNVVEGDNNI---Q--
AKKAGA-A--VRE-VEN--TIGELKA-GAQTL-KAK-------------------------------------------------------------
--------AT------IP-GTAWALDPRTNKIQVTADSTV-T----GA--G-WDT-IESTVK-SL----G-SGM-ATI---------KK-
--------T-----A-GTF--K-P-LVEGGDA-IF-----G----G----GARCSLGFNVVTGDG-T--PG-FLTAGHCGVA-AA--
E--WS-D-----AE--N-G--EPIATVEAATFPGDDFALVKYND--P-A-TV-AAST--VDVG-NG-
QTVDIAQAAEAAVGQQVFRMGSTTGLNDGEVTGLDATV-NY-A--EG-------T-VTGLIQTNVCAEAGDS-G-
GSLFT-Q--------DG--SAIGLTSGGS--------GDCTV--------------GGETFFQPVTTALAAVGATIG
>XX00000130 WP_028798215 32 143 311 MRHARR---------------------------------
----------------------------------------------------------------------------------------------------------------------
-----------------S-----L-R-R--LMR--FAAAGGVVCGSLM----------V-T--H-----A--------V-----------------------

---------------ANEP---PGAAPG---QVTYAAGPVGGGPAPAVTEVVRRLGDARTAGSWAGA-D-
GRPVVAVTDAGAAA---E--ARRAGA-T--VKR-VRH--SMDTLRA-ATESL-GAE----------------------------------
----------------------------------------PR------VP-GTAWAVDYAANQVVVRADTTV-S----AD--D-WSR-
LSGVAE-RT----G-G-V-VRM---------ER---------T-----T-GAF--T-T-RVNAAAP-IF-----A----S----
GGRCSAGFNVTDG-R-D--VY-VLTAGHCGPA-GT--A--WF-Q-----NSD-G-S--
RLLGTTVTADFPGSDFALVRYER--G-A-PLGGADV--VAIG-GG-
RGVRITGAADPVVGQRVFRSGSTTGLRDGRVTALNATV-NY-P--EG-------T-VTGLIETTVCAERGDS-G-
GPLFA-Q---------G--LALGLTSGGN--------GDCAR--------------GGVTYFQPAVKTMRALGVRPV
>XX00000141  WP_052850045         30       143      190      MTHRRT-----------------------------------
------------------------------------------------------------------------------------------------------------------------
----------------S-----HRR-RV--AA--AATG--------------IAALFA-GTFA-----VAE-----AN-------------------------
----------------AE---S----A-AVPTAAPQVLSSAEAGTLAASLSAQLTPETSGGTYYDADS-
GTLVVNVTDETAAA---T--VRSAGA-E--ARV-VRH--SLAELDE-VKDAV-AG-------------------------------------
--------------------------------------FA------VA-GTAWAVDPVSNTVRVTTDSTV-R----GA--S-LAD-LRASVA-
EL----G-A-R-ASL---------DA---------V-----A-GEF--Q-P-FIAGGEA-IY-----S----S----GARCSLGFNVSIG-G-S--
AG-FLTAGHCGSV-GS--S--WS-A-----SS--G-G--PALGTLTAGQFPGADYALVEYSA--PAP-D--HPSA--VFLY-
DG-TTQEITGAAEATVGQSVQRSGSTTGLHDGEVTAVDVAV-RY-P--QG-------T-VEGTIQTTVCAEPGDS-
G-GALFD-G---------S--NAIGLTSGGS--------GNCTS--------------GGTTFFYPVTDALAAVGATIP
>XX00000194  WP_044580095         30       144      264      MKHMRR----------------------------------
------------------------------------------------------------------------------------------------------------------------
-----------------G-----V-R-RV--VR--FAAVGALVCGGVMVSQA--GPG----G--S-----A--------------------------
-----------------------------HGTT---GSGGPRALSADRTDDVGGRLVSALGASRTAGNWIGA-D-
GRPVVAVTDDTAAD---E--VGRAGA-T--AKR-VRY--SMADLDS-ATERL-RSA----------------------------------
----------------------------------------PR------VA-GTAWMVDPKSNQVVLAGDSTV-S----TA--R-WSK-
MKDLAA-EV----G-G-A-VRA---------QR---------T-----E-GSF--T-T-RTAGASP-MF-----T----N----
GSRCSAGFNVTNG-Q-T--NF-ILTAGHCGPN-GT--P--WF-T-----DGT-G-S--
TRIGTTVQTSFPGNDFSLVRYDN--A-A-LD-QNSV--VNVG-GG-
RTVRVTGVADPVVGQEVFRSGSTTGLRSGKVTGLNATV-NY-P--EG-------T-VTGLVQTTVCAEPGDS-G-
GPLFA-Q---------G--VALGVTSGGS--------GDCDR--------------GGVTFFQPVTKALTALGVSIP
>XX00000026  WP_014173666         30       146      295      MGHTRR----------------------------------
------------------------------------------------------------------------------------------------------------------------
-----------------R-----A-H-RV--AR--LVAIGAVVCGGVMVSQAAADGS----P--A-----N-------------------------
------------------------------GSGG---DPTRALEARDTRDAQLARTLVDKLGTSRTAGSWIGS-D-
GRAVVAVTDAEAAD---Q--VNQAGA-R--AKQ-VRY--SMRQLRS-ATETL-REA---------------------------------
----------------------------------------PR------VA-GTAWAVDAMSNEVVVLGDSSV-S----AD--Q-WSR-
MRGVAS-AI----G-G-E-VRM---------ER---------T-----A-GSF--T-T-RTLGAEP-IF-----T----Q----
GGRCSAGFNVTNG-Q-A--GF-ILTAGHCGPN-GT--A--WF-A-----DTQ-G-N--
TTVGTTVRSDFPGSDFSLVQYQN--T-S-LD-QSSV--LNIG-NG-
QTVRITGAADPVVGQKVFRSGSTSGLRTGSVTALNATV-NY-P--EG-------T-VSGLIQTDVCAEPGDS-G-
GPLFA-Q---------N--VALGVTSGGT--------GDCTS--------------GGVTFFQPVTKALTALGVSLP
>XX00000061  WP_030751531         32       146      285      MKHARR-----------------------------------
------------------------------------------------------------------------------------------------------------------------
----------------T-----L-R-RI--VR--LAAV--------------G-GLLC-G--GLLVSNAVA-----EE-------------------------
-------------PGGA---RAGGNG---ASGTPGTSGASGAAATGAGLVERLGTSRTAGTWIGS-D-

GRPVVAVTDEAAAD---E--VRRAGA-R--PQM-TRY--NMRELRA-ATETL-SRS-----------------------------------
----------------------------------------SR------VA-GTAWAVDYASNKVVVRADSTV-S----AA--D-WSR-
MSAVAR-DI----G-G-S-VEM---------RR---------T-----E-GTF--T-T-RVNGAAP-IL-----T----N----
GGRCSAGFNVTNG-A-E--QF-ILTAGHCGPV-GT--G--WF-A-----EDL-A-A--
GQIGTTVAGRFPGNDFSLVRYEE--G-A-APGDANI--VDIG-NG-
QAVRVTGAADPVVGQQVFRSGSTTGLRSGQVTALNATV-NY-R--EG-------S-VSGLIETTVCAEPGDS-G-
GPLLA-E---------G--IALGITSGGS--------GDCTS--------------GGTTFFEPVTSAMNALGVRLT
>XX00000066 WP_009720379 65 146 202 MNTT--------------------------------------
--------------------------------------------------------------------------------------------------------------
-------GGPCVSTAVA----NPSPAYDPVRP--TG----------------AAAVVG-A--A-----I-V-----LP--------------------
-----------------------NANASQDKKSEAIDAKPKAFNSDSASDAIAQLASSLGE-AFGGAWYDKDQ-
QQVIVNVIGDVEVA---A--VKSAGA-V--PKA-AKN--SMKSLKA-AESTL-KKK-------------------------------------
--------------------------------------AT------MA-GTAWAINPKTNKLQVTADSTV-K----GA--K-WDT-LESTVK-
SL----G-ADM-ASV---------KR---------S-----A-GTF--K-T-FAEGGDA-IF-----G----G----GSRCSLGFNVTTESG-
D--PG-FLTAGHCGVA-AK--E--WS-E-----TQ--D-G--APVATVQDATFPGTDFALATYDD--A-A-TE-APST--
VNTG-DG-NKVNITEAAEASVGQEVQRMGSTTGLNGGSVTGLNATV-NY-A--EG-------S-
VSGLIQTDVCAEPGDS-G-GALFTEG---------G--QAIGLTSGGS--------GDCTA--------------
GGETFFQPVTTALEAVGATIG
>XX00000078 WP_045316694 22 147 188 MLLRI--------------------------------------
---VSVS----------A---------------------------------I-VLSTITIP----------------------------------
AAAASAEVVDAMSRDFGISTAQALTRIRQE-----------------------------------------------------------------------
--------------------------------------------------------------------KVALKIAPAAQEAAAD-AYGGSWFAD---
GGLQVAVTDPAKLD---A--VKAAGA-Q--PVL-RTK--SFASLTA-TKNGI-DRLSKAGA----------------------------
-------------------------------------------VS------KD-VAGWRVDVQRGAVVVDALPG--------A--D-LSG-
LPKDVV---------------V---------NT---------A-----GVQKPQTFAA-GTVGGDP-FY-----T----G----
NVRCSIGFSVQG--------G-YVTAGHCNVG-AW--N--VY-G-----W---D-R--
SWQGEFVGSSFPTNDYGWVRTGG--G-W-W--NVAV--VLGW-
GQVSDALVRGSWEAPPGTSVCRSGSTSHWHCGVVQGRNETV-NY-Q--QG-------A-
VYEMVGTSVCAEPGDS-G-GSFIT-G---------D--QAQGVTSGGY--------GNCSS--------------
GGRTWFQPVNEILRVYGLRLL
>XX00000238 WP_055501245 31 147 190 MSIRGL------------------------------------
--------------------------------------------------------------------------------------------------------------
----------------G-----T-R-RA--GL--IAA---------------A-GLLA-S--A-----LLG-----SP--------------------------------
----------A-------Q-A-----SPPPSQPGADPAAAAAAAESRLGD-RTAGSYFDQAQ-GKMVVTITDPADAQ---A--
VTDAGA-V--PKV-VPR--SGEHLAG-LMAQL-DRT---------------------------------------------------------------
----------AR------VP-GTAWAVDPAANQVVVWVDSTV-G----GG--A-RGK-VEAAAR-EA----S-P-A-VRV------
---EN---------E-----P-GRF--Q-L-FIQGGDA-IY-----G----G----GYRCSLGFNVVSG-S-T--YY-FLTAGHCGNI-
AS--T--WY-A-----NSS-R-T--TVLGTVSGSSFPGNDYALVRYTN--T-S-IA-KPGS--VNLY-SG--
SRDITGAGNAYVGQSVQRSGSTTGLHSGTVTALNATV-NY-A--EG-------T-VREMIKTTVCAEGGDS-G-
GPLFS-G---------G--TALGLTSGGS--------GNCSS--------------GGTTFFQPVTEALSAYGVSIY
>XX00000126 WP_051102134 32 148 314 MRRKRR-----------------------------------
--------------------------------------------------------------------------------------------------------------
-----------------Q-----V-Q-R--ALR--LVAV--------------A-GVVC-----------G----GVMM------------------------
-----------SQAFGAAPDGTQPRSGSRVLDAPDGARGPADAAGLAEDLAARLGDARTGGSWVDD-E-

GRPVVAVTDEDTAA---E--VARAGA-R--AEV-VRY--SMRDLQD-AAGAL-REA-----------------------------------
-------------------------------------------ER------VP-GTAWSVDPVSNEVLVLADSTV-S----AQ--E-WAR-
LQSVAD-GI----G-G-Q-VRM---------ER---------T-----E-AAF--T-T-RTAGAAP-IL-----T----R----
GSRCSAGFNVTNG-T-T--DF-ILTAGHCGPP-GT--G--WF-A-----DNQ-G-T--
RPVGTTVSTSFPGGDFSLVQYQN--D-G-SD-HSSV--VGVG-SG-
RVVRITGASDPFVGQEVFRSGSTSGLHSGQVTGLNATV-NY-P--EG-------A-VSGLIQTTVCAEPGDS-G-
GPLFA-Q---------G--LALGITSGGS--------GDCKA---------------GGITFFQPVTTAMATLGVSLP
>XX00000069 WP_030369543 21 149 192 MKRRWI------------------------------------
-----------------------------------------------------------------------------------------------------------------------------
-----------------P-------V-RV--AL--AAAS---------------AVVLAA-V---------------------------------------------------
------TLSPQQADA----SPGSPAAPARAPSAASDASLARTLAAALGG-AAAGSYYDSTL-GRLVVDVTDGRAAG-
--L--VRAAGA-A--PRV-VEH--SKAELDS-AHATL-DSR---------------------------------------------------------------
-------------IT------TP-GTAWAADPRLNKVVLTADPTV-T----AA--D-LAR-IKDTAR-SL----G-S-A-VVV---------
KR---------S-----P-TKL--T-R-FIAGGDG-IW-----G----S----QYRCSLGFNVKRA-GKP--DA-FVTAGHCGKV-EP-
-E--WR-A-----SN--G-G--AVIAKTEAWKFPGRDYAIAAYPA--TGAVD--HPSA--V----NP-
GLQAITGAGAATVGQAISRSGSSTGLKTGKVTALNATV-NY-K--EG-------Q-VTGLIETSACAEGGDS-G-
GPLFA-G---------T--TALGVLSGGS--------GSCTMP------------FPAPTTYFQPVQEILDAFGATIP
>XX00000158 KGQ05981 24 150 188 MELAKF----------------------------------------------
---------------------------------------------------------------------
LALLAVVLPVTYGAPTQAAATGELHPKILEAMKRDLGLDADQAHARVARE-------------------------------------
--------------------------------------------------------------------------------------------------------
AAAANVIEQMRGSAGE-SFAGAWFDA---DTLHIGVTDEALAG---E--VTAAGA-T--PIV-MTN--SLSKLEK-
AKADL-DKV---------------------------------------------------------------------------------F-----IP------
QANTLSGGLVIEAIGDN-H----GR--A--EE-LAAQIG-LA----S-G-E-YEV---------RT---------V-----E-ELPTLY-A-
SVQGGDN-YFI----D----N----ASRCSVGFAVTT--------G-FVSAGHCGQT-GS--S--AT-T-----P---R-G--
QPLGTFAGSVFPGSDMSYVRTVS--G-T-G--LNGS--INGY-GQ-
GNLPVSGSNVAAVGASVCRSGATTQVHCGTIRARGATI-NY-Q--EG-------S-VTGLTQTTVCAEQGDS-G-
GSFYA-G---------S--QAQGVTSGGN--------GNCRT---------------GGITYFQPVNEILETFGLTLV
>XX00000404 WP_011290931 31 151 186 MNHSSR------RTT--SL-------------LF----
--------TAALA-----------------------------------------------------------------------------------
ATALVAATTPASAQELALKRDLGLSDAEVAELRAAK----------------------------------------------------------------
-----------------------------------------------------------------------------AEAVELEEELRDSLGS-
DFGGVYLDADT-TEITVAVTDPAAVS---R--VDADDV-T--VDV-VDF--GETALND-FVASL-NAIADT------------
--------------------------------------------------------------AD------PK-VTGWYTDLESDAVVITTLRGG-T----PA--
A--EE-LAERAG-LD----E-R-A-VRI---------VE---------E-----D-EEPQSL-A-AIIGGNP-YY-----F----G----
NYRCSIGFSVRQG-S-Q--TG-FATAGHCGST-GT--R--VS----------------SPSGTVAGSYFPGRDMGWVRITS--
A-D-T--VTPL--VNRY-NG-GTVTVTGSQEAATGSSVCRSGATTGWRCGTIQSKNQTV-RY-A--EG-------T-
VTGLTRTTACAEGGDS-G-GPWLT-G---------S--QAQGVTSGGT--------GDCRS---------------
GGITFFQPINPLLSYFGLQLV
>XX00000219 WP_052534174 34 152 191 MTSR-------------------------------PSRLP-
RGSTVTPRRLL--AGLAGTTLIAVALA-VPASTA----------GAAKQSTAD----QSQN----L---------------------------
-----------------------------IAVSAAQKAS--------------------------------------------------------------------------------
--------------------------------------------------------------VQQDKAADALVKKLGK-ASAGVYYD--S-
NRLSAAVTTKAALK---Q--AKAAGA-T--AHL-VKN--STAQLTS-AVSLL-NKT--------------------------------------

-------------------------------------AK------VP-GSSWAIDPASNKVQVSIDSTV-K----GA--R-LAK-VQAAVA-
KL----G-S-K-ASL---------ER---------V-----S-GTF--T-P-RIAGGQA-IY-----G----G----QYRCSLGFNVKTS-S-A--
YY-FLTAGHCGQV-AS--T--WY-S-----NSA-K-T--ALLGSNVSYSFPGNDFALVKYSS--T-S-TV-PSGT--VYTY-
SG--TQEISSAGTPTVGQTVYRSGSTSGVHSGKVTALNATV-NY-S--DG-------TSVSQTIKTSVCAESGDS-G-
GSLYA-G---------T--VAYGLTSGGS--------GNCTS---------------GGVTYFQPVPEALSYYGVSIY
>XX00000067 WP_045315880 26 153 187 MKRMA-------------V--------------AA-----
-------VAVLA----------------------GA-------------------L-A--------T-----------------TVSPA--------AVA---
APEPSADLLAAMQRDLGLNAQQAADRLRQE--------------------------------------------------------------------
-------------------------------------------------------------------SAAAVLQHVAKGFAGD-EFGGAWFDASI-
GKLVVGVTDGAKAD---R--LRRLGA-E--PRT-VAH--KASDLDS-AKSKL-DAE-------------------------------------
--------------------------------------AP------AE-VTGWYVDTRANAVTITVKRGQ-A----DA--A-K-S-LVEKVG-
T---------L-AKV---------VE---------T-----D-EAPRVL-K-DIRGGDA-YYI----N----N----AGRCSVGFAVQG-------
-G-FVSAGHCGSP-GD--T--VA-G-----V---D-K--TALGTFEKSSFPGNDYSWVKANS--N-W-T--ATAK--VNHY-
GG-ADVVVSGHTESAVGASICRSGSTTGWHCGEVQAKSQTV-NY-Q--EG-------S-VSGLTRTNVCAEPGDS-
G-GSWVS-G---------T--QAQGVTSGGS--------GDCTS---------------GGTTYFQPVNEILSAYNLTLV
>XX00000127 WP_030433063 23 153 194 MYRTV---------------------------TG-------
-----VVLLA----------A---------------------------------G--------V-----------------LTAPV--------AAA---
QENYSPELLHAVQRDLGLDAQQARARLQND--------------------------------------------------------------------
-------------------------------------------------------------------ARAADVEHRARTAAAD-SFGGAWIQ--N-
GKLVVGITDAARAE---T--VRSQGA-E--VAV-VRH--AHSRLKA-AKQQL-DSV-------------------------------------
----------------------------------------AP------AS-VAGWRVDERRNSVVVEVARGA-R----DA-----AD-FLAAAR-
RF----S-P-S-VTV---------EE---------V-----G-EAPRLL-H-DVRGGDA-IY-----N----G----GSPCSVGFTVEG-------
-GGFVTAGHCGDK-GD--N--VK-G-----F---N-Q--VPMGSYAGSSFPANDYAWVKTNA--D-W-T--AKPW--
VNRQ-NG-TNVVVKGSAEAAVGAAVCRSGRTTGWHCGTVLAKDQTV-NY-G--GG-------Q-
VSGLTKSDACAESGDS-G-GSWVG-G--EG----FT--QAQGVLSGGS--------GTCRG---------------
GATSFYQPVQEILTAYGLTLL
>XX00000122 CTQ94560 0 154 171 MATA-----------------------------------------------------
--------------------------------------------------------------------------------
QFEAENPGLLEAMQRDLGLDVTAARDLVATQ---------------------------------------------------------------------
---------------------------------------------------------------------EIAAATENSLRTALGD-SFGGAYFDGAA-
RKLVVGVTDASKAA---T--VTATGA-T--VKM-VAR--SAAQLDS-AVAQL-NARESS------------------------------
-------------------------------------------AP------ST-VTGWYADTVTNEIVVTALPGT-A----PA--A--KS-F---A-----
---G-S-N-VRV---------VE---------A-------EAP--Q-L-FLKGGDA-YTI----G----G----SSRCSVGLTVRGG---------
FVTAGHCASA-GQ--P--------------------SNGTWGGSSFPGNDYGWVRTNS--G-V-T--LEPR--V-------
AAVTVKGSTSAATGSAVCKAGSTTGYTCGTVGAKNQTV-RY-P--QG-------S-VSGLTASNVRCQPGDS-G-
GGFITSA---------G--QGQGVVSGGN--------T-------------------SVCYFQPLGEILSVYSLTLL
>XX00000118 EWM10548 27 155 189 MNLVR-------------L--------------GL------------
SVMVT----------A---------------------------------G-LCAGLGAP-----------------------------------
LATASPVVLAAMQHDLGLTAQQAQARLTEE--------------------------------------------------------------------
-------------------------------------------------------------------SAAMRLAPQAEQRAGD-AFAGSWFDPAT-
GKLVVAVTDQHTAA---T--VRGMGV-Q--VAT-ATV--NAKTLQA-NKSAI-DNRSRAHR-------------------------
---------------------------------------------AP------AE-VASWGVDARTNSVVITVRGQ-------DA--A-VQE-
FVAAAR-K-----N-G-P-VTV---------RQ---------T-----SSPQPQVLSA-GTVGGDP-YYI----N----G----
NVRCSIGFSVSG--------G-FVSAGHCGQP-GY--S--VV-G-----W---D-G--SAMGSFAGSTFPGNDFSYIRTGN--

G-W-W--QAPV--VLGW-GTVSDALVRGDWVAPPGTSVCRSGSTSHWHCGTVEALGDTV-NY-A--QG-------
A-VYNMTRTTVCAEPGDS-G-GSFIT-G---------D--QAQGVTSGGY--------GNCSS--------------
GGETWFQPVGPILAAYGVGLV
>XX00000162 WP_009076014 29 155 189 MRRRIT--------L--SL-------------TG------
------AAVLA----------A--------------------------------S-L-AAVVPP----------------------------------
AAQASPGLVAAMQRDLGLSAGQARTRLGQE-----------------------------------------------------------------
-------------------------------------------------------------------LTASRQLPAAQQAAGA-AFGGAWFDPAL-
GKLVVGVTDPAVAD---A--VHRTGA-E--TVP-AQV--TAAALDA-AKTSL-DRAAKAKQ--------------------------
-------------------------------------------TP------AE-VSGWHADPRTGSVVVTLQPGA-H----GP--D-VDD-
FLARAR-E-----A-G-P-VTV---------ST---------G-----A--RPRTLSA-GTVGGDP-YYI----N----G----
NTRCSIGFSVHG--------G-FVSAGHCGSK-GS--S--VV-G-----W---D-N--SAMGTFAGSSFPGNDYSYITIGN--
G-W-W--TAPV--VLGW-GTVSDVIVRGSAVAPVGSSICRSGSTSHWHCGTVLGLNETI-NY-A--QG-------A-
VYQATHTNACAEPGDS-G-GSFIT-G---------D--QAQGVTSGGF--------GNCGS--------------
GGETWFQPVNEILQTYGLSLV
>XX00000197 WP_035283133 28 155 188 MRRSLV--------------------------AG------
------LVGLA----------A--------------------------------T-AVVTAGQP----------------------------------
ASAAPGALVPALQRDLGLSTDQVEQRLTRE-----------------------------------------------------------------
----------------------------------------------------------------SAARGLLPAARTAAGA-AFGGSWFD--R-
GSLVVALTDPALAG---A--VESTGA-R--STV-VAH--TARALDA-TKAAI-DEKATATG-----------------------------
----------------------------------------AP------KA-VTGWFVDPTTNSVVLTVIKGA-S----GA--E-VTS-
FVDGAK-A-----A-G-P-VRV---------EE---------T-----TA-APRTLGG-PVTGGNA-YYI----N----G----
AGRCSVGFSVSG--------G-FVSAGHCGSS-GD--S--VT-G-----D---D-D--SAMGTFQGSSFPGSDYSWVRTNS-
-S-W-S--GSPT--VNGY-GN-GNVTVTGSSVAGVGESVCRSGSTTGWHCGTIQATDQTV-NY-P--EG-------T-
VSGLTRTNACAEPGDS-G-GSWVS-G---------S--QAQGVTSGGS--------GDCTY--------------
GGTTYFQPVNEILSAYGLSLT
>XX00000200 WP_020659993 30 155 189 MRRRIA--------L--SI-------------TG------
------AAVLA----------A--------------------------------S-LAAVVAPA----------------------------------
AAQAAPGQLAAMQRDLGLSAGQAQTRLRQE----------------------------------------------------------------
---------------------------------------------------------------------LAAARQMPAAQEAAGP-
AFGGAWFDARL-GKLVVGVTDPAAAD---A--VHRTGA-E--TVP-ARI--TAVALDA-AKKAV-DATARTKR------
-------------------------------------------------------------AP------AE-VSGWHADPRTGSVVVTLQPGA-H----
GA--D-VDA-FLARAR-Q-----A-G-P-VTV---------AT---------A-----P--KPQTLSA-GTVGGDP-YYI----N----G---
-NTRCSIGFSVHG--------G-FVSAGHCGGT-GS--S--VV-G-----W---D-N--SPMGTFAGSSFPGNDYSYITIGN--
G-W-W--TAPV--VLGW-GTVSDVIVRGSAVAPAGSSICRSGSTSHWHCGTVLGLNETI-NY-A--QG-------A-
VYEATHTNVCAEPGDS-G-GSFIT-G---------D--QAQGVTSGGF--------GNCSS--------------
GGETWFQPVNEILQTYGLSLV
>XX00000214 WP_052024801 0 155 213 MA------------------------------------------
---------------------------------------------------------------------------------------------------
ADHLSQAFGLSKPEAMNRVREQ--------------------------------------------------------------------------
-----------------------------------------------------EGHSRTALTIAEKLGD-RSPGSFIDQKR-
GKLVVNVSDDEAAT---L--VGGEKV-E--ARV-VGT--TASQLAQ-IRQQA-EQRL----------------------------------
-----------------------------------------G------SM-MRSSAVDTTRNMIVLSVPAKD-L----AA--A--GQ----ATS-
DL------S-K-VEI---------QE---------S-----P-DAMDKK-A-HIGGGDR-IDYKLANG----N----VYSCSYGFSVTKG-
N-E--EG-FVTAGHCAEK-GR--K--FT-K---------E-S--KSLGEIKEFSFPGEDMAYASLDM--TQW-I--GDAS--

VNKMSGG-EMAPVKGSTEAAVGAAVCKSGLSTKWTCGTITAKNATSVNS-P--ND-P-KTRHE-
IKGLTETDVCALRGDS-G-GPWMA-G---------D--QAQGVTSSGN--------ATSFKG--K----
CQNSRGDKIKTHFQPINPILKKYGLTLK
>XX00000237 KDN17180 19 155 176 MAAA---------------------------------------------------
-------AVIGFLT-PVSAGA------TAIDGYSAAVQRE------------A-------V-----A-------------------------------
SLAAEAGISESAAVALLRTQ-------------------------------------------------------------------------------------------
--------------------------------------------------AASVDTLQRVTASLGS-RAADGYLDA-A-AKPVVNVLDQAAAE-
--Q--AVRAGA-Q--ARL-VKH--SSAALAS-AQDAL-QAL------------------------------------------------------------
---------------PA------VA-HTSIGLDPKANQVVLTVAGAA-K----GT--E---A-LLKAAG-SL----G-D-R-VRV-------
--ER---------V-----A-GEM--H-T-AIYNGEA-IT-----G----G----GTRCSAGFNTNRG-G-Q--NY-LIDAGHCTRA-
VS--Q--WN-----------------VGPSQGASFPTNDYGLIRNTT--G-S----APGA--VTLW-NG-
STQRITSAGTATVGQRISKSGSTTHLTSGSVQRTNVTV-NY-A--EG-------S-VYQLIQTNALVNPGDS-G-
GCLFA-G---------S--VGLGITSGMG---------------------------GGSSYFQPVGEALSAYGVTLN
>XX00000108 WP_012924015 24 156 202 MSGRCL------------------------------------
----------------------------------------------------------------------------------------------------------------------------
---------------------------------------------------------------VGVALTGIVALTAAGL---------------------------------
---VAANGAEADERSGTARAGRTMAPDPELLQQERIVKAGTVETQLGS-RVIGSYLSD-A-
GDVVVAVSDQATAG---L--VRAIGA-V--PKL-VRF--TAEELRS-VQRDL-DHL-------------------------------------
---------------------------------------SA------GK-VKSWYVDPITGTVVVTVPTGA-R----DA--I-TKR-FLRRAQ-
AN----G-D-R-VTV---------RT---------S-----P-GEL--TQF-GLLGGLQ-VDK----N----T----
GYTCSLGFNARTDDG-T--RI-FLTAGHCTAG-KP--S--FS-R---------N-G--YILGDTRSSSYPGNDFGTVTVID--
G-W-D--QRGY--VERW-GD-TDVPIKGQTGVTVGASVCKSGKTTRWTCGRVVAKNVTV-NY-G--SN------
QV-VKGMYKHTACVQTGDS-G-GANLT-S---------G--YALGITSGAS--------GECGQ--------------
ANESYSQPIGEALAASGASLV
>XX00000138 WP_033290876 28 157 180 MRTIRF--------L-----------------G-------
----AATAA----------A---------------------------------AVAATFAGIP--------------------------------
AAGAAPLNPGLVPAMQRDLGLTHAQALTRLARE-----------------------------------------------------------------
---------------------------------------------------------------------------DAAGQVGASLEKALGD-
SFGGTSFDAAT-GKARVGVTNPALVD---R--VRAAGA-E--AQV-VRY--SARQLDS-NVQTL-NASEKA----------
---------------------------------------------------------------AP------KS-VTGWGVDTASNRVTLTVLSGQ-R----
EA--A--DT-FVRQSG-VD----T-A-A-VDV---------VE---------T-----P-AAPTPH-Y-NVLGGDA-YYI----G----G---
-SSRCSVGFSVNG--------G-FLTAGHCAAL-TG--GGALT-G-----Y---N-R--
VAMGSFSTYRFPGSDYAFARVNS--N-W-T--PVGQ--IN---NG---
TRVTGSTEAAVGASVCKSGSTTGWTCGTIRAKNQTV-RY-Q--EG-------T-VTGMVRTNARSDHGDS-G-
GSFIS-G---------N--QAQGLLSGGD--------T-------------------VNSFYYPVNRALSATGTTLV
>XX00000234 WP_051116313 35 157 177 MRKTS------------SGRRLFAQSAT-------
-------------------TLVAAAAAIGFLA-PTTATA------TTIEGYSSLVQRE------------A-------V-----A------------------
--------------TLAANAQITEAAATKLLRVQ----------------------------------------------------------------------------
-----------------------------------------------------------------AASVETVQSLTGALGA-RAADGYLDE-A-
AKPVVNVLDKAAAD---Q--VTATGA-Q--AKL-VKH--SSAALAG-AQEAL-QAL-----------------------------------
-----------------------------------------PA------VT-HTSIGLDPKANQIVLTIADAA-K----GT--D---A-LQQAAA-
TL----G-D-R-VRV---------ER---------V-----A-GEM--H-T-AIYNGEA-IT-----G----G----GTRCSAGFNTNKG-G-
Q--NY-VVDAGHCTRA-VS--Q--WN-----------------IGPSEGASFPTNDYGLIRNTT--G-S----GPGA--VTLW-

NG-STQRISSAGNATVGQQISKSGSTTRLTSGSVQRLNVTV-NY-A--EG-------S-VHQLIQTNALVNPGDS-G-
GCLFS-G---------S--VGLGITSGKG---------------------------GGSSYFQPVGEALSAYGVALN
>XX00000243 WP_016333768 25 157 177 MARTAT-------------------------------------
------------TLVAAAAAVGFLA-PSTATA------TTIAGYSPTVQHE------------A-------V-----T----------------------
---------SLASEAGISQAAAVEFLRTQ----------------------------------------------------------------------------
--------------------------------------------------------------TAGVETLQKVAASLGS-RAADGYLDD-A-
AKPVVNVLDQAAAD---Q--VTATGA-R--ARV-VKH--STAALTS-AQNAL-QAL-----------------------------------
-----------------------------------------PT------VA-HTSIGLDPKANQVVLTISDAA-K----GA--E---A-LVKAAE-
AL----G-D-R-VRV---------ER---------T-----T-GEM--H-T-AIYNGEA-IT-----G----G----GTRCSVGFNTNRG-G-
Q--NY-ILDAGHCTRA-VS--Q--WN------------------VGPSQGASFPTNDYGLIRNTT--G-S----APGA--VTLW-
NG-STQRIASAGSATVGQRISKSGSTTRLTSGSVQRLNVTV-NY-A--EG-------S-VHQLIQTSALVNPGDS-G-
GCLFA-G---------S--VGLGITSGKG---------------------------SGTSYFQPIGEALSAYSVTLN
>XX00000025 WP_030431797 24 158 193 MKRHVF---------------------------LG------
------MSVVA----------A---------------------------------T-ALMTVPVT-----------------------------------
AQELSPGLVPAMQRDLGLTEGQALHRIRQE-----------------------------------------------------------------
-------------------------------------------------------------------A--AIRAIDAERVAGA-AYGGSWFDPAK-
GKLVIGVTDEKAAS---A--VRAAGA-E--VAR-VSR--TAAELDV-AQGKV-DAVARTRK---------------------------
-------------------------------------------AP------DA-VHSWHPDPRSGSVVVNVRAGV-N----TP--E-VDA-
FLAEAR-K-----A-G-P-VTV---------RR---------T-----TAPQPKTYAA-GTVGGDP-YYV-PVSG----G----
YVRCSIGFSVHG--------G-FVTAGHCAGA-GA--N--TQ-G-----W---D-W--
SAQGQFAGSSFPGNDYAFVRTGH--G-W-W--TVPV--VLGW-
GAVSDVLVRGQWEAPVGSSICRSGSTTHWRCGQVLAKNETV-NY-G--NG-------
QLVHQLIKTSACAQGGDS-G-GSFVT-G---------D--QAQGMTSGGW--------GDCNS---------------
GGETWFQPLTEVLQVYGLQLH
>XX00000173 WP_030432813 30 158 186 MKPKFA--------V--RY--------------VG----
-------------------SALLALGSAA-ALTL---------------------------------P---------------------A--------
AAAAPGDTFASPEMLTAMQRDLGLTAAEATARATNE-----------------------------------------------------------
-------------------------------------------------------------------------------MKAAEAEAGLRAALGD-
SFAGSHFAHGS-AKLTVSVTDAAKAD---A--VRAAGA-E--VRL-VAR--SSAQLDA-IKSTL-DKADQ--------------
-----------------------------------------------------------HA------SG-VSGWYVDTVNNRVVVTAKVVA-D----
GE------K-FVKASG-AD----A-S-A-VTV---------VQ---------S-----N-EEPTTL-Y-DVRGGDA-YY-----M----G----
GGRCSVGFSVQG--------G-YVTAGHCGTV-GT--A--TQ-G-----F---N-R--
VASGSFRGSSFPGNDYAWVGTNS--N-W-T--PRGV--VNRY-NG-
GTVAVKGSSEAAVGASICRSGSTTGWRCGTIQAKNQTV-RY-P--QG-------T-VNGMTRTNACAEPGDS-G-
GSWLS-G---------N--QAQGVTSGGS--------GNCTS---------------GGTTYFFPVNPILSRYGLRLV
>XX00000028 WP_027946180 23 159 178 MKVPI---------------------------VV--------
----MAIVA----------T---------------------------------ASLAAQPADA---------------------------------
LPSFVPLNAGLLPAMQRDLGLTHDQAVDELTAS------------------------------------------------------------------
-------------------------------------------------------------------------TRASRLEQALRSQLGD-AFGGAMFRPGA-
GRLEVAVTDPAKLG---L--VRASGA-D--VRQ-VRY--SARQLDA-MVARL-NAREAA-------------------------------
-------------------------------------------AP------RS-ITSWGVDTEHNRVTVGVEPGH-R----AE--A--ER-
FLAQSG-A-----P-D-A-FVV----------E---------R-----A-QSPRPQ-F-GVSGGDA-YYV----G----A----
NIRCSVGFAVTI--------G-FLSAGHCAAN-GR--S--IS-G-----Y---N-R--EPMGYFVTYSFPGEDYSLARVNP--N-
W-M--PLGR--LA---NG---TRVAGSIEAPVGASVCKDGSTTGWTCGRIEARNQTV-RY-E--QG-------E-

VVGAIKTDLHAGAGDS-G-SPLVA-G---------N--QAQGLLSGGD--------D-------------------
GSTYFYPVRAALFATGSTLV
>XX00000228 ACY95630 38 159 190 MRRPG----------------------------------------------
---------------------------------------------------------------------------------------------------------------------
-------AGDESRLRTVVLAAGVVVGTAGL--------------------AAGSLYAVQAGE----PAER------------------------
------------PAGTADAALTGAGGTPGEAVRAEPATADRAGADVLAARLARRLGD-RTAGAYLDD-A-
GRPVVTVTNAADAA---M--VRRAGA-V--PKP-VPR--SPATLNR-ITGTL-RRS------------------------------------
---------------------------------------GS------IP-GTGWAIDPAAGQVVMWTDDTV-T----GA--R-MAS-
VRRAAR-QM----G-P-A-VRM---------VR---------I-----P-GRL--R-T-FALGGDA-IF-----G----Q----
GARCSLGFNVRRG-R-Q--NF-FLTAGHCGNV-VR--T--WT-A-----DRR-G-A--
QVLGTTVRSSFPGNDFALVRSAP--G-A-A--GQGA--VNLY-NG-
RAQDIVRAADPVVGQRVVRSGSTTGVSTGRVTAVNATV-NF-P--EG-------T-VTGMIQTTVCAEPGDS-G-
GPLFS-G---------T--TALGLTSGGA--------GDCRI---------------GGVTFFQPVTEALRAFGVEVF
>XX00000232 WP_054290399 35 159 177 MRKIV-------------KGRRLAARSAI--------
------------------TLAAGVAAAGFAA-APVSSA------ATVDGYATEVAQS------------A-------I-----A----------------
---------------DLSAREG-VSSAEAVQILRAQ--------------------------------------------------------------------
----------------------------------------------------------------ETSIRTLDEINGTLGA-RTAGSYLDA-K-
GKPVVNVLDAAAAD---Q--VRASGA-E--AKL-VKH--STQALTS-AQDTL-QSV------------------------------------
-----------------------------------------PA------VK-HTSIGQDPKTNQVVLSVSDAA-T----GG--D-IAA-LLRTAD-
QL----G-D-K-VRV---------ER---------V-----T-GEM--R-L-AIYNGEA-IT-----G----G----GTRCSVGFNTNKG-G-
Q--NY-IVDAGHCTRA-VS--Q--WN------------------VGPSQGASFPTNDYGLIRNTS--S-S----APGA--VTLW-
NG-STQRISSHAAATVGQRIQKSGSTTRLTSGSVQRLNVTV-NY-S--EG-------S-VHQLIQTNALANPGDS-G-
GCLFA-G---------S--VGLGITSGKG---------------------------SGTSYYQPVGEALSAYGVTLN
>XX00000233 WP_042191065 35 159 177 MRKIV-------------KGRRVVARSAI--------
------------------TLIAGVTAAGFLA-APVASA------AAIDGYSPEVAQS------------A-------I-----A------------------
--------------DLTARDGVSPDAAVQILRKQ-----------------------------------------------------------------------
---------------------------------------------------------------DASVRTVGQVTDGLGL-RAAGGYLDT-N-
GNPVVNVLDSAAAE---Q--VRASGA-E--AKL-VRH--SSTALAS-AQSTL-ESV--------------------------------------
---------------------------------------PS------VT-HTSIGLDPKTNQIVLSVSDKA-T----GG--D-LTA-LLRTAD-RL--
--G-A-Q-VRV---------ER---------V-----T-GEM--R-L-AIYNGEA-IT-----G----G----GTRCSAGFNTNKG-G-Q--
NY-IVDAGHCTRA-VS--Q--WN------------------VGPSQGASFPTNDYGLIRNTS--G-S----GPGA--VTLW-NG-
STQRISSHAAATVGQRIQKSGSTTRLTSGSVQRLNVTV-NY-A--EG-------S-VHQLIQTNALVNPGDS-G-
GCLFA-G---------S--VGLGITSGKG---------------------------GSSSYFQPVGEALSAYGVALN
>XX00000245 WP_009152343 28 159 176 MRTVSR---------------------------S--------
----AI--T----LAASALALGLLA-PTVASA------TSIEGYSSDVQQA------------A-------V-----A----------------------
---------EITSTEGITPAAAEEILRVQ------------------------------------------------------------------------------
--------------------------------------------------------SDSVETLNGVLDQLGD-QQAGGYLDD-S-
GMPVVNVLDEATAA---E--LAPSGV-A--ARV-VDH--SAAELRS-AREAL-EAV------------------------------------
---------------------------------------PA------VA-HTAIAVDPETNQVVLTVAEEA-D----DA--A-AAD-LLAAAA-
RF----G-D-A-VRV---------EH---------V-----T-GGL--H-K-AIYGGEA-IT-----G----G----GSRCSAGFNVNSG-G-
Q--LY-IVDAGHCTGA-VS--Q--WN------------------VGPSVAASFPGNDYGLIRNDT--G-S----GPGA--VTLW-
DG-SAQAINSASNATVGQRICKSGSTTGLTCGVVQATNVTV-NY-S--QG-------A-VHQLIQTSASVNSGDS-G-
GSLFA-G---------S--TALGITSGMG---------------------------GGSSFFQPVVEALNAYGVSLN

>XX00000120 WP_053734875 23 160 188 MSRI------------------------------------------
--------VPALAIVLVGSTFT-APAQAA-----------------------------------------------------------------
AQELSPGLLSAMSSAFGLTEQQARVRLASE--------------------------------------------------------------------------
------------------------------------------------------------------------QRASELAPRVETEAGT-AYAGSWFEE---
NRLVVAVTDRAAAR---R--VEATGA-V--ARV-VGR--SAGSMAA-HKNRL-DALSATGA--------------------------
-------------------------------------------------VP------AA-VASWHPDPRSGTVVVNVEAGK-Q----TA--E-VAA-
FLDRAR-Q-----F-G-A-ISV---------NT---------E-----GVRQPRPFAA-GTVGGDP-YY-----T----G----
NVRCSIGFSVHG--------G-YVTAGHCGGN-GA--A--VY-G-----W---D-R--
SFQGNFAGSSFPGNDYAWVRTGG--G-W-W--TVPV--VLGW-
GKVSDVLVRGSWEAPIGTSVCRSGSTTKWHCGVIQSRNETI-RY-A--QG-------D-VHQMAGTSVCAEGGDS-
G-GSFIT-G---------D--QAQGVTSGGY--------GNCSS---------------GGRTWFQPINEILSVYGLRLH
>XX00000133 WP_020666418 30 160 188 MKHKNL--------IRALA--------------VA---
---------GVSMG----------------------SA-------------------V-A--------L-----------------SMSPA--------TAA---
GPQLSQGVLTAMQRDLGMEPGQVLTRLATE----------------------------------------------------------------------
---------------------------------------------------------------------------SAASRALPAAQSAAGA-AFGGAWFDTAA-
GKLVVGLTDQSAAD---A--VRATGA-E--TVR-VAN--TEAALDT-AKAAI-DRFAGQT------------------------------
------------------------------------------AP------KA-VTGWYVDAKSNSVVVKVRDGA-Q----DA--E-VGK-
FLDFAK-S-----A-G-P-VRV---------EQ---------A-----K-AAPRVLAG-DVVGGDA-YYI----G----G----
SARCSIGFSVNG--------G-FVSAGHCGSQ-GD--S--VT-G-----S---D-Q--SDMGSFAGSSFPGDDYSHVEVNS--
S-W-T--PTPV--VNGY-DQ-GDVTVTGSSESAVGASICRSGSTTGWHCGTVQAKNQTV-NY-A--EG-------T-
VTGLTQTDACAEPGDS-G-GSWVS-G---------T--EAQGVTSGGS--------GDCTS---------------
GGETFFQPVNEILSAYSLTLV
>XX00000191 WP_027941817 29 160 187 MKRLFA--------A--RV--------------VG----
--------AALLA----------A---------------------------------G--------L-------A---------ATPAV--------
TAQAATPMTVAPEMLQAMQRDIGLTADQAIKRMTSE--------------------------------------------------------------
-------------------------------------------------------------------------------VKASEQEQSLRQELQG-
AFGGAYFDATT-NQLVVGVTDAAKAG---V--VRAAGA-K--AVQ-VGR--SEQQLDA-VKAGL-DQNAAQ-------
-----------------------------------------------------------------AP------KT-LTSWYVDVTKNDVVLTVTPGA-V----
AD--A--RS-YLAATG-TD----A-A-A-VEV---------VE---------S-----P-EQPQPL-F-DVRGGDA-YF-----I----G----
SSRCSIGFSVQG--------G-FVTAGHCAAL-TS--G--SA-G-----S---N-G--VALGTWGGFTFPGNDYAWVRTNS-
-N-W-T--PRGV--VNRY-NG-TTVPVSGHTEAAVGSSICRSGSTTGWHCGTVQAKNQTV-TY-P--QG-------R-
VLGLTQTNVCAEPGDS-G-GSWLT-G---------N--QAQGVTSGGS--------GNCTV---------------
GGTTFFQPVNPILSRFGLTLV
>XX00000011 WP_006502626 28 161 226 MNKRI----------------------------------------
------GA----------T-----------L-------------------A-L--------L-------S-----
SAALLPAAAPLTAQAADEKKSEELSTIQEMSAEWLSKSYGLDGEEAKRRVKDQ---------------------------------
-----------------------------------------------------------------------------------------------------------
DGHAQKAKDIEGDLGD-KTAGSFIDQKR-GKLVVTVTDEKAVD---K--VKEKNA-D--VRV-VKN--SASKLND-
SKKNA-EDKV------------------------------------------------------------------------G------DK-
MAASYVDVERNVVVLTVPKDK-A----EE--A--KK----QVK-DV------Q-G-VEV---------QE---------V-----E-
ATIEAQ-A-NLYGGQE-IQ-----F----G----RSVCSVGFPATKD-G-K--NV-FITAGHCAAG-GQ--A--FR-R---------
N-G--QNLGKPVKYNFPGPDMAYSTMEN--G-W-N--GVGA--VDHW-NG-
KAVAVAGSQEAPVGATVCKSGRTTRWTCGVIQAKNVTV-RY-G--KP-G-GGQDI-

VRGMTQANVCSEGGDS-G-GSWIA-G---------N--QAQGVTSGGA--------GYGPNK--S----
CGEKVGRPNVAYFQPLNPILKDYGLKLT
>XX00000012 WP_026125632 29 161 193 MSTSPT--------V--RT--------------IG------
------GTVLA-----------LG----------------------------------------L---------------TFAVAPA--------
AHAAPEPPVASPEQLDAMQRDLGLTKAGAVDLIESE------------------------------------------------------------
---------------------------------------------------------------------------TEAQAVEKKLRKALGD-
DFGGALFDIED-QELTVQVTDTKAVK---K--VKAAGA-E--AEV-VEH--GEAELDR-VMDAL-DDSEEA-----------
-------------------------------------------------------------AD------AA-IAGWRVDPASDAVVVTVAKGE-A----
AE--A--ED-FIADAG-VD----A-D-A-VKI---------EE---------S-----S-ENPRTY-A-DLIGGNP-YYF----Q----DG-
GSWYVCSVGIPVVG--------G-YVTAGHCGDA-GS--S--TW-E-----RPN-N-S--
TQIGTVVQSNFPGQDSGFVRVTN--SSF-T--AVPQ--INDY-SG-
GTVDVTGSAEASVGASVCRSGQTTGWHCGTIQSKNQTV-RY-P--EG-------T-VRGLTRTNVCAEGGDS-G-
GSWVS-G---------T--QAQGVTSGGS--------GNCTW---------------GGTTYFQPLNPILNQWNLNLL
>XX00000074 WP_017624479 29 161 185 MRKSPF--------I--RF--------------LG------
------GSTLA----------V------------------------------------G--------L-------IV---------AAPT--------
GAQAEAFPEGTEPLAQALERDLGVASGQADDLLTAE------------------------------------------------------------
---------------------------------------------------------------------------ADARSVQKAAEEAAGA-
AFAGAVFDTET-RELTVSVADSSAVE---A--VEATGA-E--ARV-VES--SQDDLEA-VMADL-DAEAEDG---------
--------------------------------------------------------------VS------DK-VTGWYVDVESNSVVVEAVDGA-E----
GL--A--ED-LVAEAG-VD----S-S-R-VTV---------EE---------G-----A-DQPETF-A-AIVGGDA-YY-----P----G----
NSRCSIGFSVSG--------G-FVTAGHCGST-GT--S--VR-G-----S-----A--GESGRVAGSIFPGRDMGWVRASS--
G-W-T--PSPY--VNNY-RG-GRVTVRGSQEASVGASVCRSGSTTGWHCGTIQAKNQTV-NY-P--QG-------T-
VRGLTRTTACAEPGDS-G-GSWMS-G---------N--QAQGVTSGGS--------GNCSW--------------
GGTTFFQPVNPILNQWGLSLT
>XX00000209 EKC92181 35 161 188 MRRTTT--------T-----AARVGLSSLLLAGSW------
------AAFGL----------T-----------------------------------P-AATAAPDP-----------------------------------
APVASGAVLAGMERDLGLTPAEAKARLAAE------------------------------------------------------------------
------------------------------------------------------------------KEATAVEKKAQRAAGA-AYGGSWFDAEK-
GALVVAVSDRAAAD---A--VTGAGA-S--VRL-VEH--SAKTLDA-AKERI-DGLD-----------------------------------
----------------------------------------AP------RG-VAGWRVDPQANSVVVSVVGDQ-A----DK--A-VRG-
FVERAR-A-----A-G-P-VEV---------ET---------V-----A-AAPETFAA-GTVGGDP-YY-----T----G----
NVRCSIGFSVHG--------G-FVTAGHCGPT-GQ--G--VS-G-----W---D-G--
SGIGSFQGSSFPGDDMAYVSVGN--G-W-W--TVPV--VLGW-
GTVSDQLVRGSAEAPVGASICRSGSTTHWHCGTVLAKNETV-NY-S--QG-------A-VHQMTKTSVCAEGGDS-
G-GSFIS-G---------D--QAQGVTSGGW--------GNCSG--------------GGETWYQPVNEILGRYGLTLH
>XX00000213 WP_040276768 28 161 184 MRKSPQ--------F--KA--------------LG----
--------GIALS----------V------------------------------------G--------L-------IA---------AATS--------
GVAADTAPAAAAPEQLDAMQRDLGLTESQANKLLKDE---------------------------------------------------------
-----------------------------------------------------------------------------SKARSLENELRGDLGS-
DFGGAVFDAKS-GELTVSVTDAEAAE---T--VEDAGA-E--AEV-VTH--GEDALDK-VVKKL-NADAEE-----------
-------------------------------------------------------------AG------SG-VTGWYADVEADNVVMTVEKGS-A----
KA--G--KA-FLADAG-VD----K-A-A-VKV---------KE---------S-----S-EKPKTF-A-DIVGGDA-YYI----N----G----
SSRCSVGFAVTT--------G-FVTAGHCGSS-GA--S--AS-G-----A-----N--GGSGTFQGSEFPGSDMGYVGATS--
N-W-N--PTPR--VNNY-SG-GTVAVNGSSQAAVGSSICRSGSTTGWHCGTIEARNQTV-RY-P--QG-------T-

VYGLTRTSVCAEPGDS-G-GSFIS-G---------N--QAQGVTSGGS--------GNCSS--------------
GGTTYYQEVNPILQRWNLSLV
>XX00000241 WP_006238056 28 161 176 MRELLR--------------------------S--------
----AV--A----AGACALLAGALA-PATAFA---DPTVTIEGYSTDVRAE------------A-------V-----S------------------
------------ALVAERGLGTASAEALLRTQ--------------------------------------------------------------------------
-------------------------------------------------------------EAATSALDAALAQLGD-SAAGGYLDE-S-
GAPVVNVVNESAAT---S--V--EGV-T--TRL-VTR--TTAELES-ARDEL-EAL--------------------------------------
-----------------------------------PQ------VS-HTAIAVDPVTNQVVVTVGEAV-I----GA--G-TAG-LLAAAE-RM-
---G-D-T-VRI---------EH--------V-----A-GGF--E-P-AIYGGEA-IT-----G----G----GSRCSAGFNVNSG-G-Q--
DY-IVDAGHCTAA-VS--Q--WN-----------------VGPSVDASFPGDDYGLIRNDT--G-S----APGA--VSLY-DG-
SVQEIASASDAHVGQSICKSGSTTGLTCGTVEATNVTV-NY-P--QG-------A-VHELVQTSASAGSGDS-G-
GALFA-G---------S--VGLGITSGMG--------------------------EDSSFFQPLTEALGAYGVRLN
>XX00000021 AHE78425 23 162 219 MRKFS------------M-------------------------------
---------------------------------------------------------------------

SVLAATVAMTVSGAAFAADDLSPALKTAMQRDLGLSSTQLSQYLKVE-----------------------------------------
----------------------------------------------------------------------------------------------

RLAEQQEKIVQAQLGR-NFAGAWVERKADGSFGFVAASTSIKN---A--KAVAGV-E--TRQ-ARH--SLAALNS-
AKAQL-DSLLARSAK--------------------------------------------------------------------VP------KG-
VYSWTVDLPSNSVVVGVAPGA-E----EA--G--VD-FVALSG-LD----A-R-V-VRF---------ET--------L-----N-
EAPQRR-I-AVQGGRG-MLR----DPGDGY----LYACSVGFTVTKG-S-T--PG-YATAGHCGTV-GE--I--VY-
QEVSQWN---P-G--VRVGTFSASTMPGGDRGWVKVDT--T-H-T--LSPS--VYGY-GN-
GDVTVRGSTEAAVGAALCRSGRTSGWHCGAIRSKNNTV-SY-VDDAGNPDG---T-VTGLTRTSACAEGGDS-
G-GSFIT-G--------AG--QAQGVLSGGS--------GSCSGSQGN---------NGGGNSYYTPINSILSAYALTLR
>XX00000024 WP_011875063 22 162 190 MKRTI-----------RL-------------AG------
-----VAVLA----------A--------------------------------G-------------------------TIAAI--------
GAPTVGAEPVSPDLVAAMERDLGISAQQAHARLAQE----------------------------------------------------------
-----------------------------------------------------------------------------ATAMRADAELSRSLGE-
SFGGSYFDAAR-GKLVVGVTEQADAA---K--VRAAGA-E--AAV-VPN--SLRELDA-TKAAL-DAMDAA---------
--------------------------------------------------------------AP------AS-VTGWYVDVPSSSVVVSVNGRD------
AA--T--DA-FLDKAK-AA----G-D-S-VRV---------QE---------V-----A-ESPRPL-Y-NVVGGDA-YY-----M--------
GGRCSVGFSVRSSSG-Q--AG-FVTAGHCGTR-GT--A--VS-G-----Y---N-Q--
VAMGSFQGSSFPNNDYAWVSVNS--N-W-T--PQPW--VNLY-NG-
SARVVSGSSAAPVGSSICRSGSTTGWHCGSVQALNQTV-RY-A--EG-------T-VYGLTRTNVCAEPGDS-G-
GSFIS-G---------N--QAQGMTSGGS--------GNCSS--------------GGTTYFQPVNEALSAYGLSLV
>XX00000077 WP_052592027 30 162 197 MRLNPS--------------------------T-------
----IATAA----------A-----------S------------------A-G--------L-------L-----
VTALATSPSAGAATHDPTPSTERSVNVPEMTARWLAQDRGISLTEARDQVAAQ------------------------------
---------------------------------------------------------------------------------------------------------
PGQTATADTLERSLGSSRSAGSYVDA-R-GMLVVNVKDASAAD---A--VRAKGA-T--PRL-VKR--SGADLAT-
IEQTV-RRVL-------------------------------------------------------------------------G------TG-
FVSIGSEPVSNAVKVVVAGSD-L----AA--A--KA----ALA-DV------S-G-ALV---------RG--------T-----T-
AKVSTQ-A-NVYGGQQ-IE-----F----S----GYVCSLGFNARQG-S-T--PV-FVTAGHCGEG-YQ--T--FR-K---------
N-G--TTLGVTKAYSFPGNDYAYATLSS--G-W-T--GIGA--VDLY-NG-

TARSVKGSSNAAVGTAICKSGRTTGWTCGYVKAKNQQV-NY-G--NG------DV-VSGLTLTNTCTEGGDS-G-
GSWMA-G---------S--YAQGVTSGGA--------SIN--G--R----CLEVYGRENVAYFQPIGEILSAYGLTLT
>XX00000132 CEJ90055 24 162 190 MELTKF-------------------------------------------------
------------------------------------------------------------------------
LSFLAAVLPVAYGAPTQAVNELHPDILAAMQRDFGLNAEQATARVAQD------------------------------------------
--------------------------------------------------------------------------------------------------------
IQASSLIEKLRGSAGP-SFAGGWIDG---GKAVIGVTDQAMAD---Q--ITAQGG-V--PVV-MKT--PLAKLEA-
AKTAI-DN-IFLGGGAKRS---------------------------------------------------------------------AN-----D-A-
VAAFYVDEAANKVVFEALASG-H----GQ--A--AD-LAKQAG-LD----A-S-Q-YAI---------KT---------V-----S-
EMPTVR-I-TIKGGDA-YHI----N----N----QFVCSVGFSVNG--------G-FVSAGHCGKK-GA--D--VE-T-----T---
S-G--AHLGTFAQSVFPGNDMSYIKTIA--G-T-K--LTGY--INGY-GH-
ADYPVRGHQESAVGASVCRSGQTTGVHCGTIQRKNATV-NY-G-ADG-------S-VTGLTQTNACADHGDS-G-
GGWFS-G---------T--QAQGVTSGGS--------GNCKG---------------GATTYFQPVNEILSTYGLTLV
>XX00000140 WP_053616577 29 162 192 MSISPL--------G--RL--------------AA------
------TAALA----------S----------------------------------G--------L-------V-------LAPVTAA--------
SADPSATPALDPEQIAAMQEEFGLSEAGVTDLLDAQ----------------------------------------------------------------
----------------------------------------------------------------------------------AEAAELEADLREELGA-
DFGGAVFDTET-RALTVRVTDPSSGE---A--VREAGA-V--PEH-VEH--GESELEA-AVEAL-NGAEEA------------
----------------------------------------------------------------AS------DG-VHGWYADPELDTVVIEAAEGA-A----
GD--A--EA-LAAEAG-LD----V-D-T-FTV---------EE---------G-----T-EEPRTY-A-DIVGGSP-YYF----Q----QG-
SDWFVCSVGFGVVG--------G-YVTAGHCGAA-GS--S--TW-L-----QVG-G-P--
DQLGTVAESVFPGQDAAWVRSGA--G-H-T--ATPL--VDDH-NG-
GAVTVTGSQETPVGGAVCRSGQTTGWHCGTIEAKDQSV-NY-G--GD-------V-VNGLTRTTACAEGGDS-G-
GSWVT-G---------T--EAQGVTSGGS--------GNCTW---------------GGTTYFQPVNPILEQWNLTLL
>XX00000154 WP_030470942 32 162 189 MNRTSA--------A--RI--------------TG-----
-------AVLLA----------TGVVTALSVPAFGA-------------------S-A--------A-----------------PVAPT--------GSD---
TESYPAELLQAVQRDLGLNAEQARVRMASD--------------------------------------------------------------------------
------------------------------------------------------------------------QRSSELAEQAKASLGD-AFGGAWIDQAS-
GKLTVAVTKSG---------LSVTGA-D--VKV-VKN--SLTQLNG-TKSVL-DLMS---------------------------------------
--------------------------------------AP------SA-ITSWRVDEKSNSVVVEVNKNT-R----DA--A-AEK-FIATAK-SL---
-G-S-A-VTV---------EE---------V-----T-ENPTPL-Y-DLRGGDA-IY-----M----G----GGRCSLGFSVQG--------G-
FVTAGHCGTT-GT--A--VS-G-----Y---N-R--VAAGTFAGSTFPGNDYSWVRTNA--N-W-T--PRPW--VNRY-
GG-ANVIVKGSSEAAVGAQICRSGSTTGWRCGTVQAKNQTV-SY-P--QG-------T-VRGLTRTNACAEPGDS-
G-GSWVA-G--SG----FG--QAQGVTSGGS--------GNCTS---------------GGTTYFQPVNPILSRYGLRLV
>XX00000215 WP_051772033 0 162 213 MLA------------------------------------------
-------------------------------------------------------------------------------------------------------------
AMQRDLGLNAEQANRRAVQQ-----------------------------------------------------------------------------------------
--------------------------------------------------------EAADRLDDVLKARLGA-AFGGAWFDGAS-
GKLVVGVTDAGRAD---A--VTAAGA-Q--SRV-VKH--SENDLEA-VKAEL-DALAGGRGDAQRKAAPGRADK--
-----------------------------------------------------------AA------VEGLTSWSVDPVTNTVTVTALKGA-K----G-------
---LATLA-KY----G-D-A-VRV---------ET---------T-----A-TEPQTT-A-FLDGGDD-LN-----Y----P----
NGGCSAGFNMRNTTG-V--RY-VLTAGHCGAA-GS--V--VR-G-----Q---G-N--
VVIGSVQAAFFPTYDDGLVRVTN--T-G-AWTQGPWVDVNPP-NG-

GIVVVSGYSDSPVGTAICKSGRTTKLTCGTITAKRQTV-TY-T--GG------LT-VYDLTRHNACVEPGDS-G-
GSNYR-N------AGTR--TAEGVTSGAQLYNVGG-RARCGQV-----------VGVANVSWYYPAALSVGAYGATLW
>XX00000240 WP_012795865 28 162 176 MKKLMK---------------------S---TVA-----
-------AG--A-------CALVFGVVV-PSTASA---VTEPRIEGYSANVQNE-----------A-------V-----S-----------------
--------------ALIAERGVTQAEAETILRTQ----------------------------------------------------------------------------
-------------------------------------------------------------QHSVQALDTVLAQLGE-RAVDGYLDA-S-
GQPVVNVVSEAAAA---V--VERAGV-T--AKV-VEH--TAAELEA-ARAAL-ESL-------------------------------------
----------------------------------------PS------VA-DTAIGTDPTTNQVVLTIGAEA-D----DA--E-VAE-LLAEAE-
RF----G-D-R-VRV---------DR---------V-----A-GGF--D-V-AIYGGEA-IT-----G----G----GSRCSAGFNVNSG-G-
Q--NY-IVDAGHCTGA-VS--Q--WN-----------------VGPSVGASFPGDDYGLIRNDT--G-S----APGA--VSLY-
DG-TVQQINSASNAYVGQSICKSGSTTGLTCGTVQAINVTV-NY-P--QG-------A-VHELVQTSASVNSGDS-G-
GALFA-G---------S--VGLGITSGMG--------------------------GGSSYFQPLTEALSAYGVSLN
>XX00000055 EDY63586 0 163 181 APRRST---------------------------------------------
-------------------------------------------------------------------------------------------------------------------
------V----RI-K-RT--RA--LLAA---------------GAALAV-T-AA-----V-A-----PS-------TA-QA-T--P----
SPGPAQLAR-ADAAVLA------------------------------------------------------------------------------
---------------------------------------------------------------------------------------AG------VE-
GTAWYVDRTLGRVVVTADSTV-S----DA--E-ITG-IRSAAG-AD----A-G-A-ITV---------KR---------A-----P-
GVF--K-P-LLGAGDA-IY-----G----S----GYRCSLGFNVRSG-S-T--YY-FLTAGHCGDV-AP--T--WY-A-----NSA-
Q-S--TLIGPTVGSSFPSNDYALVRYDN--A-S-LS-HTGG--------------
FTAADAYVGESVKRSGSTTGTHSGTVTALNVTV-HY-S--GG-------GTVRGMIQTNVCAEPGDS-G-GALYD-
G---------T--KALGLTSGGS--------GNCST---------------GGTTFYQPVTEALAKYKVSLY
>XX00000185 WP_005443179 28 163 176 MKRLTK---------------------T-----T-------
-----VA--A-----GACALAFGALA-PAMASA--ASTPAEVEGYSAD--------VQAA----A-------V-----S---------------
-----------------ALVEERGIDEAEATALLGSQ--------------------------------------------------------------------------
--------------------------------------------------------------DAAVAALDAALQQLGD-RAAGGYLDA-S-
GKPVVNVVSDAAAEVPSL--VDGTGV-A--TKL-VEH--STAELEA-ARAAL-P--------------------------------------
----------------------------------------T------VV-GTAVATDPTTNQVVVTVADEA-N----DA--E-VAE-LTAAAQ-
RL----G-D-T-VRV---------DH---------V-----A-GGF--D-T-AIYGGEA-IT-----G----G----GSRCSAGFNVNSG-G-
Q--NY-IVDAGHCTGA-VS--Q--WN-----------------VGPSVGASFPGDDYGLIRNDT--G-S----APGA--VSLY-
NG-STQAISSAGNASVGQSICKSGSTTGLTCGTVQATNVTV-NY-A--EG-------A-VHQLVQTSASVNSGDS-G-
GALFA-G---------S--TALGITSGMG--------------------------GGSSFFQPVTEALSAYGVQLN
>XX00000196 CEJ93563 24 163 191 MELTNL---------------------------------------------
---------------------------------------------------------------------
LSFLAVMLPVAYGAPTQAADSLHPEILNAMMRDLGLNAEQAVARVSHD---------------------------------------
-----------------------------------------------------------------------------------------------
NQATELISQLSGSIGD-KFAGGWIDA---GKTYVGVTDQATAE---H--VKAAGA-V--PVV-MNT--SLSKLEA-
AKEAM-DQHLSSAIQARD------------------------------------------------------------------TD-----TTG-
IASYFVDVTANKLVVESLADS-Q----ER--A--HA-LAARGN-LS----E-G-D-YEV---------RI---------V-----K-
SLPSAA-A-SVIGGNS-YVI----N----G----KTRCSYGFAVSG--------G-FVSAGHCGNQ-GD--R--VT-AG-A-SA---
G-G--EYVGDFAGSTFPGKDMSYVKTAS--S-V-Q--LYGA--INNY-NG-
GTIPVKGSQEAAQGASVCRSGYTTGVRCGTITGKDQTV-IL-D-GVN-------K-VYGLSRSSACCDHGDS-G-
GSFYS-G---------S--QGQGVTSATN--------GGCSD--------------GGYSYFQPLNPILQNYGVSLI

>XX00000242  WP_030532933  26  163  176  MKTSRK--------------------------T--------
----AI--T----AVAGALALGMFA-PATAAS---ASVPGIEGYSDVVA------NE------A-------I-----T------------------
------------TLSSERGVSEVAAATLLQTQ----------------------------------------------------------------------------
-------------------------------------------------------------QNSVDALDELRSSLSS-DIVSEYLDA-S-
GKPVVNVATKAAAA---K--AEAAGA-T--AKV-VSR--TAEQLDA-ARAQL-ES------------------------------------
---------------------------------------VS------VA-HTSVGLDPQNNQLVVTIADQA-K----DS--A-AAK-LTQKAA-
QL----G-D-A-VRV---------EH---------V-----A-GGM--E-K-AIYNGEA-IT-----G----G----GSRCSAGFNVNDG-
S-Q--DY-ILDAGHCTAG-VA--E--WN-----------------VGPSIDASFPGNDYGLIRNDT--G-S----APGA--VTLW-
DG-TTQSISSASDATVGQDICKSGSTTQLTCGTVQATNVTV-NY-P--EG-------A-VSGLVQTSAEVNSGDS-G-
GCLFS-G---------S--TGLGITSGMG--------------------------GGSSYFQPVTEALSAYGMSLN
>XX00000009  CEJ93600  24  164  188  MELTKF---------------------------------------------
-------------------------------------------------------------------
LSFLAIALPVAYGAPAEAATTLHPNMLKAMKRDLGLDAKQAVARIAED----------------------------------------
-----------------------------------------------------------------------------------------------
LQASKVLDDLSGSLGP-AFAGGWIDA---GKSFVGVTDQAMAQ---K--VTAAGA-T--PVV-MIN--SLSRLNE-
AKTAI-DNKLKGHQTRADA----------------------------------------------------------------GS-----ITG-
VAAYYVDVAANKVVFEALAAN-H----GQ--A--KD-LAAHAG-LS----A-T-E-FEI---------RT---------V-----E-
NMPTPR-Y-NIVGGDP-YNS---IR----A----SLRCSVGFSVTG--------G-FISAGHCGSQ-GD--G--VQ-S-----T---
G-G--VDLGTFSGSSFPSVDMSYIKTLD--S-V-T--LTGY--VNDY-NG-
GSYSVSGGSEAAVGANICRSGQTTGVHCGTIKQKDVTV-NY-A--EG-------T-IYGLTQTSACSDHGDS-G-
GSFLA-G---------T--QAQGVLSGGN--------GNCNN--------------AGISYFQPLGPILSTYGVSLI
>XX00000056  EWM14380  30  164  177  MMRIAA--------A--RI-------------TG-----------
AALLS----------I--------------------------------G-LAATVSVP--------------------
AAAASAPVSVDTSTNGVVISPDVASALQRDLGLSPARAMVTFAQQ-------------------------------------------
--------------------------------------------------------------------------------------------
NRADVQVATLRTVLGP-IFAGSWFDAAA-GEAVVAVTDVTALA---R--VRAAGV-E--ARV-VTR--TDAQLTA-
VADGL-RQKQ------------------------------------------------------------------------VP------AG-
VVGWYVDLPTDQVVVQILDRS-P-----A--T--EA-FLAT----A----G-A-A-VRA---------QA---------V-----T-
ERPKPL-Y-AVRGGDA-WY-----G----S----NFRCSVGFSATDAAG-G--KH-FVTAGHCTAG-GG--T--AS-G-----
Y---N-Q--VSLGRINGSHFSGGDYGKVDVTS--GQW-T--LSGT--VA-S-GG-
GAVAVKGSAEAAVGAAVCRSGSTTGWHCGTIQAKNQTV-VY-Q--QG-------T-VTGLTRTNVCAEPGDS-G-
GAWIS-G---------N--QAQGVTSGGS--------GDCTH--------------GGTTYFSP------------
>XX00000101  WP_012922340  22  164  189  MKIL----------------------------------------
-----------VSTALVASLLA-VPA--------------------------QAMAASPE----------------------------------
QPLAVPRQTAALQKDLGLTEQQVKELLRAE----------------------------------------------------------------------
------------------------------------------------------------------AAATKLVPAAQRAAGT-AFGGTWYDAAR-
QKLVVGVTSPAAAD---A--VRATGA-E--ASV-VPV--TAAVLDR-RKAAV-DKLAGTN----------------------------
-------------------------------------------VP------AA-VSGWSADPRTGTVVVNVQAGK-R----SA--A-VDA-
FVDKVA-K-----A-G-A-VSV---------RE---------V-----AAQAPRTFAA-GTVGGDP-YY-----T----G----
NVRCSIGFSVHG--------G-FVTAGHCSGA-GA--R--TR-G-----W---D-G--
SAQGVFRGSSFPGNDYAWVQVDS--G-W-W--TVPV--VLGW-
GQVSDQLVRGSWEAPVGTSVCRSGSTTHWHCGHIRAKNETV-NY-G--NG-------
QLFYGATKTSVCAEGGDS-G-GSFIT-G---------D--QAQGVTSGGW--------GNCSS--------------
GGETWFQPVNEILGAYGLTLH

>XX00000159 WP_005464557 29 164 184 MNRKTA--------A--RL-------------IA-----
-------SVTLA----------A-------GTAVAF-------------------T-L--------P----------------ATA-----------
APATEVSTTAADPVIQAMQRDLGLTKAEAEQRLRSE-----------------------------------------------------------------------
-------------------------------------------------------------------------------------------AEAREVHKAVTKELGA-
DFAGAHYDAAL-GKLVVGVTDTADFA---E--VRAAGA-E--PRL-VEH--TVAELEK-AAKAL-DAKESS-------------
-----------------------------------------------------------------------AP------DA-VTGWYVDVEANSVVVTTAMGT-A----
EQ--A--ER-FVTRAG-VD----A-D-V-VDV---------VE---------S-----T-ESPRTF-M-DIIGGNA-YYI----G----T----
GARCSVGFAVQG--------G-FVTAGHCGST-GA--T--TS---------------SPSGRFAGSSFPGNDYAYVQTGS--G-
D-T--PRGL--VNMY-NG-SARVVSGSTVAPVGSSVCRSGSTTGWHCGTIQATNQTV-RY-A--EG-------T-
VSGLTRTNVCAEPGDS-G-GSFIS-G---------N--QAQGMTSGGS--------GNCTW--------------
GGTTYFQPVNEVLNAYNLRLV
>XX00000081 WP_052385749 28 165 204 MRKFAS-----WARRGLLA-------------------
-----------------------------------------------------------------------------------------------------
ISPLALFAAMAASAAPGDRAMLPAVREAMQRDLGLTSEQLTQYVQAE-----------------------------------------------
-------------------------------------------------------------------------------------------------------------------
RLALREEKALAKAQGE-RFAGSWLERKGNGQYQLVVATTSITP---Q--RAPAGA-E--IRS-VRH--SLDRLEA-
SKGEI-DTLLAHGAK---------------------------------------------------------------------------------SP------EG-
VYGWFVDVRNNSVTVNVAPGA-E----AA--A--IG-FIAATG-AD----A-D-T-IRF---------QR---------M-----E-
SAPMPL-A-TLQGGSE-YLS----NDG-SN----YYYCSVGFAVTRN-G-A--KG-FATAGHCGNA-GD--R--AF-
VAVN-RR---T-V--SQIGAFAASNFPGSDRAWVQVDN--N-H-T--LLPV--VDGY-SG-
SDIPVYGSTEAPIGASVCRSGRTTGLHCGVIEAKNVTV-QY-P--DA-------S-VNGLTQVKVCAEGGDS-G-
GAFIT-G--------AG--QGQGVLSGGN--------YSCKGK--Q---------AKLATSYFQPLNALLQGYGLTLS
>XX00000091 WP_051060818 0 165 201 MVSA---------------------------------------------
------------------------------------------------------------------------------------------------------------
DVEPVLRDPEQLAALQRDLGLTRTEAVDLVVRE------------------------------------------------------------------------------
------------------------------------------------------------------------------------RQARATERELRSALGP-DFGGAVFDADT-
RELTVAVTDPDAAE---A--VRDAGA-Q--ARV-VTY--GEDALGR-VMDEL-NGSVGT---------------------------------
--------------------------------------------------GY------PG-VTGWYPDVEQDAVVMTVLPGG-S----DD--A--QQ-
AIAAAG-VD----P-E-A-VVI---------SE---------V-----T-EPPRTF-V-DIVGGDP-YFI----N----D----
QSRCSIGFGVTEGSD-N--GG-FVTAGHCGSV-GS--T--AS-S-----SAE-G-S--
EATGTFVRSMFPGDDMAFVGETV--N-W-T--PTAQ--VR----S-
SPEDVAGSEEASVGASVCRSGSTSGWRCGEVLAKDQTV-IY-E--QG-------I-VFGLTHTSACAERGDS-G-
GPFMS-D---------D--QAQGVLSGGM--------GNCVE---------------GGSTFFQPVNEILDRWNLELV
>XX00000218 WP_011873078 29 165 183 MKRRLA--------A--RV---------------TG----
--------SVIFA-----------AG-------TIAAL---------------------T-V--------P-------------------A-TAN--------
PTAPAPVAAEAATLMEAMQRDLGLTAEQAAERLADE---------------------------------------------------------------------
---------------------------------------------------------------------------------------------AAATRADQSLRGTLGS-
AFGGSHYDAAL-GKLVVGVTDAGLLD---E--VRAAGA-D--GEL-VQH--NVQQLDG-VADGL-DAQSAR---------
---------------------------------------------------------------------------AP------QS-VTGWYVDSSSNSVVLTTAPGT-A----
GQ--A--TD-FVRASG-VD----A-G-A-VRV---------VE---------S-----A-EQPRTY-A-DVIGGDA-YYI----G----S----
GSRCSVGFSVQG--------G-FVTAGHCGNQ-GD--S--TS---------------QPSGTFEGSSFPGNDYGWVRTAS--
G-E-N--PVPL--VNDY-QG-GTVGVAGSSEAAEGASICRSGSTTGWHCGTVEAKNQTV-RY-P--QG-------T-
VEGLTRTNVCAEPGDS-G-GSWLS-G---------D--QAQGVTSGGS--------GDCTS--------------
GGTTYFQPVNEILQAYGLTLL

>XX00000017 WP_051771612 16 166 187 MLAF-------------GL--------------LV-------
-----PSLNS---------------------AA-------------------A-S--------D-----------------LAATS--------SVD---
APRVAAGLAEALGRDLGLTPDQVRTRLAQE----------------------------------------------------------------------
--------------------------------------------------------------------EKARAVEDTARKALGD-AYAGSWYDDRS-
GELVVAVAGVASAD---L------DA-K--VVP-VAH--TLAELDS-AKARI-DRLAGKA---------------------------------
---------------------------------------AP------AA-VNGWYVDTRTNTVVVTVNRTK-L----DP--A-ARA-FVALAK-
SA----H-G-A-VRV---------VE---------E-----D-HSPKPL-A-DVVGGWP-YWI----N----S----AGRCSVGFSVYG--
------G-YVTAGHCGRP-GD--V--TS-D-----G---N-G--AVLGSFAASSFPGNDMAWVRTNA--G-V-T--LWGY--
VEGY-DG-YWYYVRGSAQVPVGSGICRSGSTTGMWCGTIVARGQTV-NY-P--QG-------V-
VYNLTRTTVCAEPGDS-G-GSWLS-S---------N--QAQGVTSGGS--------GNCSS---------------
GGVTFYQEVNPILSTYGLSLI
>XX00000031 WP_052756301 28 166 205 MSVSKS----NLR----SR--------------------
------------------------------------------------------------------------------------
ASLVAAVAALSAGSAMATDGLNPTLKLAMQRDLGLSSAQVAQMVQAD---------------------------------------
------------------------------------------------------------------------------------------
RIATTQEAALRRTLGS-GYAGSWVERNDDGSYRVVAATSGAQK---S--VAVAGV-E--LRH-VRH--SLKQLND-
SMAAL-DRDARRR--------------------------------------------------------------------VPGISKLRSG-
VQSWYVDPTTNSVVVSVAPGA-D----EE--A--ID-FVAVSG-AD----I-S-S-IRV---------EE---------A-----V-
GTPQTT-A-TVQGGIE-YRD----G----R----VGLCSVGFPVTKG-T-I--KG-FATAGHCAKA-GQ--S--VQ-I---------
S-G--VNVGTFTASHFPNTDRAWVTIGA--A-H-T--LLGS--VTNY-TG-
GSVAVKGSTEAAIGAAVCRSGRTTQYKCGTITAKNVTV-NY-G--TG-------T-VSGLTRANNCTGRGDS-G-
GSWIT-A--------AG--QAQGLTSGGNLP-AGQ-NDNCSVP------------TSQRQTYFERINPVLSQYGLALV
>XX00000107 WP_013222213 21 166 180 MKIARL--------L--GA--------------AA-----
-------TAVMA---------------------------------------------TGALAATAVP---------------------------------
ASGQDLLNPDMVPAMQRDLGLTHDQALARLKNE--------------------------------------------------------------
--------------------------------------------------------------------------DVANKVTESLTAALGD-
AFGGATYNAAT-GKAHVTTTNAALAG---K--IREAGA-E--AEV-VKF--SARQLNS-TVDRL-NAADKA-----------
------------------------------------------------------------AP------AA-ITGWGVDDATNRVTLDVLQGQ-R----
AA--A--DA-FLAKSN-VD----K-T-A-VTI---------RE---------T-----A-TKPTTF-A-NIRGGDA-YYI----G----G----
SARCSVGFSTTT--------G-FLTAGHCGAL-TG--GGALT-G-----S---N-G--ASLGSWITYRFPGSDYAAVRTNS--
N-W-T--PVAQ--MN---NG---TAVRGSSNAATGSSVCKAGSTTGWTCGTIGAKNQSV-RY-S--EG-------T-
VNGMTATNVHSAAGDS-G-GGFIA-G---------N--YAQGLLSGGN--------S-------------------
SVTYFFPIAPALSGTGTTLK
>XX00000163 KNC97026 21 166 204 MNAANA----------------------------------------------
------------------------------------------------------------------------------------------------------------
--------------------------FTL---------------TSALLI--CSA-----I-A-------A------------------------------------
PNPHPVHAEIIASIAEAEQVHAGFLFKREQYLSTKATLLSSQSNSTVDGGHWIDRRN-GKLIVAIKSEHRHR---
RDLSTDPDI-E--HRF-VAR--STSDLED-LKSQV-DAA------------------------------------------------------------------
------------HQ------PT-GVTWYVDVPANNIAVSIWDIKRN----NT--N-TDV-FINEVS-GL----G-P-G-INL--------
-IH---------G-----S-PEIKPR-F-GILDGEE-LTS----A----N----FMSCSMGWWVKDGSG-A--DL-LMSAGHCLKS-
GH--N--WF-R-----N-----A--LLIGGNTKNNFGPMDWGIIHVDD--I-A-AP-RPTQ--ITLH-NG-
KTLKISGYGRAPIGATVCKSGTTTKYTCGPVKAYDATA-NY-Q--SA-----GVQ-VKGMTLADVCTRPGDS-G-
GPLFQ-P-DPNRPTTHA--IAQGIVSGGP--------TEG-----------------CANSIYQPIDTALTDSQTILI

>XX00000201 WP_006240664 29 166 183 MNRKNA--------A--RL-------------IA----
--------SVTLA----------A-------GTAVAF-------------------T-L--------P----------------ATAAP--------
AADAVVPATAADPVVQAMQRDLGLTKQEAEQRLRSE------------------------------------------------------------
------------------------------------------------------------------------------------AEAREVHETVSERLGS-
DFAGAHYDAER-GTLVVGVTDAAEFS---E--VREAGA-T--PRL-VEH--TVADLES-AAEKL-DAKESR------------
--------------------------------------------------------------AP------ES-VTGWYVDIEANSVVVTTKPGT-A----GQ--
A--ER-FVSRAG-VD----A-D-A-VDV---------VE---------S-----K-ESPRAL-M-DIIGGNA-YYM----G----S----
GGRCSVGFSVQG--------G-FVTAGHCGTT-GT--T--TS---------------SPTGRFAGSSFPGNDYAFVRTGS--G-
D-T--LRPW--VNMY-NG-SARVVSGSSEAPVGSSICRSGSTTGWHCGTVEAKNQTV-RY-P--QG-------T-
VYGLTRTNVCAEPGDS-G-GSFIS-G---------N--QAQGMTSGGS--------GNCTW--------------
GGTTYFQPVNEVLNAYGLRLI
>XX00000236 WP_015101633 25 166 177 MRKI----------------VNGRRLAAR---------
--------------SAITLVAGVLTAGITA-APA---------AGIEGYATEVAQS------------A-------I-----A-------------------
------------DMSTREGISPAAAAEILRTQ------------------------------------------------------------------------
-----------------------------------------------------------------DASARTLGQLSDRLGD-RVAGGYLDA-N-
GKPVVNVLDAATAE---Q--VRASGA-E--AKL-VRH--SSQALAS-AQSTL-ESV-------------------------------------
--------------------------------------PA------VT-HTSVGLDPKTNQVVLSVSSAA-K----GG--D-LAA-LLRTAR-
QL----G-D-R-VRV---------ER---------V-----T-GEM--T-L-AILNGEA-IT-----G----G----GTRCSVGFNTNKG-G-
Q--NY-IVDAGHCTRA-VS--Q--WN------------------VGPSQGASFPTNDYGLIRNTT--G-S----APGA--VSLY-
NG-SSQKISSHAPATVGQRIQKSGSTTRLTSGSVQRLNVTV-NY-S--EG-------A-VRELIQTNALANPGDS-G-
GCLFA-G---------S--VGLGITSGKG---------------------------SGTSYYQPVAEALTAYGVTLN
>XX00000244 WP_024876375 22 166 176 MRKLMR---------------------A-----T------
------VA--A-----GACALAFGAVA-PVAGAT----SGPEIEGYSADVQ------SE------A-------V-----S-----------------
--------------SLGAERGITDTAASRILDTQ----------------------------------------------------------------------
-------------------------------------------------------------DDSVRALDAALAALGD-RAAGGYLDA-S-
GAPVVNVVSEVAASQ-AR--TAQAGV-V--TRV-VEH--STADLEA-ARDAL-P-------------------------------------
-----------------------------------------S------VA-DTSVATDPTSNQLVVTVGSDA-P----AA--E-VAD-LRAAAD-
RL----G-D-A-VRV---------EQ---------V-----A-GGF--D-A-AIYGGEA-IT-----G----G----GSRCSAGFNANSG-G-
Q--NY-IVDAGHCTGA-VS--Q--WN------------------VGPSVGASFPGNDYGLIRNDT--G-S----APGA--VSLY-
NG-SAQAINSAANAYVGQSICKSGSTTGLTCGTVQATNVTV-NY-P--QG-------A-VYQLVQTSASVNSGDS-
G-GALFA-G---------S--TGLGITSGMG---------------------------GGSSFFQPLTEALSAYGVTLN
>XX00000402 ABI54136 33 166 198 MYVSNRSRRIARA----SV---------------------------
----------------------------------------------------------------------------
FCVAAALAAMSSGAALASEQVGPELKFAMQRDLGIFPAQLPQYLRTE------------------------------------------
-----------------------------------------------------------------------------------------
KLARTQAAAIEREFGA-QFAGSWIERNEDGSFKVVAATSGARK---S--SALGGV-E--VRN-VRY--SLKQLQA-
SMDQL-DAGANAR----------------------------------------------------------------------VKGISKPLDG-
VQSWYVDARTNSVVVKVDEGA-T----DA--G--VD-FVALSG-AD----S-A-Q-VRI---------EH---------S-----P-
GKLQTT-A-NIVGGIE-YSI----N----N----ASLCSVGFSVTRG-A-T--KG-FVTAGHCGTV-NS--V--AR-I---------
G-G--AQVGTFAARVFPGNDRAWVSLTS--A-Q-T--LLPR--VV-N-GG-
SYVTVRGSAEAAVGAAVCRSGRTTGYQCGTITAKNVTA-NY-A--EG-------A-VRGLTQGNACMGRGDS-G-
GSWIT-S--------AG--QAQGVMSGGNVQSN---GNNCGIP------------ASQRSSLFERLTPILSQYGLTLV
>XX00000409 P00778 33 166 198 MYVSNRSRRVAR--V--SV--------------------------
----------------------------------------------------------------------------

SCLVAALAAMSCGAALAADQVDPQLKFAMQRDLGIFPTQLPQYLQTE------------------------------------------
-------------------------------------------------------------------------------------------------
KLARTQAAAIEREFGA-QFAGSWIERNEDGSFKLVAATSGARK------SSTLGGVE--VRN-VRY--SLKQLQS-
AMEQL-DAGANAR----------------------------------------------------------------------VKGVSKPLDG-
VQSWYVDPRSNAVVVKVDDGA-T----EAGVD-----FVALSG-AD----S-A-Q-VRI---------ES---------S-----P-
GKLQTT-A-NIVGGIE-YSI----N----N----ASLCSVGFSVTRG-A-T--KG-FVTAGHCGTV-NA--T--AR-I---------
G-G--AVVGTFAARVFPGNDRAWVSLTS--A-Q-T--LLPR--VA-N--
GSSFVTVRGSTEAAVGAAVCRSGRTTGYQCGTITAKNVTA-NY-A--EG-------A-VRGLTQGNACMGRGDS-
G-GSWIT-S--------AG--QAQGVMSGGNVQSN---GNNCGIP-----------ASQRSSLFERLQPILSQYGLSLV
>XX00000018 AHE78423 32 167 207 MSLSQSARRACRR----SL---------------------------
---------------------------------------------------------------------------
FLALASAASLVSGAAFAADSIDPRLQLAMQRDLGISAKQLPQYLKTQ---------------------------------------------
-----------------------------------------------------------------------------------------SLSLRQGASAKRALGS-
NFAGSWIERKADGSFGFVVATSGTGK---A--ARSGGV-E--VRS-VRH--SLGQLEN-AFAAL-ESQVRSR---------
------------------------------------------------------------VAGVSKPLGG-VHSWRIDPVTNSVVVSLAPGA-T---
-ED--G--VD-FVALSG-AD----A-G-A-VRF---------ET---------D-----E-GTPQLL-A-NVIGGDQ-YLT----E----G---
-YYPCSVGFSVTKG-E-T--KG-FVTAGHCGQV-GR--V--VL-TG-D-VE---A-L--
VPLGTFTAVNHPGTDMAWVTVNP--E-H-T--LLGQ--VKDY-AD-
GLLSVKGSVEAPVGAALCRSGRTTGYKCGTIRAKNVSV-RV-G--FD-------R-YTGMTETNVCTGRGDS-G-
GAWLT-A--------DG--QAQGVTSTGLLP-FGA-EDNCAVA-----------ESTRKTWFQPLNPILSKYGLTLL
>XX00000035 WP_017616658 28 167 191 MRTSPI--------F--SA-------------LG------
------ATALT----------F---------------------------------G--------L-------I---ASPAPASADSP--------
TDAPQGAPDRADTQSEALQRDLGLSSAEAKKLLANE-------------------------------------------------------------
-------------------------------------------------------------------------------TKARKLERTARKAAGD-
AFGGAVYDQSS-GKLTVSLTDASATE---K--VEAMGM-R--TRV-VEH--GEEQLDE-VVDEL-NGAEDT----------
------------------------------------------------------------AP------AE-VTGWYADLEDDSVVVTALPGE-T----
KA--A--KD-LVADAG-VD----A-D-T-VRI---------EE---------A-----S-ETPELY-A-DIIGGLA-YYI----G----N----
RSRCSVGFTATNSSG-Q--HG-FVTAGHCGNV-GD--R--VT-L----------G--
NGTGVFQYSRFPGDDMAFVRATS--N-L-S--PTNL--VSMY-NG-
SYREVRGSNEANVGASVCRSGSTTGWHCGTIEAKNQTV-RY-P--QG-------T-VYGMTRTTVCAEPGDS-G-
GSYIS-G---------Y--QAQGVTSGGS--------GNCRG--------------GATTFFFPVNDILQQFNLDLY
>XX00000040 WP_026417552 29 167 186 MNRRIA--------A--RV-------------TG----
--------AVVLA----------AG-------TTVGL-------------------T-L--------P-------------------ATA--------
DQEAPAPVSADQEVLLAMQRDLGLDAAEAELRLAQE-------------------------------------------------------------
-------------------------------------------------------------------------------AAAATAEETLTRTLDG-
VFAGAVFDAER-GALVVAVTDEAEAS---A--VRAAGA-V--PRV-VEH--TLVELES-VKSAI-DQTASSQG---------
------------------------------------------------------------AP------AG-VSSWAVDVETNSVVVTVDPEL-R----
DA--E-VDA-FLDRAA-SD----G-T-V-RVV---------EE---------A-----N-STPVPF-A-DIVGGDA-YY-----P----G----
ASRCSVGFSVDG--------G-FVTAGHCGTV-GT--S--TS-G-----V---N-Q--
QSQGTVAGSIFPGSDMGWVETNS--S-W-T--PTAA--VNDY-DG-
GTVAVSGSSEAAIGASVCRSGSTTGWHCGTIEAKGVTV-NY-Q--EG-------A-VHEMTQTSACAEPGDS-G-
GSWLS-G---------S--QAQGVTSGGS--------GNCTL--------------GGTTFFQPLNPILSEFGLSLT
>XX00000124 WP_044849715 21 167 176 MKITKL--------L--GV-------------AA-----
-------TAALA----------T---------------------------------GALVTGTPAA--------------------------------

ASSDTLLNAEMVPALQRDLGLTAGQALARIQSE--------------------------------------------------------------------
--------------------------------------------------------------------------TAAGTLEQSLASALAG-SFGGASYNEAT-
GKLRVGVTDAAKLG---Q--VVASGA-E--AEL-VRF--SAAQLDS-TVDKL-NAGERA-------------------------------
------------------------------------------AD------GA-ITGWGVDTKTNRVTLNVLPGK-S----GT--V--DS-FLEKTG-
VD----K-S-A-VAV---------VE---------T-----A-AVPTLK-Y-DVRGGDA-YYI----N----N----SSRCSIGFSVNG------
--G-FLTAGHCGPG-TV------T-G-----S---N-R--VAMGSFARASFPGNDYGHVRVNS--N-W-V--PRGI--IN---
NG---TRVSGSSEASTGASICKSGSTTGWTCGTVGAKNQTV-RY-A--EG-------T-VYGMTATNVRSQAGDS-
G-GSFIA-G---------N--QAQGMLSGGN--------S--------------------TVTYFFPVRPALSATGTSLV
>XX00000150 WP_025354149 29 167 192 MNRTRA--------V--RV--------------VG---
---------AALLA----------L----------------------------------G-VASAISVP---------------------
ALGASVPASPTPSSDDFSLSPDVVTSLQRDLGLDPAQAQQLFAQQ-----------------------------------------------
------------------------------------------------------------------------------------------------DHAARLADSLRGKLGG-
AFADSWFDQVS-RRAVVDVTDARQAA---Q--VTAAGA-K--ARV-VRH--TEAQLNA-VKDRL-DALA-----------
------------------------------------------------------------------AP------DS-VTGWYVDLPGNSVVVETTGTD-Q-----
R--V--SA-FLDQAR-SL----G-D-S-VRV---------VT---------G-----S-AKPELF-Y-NVRGGDA-WY-----G----S----
NFRCSVGFSATDSSG-G--KH-FVTAGHCTKS-GG--S--AS-G-----Y---N-R--
AALGTINGSHFTGGDYGKVDVTS--SQW-T--LQGL--VT-N-GS-
STVAVKGSTEAAVGAAVCRSGSTTGWHCGTIQAKNQTV-RY-A--EG-------T-VTGLTKTNVCAEPGDS-G-
GAWIS-G---------S--QAQGVTSGGS--------GNCSS---------------GGTTYFSPVGPALSTWGLRLT
>XX00000189 WP_014909500 29 167 188 MRPSPV--------V--SA--------------IG-----
-------TAALA----------F----------------------------------G---------------LALTTSPAALAATGPL--------
PQSPTPDEAEATTMVEALQRDLGLSPSQADELLEAQ--------------------------------------------------------------
--------------------------------------------------------------------------------AESFEIDEAATAAAAD-
SYGGSIFDTDS-LTLTVLVTDASAVE---A--VEAAGA-E--AKV-VSH--GMEGLEE-IVADL-NAADA---------------
-------------------------------------------------------------Q------PG-VVGWYPDIHSDTVVLEVLEGS-G----AD--
V--DS-LLADAG-VD----T-A-D-VKV---------ES---------T-----T-EQPELY-A-DIIGGLA-YT----------M----
GGRCSVGFAATNASG-Q--PG-FVTAGHCGTV-GT--P--VS-I-----------G--
NGQGVFERSVFPGNDSAFVRGTS--N-F-T--LTNL--VSRY-NG-
GYATVSGSSQAAIGSQICRSGSTTGWHCGTVQARGQTV-SY-P--QG-------T-VQNLTRTNVCAEPGDS-G-
GSFIS-G---------S--QAQGVTSGGS--------GNCSF---------------GGTTYYQEVNPMLSSWGLTLR
>XX00000235 WP_005461306 26 167 176 MRKLMK----------------------STVAAGA--
----------CA--L----------VFGAVA-PAMASETDTASTAGIEGYSANVEAE------------A-------V-----S---------------
-----------------TLVAERGLDRAEAVELLRKQ--------------------------------------------------------------------------
--------------------------------------------------------------------DAAVRSLDAALAQLGD-RAAGGYLDD-A-
GNPVVNVVSKTAAA---A--VEGTGV-T--AKL-VSH--TTAELEA-AKAAL-EAL--------------------------------------
---------------------------------------PA------VA-DTTIATDPTTNQVVLTIGTEA-D----DT--E-AAD-LVAEAR-KL-
---G-D-K-VRV---------EY---------V-----A-GGF--D-V-AIYGGEA-IT-----G----G----GSRCSAGFNVNSG-G-Q--
NY-IVDAGHCTGA-VS--Q--WN------------------VGPSVDASFPGNDYGLIRNDT--G-S----APGA--VSLY-DG-
SAQPITSASNAYVGQSICKSGSTTGLTCGTVQATNVTV-NY-A--EG-------A-VHQLVQTSASVNSGDS-G-
GALFA-G---------S--VGLGITSGMG----------------------------GGSSFFQPLTEALSAYGVQLN
>XX00000403 Q6K1C5 29 167 188 MRPSPV--------V--SA--------------IG-----------
TGALA----------F----------------------------------G---------------LALAAAPGALAASGPL--------
PQSPSPDSDVATTMAEALERDLNLTSAEAQELLTAQ---------------------------------------------------------------
-------------------------------------------------------------------------------EAAFEADEAAAQAAGD-

AYGGSVFDTET-LDLTVMVTDAAAVQ---A--VEATGA-K--ADV-VSH--GLDGLEA-VIDDL-NEAAH------------
------------------------------------------------------------Q------PG-VVGWYPEITSDTVVLEVLEGA-E----AD--
A--EA-LIADAG-VD----A-S-T-VTI---------ET---------T-----T-QTPELY-A-DIIGGLA-YT----------M----
GGRCSVGFAATNASG-Q--PG-FVTAGHCGSV-GT--Q--VS-I-----------G--
NGRGVFERSVFPGNDAAFVRGTS--N-F-T--LTNL--VSRY-NG-
GYATVSGSSTAPIGSQVCRSGSTTGWYCGTIQARNQTV-SY-P--QG-------T-VHSLTRTSVCAEPGDS-A-
GSFIS-G---------T--QAQGVTSGGS--------GNCRT---------------GGTTFYQEVNPMLNSWNLRLR
>XX00000036 AHE78426 30 168 199 MQSTGFSSRRSGA----HV--------------------------
--------------------------------------------------------------------------
CALSLLCMVAFDAAAATDLQVSPTLHQAAQHDLGLSSAQVDQLLQVQ-----------------------------------------
----------------------------------------------------------------------------------------------
RDAPAQEALARRLGA-HFGGAWVERASDGRYRFVVGSSDPRG---P--LKLDGV-Q--LRQ-VRY--SLSELEA-
AKARL-DRSVRAR---------------------------------------------------------------------VSGISRPIDG-
VHSWYVDPGSNSVLVSVAPDA-M----ER--A--ID-LAAVSG-AD----S-G-L-LRF---------QQ---------T-----P-
GVPQPT-S-SVYAGRS-YEK----N----G----ALSCSVGFAVMQG-A-T--KG-FVTAGHCGTA-GE--T--IG-L--------
-D-G--APAGYFAASEFPGTDRAWVALNA--N-H-E--LFPL--ITNY-AG-
GFFAVRGSVEAGYSAVVCRSGWKTGYQCGLITAQNVTV-NS-T--RG-------V-VFGLTQSNACAGFGDS-G-
GGWID-V--------HE--QAQGVLSLASIA-PGA-QNNCGTG------------I-FQVSWFQPVNPILQQYGLTLV
>XX00000175 XP_006667902 19 168 187 MELTKF-------------------------------------
----------------------------------------------------------------------------
LAFLAVVLPAAYGVPTPSGELHPKILEAMKRDLGLDAEQATARVARD--------------------------------------------
-----------------------------------------------------------------------------------------------
LQAAEVIEQVRTSAGE-SFAGAWIDGE--AGLQIGITDEALAG---Q--VTAAGA-T--PVV-MAN--SLSKLEE-
AKANL-DK-FFTDETKKRS-----------------------------------------------------------------AS-----AEG-
IASFFVDEASNKLVLEVLADS-R----AH--A--EE-LAAQVG-LT----A-A-E-FEI---------RT---------V-----D-
EMPTIM-A-TVQGGDA-YYI----N----N----QFRCSVGFSVTT--------G-FVTAGHCGTV-GS--S--AT-S-----S---
G-G--QPLGTFAGSNFPGADMAFVRTVS--G-T-V--LRGA--INGY-GQ-
GNLPVSGSTEAAVGASVCRSGSTTQVHCGTIRAKGATV-NY-A--EG-------S-VTGVTQTNVCAQQGDS-G-
GSFYS-G---------A--QAQGVTSGGN--------GDCTR---------------GGTTYFQPVNRILQTYGLTLV
>XX00000207 WP_034384145 25 168 213 MAMSRK-----------------------------------
-------------------------------------------------------------------------
LLPLVLAATTLWVPQDANADPVRPPGPDGLAEVIKAMSRDLHIDEATARRRLVQE-----------------------------
------------------------------------------------------------------------------------------------------
EAAATAEAALVPALGT-AYGGSWFDART-GKLVVNVTDPAKAP---V--VSAQGA-T--PKL-LKNGYSMARLTA-
VKAKL-DHLARTSP------------------------------------------------------------------SS------
VRGVRSWSIDPVANRVVVTRLAGA-E----GG--A-----LAEAAL-AQ----G-D-A-VRF---------ET---------T-----T-
QPAVTG-V-YLDGGEG-IN----------S----GKDCSLGFNARLTVS-D--PV-VLSAGHCANL-PT--V--T--G-----Y---
G-N--TRSDQPTRTSYPGLDWGVIGSLD--L-Q-RWAQGPYIALHTP-GD-
HYLNVTGTQQRPEGSSICKSGFKTRITCGVIVRRGEPI-WLWD--AG------RT-VFNLTRHTACTDPGDS-G-
GPVYT-A--------DG--HAQGVVSSSG--------VGCSQ---------------PGDGWYVEINHIQGASGARVL
>XX00000156 WP_030731666 27 169 189 MRRTRS--------F-----
TLGAAALMVLTAGVG------------GAAVA----------A--------------------------------D-SSTPAPGG---------------
---------------------EPVAEPALIEALQRDLGLTPEQAVDRLATE-----------------------------------------------------
---------------------------------------------------------------------------------AEAMATDEKAQEAAGD-

AYAGSWIDAAT-GTLVVAVTSAAAES---A--VESTGA-A--TER-VTH--SQEALDA-ALAAL-DELE----------------
---------------------------------------------------------AP------AG-VSSWYADVQDNGIGVDVVAGS-E----ND-
-Q-VAE-FLAEAEAV-----Q-SIP-LTV---------RT---------V-----E-EAYTTSAA-GVVGGDP-YY-----T----G----
NVRCSVGFSVHG--------G-FITAGHCGSG-SA--A--VY-G-----W---D-N--
SLMGNFAGRSWPGGDHAWVRTGH--G-W-W--TVPV--VLGW-
GTVSDALVRGSNEAPTGASVCRSGSTTGWRCGTIGAKNVTV-NY-G--NG-------
NVVYGMTNSTACAEGGDS-G-GSFIS-G---------D--QAQGVTSGAA--------GNCSS--------------
GGSSVFYPIRPILSAYGLTLH
>XX00000157 WP_026424317 20 169 188 MKRTLV------AAA--VL-------------TA---
---------SAGVV----------V--------------------------------A-------------------------APAFA--------
GQDAAKTEGVSNVVLDTMQRDLGLSHDAALARIASE-----------------------------------------------------------
----------------------------------------------------------------------------NRATALESTLRASLGD-
TFGGAYYDAPS-ATLVVGVTDQAKIA---A--AQSAGA-T--AKF-VSH--SARQLDT-TADTL-NAKEKS------------
---------------------------------------------------------AP------AG-ITGWYVDVQHNTVTVTTARGT-L----AA-
-A--HQ-FLTTTG-VD----L-S-T-TQV---------VE---------S-----T-ESPRVL-Y-DVRGGDA-YY-----I---GS----
GARCSVGFSVQG--------G-FVSAGHCGNR-GD--S--TS-G-----S---N-R--
VAQGTFQASTFPGRDYSWVKTNA--N-W-T--PRGV--VNRY-SG-
STVAVQGSTESAVGAAICRSGSTTGWHCGTVQAKNQTV-RY-S--EG-------A-VNGLTRTNVCAEPGDS-G-
GSWLS-G---------T--QAQGVTSGGS--------GNCTS---------------GGTTYFQPVNPILSTYSLRLV
>XX00000180 WP_012243860 28 169 224 MKMAKT----------------------------T----L-
R-A---CLSSL----------A------------A-------------------T-G--------L-------L-----V----
SAVAGAATAAPATTPANLHPESATEAAVKYLESTGLSSNDAADRVVKQ-----------------------------------------
---------------------------------------------------------------------------------------------
PALSLAAERLDGVLGA-TGSGAFIDQAT-GKLTVGITNPTAAL---T--AFGNGA-E--KVVQMSR--SAAQSEQ-
LRTAL-SPLL------------------------------------------------------------------------SE------VA-
GSSWGVNPSSNTVDVELPANT-P----AA--V--SQ----ALT-QY----G-S-A-VKL---------TE---------T-----A-
ATTAGT-T-NAYGGQE-YKP---------Y----WQVCSLGFAAVDG-G-A--SF-DVTAGHCTTK-IG--A--KEAK-------
--G-T--TILGTHEKSNFPGTDYGLIRTNT--AAI-S--LQAQ--VDRQ-NG-
TYVNVTGSTESAVGGTSCKSGRTTQWTCGTIKAKNVSV-NY-S--RE-D-GGTDR-VNGLVQYNACVEGGDS-
G-GAILS-G---------T--QAQGLTSGGQ--------GYSQGP--GKPAVCGEKVGKPNVAYYQPVNPALSAYGMTLV
>XX00000001 WP_020142554 26 170 204 MIRTSL--------T--TL-------------AA------
------TAAIA----------T------------A-------------------I-T--------V-------M-----
PAQASTLASDPTPAPTPAASTDSGQSVAEMSARWLAKDRAISLATARQRVAAQ--------------------------------
------------------------------------------------------------------------------------------------------
DGQTRTAASLERALGA-RAAGSYIDATS-GALVVNVVDTASVA---R--VLSAGA-V--AKV-VDR--STSELSA-
TERAA-RARA----------------------------------------------------------------------------G------SA-
VVSSYTDPVTNGVVLTVPSAR-V----SE--V--RS----EVV-GL------D-G-VTV---------AG---------T-----D-
ARTTTQ-A-NVYGGQQ-IE-----F----S----GYVCSLGFNATRG-G-A--PV-FVTAGHCGEG-YQ--T--FS-K---------
G-G--TTLGSTQAYSFPGNDYAYSTLTS--S-W-T--GVGA--VDLY-DG-
VARRVSGYSNAPVGTAICKSGRTTGWTCGSVQAKNVTV-NY-S--NA-D-GSTST-VSGLTKSNTCTEGGDS-G-
GSWMA-S---------T--SAQGVTSGGA--------GYGANS--V----CGQKVGQPNIAYFQPVDEIVSAYGLTLK
>XX00000092 WP_026419506 28 170 189 MRPVR---------------------------------------
------------------LTGLVA-TAAVVT------ATLTGVGAASAGE----RPADGPD-------------------------------------
RQDYSPGLLTSMADEFGITPLEAETRLGQE--------------------------------------------------------------------

-----------------------------------------------------------------FAAMDAVDPAEEAAGS-AYAGAWFDAER-
GALVVGVTDAGAAD---A--VRAAAEVD--VEV-VEH--SAAVLDG-VVAEL-NAVAEEEA-------------------------
---------------------------------------------PP------AD-VTSWYVDPAANEVVVTVLAGA-D----DS--T-TAA-
FLAEAR-A-----L-G-P-VTV---------TE---------V-----S-ERPETFAA-GTVGGDP-YYI----N----S----
NTRCSIGFSVHG--------G-FVTAGHCGGA-GS--T--VH-G-----W---D-W--
SLMGNVAAASFPGNDHGWVRIGH--G-W-W--TEPV--VLGW-
GTVHDVLVRGSSVAPVGSSICRSGSTTGWRCGTVEALNVTV-NY-Q--QG-------P-VYGTTQTSACAQGGDS-
G-GSFIT-G---------D--QAQGVTSGGS--------GNCAT---------------GGRTFFAPVNPILSAYGLSLV
>XX00000106 WP_046471059 28 170 187 MRKSSI--------T--RA--------------LG------
------GGALA----------L---------------------------------G--------L-------VAAGTVAAAAEDQEP--------
VADQGSGQDQAQDLLAAMERDLGLTTAEAEQRMADE----------------------------------------------------------
-------------------------------------------------------------------------------AEARSIDNELRTELGE-
AWGGSYFDGDS-GELTVAVTDESATD---E--VAAAGA-N--PEI-VTY--GTDELEA-IVDEF-NAQGSQ----------
-----------------------------------------------------------LG------NG-ITGWYADSSADAVVVEVTSGN-E----
DA--V--DD-LVSETG-VE----A-G-A-VQA---------AE---------T-----D-EAPELY-A-DIVGGDP-YMI----G----G---
-TGRCSIGFAVEG--------G-FATAGHCGDV-GT--E--VS-S-----E---D-G--SGTGTVGGSEFPGTDMGFVEADD-
-N-W-T--PTST--VNDY-EG-GTLEVAGSEQAPEGASICRSGSTTGFHCGEIEAHDQTV-EY-P--QG-------Q-
VQGLTQTNVCAEPGDS-G-GSWLA-D---------D--QAQGVTSGGS--------GNCTI---------------
GGTTFYQPIEPILDEFNLTLV
>XX00000212 WP_047135691 28 170 195 MHTLHA----AVRRTLALA-----------------
-----------------------------------------------------------------------------------
LLSAAAAGGSALAAQPGLPSLDPGMVAALERDLGMTQDQYRVTVDAE----------------------------------------
---------------------------------------------------------------------------------------------
RRVPTVETAARQLYGD-SYAGTWIEHDADGQPRLVVATAARDK---RL-PAVDGA-D--VRQ-VRY--SLYELDQ-
AVHAL-DQSA------------------------------------------------------------------TSRIMRRLDG-
VQSWGVDLPNNRVVVTLSPGA-G----RA--EAVLD-FLARSD-VD----T-G-A-LHF---------EV---------S-----E-
QVPSTL-I-NIYGGIQ------------------YNTCSIGFPVTRG-S-T--KG-FVTAGHCGSA-GT--A--VR-I---------G-G--
TTVGSFQASNFPGSDAAWATVRQ--Q-D-T--LFGR--VWQY-SG-
STTPVVGSAEAAVGAAICRSGYRTGWRCGTITRTNVSV-TY-A--QG-------T-VNGLRESNACAGQGDS-G-
GSWIS-S--------AG--HAQGVTSGGALG--TD-GTNCGIA------------QSQRRSWYQRLTPMLSSYGVTLV
>XX00000030 WP_026066477 24 171 187 MRNRSL--------A--RI--------------AG-----
-------ALVAA---AGTATALA---------------------------------L--------P-------A---------GADTP--------
SPDGADATVASPEMLSAMQRDLGLTEQEALTRVAVE------------------------------------------------------------
-------------------------------------------------------------------------------ATAVETEDELRASLGP-
AFGGAHFDGDT-NTLVVGVTSAAKAD---E--VRAAGA-T--PEV-VAF--SADTLDG-VVSTL-NETSE------------
-----------------------------------------------------------VP------DG-VTGWYVDTADNTVVVTTALGS-G----
EA--A--AD-FVAESG-VN----A-D-A-VTV---------VE---------S-----T-EQPRTL-Y-DIIGGDA-YY-----F----G----
GSRCSVGFSVSV--------G-YVTAGHCGGV-GT--A--TQ-G-----Y---N-R--
VSSGQVAGSVFPGSDMGYVRTNA--N-W-T--PRPL--VNRY-
SGGATVTVSGSNEAAVGASICRSGSTTGWRCGTVQAKNQTV-FY-A--QG-------A-VSGLTRTNACAEGGDS-
G-GSWLS-G---------S--QAQGVTSGGS--------GNCTW---------------GGTTYFQPLNPILSRWGLSLT
>XX00000082 WP_043576116 23 171 181 MKLGI---------A--CR--------------LV------
------GTLLL----------A---------------------------------T--------A-------T---AATAAPASSAS--------
EPGTPEKPDGTASLLEAMQRDLGLSARESRLRLYRE-------------------------------------------------------------

------------------------------------------------------------------------------TIAEQVRHTLRRTLGN-
SYGGAHYDAAS-GRLVVGVTDRSRLA---D--ARAAGA-H--AQL-VEY--SSRRLTG-IAERL-GESLRS-------------
------------------------------------------------------------AP------DG-VTGSYVDTADNSVVLTAERGS-A----AD--
A--RN-YVRSTG-VA----A-D-A-VRV---------VE---------T-----E-RTPRLY-A-DVIGGNP-YYV----G----G----
NTRCSVGFAVQG--------G-FLSAGHCAAT-GD--S--TS----------------NPAGWAEGSSFPGNDFSYIRSSA--R-
G-----RPL--VNDY-AG-GTVNIAGSTEAPVGASVCRSGSTTGWHCGTIRGKRQSV-RY-P--SG-------V-
VEGLTRTDVCAESGDS-G-GSFVS-G---------Q--QAQGITSGGW--------GNCTD--------------
GGTTFFQPVNEALRHYDVDLL
>XX00000186 GAB77127 36 171 214 MPRHTT--------T--TH-------------RH-----------
VRTAM----------S------------L-------------------A-V--------T-------T-----
ALTLTVAPLLAHAGEKSADRAEEISDITKMSADHISRAFGLSAEDALSRAHEQ------------------------------------
----------------------------------------------------------------------------------------------------
EKHAGTARDIQAKLGE-RTSGSFIDQKR-GKLVVNVVDDATAK---Q--ISGDKV-E--AKV-VKT--SRREIET-
NRRKA-EETL------------------------------------------------------------------------G------DL-
AKATSLDTRRNTIVLTVSNKD-G----DA--A--RK----AVE-GL------D-N-VEI---------QT---------S-----D-
DEDTPR-T-SVYGGQK-ITF-TREA----K----RFKCSLGFNAQKD-G-K--DV-FITAGHCGRG-NA--T--FT-K--------
-N-G--KKLGVTQEFAYPGHDMAYATLDP--A-W-E--GKDA--VDKW-NG-
KVVVVKGSTEAPVGTSVCKSGQTTGWTCGTITAKDVTA-KY-R--NK-KTGQVAR-ITGLTQTDACSAKGDS-G-
GAWMA-G---------N--QAQGVHSGGN--------GRKIIR--A----CRHQDGKPNVAYFQPLNPILETYGLTLK
>XX00000116 WP_006503189 27 172 209 MKLPVT-----------TS-------------LA------
------VTTSL----------L------------L-------------------L-A--------P-------M-----
ATVAQAAEDKSEKAGTAASIESDMEALQKITAEYLSKHYGLDNDTAKKRVAAQ---------------------------------
------------------------------------------------------------------------------------------------------
DEFSKEAKKASDELSG-DSAGSYIDQKR-GVLVVQATKDEAKN---K--VKGKDT-E--VKV-VSR--SMSTLEN-
QAKQL-TDKL------------------------------------------------------------------------G------DK-
LVSARIDVQNNRVVATVKKDQ-V----DP--A--KD----SAK-GM------D-G-VEV---------EG---------T-----D-
AVITPQ-K-NIYGGQQ-IE-----F----N----GGRCSIAFNATKD-G-K--DV-FITAGHCMSG-RS--P--FT-H---------
N-N--QPLATPVAGEFPGADMGYAALNQ--G-W-T--GQPG--VDKW-NG-
RGVAVQGSEEAPVGAAVCKSGRTTQWTCGTIQAKNVTV-NF-S--RLKG-GGTDR-MTGLTQASACTEPGDS-
G-GAWIS-G---------T--QAQGITSGGA--------GRPGPQ--GR-NLCMEKFGRPNVAYFQPLKPALQKYGLQLK
>XX00000125 WP_017620102 32 173 187 MRKSPT--------F--RV-------------LG-----
-------GGALA----------L---------------------------------G--------LVAVGTTIATTSGGSEISPVSSD--------
SADSADTATTADDQLTALQRDLDLSATEATRLLDTE-----------------------------------------------------------
------------------------------------------------------------------------------AKARKLDSELRRNLGD-
DFGGSRFDIDS-GDLTVAVTDKAAVD---T--VEKAGA-T--AQV-VTY--GESGLDK-IVDDL-NEEGAS-----------
-------------------------------------------------------------LS------EG-VTGWYPSATDDAVVITVLKGK-E----AA--
A--EK-LIADAG-VE----T-G-A-VTV---------EE---------T-----T-DKPRTY-A-DILGGDAYII-----N----G----
SGRCSIGFAVEG--------G-FATAGHCGAK-GT--P--VA-A-----Q---S-G--
GGTGTVAESHFPGRDMGYVQTDA--G-W-T--PTGF--VNDH-NG-
GTVAVTGSQEAPEGSSVCRSGSTTGWHCGTIGAKNQTV-VY-P--EG-------R-VEGLTQTDVCAEPGDS-G-
GSWIS-D---------D--QAQGVTSGGS--------GNCTA---------------GGVTYFQPLNPILQEWNLTLS
>XX00000161 WP_026128803 28 176 186 MRKSTI-----------VR-------------AA------
------GGGAL----------A---------------------------------L--------GLVGAGVAVLSDGSPQEAATVAA--------
DTGGTDAGGPDAERIEAMQRDLGLSEAEALELVDTE---------------------------------------------------------

------------------------------------------------------------------------------------AQARSTDETLRSALGE-
DFGGSWFDIDS-GTLTVAVTDASAAP---E--VTAAGA-E--AEV-VEN--GRDGLDA-MIDDL-NAEGAG---------
--------------------------------------------------------------------LS------EG-VAGWYPDIRQDSVVITALEGE-E----
AA--A--EE-FARAAG-VP----E-D-A-YTV---------ES---------T-----R-DEPRLY-A-DIVGGDA-YS-----T----G----
GGRCSIGFATES--------G-FVTAGHCGST-GT--Q--VS-S-----Q---D-G--SGTGTVTDAEFPGADMAAVQADA-
-G-W-N--PTPA--VNDY-AG-GTVAVAGSEEAPEGSSVCRSGSTTGWHCGTIQAKNQSV-SY-P--QG-------T-
VNGLTQTDVCAEPGDS-G-GSWVS-D---------D--QAQGVTSGGS--------GDCTS--------------
GGTTFFQPVNPILDTYGVTLL
>XX00000071 AEP18660 26 177 205 MPQAS--------------------------------------------------
--VRASALSILGLLA------------------------------------------------------------
LSAHSAHAQPPAAPAERASNPIAGVGAAALRDGEVSPDRARQQWRAQ-----------------------------------------
------------------------------------------------------------------------------------------------------------
RDAVAGEAAARRALGD-DFAGSWIESEPGGGYRLVVASAGAVA---G--ASVAGV-E--IRP-VAR--
NWRRLQA-SKQRL-DQALQSR-----------------------------------------------------------------------------
RPGLHKPLTG-VYTWHVDPASNRVVVRIARDA-Q----AQ--A--VD-FAAASG-AD----A-D-A-LRF---------EY---
------L-----D-GAPQTT-N-WIFGGLS-YIT----A----Y----GGGCTAGFVAYKA-G-V--PG-VVTAGHCGKA-PD--
K--ID-VDWS-GI---G-R--TPFGQFVHSRFPTADRGWIASSS--S-F-A--VSPQ--IYDW-AG-
NYITVKGSIEAPVGATVCRAGPKTGYRCGQITAKQVTV-NY-P--QG-------A-VYGMTESNACSGFGDS-G-
GPWID-G--------GG--QAQGIQSGNQG--------NSSAG--HN---CDT-PAAQRKSWFDPLNSILGEYGLVLT
>XX00000065 EXG79206 43 178 194 MTTPR------------HRRSPTARRRNI----------
TISSVVAGAVAAALAAFMATAPTAGASDKTGTDAARPAVEAADTGPGAYTAEIRK------------------------------
---------------------------LAVSALAAQLGISEQDAAARLDQL-------------------------------------------------------------
----------------------------------------------------------------------------------------DKRTRTAASLQNSLGD-
SGAGTWISKTT-GELMVGVTDQKAAD---A--VEAAGA-T--PKL-VSR--DLNDLQQ-VKTQL-DAI----------------
----------------------------------------------------------------GR------IP-GTSWAIDPSNNAVTVQVSRKA-A----ADP-
R-ADA-WLDKLA-GY----G-D-A-VNV---------QR---------T-----T-AAF--A-THAFFGGQA-IEA---EG----G----
AGRCSSAFNATSG-Q-N--AF-IITAGHCTAA-ID--T--WT-D-----G-----Q--EVIGQSALTQFPGNDYGVIRVDD-
-A-Q-AL-DPQP--AVIN-QD-QAQPITGTDQVPVGSPVCKTGSTTGTTCGVVLAFDTTV-VY-P--EG-------A-
VQGLIQTDVCSQPGDS-G-GSLFA-G---------D--QGQGIVSGGS-------IGDCET--------------
PFVSFFQPINEVLEDTGLQLI
>XX00000229 WP_005453178 26 179 176 MRKLVR----------
STVAAGACALVFGAVAPAM------------AS--E----ASEASKASETPE-ASTAST------SGVEGYSADVAAE---------
---A-------V-----S---------------------------------ALVAERGVDQEAAKALLRTQ----------------------------------------
-------------------------------------------------------------------------------------------------------------
DAAVRTLDAALSQLGD-RAAGGYLDA-S-GNPVVNVVSQAAAT---S--VESSGV-V--TRL-VSH--TAAELES-
ARAEL-ESL---------------------------------------------------------------------------------PS------VS-
DTTIATDPITNQVVVTVSDAA-D----DA--Q-AAD-LVAAAE-RL----G-D-K-VRV---------EY---------V-----A-
GGF--E-A-AIYGGEA-IT-----G----G----GSRCSAGFNVNSG-G-Q--NY-IVDAGHCTGA-VS--Q--WN-----------
-------VGPSVGASFPGNDYGLIRNDT--G-S----APGA--VSLY-NG-
SAQQINSASNAYVGQSICKSGSTTGLTCGTVRATNVTV-NY-P--QG-------A-VYQLVQTSASVNSGDS-G-
GALFA-G---------S--VGLGITSGMG---------------------------GGSSFFQPLTEALSAYGVRLN
>XX00000097 XP_006673484 19 180 285 MELTKL----------------------------------------
------------------------------------------------------------------------------------
LTFLAAVLPAAYGAPQKMANSMHPNILAAMKRDLGLNTRESMARVERD----------------------------------------

--------------------------------------------------------------------------------------------------------------------------------
QKAADVIENVRRSTGD-AFAGGWLHG---DEVRIGVTDKASAE---H--VTAAGG-T--PIM-MRN--SLSKLEK-
IKAGF-DD-IKQGNATASDDTGVKAREDL------------------------------------------------------------------------GN-----YAD-
IASYFVDVASNKIVFEVVNGN-Q----DS--A--DK-MAKDAG-LA----A-E-E-YEV---------RK---------V-----D-
DYAVPY-A-QLQGGDS-YGI----N----N----QFLCSVGFAVNG--------G-FISAGHCGQV-GS--T--AA-------V---
N-G--QVIGRFARSNFPGGDMSYVQTEQ--G-T-T--VAPT--VNGY-SG-
QSYEIKDSQQLPIGAAVCRSGQTSGVSCGTITEMGKTV-QY-G--PG------QV-VAGLTQTDACSQQGDS-G-
GSFWN-A---------G--HAQGVLSGGP--------KDPGG---------------PCYSIFQPIGPILQNYGLTLT
>XX00000142 WP_033301087 23 181 191 MQKSIR------KSPFLAA--------------LS---
---------TGVLA-------------------------------------------------------------------------LGLTGTTPGTATA--------
DQSTPAPSRLSAGHLQALQRDLGVDPQEAAMLSQQT--------------------------------------------------------------------------------
------------------------------------------------------------------------------------------------------DAARTLESQARAVAGD-
AFGGAVFDIET-GGLTVAVTDRKAAD---A--VDDLGA-T--TEL-VER--SEDELTD-VVTKL-NEAEDA-------------
-----------------------------------------------------------------------------AD------AS-VTGWYLDPRANGVVVTVLDGG-Q----
AA--A--AD-LIAQAG-VD----A-E-A-VTI---------DE---------G-----A-DRPRTA-T-DIHGGNP-YY-----N----PP-
DRTYVCSVGFGVEG--------G-YVTAGHCGER-GK--G--AF-H-----NAG-L-T--
VRYGKVADADFPGTDMGWVKTTG--R-W-T--PTRR--VNDY-NG-
GSVTVAGAEEAPVGAAVCRSGQTTGWHCGTIQAKNQTV-YV-P---H-------T-VHGLTRTTVCAEPGDS-G-
GPYLS-G---------D--QAQGVTSGGA--------GNCTS---------------GGTTFFQPLRPIIEKYDLTLL
>XX00000112 WP_026218628 0 190 176 MAHSGM--------K--RT--------------TV---
---------TRSML----------S-------------------------------------------------------------M-------------------------------------------------------------
------------------------------------------------------------------------------------------------------------------------------------------------------------------------
---------------------------------------------------------------------------AAAAVLVLTTQVGPA-----ASGPTAGG------GCLVD--
RLDELRV-LRHRL-DGLAL----------------------------------------------------------------------------------------------DG----RAGN-
SQYWYIDHQRCTLTIATLRGA-A----DP--A-TSA-FLNTAR-LM----P-A-L-VKIVELDAAIRSRVAKGP-AADP---
---N-SHA--A-S-GFHGGSA-IY-----S----GLTGTVLICTAGFNGYRP-N-----T-TGTAGHCARD-AP--T--WY-D---
--G---T-G--ALVGTVDQSRFPGADWAVIPAAE--G-L-T--LTAD--I-VD-GD-
STTAITGFARPRLRDRVCATGASSGTRCGTIDATDVTV-NY-P--EG-------A-VTGLARSTQSGAAGDS-G-
GPVHI-G---------P--TGVGLISGGP--------P---Q---------------GAPTFIQPLDF----------
>XX00000053 WP_006502134 36 214 208 MRRHTK--------T--AS--------------TQ----
--------ARQAL----------S-----------I-------------------A-A--------V-------A-----
ALTLTGTSALAYADDKKGDALDEISAVEKMSAEYLTKEFGLSEGDALGRVRDQ-----------------------------------------------
-----------------------------------------------------------------------------------------------------------------------------------
KKHATTARELESKLGD-RTSGSYIDQKR-GKLIVNVADEESAK---E--VKGDNV-E--AKV-VKT--SAKEVSE-
NRRKA-EEKL-------------------------------------------------------------------------------------------G------DK-
LTSSSLDTSRNLIVLNVPVKE-S----EG--A--KG----GGV-KG------G-D-NGG---------GV---------K-----G-
GGGAKG-Q-AIYGGQE-IR-----A----N----KFMCSLGFNAKKG-N-Q--DV-FITAGHCTGG-GV--S--FT-R--------
-N-N--KPLGKAQASNYPGADMGYATLES--G-W-S--GKEA--VDKY-NG-
KAVVVKGSEEAPVGAAVCKSGRTTGWTCGTITAKNVTV-NY-R--SK-N-GSTDR-ITGLTQTNVCTSSGDS-G-
GAWIA-G---------E--QAQGVHSGGN--------GNKIVG--V----CKHRTGQPNIAYFQPIKPILEKYGLTLK
>XX00000119 WP_019676482 0 224 279 MKLI-------------------------------------------------
-------------------HVVVLFNAMKTF---------------------Q-
ISRLLLKIIMKKVINRKHFSCYKAFTYLAVATGLALSFSRAAALDSMPALDPKAVTEIQKQLGLSEAETRARLHAE
---------------------------------------------------------------------------------------------------------------------------------------------------------------------

----------------AHANNIHQPLQEFLGD-AYAGAWFDVDR-MKLAVAYTEPRLRN---W--IKVAGA-Y--PVL-
VKH--NRSQLLN-IQQQL-LLEAQADPR---------------------------------------------------------------------WL----
--AG-VSSWYIDDKTNSVVLEALPED-W----DY--I--AG-RLPNIG-RA----N-D-V-LRL---------KR---------S-----
H-KKPQLN-S-DIRGGES-----------------ANGCSVGFSVVG--------G-FVWAGHCGLT-ND--T--VE-N-----A---
S-G--QVIGSIAYTTAPGYDHGWVQTTS--G-W-T--PQPV--VQGY-SD-
GLLPVKGQKTPIQGMSVCRYGASSQPFCGSIHSTANTI-PI-T-ATA-------S-IEGLSRVNIFSFPGDS-G-GPYLT-
A--------AG--DGLGTHSASL--------YGPTD--------------YPESYFEPLSHSLTAMGLTLL
>XX00000148  WP_012244023          0          229       212       MSVA-----------------------------------
DLRFAMHRLQSLAR-SKTIFAVKSQNIQGVSVKTSTNVRKACLSGLAT----------------V-------S-----L----
LAVTFGATAFGAANANAADPADPVQAAVNYLVEQGVNRDAAQHRIDTQ------------------------------------
-----------------------------------------------------------------------------------------------------
AARGELATKLDSILGS-HGAGAYTDFAS-GELTASVTDDAGAA---L--ASAQGI-R--SVR-VAN--SLAQLQS-
VSDQL-AATF-------------------------------------------------------------------------DS------AS-
GARWSISPVTNSVNVTLPTAT-T----ATPAQ--QA----VLD-AS----S-S-V-IKI---------SK---------A-----A-
NAASAG-T-AAYGGQR-YLF---DS----D----RYTCSIGFFSKSA-S-D--IK-IVQAGHCTTT-GG--P--VS-K---------
D-K--VNLGNDEDSSFGGTDYGLIDVNT--ANF-T--GKGA--VDKY-DG-
TYLSVKGSTNVAVGDTICKSGQKTKWKCSKVLSTNASV-SY-Q--DE-----GVT-LKNMIQTNGCTIPGDS-G-
GSNLS-G---------N--YAAGMTSGGN--------LHKTSQ--GS-PYCASQDGDTDETYLQKVGPALQAYGVSLL
>XX00000023  ADI04060        28        285       303       MNKRTV--------------------------------------------
-------------------------------------------------------------------------------------------------------------------------
---------------------VI--AGAA--------------AAALAG-A--A-----V-L-----LP---------------------------------------
-NA---N----ASPDKSAEAKTFSSQAAVKLARGLASDLGG-NAAGWYYQGDS-KQLVMNVLNEDAAN---D--
VRAKGA-V--AKI-VAN--SLTELKA-TTKTL-SDN--------------------------------------------------------------------
-------AS------IP-GTAWSIDPKTNKVRVLADRTV-T----GT--K-WAT-LTKAVD-GL----A-G-K-ATV---------QR-
--------T-----N-GEF--KAG-ALDGGDA-IF-----G----G----GARCSLGFNVTVD-G-A--PA-FLTAGHCGNI-SE--T-
-WA-T-----DQA-G-A--QQVGTTTESQFPVSDFALVTYDD--P-NTQ--ASST--VDLQ-DG-
TTQQITQVAEASVGLAVQRSGSTTGVTDGTVTGLNATV-NY-G--NG-------DIVNGLIQTDVCAEPGDS-G-
GAMFA-E---------D--AAVGLTSGGS--------GDCTQ--------------GGETFFQPVSTALQATGAVIG

Supplement 2. Phylogenetic trees of the αLP family from protease sequences only with rooting by outgroup, midpoint, and minimal ancestral deviation

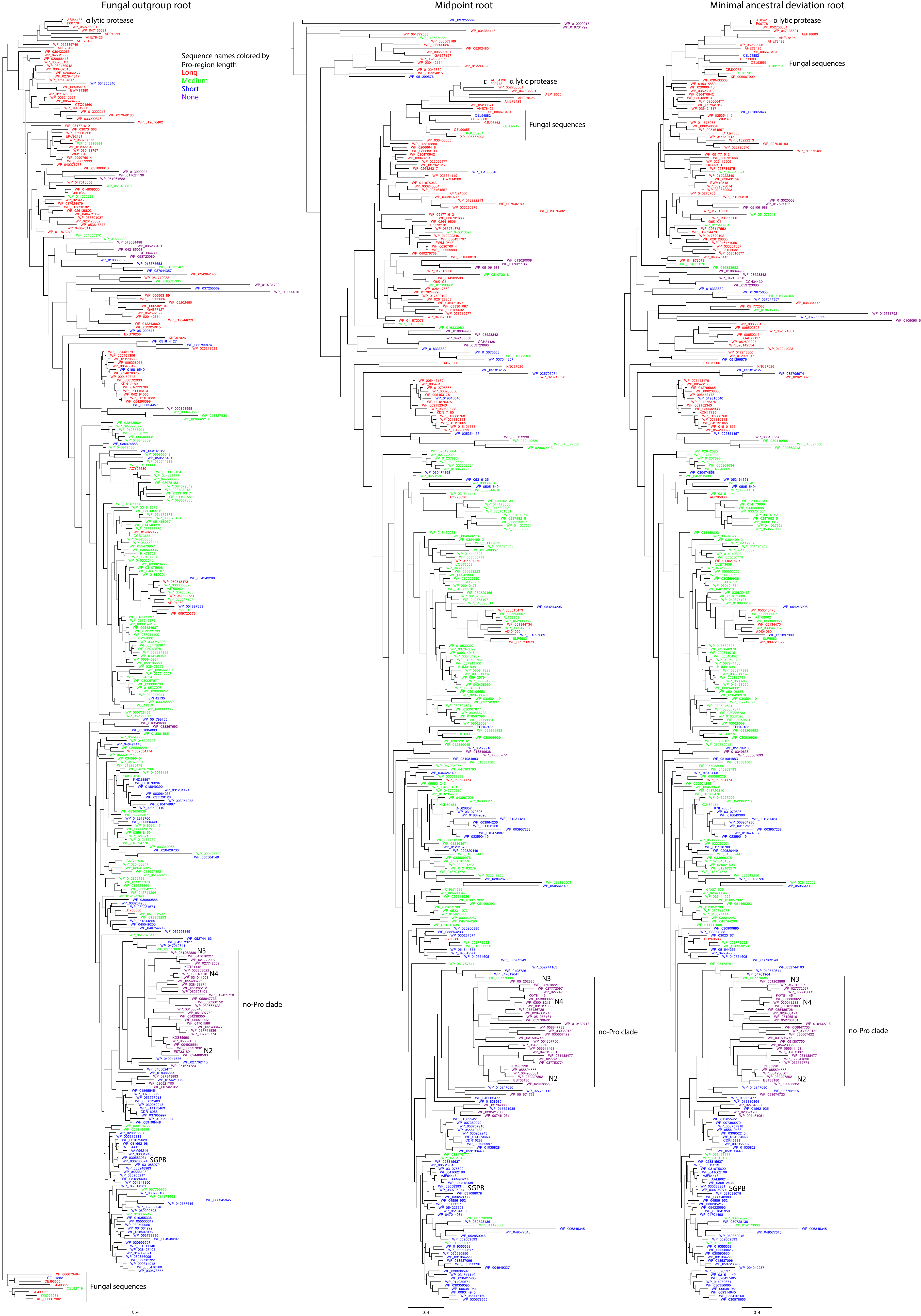

Fungal outgroup root
Midpoint root
Minimal ancestral deviation root
Sequence names colored by
Pro-region length
Long
Medium
Short
None
α lytic protease
Fungal sequences
no-Pro clade
N2
N3
N4
SGPB
0.4

Supplement 3. Phylogenetic trees of the αLP family from full-length pro-protease sequences with rooting by outgroup, midpoint, and minimal ancestral deviation

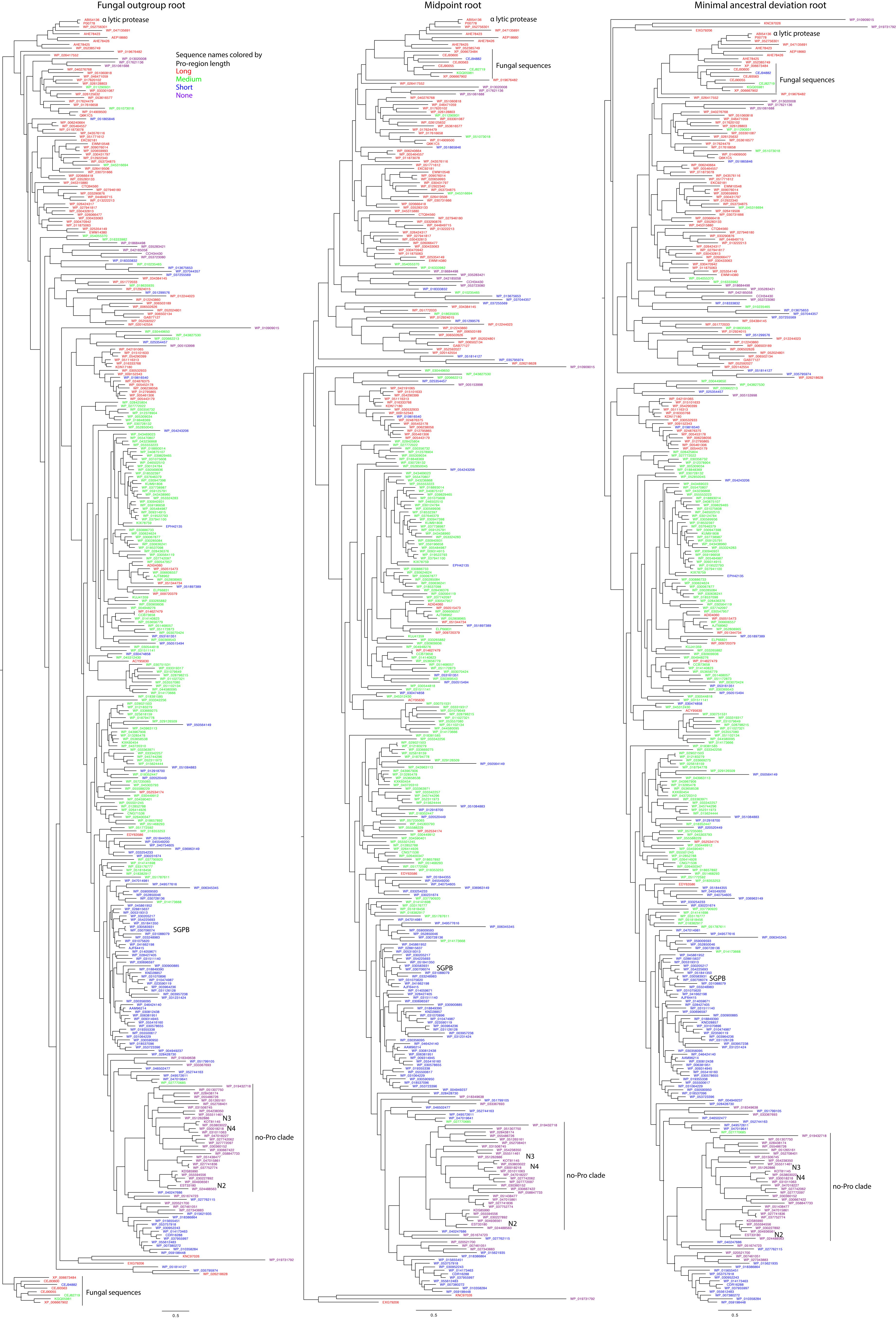

Fungal outgroup root
Midpoint root
Minimal ancestral deviation root
α lytic protease
Sequence names colored by
Pro-region length
Long
Medium
Short
None
Fungal sequences
SGPB
N3
N4
N2
no-Pro clade
0.5

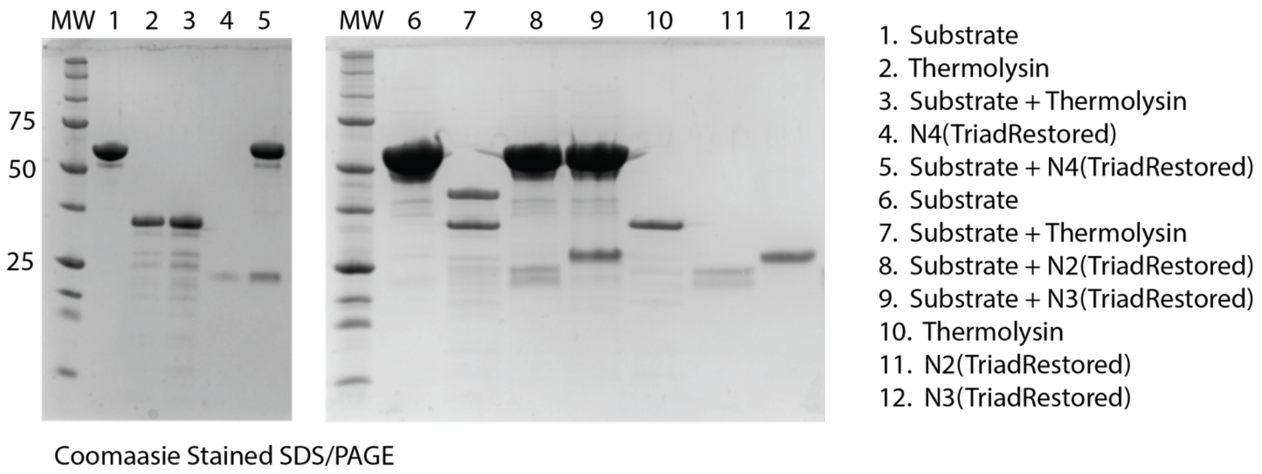


**Supplement 4.** Protease activity assay of engineered No-pro homologs

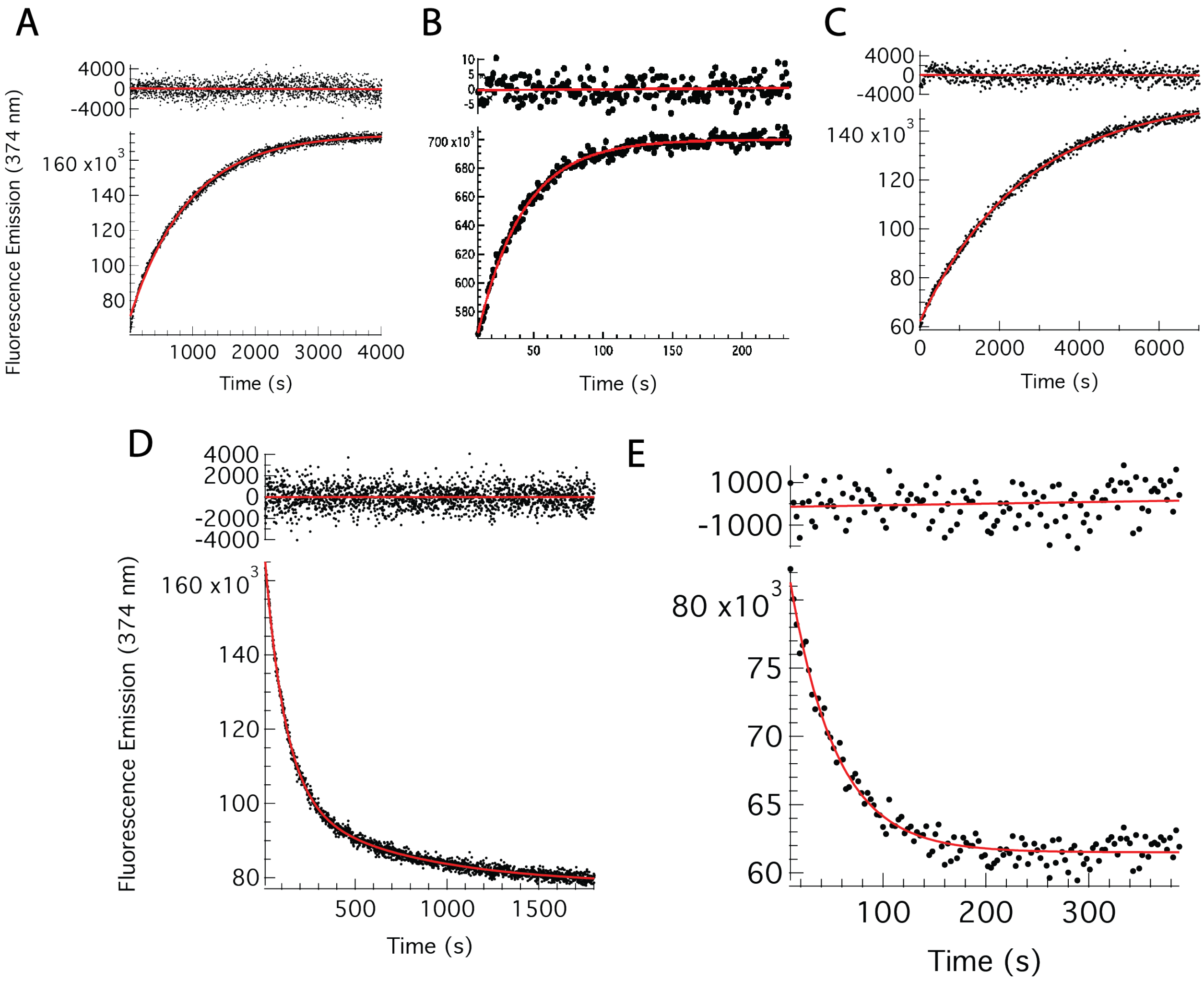


**Supplement 5.** Kinetic traces of A) N2 unfolding at 3.12 M GdmCl, B) N3 unfolding at 5.33 M GdmCl, C) N4 unfolding at 1.99 M GdmCl, D) N2 folding at 0.22M GdmCl, and E) N4 folding at 0.15M monitored by fluorescence emission at 374 nm, fit to single exponential curves.
